# Novel nitrifying symbiont lineages are vertically inherited and widespread in marine sponges

**DOI:** 10.1101/2023.12.23.573102

**Authors:** Bettina Glasl, Heidi M. Luter, Katarina Damjanovic, Katharina Kitzinger, Anna J. Mueller, Leonie Mahler, Joan Pamela Engelberts, Laura Rix, Jay T. Osvatic, Bela Hausmann, Joana Séneca, Holger Daims, Petra Pjevac, Michael Wagner

## Abstract

Ammonia-oxidising archaea and nitrite-oxidising bacteria are common members of marine sponge microbiomes. They derive energy for carbon fixation and growth from nitrification - the oxidation of ammonia to nitrite and further to nitrate - and are proposed to play essential roles in the carbon and nitrogen cycling of sponge holobionts. In this study, we characterise two novel nitrifying symbiont lineages, ‘*Ca*. Nitrosokoinonia’ and ‘*Ca*. Nitrosymbion’ in the marine sponge *Coscinoderma matthewsi* using a combination of molecular tools, single-cell imaging techniques, and physiological rate measurements. Both represent a new genus in the ammonia-oxidising archaeal class *Nitrososphaeria* and the nitrite-oxidising bacterial order *Nitrospirales*, respectively. Furthermore, we show that larvae of this viviparous sponge are densely colonised by representatives of ‘*Ca*. Nitrosokoinonia’ and ‘*Ca*. Nitrosymbion’ indicating vertical transmission. In adults, the representatives of both symbiont genera are located extracellularly in the mesohyl. Comparative metagenome analyses and physiological data suggest that ammonia-oxidising archaeal symbionts of the genus *Ca*. Nitrosokoinonia strongly rely on endogenously produced nitrogenous compounds (i.e., ammonium, urea, nitriles/cyanides, and creatinine) rather than on exogenous ammonium sources taken up by the sponge. Additionally, the nitrite-oxidising bacterial symbionts *Ca*. Nitrosymbion may reciprocally support the ammonia-oxidisers with ammonia via the utilisation of sponge-derived urea and cyanate. Interestingly, comparative analyses of published environmental 16S rRNA amplicon data revealed that ‘*Ca*. Nitrosokoinonia’ and ‘*Ca*. Nitrosymbion’ are widely distributed and predominantly associated with marine sponges and corals, suggesting a broad relevance of our findings.

## Introduction

Marine sponges (phylum *Porifera*) are among the oldest metazoans (1). They evolved around 650 million years ago (2). The sessile filter-feeding lifestyle and the ability to take up and recycle dissolved organic matter (DOM) make sponges important members of almost all marine benthic ecosystems, particularly in oligotrophic regions such as coral reefs (3–5). Their evolutionary and ecological success can partly be attributed to the diverse metabolic interactions with the abundant microbiome members they harbour in their tissue (6–8). Despite their ability to filter large amounts of seawater resulting in constant exposure to environmental microbes (9, 10), sponges discriminate between symbionts and food extremely well (11). Consequently, each sponge species is equipped with a distinct and relatively stable microbiome (12), which is mostly passed vertically from one generation to the next, but horizontal acquisition has also been reported (11, 13). In an adult sponge, the microbiome can contribute up to 35% of the sponge volume (14, 15) and facilitates key processes such as the removal of nitrogenous waste products, the production of vitamins and amino acids, and the uptake of DOM (6, 16–19). Overall, the symbiotic interactions of marine sponges with their microbiomes can be seen as one of the earliest examples of an animal holobiont, providing key insights into the origin of animal-microbe symbioses (20).

Nitrification, the oxidation of ammonia (NH_3_) to nitrite (NO_2_ ^-^) and further to nitrate (NO_3_ ^-^), is a key process in many marine sponge holobionts and is exclusively mediated by their microbial symbionts (6, 21, 22). Ammonium (NH_4_ ^+^) is a metabolic waste product of the sponge and has a higher toxicity to aquatic animals than nitrite and nitrate (23). The symbiont-driven transformation of ammonium is thus beneficial for the host. Additionally, the growth of nitrifiers might also represent an efficient recycling strategy to secure microbial food biomass for the sponge (17). Ammonia-oxidising archaea (AOA), class *Nitrososphaeria* (syn. *Thaumarchaea*), are key players in the nitrification process in marine sponges (24, 25). Ammonia-oxidising bacteria (AOB) can also be found in sponges, yet they play only a minor role compared to AOA (26, 27). The second step of nitrification in sponges is mediated by nitrite-oxidising bacteria (NOB), predominantly of the order *Nitrospirales* (6, 16, 28). Both ammonia- and nitrite-oxidisers are chemolithoautotrophs and use the energy gained from the oxidation of ammonia or nitrite, respectively, to fix dissolved inorganic carbon (DIC; 29, 30). Despite the frequent detection of putative nitrifying sponge symbionts using amplicon and metagenome sequencing (6, 16, 21, 22), studies investigating their contribution to the health and nutrition of the host, and their activity and localisation within the sponge tissue over different life stages remain scarce.

In this study, we characterised the environmental distribution, metabolic potential, metabolic function, and transmission mechanisms of nitrifying symbionts associated with the tropical marine sponge *Coscinoderma matthewsi,* using a combination of physiological rate measurements, molecular tools, and single-cell imaging techniques. The ammonia- and nitrite-oxidising symbionts of *C. matthewsi* belong to yet uncharacterised novel genera of the archaeal family *Nitrosopumilaceae* and the bacterial order *Nitrospirales*, respectively. Interestingly, members of both genera seem to be predominantly associated with marine sponges and corals, frequently co-occur within the same host species, and appear to be vertically transmitted. This study provides a comprehensive assessment of the symbiotic association and metabolic function of these novel AOA and NOB lineages ubiquitously associated with marine sponges.

## Materials and Methods

### Sponge collection

Three sponge individuals of the species *Coscinoderma matthewsi* were collected at 8 m depth from the reef surrounding Falcon Island (18° 45’ 59.3” S, 146° 32’ 7.8” E, Great Barrier Reef, Australia) in October 2021. Sponge species identification was confirmed by amplification and sequencing of two host phylogenetic marker genes (see Supplementary Methods and Supplementary Figure S1-S2). To ensure minimal impact on the natural sponge population, only parts of each sponge individual were removed, leaving the remaining sponge (at least one-third) on the reef. Sponges were collected under permit G21/38062.1 issued by the Great Barrier Reef Marine Park Authority. Removed sponge fragments were immediately relocated to a flow-through aquaria system at the National SeaSimulator at the Australian Institute of Marine Science (Townsville, Australia). After a one-month acclimation period, each sponge individual was fragmented into equally sized explants (approximately 5 cm in diameter) using sterile scalpel blades. Sponge explants were allowed to heal in an outdoor flow-through aquaria system (2,500 L) under natural lighting and ambient seawater temperatures (approximately 27°C) for two weeks before further experiments.

### Experimental setup

Nitrification activity of the *C. matthewsi* sponge holobiont was assessed during a six-hour incubation experiment. Sponge explants (each containing at least one visible osculum) were placed in 1.5 L acid-washed glass jars prefilled with 1 L of 0.1 µm filter-sterilised (Sawyer, hollow fibre membrane filter) seawater. Individual sponge explants were subjected to three different treatment conditions: i) ambient NH_4_ ^+^, ii) 5 µM NH_4_ ^+^, and iii) 25 µM NH_4_ ^+^. In the ‘ambient NH_4_ ^+^’ treatment, no additional ammonium source was added, and the sponge explants were exposed to ambient ammonium concentrations of in-shore reef sites (approximately 0.10 µM). In the 5 µM and 25 µM NH_4_ ^+^ treatments, the ammonium concentration was adjusted accordingly by addition of ammonium chloride (NH_4_Cl) from a sterile 25 mM stock solution before sponge explants were placed into the incubation jars. The elevated ammonium treatments reflect the approximate toxicity threshold for aquaculture species (100 µg L^-1^ ammonia) and the toxicant trigger value in marine water (900 µm L^-1^ ammonia) (31). In addition, abiotic control incubations (1 L of 0.1 µm filter-sterilised seawater without a sponge explant) were carried out in triplicates for each of the three treatment conditions. All incubation jars (n = 18) were randomly distributed among three chest-type orbital shaker incubators (Thermoline Scientific). Each incubator was set to 60 rpm and 27.5°C, matching the *in-situ* seawater temperature. All jars were kept in the dark throughout the experiment, as *C. matthewsi* is a non-phototrophic sponge. A HQ30D portable multimeter and an Intellical LDO101 oxygen probe (Hach, Colorado, USA) were used to measure oxygen concentrations at 0h and 6h. Oxygen levels at the end of the experiment (6h) dropped to 79% (± 5% SD) and 99% (± 1% SD) saturation in the sponge and abiotic control incubations, respectively.

Samples for determining nitrification activity were collected at four time points during the incubation (0h, 1h, 3h, and 6h). At each time point, 30 ml of seawater was sampled using a 50 ml syringe and subsequently filtered through a 0.45 µm filter (Sartorius Minisart, cellulose acetate membrane). Discarding the initial 10 ml of filtrate, the remaining 20 ml were divided into duplicates and stored at -20°C until further processing. Nitrite, nitrate, and ammonium concentrations in the samples were submitted to the AIMS analytical centre (Townsville, Australia) and analysed according to their standard operating procedures against OSIL standards and in-house reference samples using a Seal AA4 segmented flow analyser.

After the last sampling time point (6h), the wet weight (ww) of each sponge explant was measured, and the sponge tissue was then rinsed with 0.1 µm filter-sterilised seawater to remove loosely attached microbes. Subsequently, the tissue was subsampled for molecular analyses and microscopy. In addition, the explants of two out of three sponge individuals released larvae during the incubation experiments (approximately 2h into the experiment). Newly released larvae from the ‘ambient NH_4_ ^+^’ and ‘5 µM NH_4_ ^+^’ incubation jars were collected (approximately 2h after the release) using 1 mL plastic pipettes, pooled per individual, and processed for molecular analysis and microscopy.

### 16S rRNA gene amplicon sequencing and analysis

Subsampled sponge tissue incubated under ‘ambient NH_4_ ^+^’ conditions (n = 3) was snap-frozen in liquid nitrogen and larvae were preserved in molecular grade absolute ethanol. In addition, 1 L of freshly 0.1 µm filter-sterilised seawater (n = 2) was filtered onto a 0.2 µm Sterivex (Merck Pty Ltd) filter and snap-frozen in liquid nitrogen. All samples were stored at -80°C until the DNA was extracted using the DNeasy PowerSoil Pro Kit (Qiagen). A blank DNA extraction was included to account for possible contaminations. The V4 region of the 16S rRNA gene was amplified and sequenced at the Joint Microbiome Facility (JMF, Medical University of Vienna, Austria) under JMF project ID JMF-2203-05, using the primer pair 515F 5′-GTG YCA GCM GCC GCG GTA A-3′ (32) and 806R 5′-GGA CTA CNV GGG TWT CTA AT-3′ (33), as previously described by Pjevac *et al.* (34; see Supplementary Methods for further details). Amplicon sequence variants (ASVs) inferred from demultiplexed amplicon data in pooled mode with default settings with the DADA2 R package v1.20.0 (R 4.1.1) (35, 36) and classified with the SINA classifier v1.6.1 (37) against the SILVA reference database (SILVA release 138). ASVs classified as chloroplast and mitochondria were removed from the ASV table. Singletons (reads that occur only once) were removed from the ASV table and reads were rarefied to an equal sequencing depth of 4,424 sequencing reads for the subsequent data analysis.

Alpha diversity measures (i.e., richness, evenness, and Shannon Index) were calculated using the ‘phyloseq’ package (38). Data analysis and figure generation were performed in R (39) using the packages ‘dplyr’ (40) and ‘ggplot2’ (41), respectively.

### Metagenome sequencing, *de novo* assembly, and binning

DNA of one pooled larvae sample (larvae #2a) was submitted to the Joint Microbiome Facility (JMF, Medical University of Vienna, Austria) for metagenome sequencing, under JMF project ID JMF-2203-05 (see Supplementary Methods for further details). Metagenomic data were assembled using SPAdes v3.15.5 (42) in ‘meta’ mode with kmers set to 21, 31, 41, 51, 61, 71, 81, 91, 101, 111, 121. Contigs with less than 1,000 bp were removed from the assembly.

The assembly was binned into metagenome assembled genomes (MAGs) using MetaBAT2 v2.15, run with a minimum contig length of 1,500 bp and default workflow was used in concoct (43). Coverage information was calculated using BBMap v39.01 with 98% identity, and the resulting bam file was sorted using samtools v1.17. MAGs were also binned using MetaBAT2 with no coverage information. Both resulting MAG sets were dereplicated using dRep v3.4.2 (44). Completeness and contamination of the MAGs were calculated using dRep’s checkM output (45). Coverage for high-quality MAGs were calculated using BBMap at 95% mapping identity.

### Phylogenetic placement of metagenome-assembled genomes

The taxonomy of recovered high-quality (> 90% complete and < 5% contamination) MAGs (46) was assigned using GTDB-tk v2.3.2 (47). MAGs belonging to the archaeal family *Nitrosopumilaceae* (n = 1) and the bacterial order *Nitrospirales* (n = 1) were screened for 16S rRNA gene reads using Barrnap v0.9 and the recovered genes were aligned against the representative ASV sequences of the 16S rRNA gene amplicon dataset with the nucleotide blast tool (48). In case no 16S rRNA gene was detected in a MAG, the 16S rRNA gene amplicon reads of the dominant ASVs were placed into a full-length 16S rRNA gene tree of publicly available genomes with pplacer v1.1alpha19 (49). Phylogenomic trees of MAGs generated in this study and of publicly available genomes of closely related organisms (> 90% complete and < 5% contamination; Supplementary Table S1) were constructed using concatenated amino acid sequences of marker proteins extracted using checkM v1.1.3 (45). Phylogenetic trees were constructed in IQ-TREE v2.1.2 (50), see Supplementary Methods for details, and visualised in iTOL v5 (51).

### Global distribution and co-occurrence of novel nitrifying symbiont lineages

The global distribution and prevalence of the novel nitrifying archaeal and bacterial lineages were investigated by screening publicly available 16S rRNA gene amplicon sequencing datasets with the integrated microbial next-generation sequencing (IMNGS) platform (52). Full-length 16S rRNA gene sequences of the recovered MAGs (or their close relatives) were submitted as a ‘parallel similarity’ query against all available sequences in IMNGS (as of October 2023) with a similarity threshold of 97%, a minimum length of 200 nucleotides, and a minimal relative abundance of 0.01%. The similarity threshold was chosen based on the 16S rRNA gene sequence similarities among members of the novel nitrifying archaeal and bacterial lineages (Supplementary Table S1). IMNGS habitat classifications were manually corrected by cross-checking to original NCBI Sequence Read Archive entries. The following three categories were then assigned to the samples: ‘sponge’, ‘coral’, and ‘other’ (Supplementary Table S2). The data were analysed and visualised using the R packages ‘dplyr’ (40) and *‘*ggplot2’ (41).

### Metagenome assembled genome annotation

Nitrifier MAGs from this study and a dereplicated set of related, publicly available MAGs and isolate genomes (Supplementary Table S1) were functionally annotated using the eggnog mapper v2.1.10 (53) and the MicroScope platform v3.16.0 (54). Genome features (i.e. completeness, contamination, genome size, GC content, coding densities, and the number of predicted genes) were assessed using checkM v1.1.3 (45). The annotations were manually screened for genes of key metabolic features of AOA and NOB (i.e., CO_2_ fixation, ammonia and nitrite oxidation, electron transport, and alternative substrate utilisation). Unique and shared orthologs (based on ‘eggNOG OGs’) among MAGs/genomes were identified and visualised using the R packages ‘dplyr’ (40) and ‘Venn Diagram’ (55).

### Absolute quantification of nitrifying symbionts

Absolute abundances of the dominant AOA and NOB in the sponge tissue and larvae samples were quantified with droplet digital PCR (ddPCR) and symbiont specific primer sets using the Bio-Rad QX200 system (Bio-Rad Laboratory). Detailed description on the primer design, ddPCR optimisation, and ddPCR reaction is provided in the Supplementary Methods. The resulting concentrations (copies µl^-1^) were corrected for the DNA template dilution factor and normalised to the wet weight (copies g^-1^ ww) of the sponge tissue and larvae used for DNA extraction (Supplementary Methods).

### Visualisation of nitrifying symbionts using fluorescence *in situ* hybridisation (FISH)

Freshly collected sponge tissue and larvae were fixed in 4% paraformaldehyde (ProSciTech) overnight at 4°C, rinsed three times with ice-cold 1x phosphate-buffered saline (PBS), and subsequently stored in PBS:ethanol at -20°C. Thin sections of sponge tissue were prepared at the Histology Facility of the Vienna Biocenter (Vienna, Austria). In brief, fixed samples were washed for 1 h in 1x PBS and stored overnight at 4°C in 30% sucrose to preserve tissue morphology. Samples were subsequently transferred to 50% sucrose:Tissue-Tek O.C.T compound embedding medium (Sakura) for embedding, and stored overnight at 4°C. The embedded samples were transferred to isopentane and frozen in liquid nitrogen. Embedded tissue blocks were kept either on dry ice or at -80°C, until 5 µm-cryosections were prepared.

The 16S rRNA-targeted oligonucleotide probe Arch915 (56), tetra-labelled (57) with fluorochrome Atto565 (Biomers), was used for visualisation of AOA in tissue sections of *C. matthewsi*. Bacteria belonging to the genus *Nitrospira* were visualised using the 16S rRNA-targeted probe Ntspa662 (58) double-labelled (59) with Cy5 (Biomers). Hybridisations (4h at 46°C) were performed with an equimolar mixture of both probes and 30% formamide (v/v) in the hybridisation buffer. The stringency of the washing buffer was adjusted accordingly (60). Additional tissue sections were hybridised with probe non-EUB338-I (61) tetra-labelled with Atto565 or double-labelled with Cy5 and served as negative controls (Supplementary Figure S3). Sequences of FISH probes are provided in the Supplementary Methods. All hybridised tissue sections were DAPI stained and subsequently visualised at 1000x magnification on a THUNDER Leica Imaging System.

## Results

### Microbiome composition and vertical symbiont transmission in *C. matthewsi*

On average 171 (± 9 SD) unique ASVs were detected by 16S rRNA gene amplicon sequencing in the tissue of the sponge *C. matthewsi*. An equally high richness of unique ASVs was present in the freshly released larvae (169 ± 3 SD) and most ASVs were detected in the adult sponge as well as corresponding larvae (Supplementary Figure S4). The skewed evenness of the larval microbiome, however, led to a reduced alpha diversity (based on the Shannon Index) in the larvae (3.51 ± 0.27; mean ± SD) compared to the adult tissue (4.29 ± 0.18) samples (Figure 1A). Overall, 21 ± 1 distinct microbial phyla were detected in the adult sponge tissue and 20 ± 1 microbial phyla in the larvae (Figure 1B) samples. *Cyanobacteriota* was the only microbial phylum that was consistently absent in the larvae when compared to the adult sponge microbiome (Figure 1B and Supplementary Figure S4). Furthermore, the larval microbiome was dominated (39.7% ± 4.7%) by ASVs belonging to putative ammonia-oxidising archaeal symbionts of the phylum *Thermoproteota* (syn. *Crenarchaeota*), family *Nitrosopumilaceae* (Figure 1B-C). In contrast, the same ASVs only made up on average 5.0% ± 2.6% of the microbiome associated with the tissue of adult sponges.

**Figure 1.**
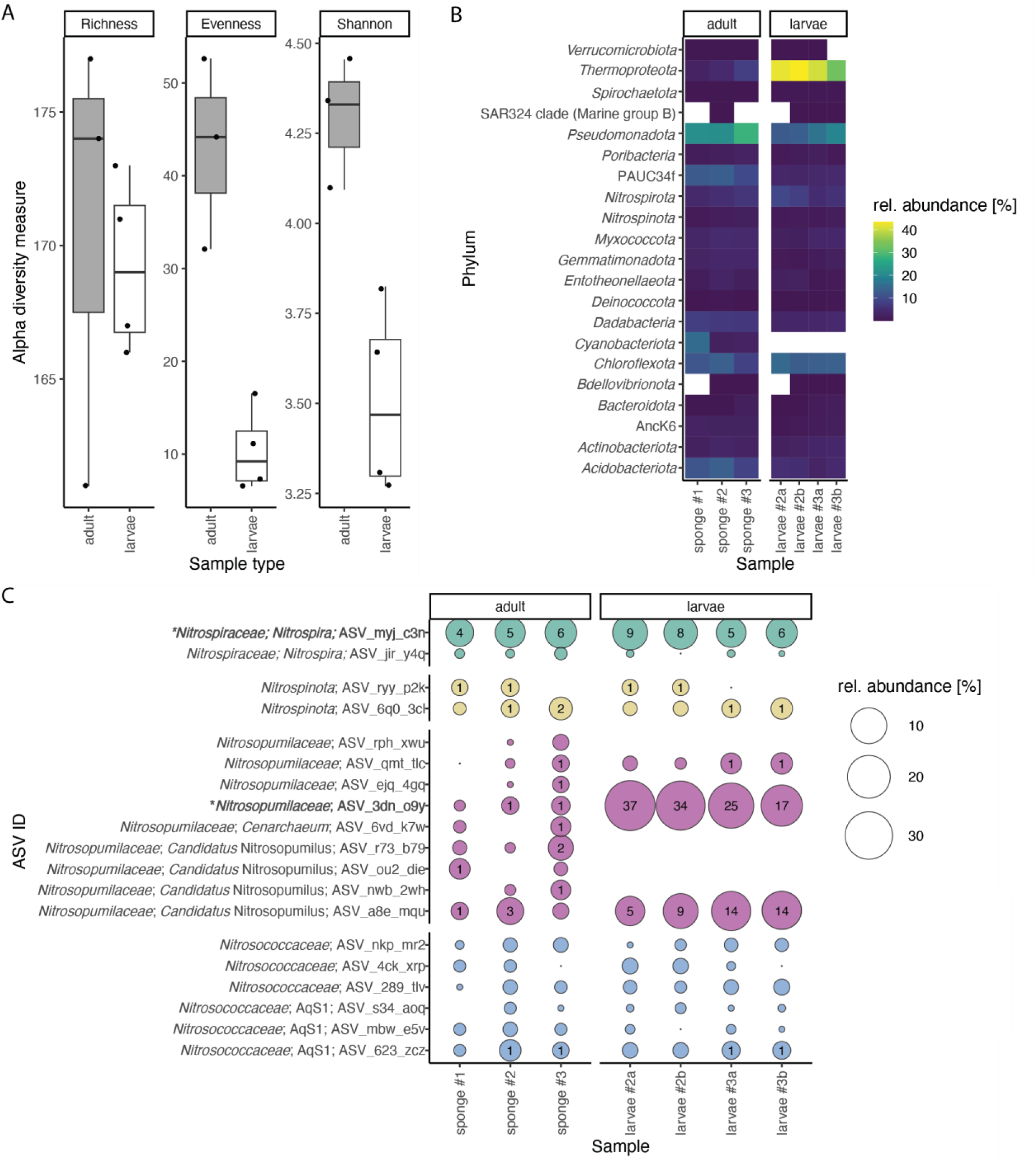
Alpha diversity and taxonomic composition of the *Coscinoderma matthewsi* microbiome. A) Microbiome richness, evenness, and Shannon diversity associated with adult and larvae samples. B) Bacterial and archaeal phyla (based on GTDB taxonomy) detected by 16S rRNA gene amplicon sequencing of adult sponge tissue (individual #1-#3) and pooled larvae samples (recovered from sponge individual #2 and #3). C) Relative abundance of individual amplicon sequence variants (ASVs) of putative nitrifying symbionts associated with adult sponge tissue and larvae samples. The asterisk (*) highlights ASVs for which representative high-quality metagenome-assembled genomes (MAGs) were obtained.

Cumulatively, putative nitrifying ASVs (function inferred based on their taxonomy; ‘*Nitroso*‘ = ammonia oxidisers, ‘*Nitro*‘ = nitrite oxidisers) made up on average 13.1% ± 4.7% and 49.1 ± 5.7% of the microbiome associated with adult sponges and larvae, respectively (Figure 1C). In general, the putative ammonia-oxidising community (‘*Nitroso’*) was dominated by two ASVs (ASV_3dn_o9y and ASV_a8e_mqu) belonging to the archaeal family *Nitrosopumilaceae*; both were present in all sequenced adult and larvae samples and were found in high relative abundances in all larvae samples. Absolute quantification with symbiont-specific ddPCR primers targeting the archaeal symbiont with the highest relative abundance (ASV_3dn_o9y) revealed an average absolute abundance of 1.02 ± 0.83 (SD) x 10^6^ *amoA* gene copies g^-1^ ww sponge tissue and 1.97 ± 1.54 (SD) x 10^9^ *amoA* gene copies per g^-1^ ww larva. The nitrite oxidising community (‘*Nitro’*) was dominated by one ASV (ASV_myj_c3n) of the bacterial order *Nitrospirales*, which also was present in all adult and larvae samples. Absolute quantification using symbiont-specific ddPCR primers targeting the 16S rRNA gene of the dominant *Nitrospira* symbiont revealed an average absolute abundance of 4.90 ± 4.00 (SD) x 10^7^ 16S rRNA gene copies g^-1^ ww sponge tissue and 1.43 ± 0.94 (SD) x 10^9^ 16S rRNA gene copies per g^-1^ ww larva. Furthermore, the AOA as well as nitrite-oxidising symbionts of the order *Nitrospirales* were visualised using FISH (Figure 2). Both symbiont clades were present in high abundance in the extracellular medium (mesohyl) of adult sponges, and in the central mesohyl of freshly released larvae.

**Figure 2.**
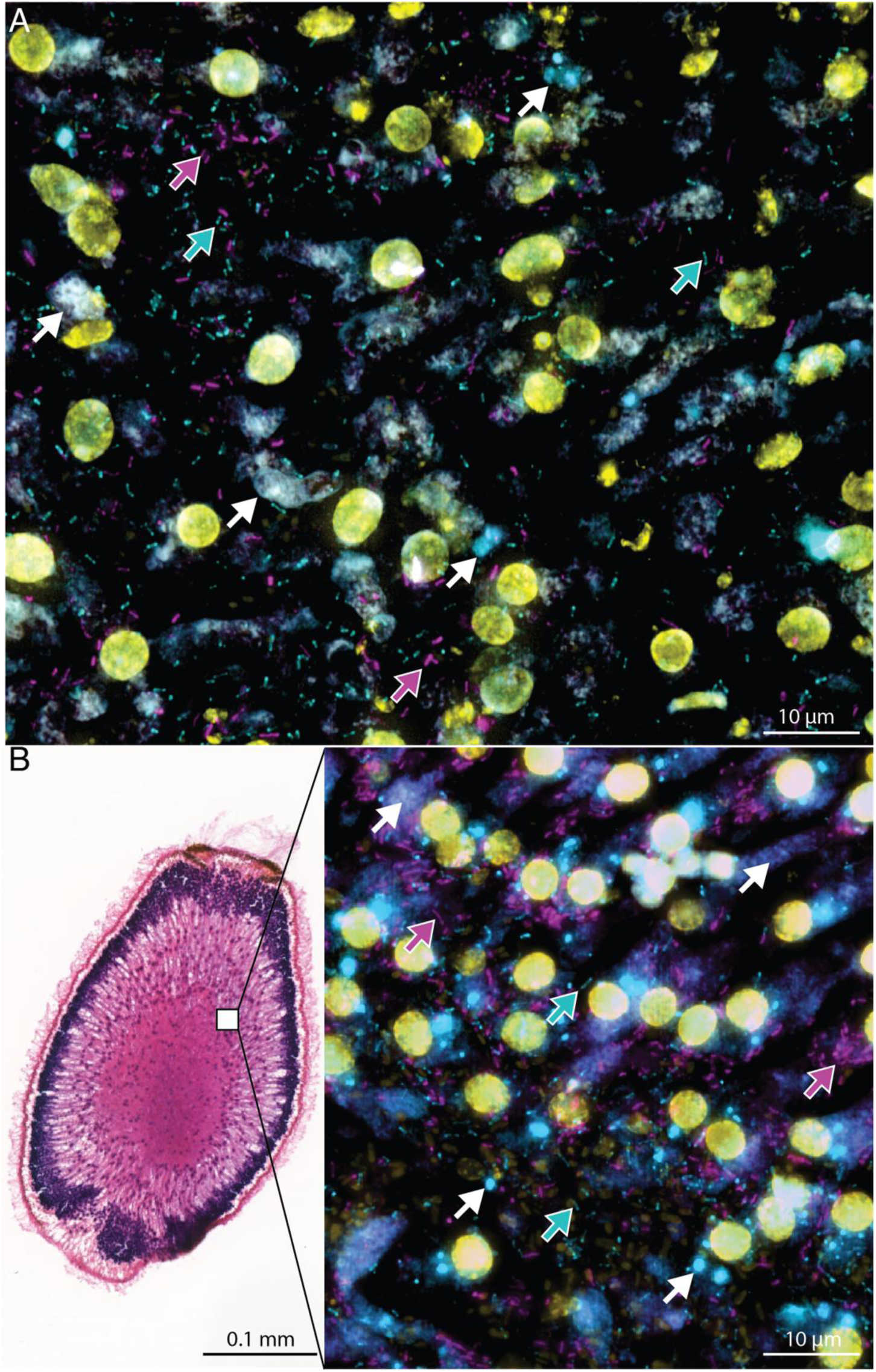
Visualisation of nitrifying symbionts in the sponge *Coscionderma matthewsi* using fluorescence *in situ* hybridisation (FISH). Symbionts were visualised in 5 µm cryosections of A) adult sponge tissue and B) freshly released larvae (longitudinal section of entire larva). Archaeal symbionts (based on 16S rRNA amplicon sequencing all belonging to the ammonia-oxidizing family of *Nitrosopumilaceae*) were hybridised using a tetra-labelled Arch 915 probe (Atto 565, shown in magenta, see magenta arrows). Putative nitrite-oxidising symbionts of the order *Nitrospirales* were hybridised using the double-labelled Ntspa 662 probe (Cy5, shown in cyan, see cyan arrows). Eukaryotic and prokaryotic DNA (if not additionally stained with a FISH probe) was stained using DAPI (shown in yellow). Purple/blue and white structures represent the autofluorescence of sponge tissue structures (see white arrows).

### Nitrification rates of the sponge holobiont

Nitrification rates of the *C. matthewsi* holobiont were experimentally determined to verify the ammonia- and nitrite-oxidising capacity of the associated microbiome. Nitrite accumulation was not observed in any of the sponge incubations (Supplementary Table S3). Nitrate concentrations increased significantly over time in all sponge incubations, suggesting ammonia was oxidised to nitrate without intermediate accumulation of nitrite (Figure 3A), irrespective of the initial ammonium concentration in the incubation jars (linear regression p-value = <0.05; Supplementary Table S3). In contrast, no nitrite and nitrate was produced and ammonium concentration remained stable over time in the abiotic control incubations under all three treatment conditions (Figure 3A; Supplementary Figure S5; Supplementary Table S3). Sponges kept under ‘ambient NH_4_ ^+^’ treatment conditions represented an ammonium source (Figure 3B). In the ‘5 µM NH_4_ ^+^’ treatment, the NH_4_ ^+^ production by the sponge holobiont met its consumption rates, however, in the ‘25 µM NH_4_ ^+^’ treatment, ammonium consumption exceeded the endogenous production rate (Figure 3B). Moreover, the nitrification rate was significantly higher in the ‘25 µM NH_4_ ^+^’ treatment (nitrate production rate = 0.1362 µM g^-1^ ww h ± 0.202 SE) when compared to the ‘5 µM NH_4_ ^+^’ treatment (nitrate production rate = 0.1083 µM per g^-1^ ww h ± 0.008 SE; t-test Bonferroni adjusted p-value = 0.0001), and ‘ambient NH_4_ ^+^’ (nitrate production rate = 0.096 µM g^-1^ ww h ± 0.008 SE; t-test Bonferroni adjusted p-value = 0.0001) condition.

**Figure 3.**
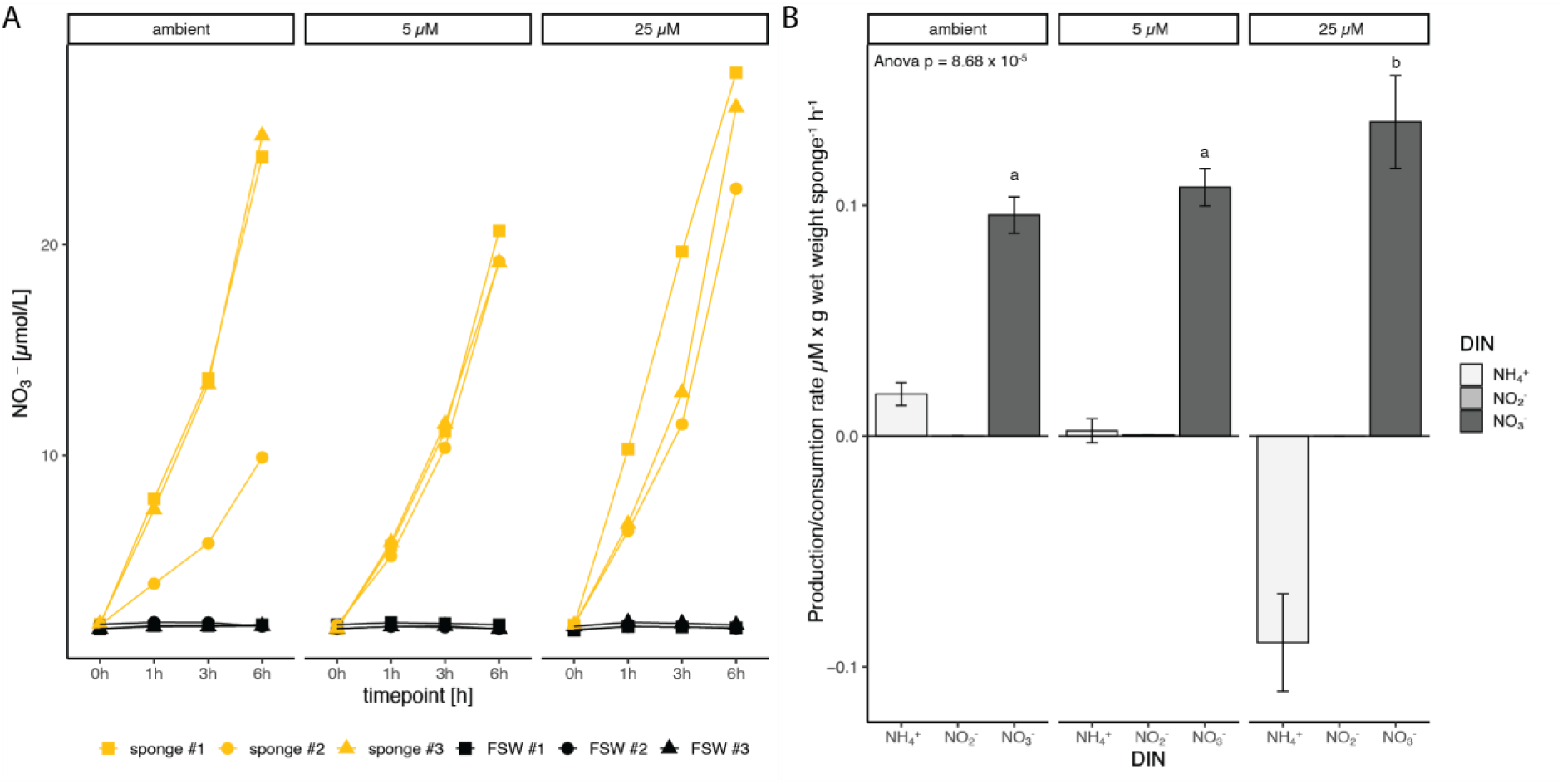
Nitrification activity of the *Coscinoderma matthewsi* holobiont. A) Production of nitrate (NO3^-^) per sponge individual (yellow) and in abiotic control incubations (black) in 0.1 µm filter-sterilized seawater (FSW) during the 6h incubation experiment under ambient and amended ammonium (NH4^+^) concentrations (5 µM and 25 µM, respectively). B) Dissolved inorganic nitrogen (DIN) production (> 0) and consumption (< 0) rates per gram wet weight (ww) sponge per hour. Rates and standard errors (represented as error bars) were calculated with a linear regression model across all time points. Ammonium (NH4^+^), nitrite (NO2^-^), and nitrate (NO3^-^) production/consumption rates are displayed for each of the three treatment conditions (i.e., ambient, 5 µM, and 25 µM NH4^+^).

### Phylogeny of dominant nitrifying symbionts

Two high-quality nitrifying symbiont MAGs (>90% completeness and <5% contamination) were recovered, belonging to the archaeal family *Nitrosopumilaceae* and the bacterial order *Nitrospirales*, respectively (Supplementary Table S1).

In a concatenated marker protein tree, the archaeal MAG was most closely related to four MAGs previously recovered from the marine sponges *Ircinia ramosa, Theonella swinhoei, and Petrosia ficiformis* (Figure 4A). The genome-wide average nucleotide identity (ANI) with these four MAGs ranged from 77% to 97% (Figure 4A and Supplementary Table S1), and based on the GTDB-tk analysis (Supplementary Table S4), these sponge-associated MAGs form the yet unnamed Genome Taxonomy Database (GTDB) genus ‘VYCS01’. Therefore, we propose to name this genus ‘*Candidatus* Nitrosokoinonia’ gen. nov. (see Taxonomic Consideration). Furthermore, based on a species-delineation threshold of 96.5% ANI (62), one MAG (GCA 009843815.1) previously recovered from the closely related sponge *Ircinia ramosa* represents the same species (Figure 4A). Here, we propose the name ‘*Candidatus* Nitrosokoinonia keratosae gen. nov. sp. nov. (see Taxonomic Consideration). Since no 16S rRNA gene was present in either *Ca.* N. keratosae MAG, the 16S rRNA gene amplicons of the two dominant archaeal ASVs (ASV_3dn_o9y and ASV_a8e_mqu) were placed into a full-length 16S rRNA gene tree of publicly available AOA genomes. The best placement location of ASV_3dn_o9y was with other members of the here-described genus *Ca*. Nitrosokoinonia, and for the ASV_a8e_mqu with the previously described sponge symbiont *Ca*. Nitrosospongia ianthellae (Supplementary Figure S6). This suggests that the dominant archaeal ASV_3dn_o9y can be linked to the MAG of *Ca*. Nitrosokoinonia keratosae.

**Figure 4.**
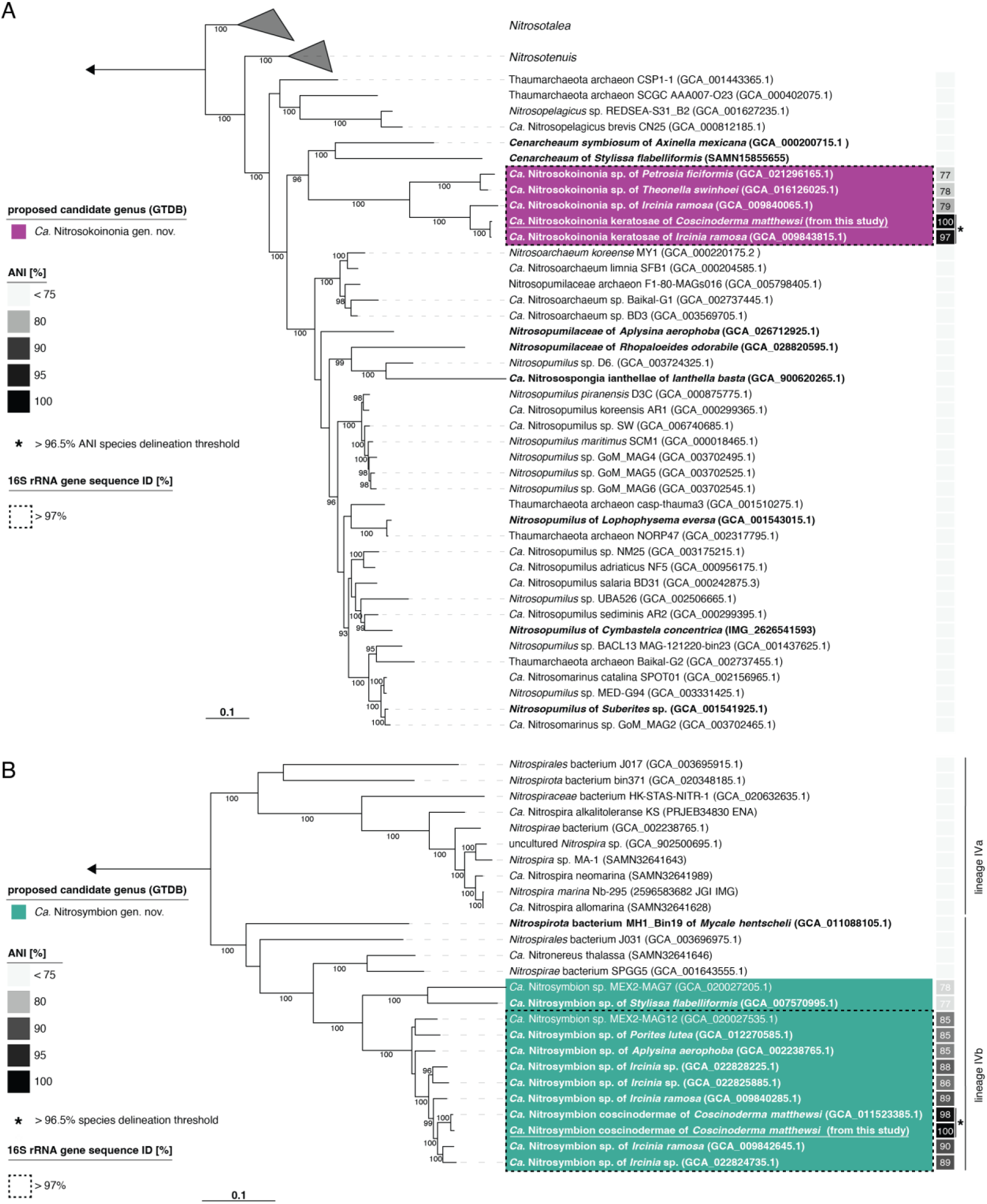
Phylogeny of nitrifying symbionts associated with the sponge *Coscinoderma matthewsi*. Maximum-likelihood tree of A) Ammonia-oxidising archaea (AOA) belonging to the family of *Nitrosopumilaceae* and B) the nitrite-oxidising bacteria (NOB) lineage IV *Nitrospirales*. Both phylogenomic trees are based on concatenated marker proteins identified with checkM v1.1.3 and depict the positions of the recovered metagenome-assembled genomes (MAGs) from the sponge *C. matthewsi*. Genomes and MAGs of previously described sponge symbionts are depicted in bold. The proposed candidate genera are highlighted in colour. The average nucleotide identities (ANI) of the MAGs from the dominant symbionts associated with the sponge *C. matthewsi* (recovered in this study) to other publicly available genomes and MAGs are displayed as a heatmap. MAGs highlighted with an asterisks (*) have an ANI > 96.5% (i.e. above the species delineation threshold) with MAGs recovered in this study. The dashed box highlights the boundaries of 16S rRNA gene sequence similarities > 97% (see Supplementary Table S1). Numbers at the branches indicate ultrafast bootstrap (n = 1,000) support. The scale bar corresponds to 0.1 estimated amino acid substitution per site.

The concatenated marker protein tree of lineage IV *Nitrospirales* showed a clear phylogenetic placement of the MAG described in this study among MAGs of the yet unnamed GTDB genus ‘bin75’ within the lineage IVb (Figure 4B). MAGs of this genus are predominantly associated with marine sponges (i.e., *Aplysina aerophoba*, *Coscinoderma matthewsi*, *Ircinia ramosa*, *Ircinia* sp., and *Stylissa flabelliformis*). Interestingly, one of the MAGs within this genus was previously retrieved from the coral *Porites lutea*. Only two of the 11 MAGs within this genus were not recovered from an animal host but rather originated from cold seep environments, which often harbour encrusting sponges and cold-water corals (63). However, given that most of the MAGs belonging to this genus were recovered from marine sponges, we propose to name this genus ‘*Candidatus* Nitrosymbion’ gen. nov. (see Taxonomic Consideration). Moreover, based on a species-delineation threshold of 96.5% ANI (62), the MAG retrieved in this study represents the same species as one MAG previously assembled from samples of the same sponge species (GCA 011523385.1, Figure 4B). Thus, we propose the name ‘*Candidatus* Nitrosymbion coscinodermae’ gen. nov. sp. nov. (see Taxonomic Consideration). Both *Ca*. N. coscinodermae MAGs contained the 16S rRNA gene, which showed a 100% nucleotide sequence identity with the dominant *Nitrospirales* ASV of the amplicon dataset (ASV_myj_c3n), implying that the *Ca*. N. coscinodermae MAG and the ASV_myj_c3n represent the same symbiont.

### Global occurrence of Ca. Nitrosokoinonia and Ca. Nitrosymbion

The occurrence of the archaeal candidate genus Nitrosokoinonia and the bacterial candidate genus Nitrosymbion in publicly available amplicon datasets was explored using IMNGS (52). In total, 1,070 samples (> 97% sequence similarity and > 0.01% relative abundance) were identified to contain either a member of *Ca*. Nitrosokoinonia (72 samples), or *Ca*. Nitrosymbion (881 samples), or both (117 samples). Both genera predominantly occur in samples (67% to 88%) of marine sponges and corals (Figure 5), yet, they were also detected in seawater and sediment samples which were often collected in close proximity to coral reefs (Supplementary Table S2). Overall, this analysis suggests that members of the genera *Ca*. Nitrosokoinonia and *Ca*. Nitrosymbion occur and also co-occur in marine sponges and corals around the globe.

**Figure 5.**
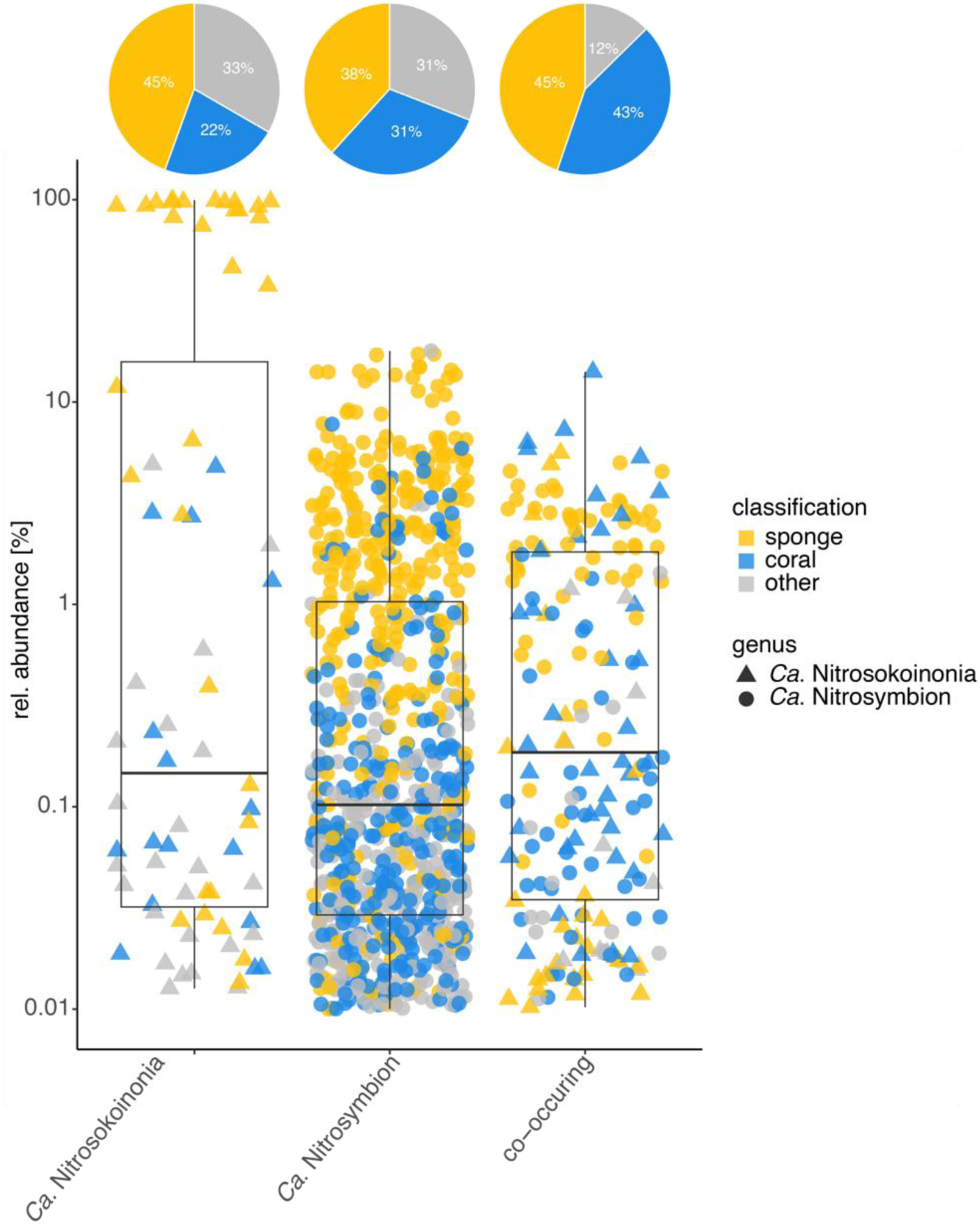
Environmental distribution of *Candidatus* Nitrosokoinonia and *Candidatus* Nitrosymbion. Relative abundances of the 16S rRNA genes grouped at 97% sequence identity level from publicly available amplicon sequencing datasets obtained from IMNGS. It is important to note that the numbers of the archaeal *Ca*. Nitrosokoinonia might be heavily under- or overestimated in publicly available amplicon datasets depending on the primer selection. Hence, differences in the primer selection might explain the detection discrepancy of the here-described bacterial and archaeal nitrifying genera in publicly available datasets. Each point represents an environmental sample where a closely related organism (> 97% sequence similarity) was detected, with a minimal relative abundance of 0.01%. Colours represent the environmental source of the individual samples and shapes represent the respective genus. The y-axis is log10 transformed. Pie charts show the relative frequencies (in percent) of samples from sponges, corals, and other environments.

### Genome characteristics of the novel candidate genera

MAGs of the archaeal genus *Ca*. Nitrosokoinonia are on average 1.61 Mbp (± 0.18 Mbp SD) in size and have a median GC-content of 51.90% (± 8.78% SD; Supplementary Table S1), which is smaller than the genome size of other sponge-associated AOA genomes (1.87 Mbp ± 0.30 Mbp SD). Generally, most described sponge-associated AOA have larger genomes (1.77 Mbp ± 0.28 Mbp SD) and a higher GC content (50.00% ± 12.79% SD) than free-living relatives (1.47 Mbp ± 0.25 Mbp SD and 35.40% ± 3.96% SD) of the family *Nitrosopumilaceae* (Supplementary Table S1). A pan-genome analysis including the new genus *Ca*. Nitrosokoinonia (n = 5), other sponge-associated AOA (n = 8), and closely related free-living AOA (n = 29; Supplementary Table S1) showed that a considerable number of unique orthologs are exclusively found in *Ca*. Nitrosokoinonia (34) and other sponge-associated AOA (135) genomes, or were shared amongst all symbiont genomes (29; Supplementary Figure S7A and Supplementary Table S5). For example, the unique orthologs include genes encoding for lipid A (endotoxin) biosynthesis, eukaryotic-like-proteins (Toll-interleukin-1 receptor (TIR)–like domain DUF1863), and restriction-modification systems (including a type II restriction endonuclease).

Genomes of the genus *Ca*. Nitrosymbion are on average 2.36 Mbp (± 0.29 Mbp SD) in size and have a GC-content of 56.90% (± 1.63% SD; Supplementary Table S1). In general, the average genome size of all lineage IVb *Nitrospirales* is smaller (2.44 Mbp ± 0.57 Mbp SD) when compared to genomes of the lineage IVa (3.84 Mbp ± 0.12 Mbp SD; Supplementary Table S1). However, within Lineage IVb, there is a large range in genome sizes. Some of the *Ca*. Nitrosymbion MAGs have the smallest genomes within this lineage, while the only cultivated representative, *Ca.* Nitronereus thalassa, has an unusually large genome (4.02 Mbp; Supplementary Table S1). A pan-genome analysis including the new genus *Ca*. Nitrosymbion (n = 12), other members of lineage IVb (including *Ca*. N. thalassa; n = 4), and members of lineage IVa (n = 9; Supplementary Table S1) showed that 385 identified orthologs are exclusively present in MAGs of the genus *Ca*. Nitrosymbion (Supplementary Figure S7B and Supplementary Table S6). For example, the unique orthologs include genes that encode for multiple toxin-antitoxin modules, bacteriocin protection mechanism, and a solute/sodium symporter.

### Genomic potential of ‘*Candidatus* Nitrosokoinonia keratosae’ gen. nov. sp. nov

The representative MAG of *Ca*. Nitrosokoinonia keratosae (97.57% completeness and 0.00% contamination) contains all the hallmark genes for chemolithoautotrophic ammonia oxidation and CO_2_ fixation (Figure 6A and Supplementary Table S7). This includes genes encoding for all known and putative subunits of the ammonia monooxygenase gene cluster (*amo*AXBCYZ), the 3-hyroxypropionate/4-hydroxybutyrate (3HP/4HB) cycle for CO_2_ fixation, and the electron transport chain (complex I to IV). Like all other AOA genomes of the family *Nitrosopumilaceae*, except for *C. symbiosum*, *Ca*. N. keratosae also encodes for the NO-forming Cu-containing nitrite reductase (*nirK*), which has been hypothesised to play a critical role in the currently postulated archaeal ammonia oxidation pathways (64, 65). Furthermore, *Ca*. N. keratosae encodes for the candidate enzyme ‘*Cu-Hao’*, which has been suggested to be putatively involved in hydroxylamine oxidation (66, 67). The *Ca*. N. keratosae genome also encodes for a high-affinity ammonia transporter (*amt*2) (68) and, like almost all other sponge-associated AOA genomes, lacks the low-affinity ammonia transporter (*amt1*) and only contains one nitrogen regulatory protein PII gene. In the genomes of their free-living relatives, *amt1* transporters and additional copies of the nitrogen regulatory protein PII genes can be found. Interestingly, *Ca*. N. keratosae might be able to directly use sponge-derived urea, nitriles/cyanides, and creatinine but not cyanate as alternative ammonia sources. Its genome contains a urease gene cluster (*ureABC*), a gene encoding for nitrilase/cyanide hydratase (*nitA*), and for a creatinase/creatinine amidohydrolase as well as for creatinase/creatin amidohydrolase (Figure 6A). Additionally, the genome of *Ca*. N. keratosae encodes for branched-chain amino acid transporters (*liv*FGM) - a genomic feature unique to sponge-associated AOA (17).

**Figure 6.**
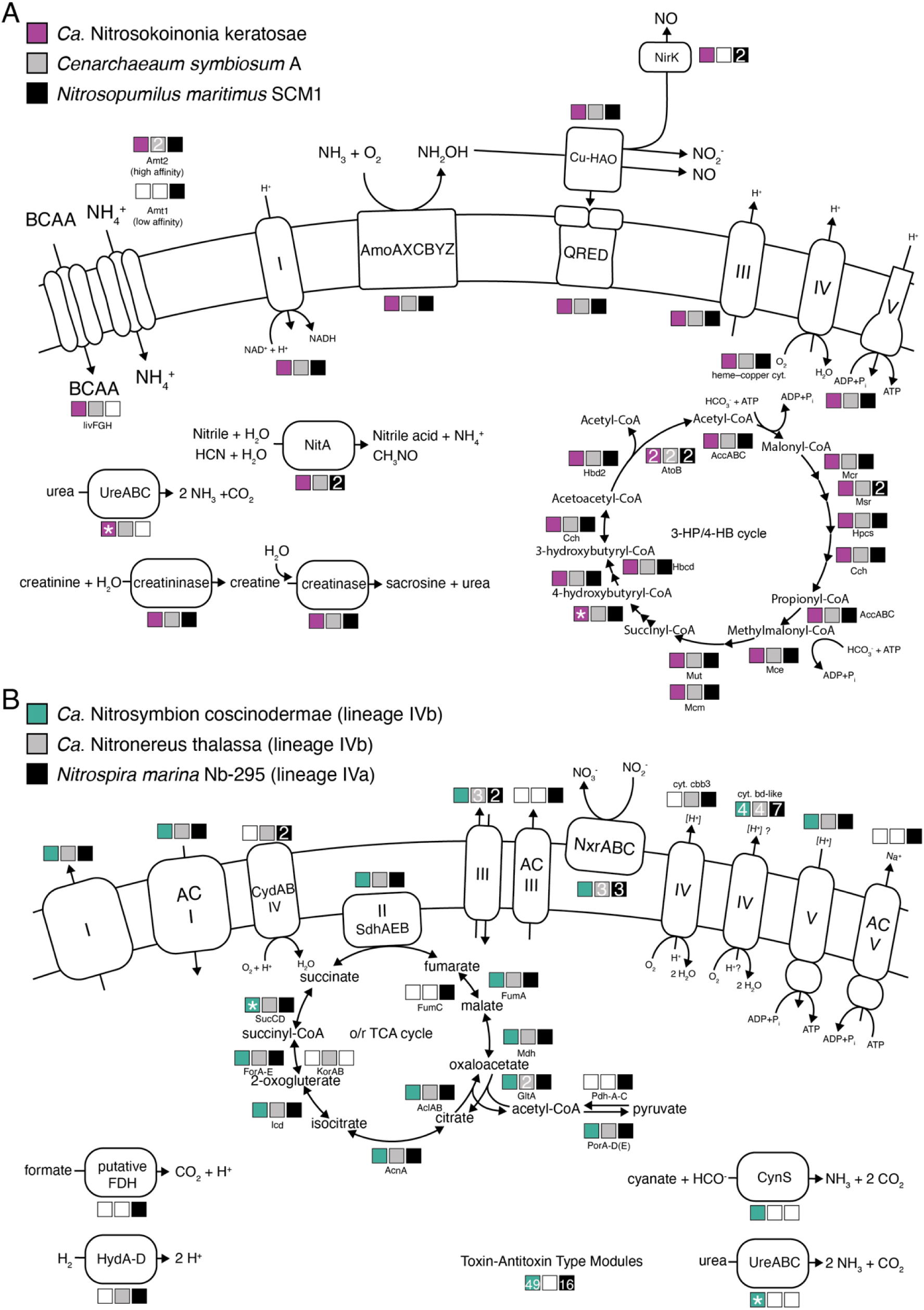
Schematic overview of selected metabolic key pathways. A) Genomic repertoire of the ammonia-oxidising archaeal sponge symbionts *Ca*. Nitrosokoinonia keratosae and *Cenarchaeum symbiosum* A as well as the marine *Nitrosopumilus maritimus* SCM1. The illustration depicts key steps of the putative ammonia-oxidation pathway i.e., the ammonia-monooxygenase enzyme (AmoAXCBYZ), the putative copper-containing hydroxylamine oxidase (Cu-Hao), the putative quinone reductase (QRED) and the nitrite reductase (NirK). It further highlights the electron transport chain (complex I-V) and the 3-hyroxypropionate/4-hydroxybutyrate (3-HP/4-HB) cycle mediating CO2 fixation. In addition, selected key functions such as ammonia transporters (Amt1 and Amt2), branched-chain amino acid (BCAA) transporters, as well as pathways facilitating the utilisation of alternative substrates such as urea (UreABC), nitrile/cyanide (NitA), and creatinine (creatininase and creatinase). B) Genomic repertoire of the nitrite-oxidising bacterial sponge symbiont *Ca*. Nitrosymbion coscinodermae, as well as the marine *Ca*. Nitronereus thalassa and *Nitrospira marina* Nb-295. The illustration depicts hallmark genes of nitrite-oxidising bacteria such as the nitrite oxidoreductase (NxrABC), the membrane-bound respiratory chain (complex I-V) including the alternative complexes (AC I-V), and the oxidative and reductive tricarboxylic acid (o/r TCA) cycles. In addition, selected key functions such as toxin-antitoxin (TA) modules, formate metabolisation (putative FDH), putative hydrogen oxidation (HydA-D; Group 3b [NiFe] hydrogenase), and cyanate (CynS) and urea (UreABC) utilisation. Coloured boxes indicate the presence of genes coding for the shown proteins in the respective genomes. If more than one copy of a gene was present, the copy number of this gene is displayed in the box. Asterisk (*) indicates that genes were missing in the metagenome assembled genomes (MAG) of the species representative but were present in the other MAG(s) of the newly described genus.

### Genomic potential of ‘*Candidatus* Nitrosymbion coscinodermae’ gen. nov. sp. Nov

The genome of the nitrite-oxidising sponge symbiont *Ca*. Nitrosymbion coscinodermae (completeness 98.64% and contamination 3.86%) encodes for all hallmark genes for chemolithoautotrophic nitrite oxidation and CO_2_ fixation. This includes genes for the known subunits of nitrite oxidoreductase (*nxr*ABC), the oxidative (oTCA) and reductive (rTCA) tricarboxylic acid cycles, and the membrane-bound respiratory chain (Figure 6B and Supplementary Table S8). Similar to other genomes of the order *Nitrospirales*, *Ca*. N. coscinodermae encodes for four putative cytochrome *bd*-like oxidases (complex IV); however, it lacks the high-affinity, heme-copper cytochrome *cbb_3_*-type terminal oxidases found in some other lineage IV *Nitrospirales* (Figure 6B and Supplementary Table S8). Moreover, the genome of *Ca*. N. coscinodermae also lacks the putative alternative complexes (AC) III and V (Figure 6B and Supplementary Table S8).

The order *Nitrospirales* likely emerged from anaerobic ancestors (69). Based on the observed genome adaptations we propose that *Ca*. N. coscinodermae, like all other members of the order Nitrospirales, has adapted to life under oxygenated conditions. For example, the *Ca*. N. coscinodermae genome encodes for the presumably oxygen-tolerant form of pyruvate:ferredoxin oxidoreductase (*porA-D*) (69) and the five subunits of oxoglutarate:ferredoxin oxidoreductase (*forA-E*), key enzymes of the o/rTCA cycle that are universally present in genomes of the order *Nitrospirales*. In contrast to the closely related *Ca*. Nitronereus thalassa (70), the *Ca*. N. coscinodermae genome lacks an isoenzyme Kor (*korAB*), which was previously described to facilitate growth under anaerobic conditions (71).

Alternative electron donors, such as hydrogen and formate, have been described to be used by some *Nitrospirales* (69, 72–74). In the genome of *Ca*. N. coscinodermae, putative formate dehydrogenase genes (putative FDH) or genes related to hydrogen metabolism (3b [NiFe] hydrogenase, 2a [NiFe] hydrogenase, and related genes involved in hydrogenase maturation) were not detected (Figure 6B and Supplementary Table S8). Thus, alternative metabolisms based on formate and hydrogen seem unlikely. Moreover, the *Ca*. N. coscinodermae genome also lacks the canonical cyt. *bd* quinol oxidase (*cydAB*, complex IV), which accepts electrons from electron donors with lower reduction potential than nitrite and may play a role in the oxidative stress response in other *Nitrospirales* (69, 72–74).

Based on its genomic repertoire, *Ca*. N. coscinodermae may be able to take up small dissolved organic nitrogen compounds, such as cyanate and urea, for nitrogen assimilation. The genome of *Ca*. N. coscinodermae encodes for the enzymes cyanase (*cynS*) and urease (*ureABC*), both of which can cleave ammonia from the original substrate and release CO_2_. Thus, *Ca*. N. coscinodermae potentially indirectly gains energy from cyanate and urea via reciprocal feeding (73, 75) with the AOA symbiont population within the sponge holobiont. Interestingly, the genome of *Ca*. N. coscinodermae as well as other members of the genus *Ca*. Nitrosymbion are enriched in genes encoding for toxin-antitoxin (TA) modules, especially of type II and type IV, which may reflect an adaptation to a life in close proximity to other microbes, a defence mechanism against phages and/or play a role in evading host phagocytosis (76).

## Discussion

Marine sponge microbiomes can be remarkably diverse (12, 77). Yet, nitrification appears to be a ubiquitous functional trait in marine sponge microbiomes (6, 21, 22, 78). In this study, we characterise two novel genera of nitrifiers, *Ca*. Nitrosokoinonia and *Ca*. Nitrosymbion, which frequently co-occur in the microbiome of many marine sponges, and intriguingly, corals (Figure 5 and Supplementary Table S2). The prevalent symbiotic lifestyle and co-occurrence of *Ca*. Nitrosokoinonia and *Ca*. Nitrosymbion may indicate a co-evolutionary relationship between the two symbiont lineages and their shared habitat preferences. Notably, sponges and corals are among the most basal metazoan phyla (1), and thus the diversification of these two symbiont lineages may be intertwined with the early evolution of animals.

Vertical symbiont transmission is common in marine sponges (11, 13) and has also been observed for nitrifying symbionts (79). Viviparous sponge species (‘brooders’) predominantly transfer their symbionts vertically from one generation to the next (11). *C. matthewsi* is a viviparous sponge (80), and we detected the symbionts *Ca*. N. keratosae and *Ca*. N. coscinodermae in high abundance in the mesohyl of freshly released larvae (Figure 1 and Figure 2). Interestingly, MAGs with the highest ANI (> 88% ANI) to the representative symbionts of *C. matthewsi* were associated with the closely related viviparous sponge *Ircinia ramosa* (Figure 4 and Supplementary Table S1). Symbionts belonging to the novel genera are not restricted to viviparous sponge species of the order Dictyoceratida but can also be detected in tissue samples of oviparous sponge such as *Aplysina aerophoba* (Figure 4 and Supplementary Table S3). Larvae of oviparous sponges have been described to already contain symbionts during their development outside of the mother sponge (79, 81). Since currently known sponge larvae are lecithotrophic, i.e., they do not feed during their short planktonic life stage (82), the symbionts of the herein described genera are likely vertically transmitted in both viviparous and oviparous sponges. However, the precise transmission strategy of both symbiont lineages remains to be determined.

The ammonia-oxidising archaeal symbiont *Ca*. N. keratosae dominated the larval microbiome (Figure 1). The overrepresentation of AOA in the larval microbiome may suggest that they play an essential role in the early life stages of a sponge. For example, vertically inherited AOA of the genus *Ca*. Nitrosokoinonia may induce early metamorphosis of their sponge host via the production of nitric oxide (NO). Nitric oxide is a well-known signalling molecule in marine invertebrates (83). For instance, in the sponge *Amphimedon queenslandica,* nitric oxide has been shown to trigger larval settlement and metamorphosis (84). Nitric oxide production by AOA (Figure 6A) in sponge larvae could potentially induce settlement and therefore either help to sense a suitable substrate (i.e. ammonia and oxygen availability) or function as an indicator of nitrogenous waste accumulation. Elucidating the function of AOA symbionts in sponge larvae and their ability to interfere with the eukaryotic signalling pathway of their host is an interesting avenue for further research.

Nitrification activity was measured in the adult sponge holobiont. Linking physiological data (Figure 3) with the genomic potential of *Ca*. N. keratosae and *Ca*. N. coscinodermae (Figure 6) indicates that the dominant nitrifying symbionts actively contribute to the removal of endogenously produced ammonia from the sponge holobiont. However, based on 16S rRNA gene amplicon sequencing (Figure 1), *Ca*. N. keratosae and *Ca*. N. coscinodermae are not the only nitrifying symbionts residing within the sponge holobiont. Addressing microdiversity and niche differentiation in terms of spatial distribution and substrate utilisation will be crucial to understanding co-occurrence of functionally redundant symbionts within a single sponge host. It also remains to be determined how symbiont shuffling and/or switching of functionally redundant symbionts can affect the phenotypic plasticity of the holobiont and, thus, facilitate adaptation.

Nitrogenous waste products of the sponge holobiont fuel the symbiont-mediated nitrification activity. Nitrification rates of the *C. matthewsi* microbiome seem to rely on endogenous ammonia sources and are not drastically affected by external ammonia concentrations (Figure 3). Similar observations have recently been described for the Mediterranean sponge *Chondrosia reniformis*, where exogenous ammonia concentrations did not significantly affect the ammonia and oxygen consumption nor the exhaled nitrate concentrations (85). Ammonia assimilation in AOA is regulated by nitrogen regulatory proteins PII, which are overrepresented in free-living AOA genomes (86). Nitrogen regulatory proteins receive information about the carbon/nitrogen ratio and energy status of the cell and regulate ammonia uptake and utilisation in response to changes in extracellular nitrogen availability (87). Since sponge-associated AOA are universally equipped with the high-affinity ammonia transporter (*amt2*) and reduced copy numbers of nitrogen regulatory protein PII genes (Supplementary Table S7), the symbiotic AOA seem to be adapted to low but stable ammonium concentrations. Besides ammonium, alternative nitrogenous substrates such as urea, cyanide, creatinine, and branched-chain amino acids may play a crucial role in fuelling the ammonia oxidation rate of *Ca*. Nitrosokoinonia, as they are ubiquitously equipped with genes encoding for urease (*ureABC*), nitrilase/cyanide hydratase (*nitA*), creatinase (*arfB*), and branched-chain amino acid transporters (Figure 6A). Interestingly, nitrogenous terpenoids, which feature nitrogen-containing functional groups like isonitriles, isothiocyanates, and formamides that originate from inorganic cyanide, are a secondary metabolite class ubiquitously produced by marine sponges (88). Thus, sponge-produced nitrogenous terpene (88), and other nitrogen-containing secondary metabolites (89), as well as creatin/creatinine (28) might function as additional ammonia sources. Moreover, *Ca*. Nitrosymbion genomes also encode genes for enzymes involved in utilising simple organic nitrogen compounds - urease (*ureABC*) and cyanase (*cynS;* Figure 6B). This suggests nitrite-oxidising symbionts of the here-described genus *Ca*. Nitrosymbion can convert urea and cyanate to ammonia and CO_2_, providing an additional energy source to the resident AOA symbiont population within the holobiont, and receiving nitrite in return. Reciprocal feeding would be particularly relevant if the co-occurring AOA population lacks an urease (73, 75), which is currently uncertain (Figure 6B).

Overall, this study characterises two new genera of AOA and NOB that frequently occur and co-occur in marine sponges and corals. The predominant association of these genera with basal metazoan hosts and the vertical transmission mode suggest that both genera are well adapted to their symbiotic lifestyle and may even have co-evolved alongside their hosts. Furthermore, the high number of unique orthologs suggest an immense source of genetic novelty. We therefore propose that both genera, *Ca*. Nitrosokoinonia and *Ca*. Nitrosymbion, are an ideal model system to study co-evolution and co-diversification of symbionts associated with marine sponges and corals. Marine sponge holobionts are currently emerging as an interesting model system due to their complex microbe-microbe and host-microbe interactions. Here, we particularly emphasise the importance of studying niche partitioning within the sponge holobiont, single-cell interactions among holobiont members, and the effect of symbiont-produced metabolites on the host signalling pathways.

### Taxonomic consideration of ‘*Candidatus* Nitrosokoinonia keratosae’ gen. nov. sp. nov

niˈtro.so.koi.noˈni.a L. n. nitroso: nitrosus, full of natron, here intended to mean nitrous; Gr. fem. n. koinonia: refers to concepts such as fellowship, joint participation, partnership; L. fem. n. Nitrosokoinonia partner that consumes ammonia. N.L. n. keratosa: subclass of Demospongiae comprising horny sponges with a spongin skeleton and without spicules.

An ammonia-oxidising, nitrite-producing archaeon associated with keratose sponges. Phylogenetically affiliated with the order *Nitrosopumilales*, phylum *Thermoproteota (syn. Thaumarchaeota)*. The metagenome-assembled genome (MAG) is 97.57% complete and has 0.00% contamination. It consists of 162 scaffolds, with a total of 1,505,883 bp. The DNA G+C content is 45.56%. *Ca*. Nitrosokoinonia keratosae was recovered from the tissue of two closely related marine sponge species (*Ircinia ramosa* and *Coscinoderma matthewsi)*, collected at Great Barrier Reef in Queensland (Australia).

### Taxonomic consideration of ‘*Candidatus* Nitrosymbion coscinodermae’ gen. nov. sp. nov

Ni.tro.sym·bi·on L. n. nitrum: nitrate, Gr. n. symbion: living together; L. fem. n *Nitrosymbion* nitrate-forming symbiont. cos.cino.dér.mae Gr. fem. n. koskinon: sieve, Gr. fem. n dermae: skin; Gr. fem. n. a sieve-like skin, in this case referring to the sponge genus *Coscinoderma;*

A nitrite-oxidising, nitrate-forming symbiont obtained from a marine sponge. Phylogenetically affiliated with the order *Nitrospirales*, phylum *Nitrospirota*. The metagenome-assembled genome (MAG) is 98.64% complete and has 3.86% contamination. It consists of 38 scaffolds, with a total of 2,506,777 bp. The DNA G+C content is 56.70%. *Ca*. Nitrosymbion coscinodermae was recovered from the tissue of the marine sponge *Coscinoderma matthewsi*, collected at the in-shore coral reef location Falcon Island (Great Barrier Reef, Queensland, Australia).

## Supporting information

Supplementary Methods and FiguresS1-7

Supplementary TablesS1-8

## Acknowledgments

This research was funded by the Austrian Science Fund (FWF) [T 1218] Hertha-Firnberg Fellowship awarded to BG and the Wittgenstein Award of the FWF (Z-383-B to MW) For the purpose of open access, the BG has applied a CC BY public copyright license to any Author Accepted Manuscript version arising from this submission. We thank Muhammad Azmi Abdul Wahab for his help with the sponge species identification and field collections. We also thank the AIMS Analytical Technology Team for their help with analysing the dissolved inorganic nitrogen concentrations and the CeMESS Technical Assistant Team, Microscopy Team, JMF Laboratory Team as well as the Life Science Compute Cluster for their support throughout the project. We further acknowledge the traditional owners of land and sea country, the Bindal, Manbarra and Wulgurukaba people, where parts of the research has been conducted and we respect their Elders past, present and emerging.

## Data availability statement

The assembled genomes, raw sequencing reads, and amplicon sequencing (16S rRNA gene, 28S rRNA gene and CO1 gene) data are available upon request under the Join Microbiome Facility project ID JMF-2203-05 and JMF-2305-01, and will be available at the NCBI BioProject PRJNA1037309. Furthermore, all assembly IDs of publicly available genomes and MAGs used within this study are summarised in Supplementary Table S1. Nitrification data, including NH_4_^+^, NO_2_^-^, and NO_3_^-^ concentrations, are available in the Supplementary Table S3.

## Authors’ contributions

BG prepared the samples for sequencing, analysed the data, prepared the figures, and drafted the first version of the manuscript. HML and KD collected samples and conducted the experiments. KK, AM, PP, HD, MW, and BG were involved in the conceptual design of the study and the interpretation of the data. JO performed the metagenome read assembly and binning. BH and JS performed the amplicon data processing and analysis. BG and LM optimised the ddPCR protocol for the absolute quantification of the symbionts. BG, LM, KK, JPE, and LR optimised the FISH protocol to visualise nitrifying symbionts in the sponge tissue. All authors read and approved the final manuscript.

