## Supplementary Methods and FiguresS1-7 for "Novel nitrifying symbiont lineages are vertically inherited and widespread in marine sponges"

<u>Content:</u>

Supplementary Methods

7 Supplementary Figures

8 Supplementary Tables

**Supplementary Methods**

*Sponge phylogeny*

During sampling, the sponge species was identified based on its visual appearance. Species identification was later confirmed by phylogenetic marker gene sequencing. In brief, subsampled tissue of all three sponge explants incubated under 'ambient $NH_4^{+}$' conditions was snap frozen in liquid nitrogen and stored at -80°C until DNA was extracted using the DNeasy Blood & Tissue Kit (Qiagen). DNA extracts (n = 3) were sent to the Joint Microbiome Facility (JMF, Medical University of Vienna, Austria; project ID JMF-2305-01) for amplicon sequencing of two commonly used phylogenetic marker genes (i.e. 28S rRNA and CO1) following their standard procedure (1). The C-region of the 28S rRNA gene was targeted using the C2-forward: 5′-GAA AAG AAC TTT GRA RAG AGA GT-3′ and D2-reverse: 5′-TCC GTG TTT CAA GAC GGG-3′ primers (2). The 5' end of the mitochondrial cytochrome oxidase subunit 1 (CO1) gene was amplified with the forward primer dgLCO1490: 5′-GGT CAA CAA ATC ATA AAG AYA TYG-3′ and the reverse primer dgHCO2198: 5′-TAA ACT TCA GGG TGA CCA AAR AAY CA-3′ (3). Amplicon pool extraction from raw sequencing data, PhiX contamination filtering and demultiplexing were performed similarly to the 16S rRNA gene dataset. Extracted 28Sr rRNA and CO1 FASTQ reads were trimmed (230 nt, for the forward and reverse read, allowing 4 and 6 errors per read, respectively. The derived 28S rRNA gene amplicon sequence variants (ASVs) were blasted against the NCBI nucleotide collection with an optional organism filter set to 'Porifera (taxid:6040)'. The taxonomy of the CO1-ASVs was assigned using the BOLD database (4) and only animal-derived ASVs were kept in the dataset. ASVs with the highest relative abundances were subsequently aligned with mafft (5) against the 28S rRNA gene and CO1 gene sequences of sponges belonging to the order Dictyoceratida (6). Phylogenetic trees (see Supplementary Figure S1 and S2) were calculated using IQ-Tree (7) and visualised in iTOL (8).

*16S rRNA gene amplicon sequencing*

Sequencing libraries of barcoded amplicon pools were prepared using the TruSeq Nano DNA Kit (Illumina), and sequenced on the Illumina MiSeq platform (V3 chemistry, 600 cycles). Downstream data processing included amplicon pool extraction from raw sequencing data (FASTQ workflow, default parameters; BaseSpace; Illumina), PhiX contamination filtering (BBDuk; BBTools v39.01) (9), and demultiplexing with the Python package demultiplex (Laros JFJ, github.com/jfjlaros/demultiplex), allowing for one mismatch per barcode and two mismatches for linker and primer sequences. Extracted FASTQ reads were trimmed (220 and 150 nt, for the forward and reverse read, respectively), allowing for two errors per read.

*Metagenome sequencing*

Metagenomic sequencing libraries were prepared using the NEBNext Ultra II FS DNA Library Prep Kit for Illumina, and sequenced in paired-end mode (2x 100 bp) on the Illumina NovaSeq6000 platform. Sequencing data were quality-checked using fastQC v0.12.1. Adapters were trimmed and phiX contamination was removed using BBDuk (BBTools v39.01; (9) to a minimum length of 50 bp. Reads were k-trimmed from the right with a kmer of 21, minimum kmer of 11, and hamming distance of two along with the 'tpe' and 'tbo' options, and q-trimmed from the right at a Q-score of 15. Reads were interleaved using the reformat.sh script of BBMap v39.01.

*Phylogenetic trees of marker proteins*

Phylogenetic trees concatenated amino acid sequences of marker proteins were constructed in IQ-TREE v2.1.2 (7) using the best-fit model identified by ModelFinder (10) and 1,000 ultrafast bootstraps. The AOA tree was calculated using the best-fit model (LG+F+R5) according to the Bayesian Information Criterion (BIC); the genomes of *Nitrososphaera viennensis* and *N. gargensis* used as the outgroup. The NOB lineage IV *Nitrospira* tree was calculated using the best-fit model Q.plant+F+I+R4 based on the BIC. *N. moscoviensis* and *Ca*. Nitrospira inopinata were used as the outgroup. Genome-wide average nucleotide identities (ANI) of genomes and MAGs included in the concatenated marker gene trees were calculated with fastANI v1.33 (11) to assess species delineation (>96.5% ANI). Trees were visualised and annotated in iTOL v5 (8) and Adobe Illustrator.

*ddPCR Primer design, optimisation, and reaction*

The dominant AOA was quantified using a symbiont-specific primer set that targets a 175 bp fragment of the archaeal *amo*A gene (forward primer 5'-GCA GCA CCA TCC AGT CCT TT-3' and reverse primer 5'-TGT CCA AGC CCA ATC AGT GT-3'). The dominant NOB was quantified using a symbiont-specific primer set (forward primer 5'-TAA CGG CCT ACC AAG GCA AC-3' and reverse primer 5'-GTG GAA ACC CAC CCC ATC TT-3') targeting a 207 bp fragment of the 16S rRNA gene of the symbiont. Both primer sets were designed with the NCBI Primer-Blast tool (12).

Optimal primer annealing temperatures were assessed with temperature gradient PCRs, ranging from 55-64°C. Highest possible primer annealing temperatures at which a clear separation between positive and negative droplets was obtained were used for the subsequent ddPCR approach. Furthermore, to obtain optimal positive and negative droplet ratios, DNA extracts of all samples were diluted with PCR-grade water in a 1:10 ratio. Primer specificity was verified by adding DNA from a negative control sample containing a close relative to the target symbiont (i.e., AOA = *Ca.* Nitrosospongia ianthellae - DNA of the sponge holobiont *Ianthella basta*, NOB = *Ca*. Nitronereus thalassa).

Each ddPCR reaction consisted of QX200 ddPCR EvaGreen Supermix, 0.1 µM forward and reverse primers, 2 µl of DNA template, and PCR-grade water was used to adjust for the final volume of 22 µl. Next, 20 µL of each PCR reaction and 70 µL of QX200 droplet generation oil were loaded onto the designated wells of DG8 cartridges. Fully loaded cartridges were placed into the QX200 Droplet Generator, and 40 µl of the resulting droplets (minimum of 14,000 droplets) were transferred to 96-well PCR plates. For both AOA and NOB primer sets the following two-step PCR protocol was performed: 'initial denaturation' at 95°C for 5 min, 40 cycles of 'denaturation' at 95°C for 30 s and 'annealing and elongation' 62°C annealing temperature for 1 min, followed by a final signal stabilisation step at 4°C for 5 min and 90°C for 5 min. The ramp rate for each step was set to 2°C/s, and cycler lid temperature was set to 105°C throughout. ddPCR products were subsequently quantified using a QX200 Droplet Reader (Bio-Rad) in combination with the QX Manager Software (version 1.2 Standard Edition, Bio-Rad). The detection threshold was set automatically using the standard deviation function.

The archaeal symbiont with the highest relative abundance (ASV_3dn_o9y) revealed an average absolute abundance of 7.47 ± 6.6 (SD) x$10^4$ *amoA* gene copies per larva and the dominant *Nitrospira* symbiont (ASV_myj_c3n) revealed an average absolute abundance of 5.20 ± 3.38 (SD) x $10^4$ 16S rRNA gene copies per larva. Copy numbers per larva were used to calculate the copy numbers per wet weight larva (see below).

*Wet weight larva calculations*

Larvae of the sponge *C. matthewsi* are prolate spheroid in shape and are on average 572.39 µm (± 3.05 µm SE) long and 345.86 µm (± 3.40 µm SE) wide (13). Based on the average measurements the volume (in $\mu m^3$) of a larva was calculated with the following formula:

$$V = \frac{4}{3} \times \pi \times a \times b \times c$$

Next, the mass of a single larva (in g) was calculated based on the assumption that sponge larvae have the same density as water (1 $\mu m^3$ = 1 x $10^{-12}$ g).

*Fluorescence in situ hybridisation (FISH) probes*

The 16S rRNA-targeted oligonucleotide probes Arch915 (14) and Ntspa662 (15) were matched against full-length 16S rRNA gene sequence of *Ca*. Nitrosokoinonia kreatosae and *Ca*. Nitrosymbion coscinodermae, respectively, using probeBase (16).

- **Arch915:** 5’ - GTG CTC CCC CGC CAA TTC CT - 3’ (14)
  A tetra-labelled FISH probe was used for the visualisation of AOA symbionts in the tissue of *Coscionderma matthewsi* due to the low ribosome numbers of AOA (17). The probe had an exact match with the 16S rRNA gene sequence recovered from the *Ca*. Nitrosokoinonia kreatosae MAG.
- **Ntspa662:** 5’ - GGA ATT CCG CGC TCC TCT - 3’, used together with the unlabelled competitor probe 5'-GGA ATT CCG CTC TCC TCT -3' (15)
  The probe had one mismatch with the 16S rRNA gene sequence recovered from the *Ca*. Nitrosymbion coscinodermae MAG.
- **non-EUB338-I** 5'- ACT CCT ACG GGA GGC AGC -3' (18)

**Supplementary Figures**

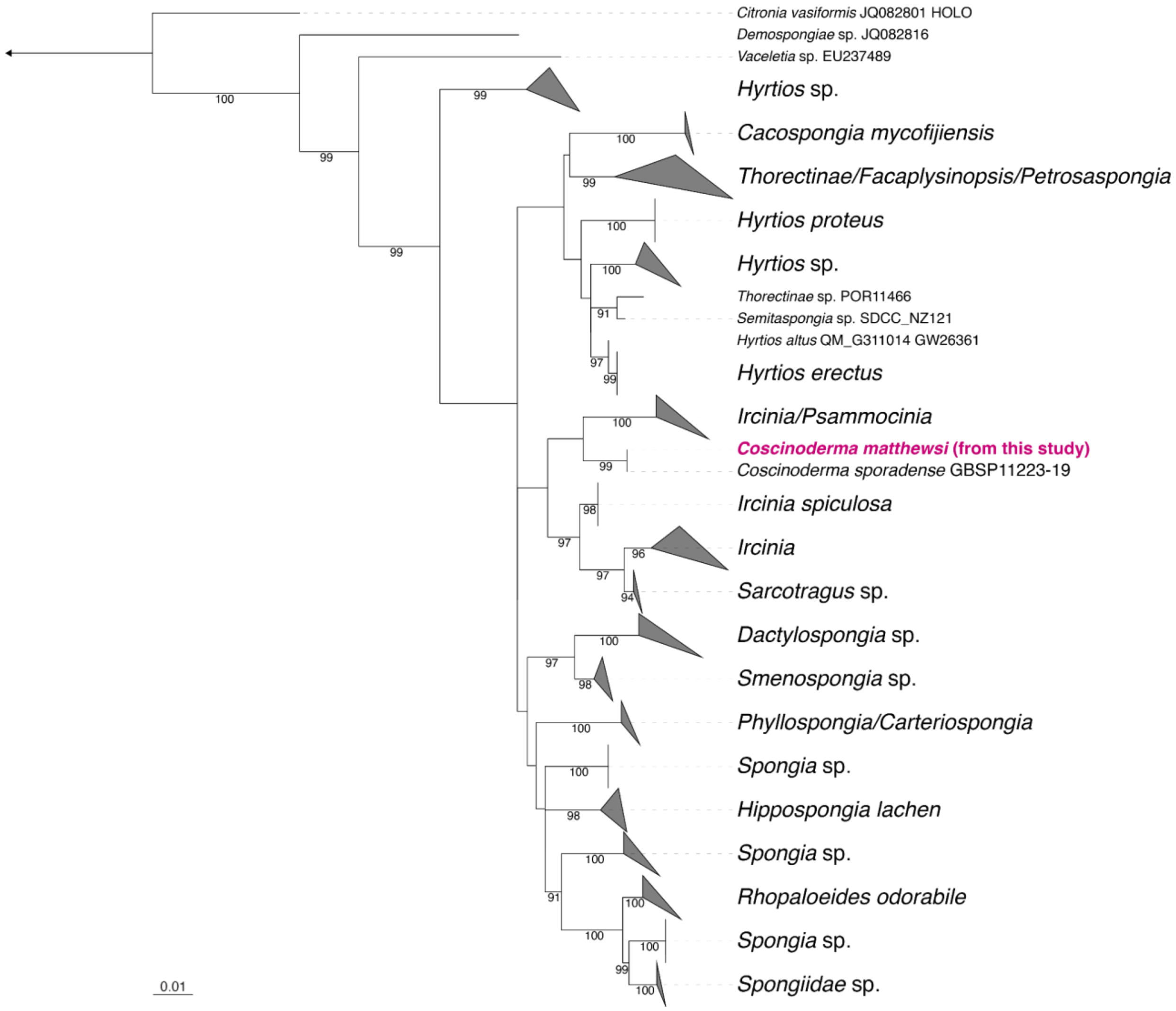


**Figure S1:** Phylogenetic analysis of the sponge host. Phylogenetic tree based on the eukaryotic marker gene CO1. Numbers at the branches indicate ultrafast bootstrap (n = 1,000) support. The scale bar corresponds to 0.01 estimated nucleotide substitution per site.

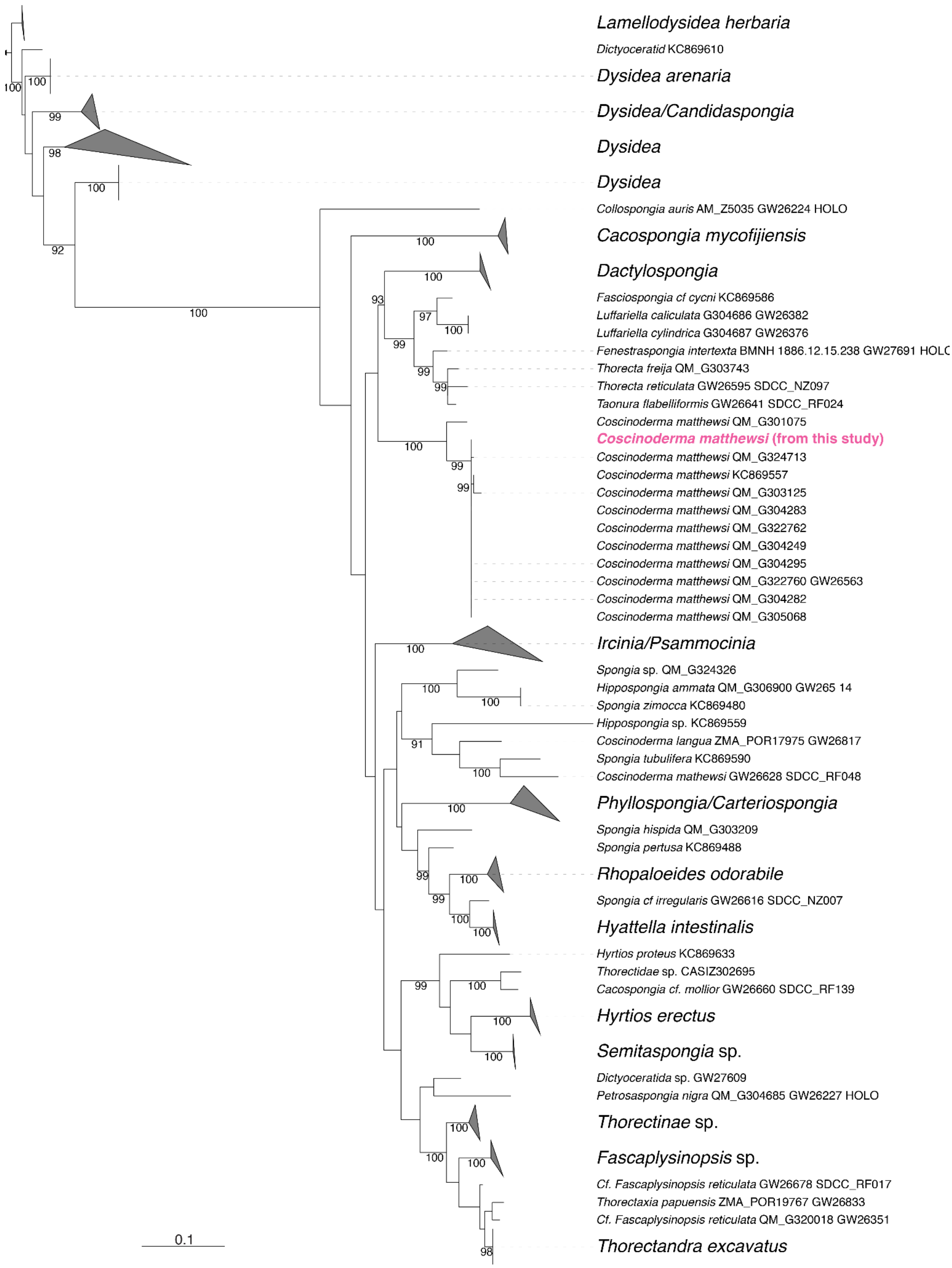


**Figure S2:** Phylogenetic analysis of the sponge host. Phylogenetic tree based on the 28S rRNA gene. Numbers at the branches indicate ultrafast bootstrap (n = 1,000) support. The scale bar corresponds to 0.1 estimated nucleotide substitution per site.

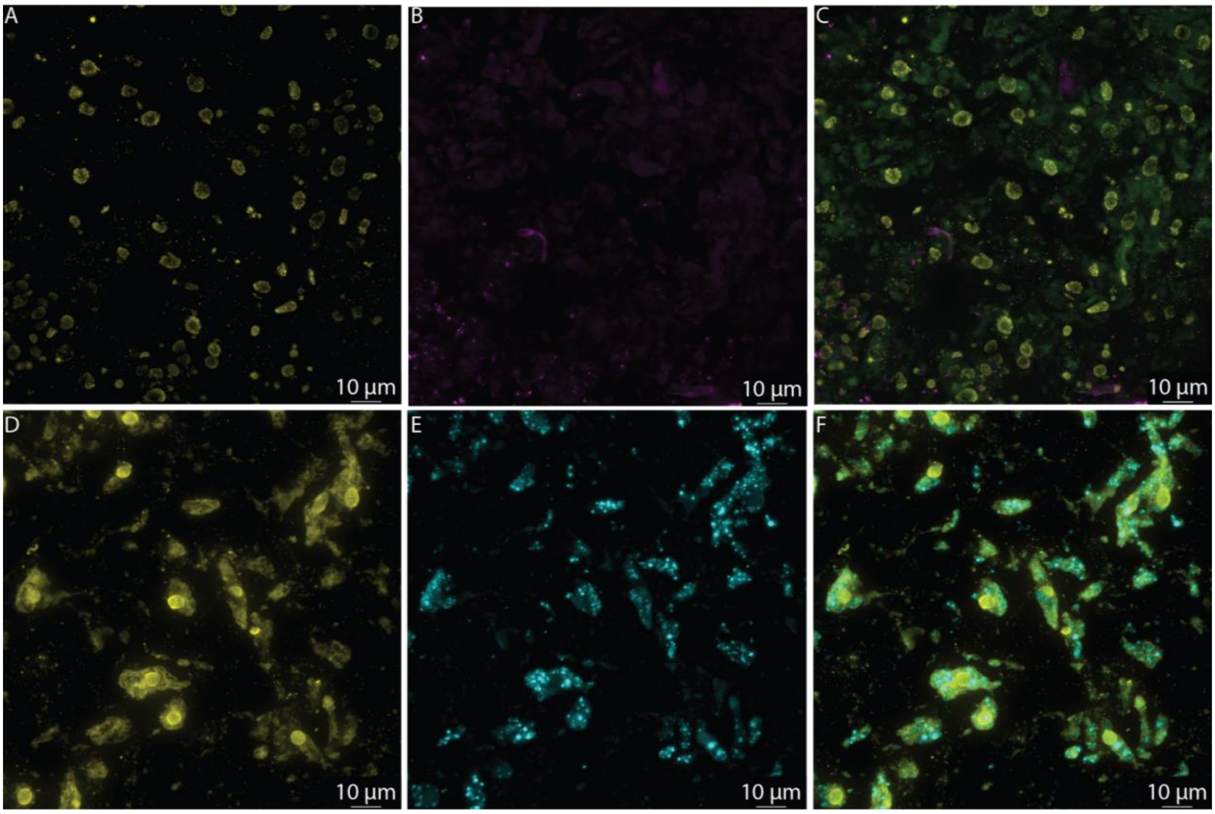


**Figure S3.** Fluorescence *in situ* hybridisation (FISH) in 5 µm cryosections of adult sponge tissue. A-C) Eukaryotic and prokaryotic DNA was stained using DAPI (in yellow), a tetra-labelled non-EUB338-I probe in Atto565 (magenta) was used to control for non-specific binding, and an overlay of the two channels. D-F) Eukaryotic and prokaryotic DNA was stained using DAPI (in yellow), a double-labelled non-EUB338-I FISH-probe in Cy5 (cyan) was used to control for non-specific binding, and an overlay of the two channels.

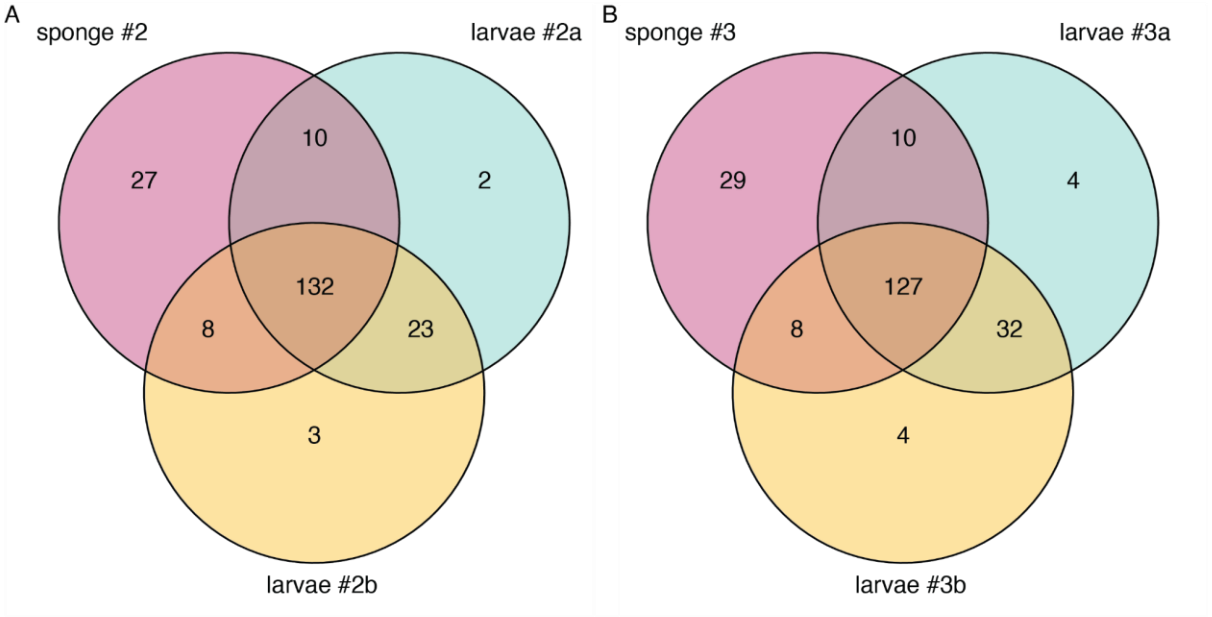


**Figure S4.** Number of unique and shared amplicon sequence variants (ASVs) between adult sponges and their freshly released larvae. A) Sponge individual #2 and its larvae shared 132 ASVs, representing up to 93% and 95% of the adult and larval microbiome, respectively. A total of 28 ASVs (mainly belonging to the phylum *Chloroflexota*) were detected only in the larvae samples but not in the adult mother sponge. B) Sponge individual #3 and its larvae shared 127 ASVs, contributing up to 87% and 94% of the adult and larval microbiome, respectively. 40 ASVs (mainly belonging to the phylum *Chloroflexota*) were detected solely in the larvae but not in the adult mother sponge microbiome.

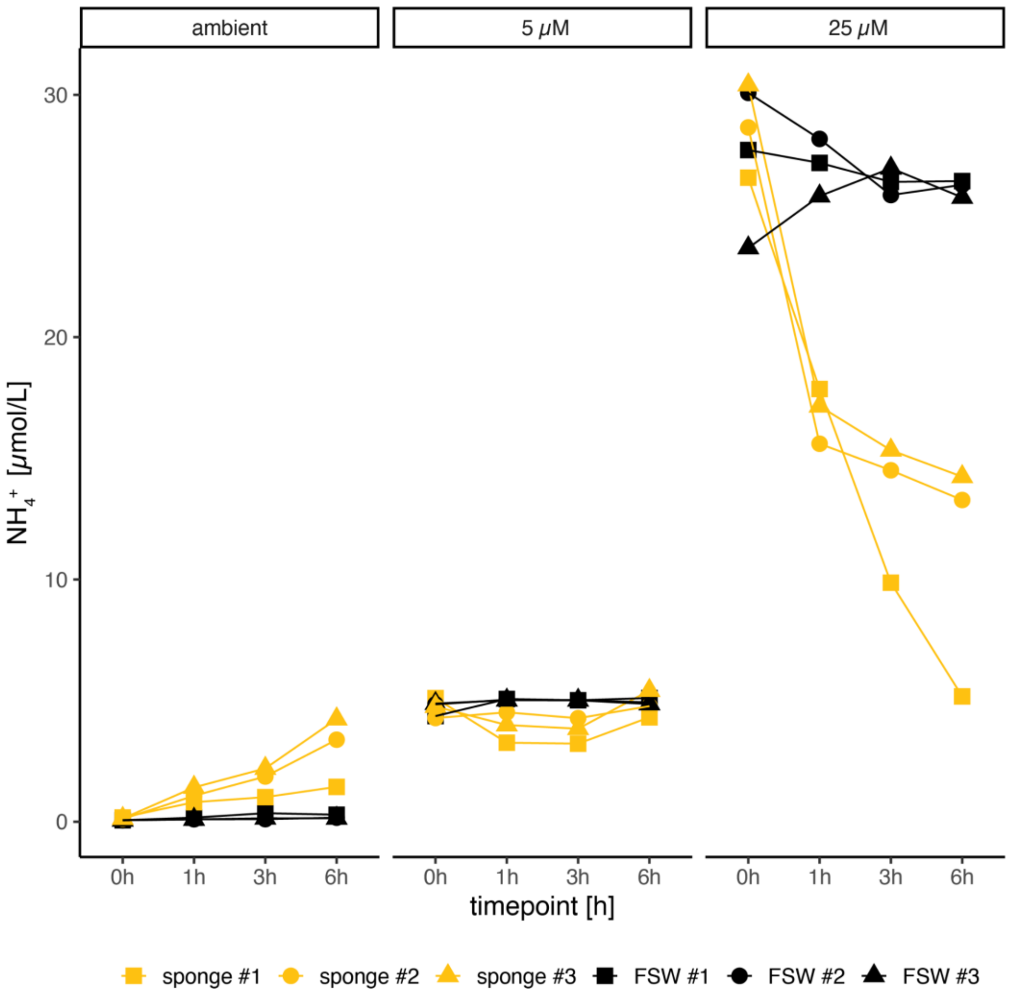


**Figure S5.** Ammonium ($NH_4^+$) concentrations per sponge individual incubations (yellow) and in abiotic control incubations (black) in 0.1 μm filter-sterilised seawater (FSW) during the 6h incubation experiment under ambient and amended ammonium concentrations (5 μM and 25 μM, respectively). The higher variability of the ammonium concentration at t0 in the 25 μM treatment may be due to a pipetting error.

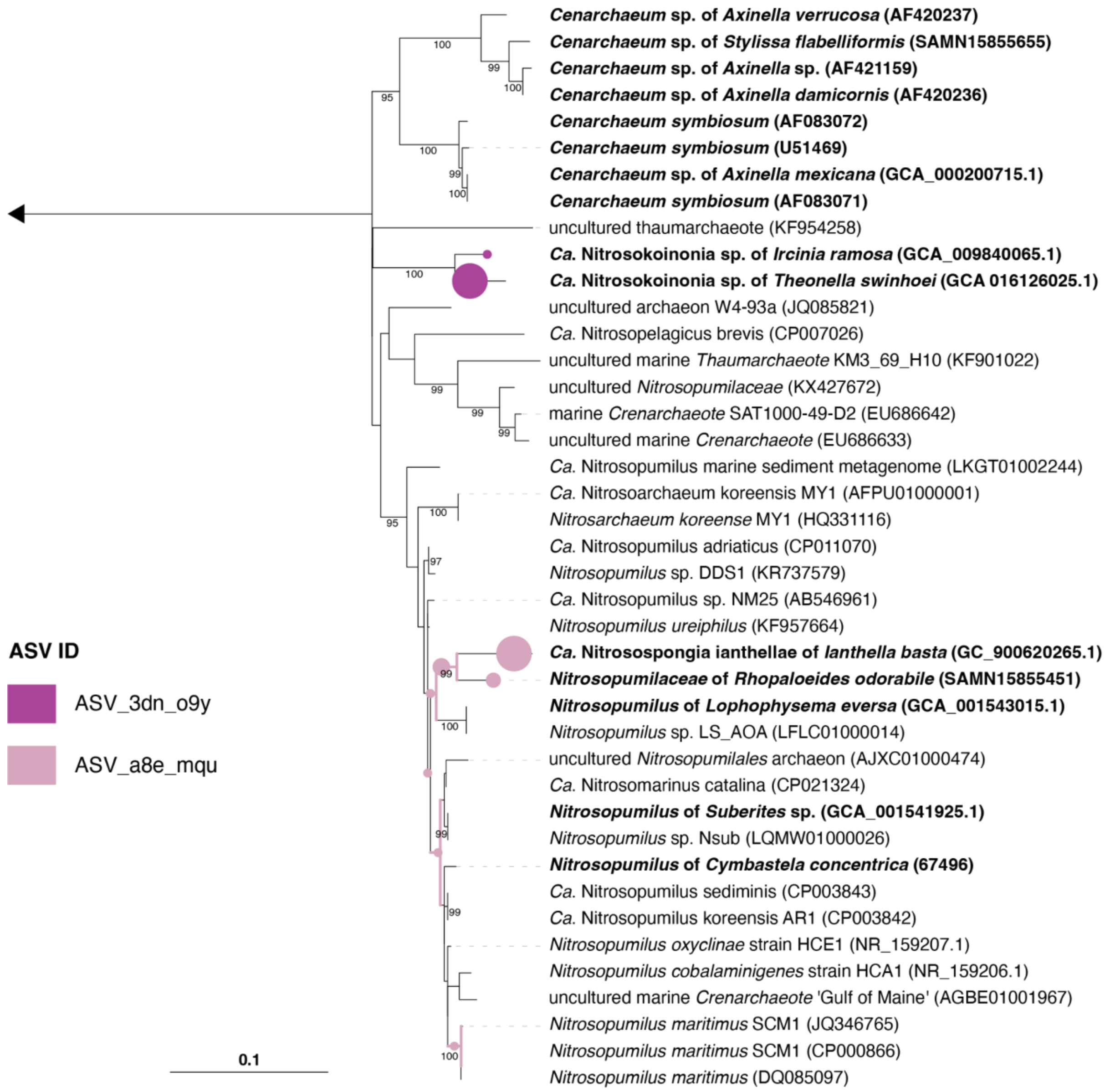


**Figure S6.** Phylogenetic tree of full-length 16S rRNA genes recovered from publicly available ammonia-oxidising archaea (AOA) genomes. Short 16S rRNA gene reads of the two dominant archaeal symbionts (amplicon sequence variants = ASV_3dn_o9y and ASV_a8e_mqu) were placed into the tree using pplacer v1.1alpha19. Numbers at the branches indicate ultrafast bootstrap (n = 1,000) support. The scale bar corresponds to 0.1 estimated nucleotide substitution per site.

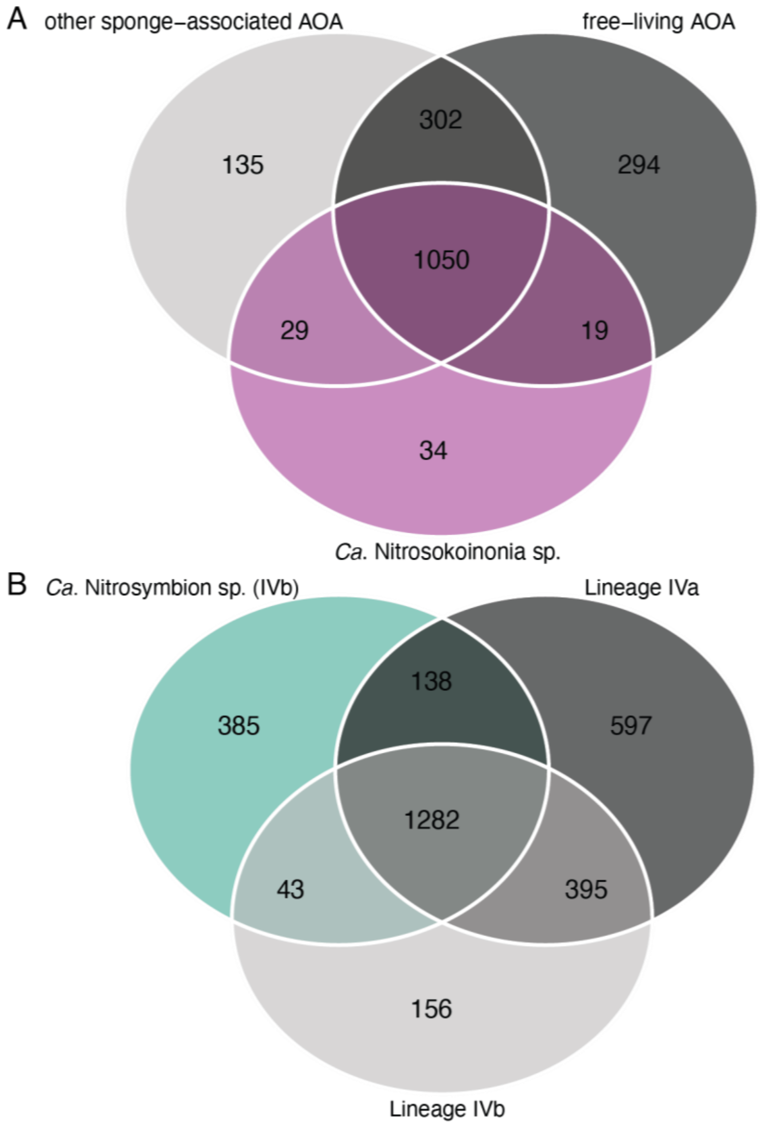


**Figure S7.** Number of unique and shared orthologs identified using eggnog mapper (eggNOG OGs - broadest hierarchical level). A) Shared and unique orthologs of selected ammonia-oxidising archaea (AOA) genomes. The selected genomes of the family *Nitrosopumilaceae* include other sponges-associated AOA (n = 8), free-living AOA (n = 29), and MAGs of the novel genus *Ca.* Nitrosokoinonia (n = 5). B) Shared and unique orthologs of selected nitrite-oxidising bacterial (NOB) *Nitrospirales* genomes. The compared genomes include representatives of the lineage IVa *Nitrospirales* (n = 9), the here described mostly marine-host associated genus *Ca*. Nitrosymbion (lineage IVb, n = 12) as well as other members of lineage IVb *Nitrospirales* (n = 4), including the recently described lineage *Ca*. Nitronereus thalassa. For a detailed overview of the included genomes please refer to the Supplementary Table S1. A description of the unique orthologs associated with the here described genera are provided in Supplementary Table S5 and S6.
