## Supplementary TablesS1-8 for "Novel nitrifying symbiont lineages are vertically inherited and widespread in marine sponges"

**Table S1:** Overview of genomes and metagenome assembled genomes (MAGs) included in this study. Asterisk (*) highlights MAGs that were dereplicated (based on 95% average nucleotide identity - ANI) with MAGs retrieved from this study.

| | | MAG_ID | Assembly | original publication | environment | host species | genome name | ANI [%] with MAG recoverd in this study | 16S rRNA sequence identity | included in genome tree | included in comperative analysis | GC | GC std | Genome size | # ambiguous bases | # scaffolds | # contigs | Longest scaffold | Longest contig | N50 (scaffolds) | N50 (contigs) | Mean scaffold length | Mean contig length | Coding density | Translation table | # predicted genes | Completeness | Contamination | Strain heterogeneity |
|---|---|---|---|---|---|---|---|---|---|---|---|---|---|---|---|---|---|---|---|---|---|---|---|---|---|---|---|---|---|
| ammonia-oxidizing archaeal genomes and MAGs | | **metabat_nocov.1** | **TBA** | **from this study** | **sponge** | ***Coscinoderma matthewsi*** | **Ca. Nitrosokoinonia kreatosae** | **100** | **NA** | **yes** | **new_genus** | **44.38%** | **2%** | **1400504** | **240** | **221** | **245** | **34155** | **34155** | **7949** | **7604** | **6337** | **5715** | **86.06%** | **11** | **1636** | **91.26** | **0.97** | **100** |
| | * | **IRC_PAM_SB0663_bin_5** | **GCA_009843815.1** | **Engelberts et al. (2020) ISME J** | **sponge** | ***Ircinia ramosa*** | **Ca. Nitrosokoinonia kreatosae** | **97.47** | **NA** | **yes** | **new_genus** | **45.56%** | **4%** | **1505883** | **860** | **162** | **194** | **27957** | **25073** | **12594** | **11225** | **9296** | **7758** | **86.51%** | **11** | **1716** | **97.57** | **0** | **0** |
| | | IRC_PAM_SB0662_bin_33 | GCA_009840065.1 | Engelberts et al. (2020) ISME J | sponge | *Ircinia ramosa* | Ca. Nitrosokoinonia sp. of *Ircinia ramosa* | 79.78 | 100 | yes | new_genus | 46.49% | 3% | 1573815 | 1350 | 209 | 227 | 34024 | 28289 | 9936 | 9524 | 7530 | 6927 | 82.27% | 11 | 1789 | 96.6 | 0 | 0 |
| | | GCA_016126025.1_ASM1612602v1_genomic | GCA_016126025.1 | Haber et al. (2021) Front Microbiol | sponge | *Theonella swinhoei* | Ca. Nitrosokoinonia sp. of *Theonella swinhoei* | 77.78 | 97 | yes | new_genus | 61.25% | 3% | 1859711 | 0 | 158 | 158 | 52333 | 52333 | 14074 | 14074 | 11770 | 11770 | 83.00% | 11 | 2028 | 95.63 | 0 | 0 |
| | | GCA_021296165.1_ASM2129616v1_genomic | GCA_021296165.1 | Burgsdorf et al. (2019) mSystems | sponge | *Petrosia ficiformis* | Ca. Nitrosokoinonia sp. of *Petrosia ficiformis* | 77.02 | NA | yes | new_genus | 61.65% | 3% | 1694402 | 11173 | 205 | 268 | 34955 | 34955 | 11482 | 9045 | 8265 | 6281 | 81.79% | 11 | 1859 | 90.78 | 0 | 0 |
| | | IB2_shallow_metabat_bins_Nitrosopumilus | GCA_900620265.1 | Moeller et al. (2019) Envir Microbiol | sponge | *Ianthella basta* | Ca. Nitrosospongia ianthellae | <75 | 92 | yes | sponge | 65.21% | 3% | 1778192 | 2120 | 34 | 57 | 261843 | 261843 | 133442 | 53670 | 52300 | 31159 | 89.61% | 11 | 1886 | 98.54 | 0 | 0 |
| | | STY3_bin_6 | SAMN15855655 | Robbins et al. (2021) ISME J | sponge | *Stylissa flabelliformis* | Cenarcheaum of Stylissa flabelliformis | <75 | 91 | yes | sponge | 67.38% | 2% | 1488362 | 4639 | 100 | 123 | 93182 | 76110 | 26624 | 20677 | 14884 | 12063 | 88.69% | 11 | 1622 | 98.54 | 0 | 0 |
| | | AXIM_hallam_GCA_000200715.1_genomic | GCA_000200715.1 | Hallam et al. (2006) PNAS | sponge | *Axinella mexicana* | Cenarchaeum symbiosum | <75 | 93 | yes | sponge | 57.38% | 0% | 2045086 | 230 | 1 | 6 | 2045086 | 1281157 | 2045086 | 1281157 | 2045086 | 340809 | 88.72% | 11 | 1924 | 99.03 | 0 | 0 |
| | | GCA_000220175.2_ASM22017v1_genomic | GCA_000220175.2 | Kim et al. (2011) J Bacteriol | soil | - | Nitrosoarchaeum koreense MY1 | <75 | 93 | yes | free-living | 32.67% | 0% | 1607695 | 0 | 1 | 1 | 1607695 | 1607695 | 1607695 | 1607695 | 1607695 | 1607695 | 90.08% | 11 | 1925 | 100 | 0 | 0 |
| | | APA_bin_56 | GCA_026712925.1 | Robbins et al. (2021) ISME J | sponge | *Aplysina aerophoba* | Nitrosopumilaceae of Aplysina aerophoba | <75 | NA | yes | sponge | 38.85% | 2% | 2186878 | 0 | 330 | 330 | 43697 | 43697 | 9798 | 9798 | 6627 | 6627 | 81.32% | 11 | 2609 | 91.59 | 0 | 0 |
| | | RHO2_bin_25 | GCA_028820595.1 | Robbins et al. (2021) ISME J | sponge | *Rhopaloeides odorabile* | Nitrosopumilaceae of Rhopaloeides odorabile | <75 | 93 | yes | sponge | 60.37% | 2% | 1990914 | 2420 | 140 | 175 | 80611 | 48392 | 20213 | 17585 | 14221 | 11363 | 83.79% | 11 | 2135 | 94.17 | 0 | 0 |
| | | GCA_000018465.1_ASM1846v1_genomic | GCA_000018465.1 | Walker et al. (2010) PNAS | marine | - | Nitrosopumilus maritimus SCM1 | <75 | 93 | yes | free-living | 34.17% | 0% | 1645259 | 0 | 1 | 1 | 1645259 | 1645259 | 1645259 | 1645259 | 1645259 | 1645259 | 91.96% | 11 | 1947 | 100 | 0.97 | 0 |
| | | CYMC_moitinhoThomas_67496.assembled | IMG_2626541593 | Moitinho-Silva et al. (2017) ISME J | sponge | *Cymbastela concentrica* | Nitrosopumilus of Cymbastela concentrica | <75 | 93 | yes | sponge | 38.42% | 5% | 2161854 | 3 | 637 | 637 | 89589 | 89589 | 12783 | 12783 | 3394 | 3394 | 83.48% | 11 | 2759 | 97.09 | 4.05 | 0 |
| | | LOPHE_tianQian_GCA_001543015.1_ASM154301v1_genomic | GCA_001543015.1 | Tian et al. (2016) Envir Microbiol | sponge | *Lophophysema eversa* | Nitrosopumilus of Lophophysema eversa | <75 | 93 | yes | sponge | 31.64% | 2% | 1925182 | 0 | 67 | 67 | 218566 | 218566 | 41513 | 41513 | 28734 | 28734 | 88.80% | 11 | 2269 | 100 | 0 | 0 |
| | | SUB_tianQian_GCA_001541925.1_ASM154192v1_genomic | GCA_001541925.1 | Tian et al. (2017) mSystems | sponge | *Suberites sp.* | Nitrosopumilus of Suberites sp. | <75 | 93 | yes | sponge | 31.40% | 1% | 1383621 | 0 | 26 | 26 | 200414 | 200414 | 91269 | 91269 | 53216 | 53216 | 88.44% | 11 | 1735 | 100 | 0 | 0 |
| | | GCA_000875775.1_ASM87577v1_genomic | GCA_000875775.1 | Bayer et al. (2015) ISME J | marine | - | Nitrosopumilus piranensis D3C | <75 | 93 | yes | free-living | 33.82% | 0% | 1713078 | 0 | 1 | 1 | 1713078 | 1713078 | 1713078 | 1713078 | 1713078 | 1713078 | 89.91% | 11 | 2145 | 100 | 0.97 | 0 |
| | | GCA_000204585.1_ASM20458v1_genomic | GCA_000204585.1 | | | | Ca. Nitrosoarchaeum limnia SFB1 | <75 | 93 | yes | free-living | 32.46% | 0% | 1772718 | 29483 | 1 | 76 | 1772718 | 230803 | 1772718 | 66478 | 1772718 | 22937 | 85.76% | 11 | 2162 | 98.06 | 0 | 0 |
| | | GCA_000242875.3_ASM24287v3_genomic | GCA_000242875.3 | | | | Ca. Nitrosopumilus salaria BD31 | <75 | 93 | yes | free-living | 33.79% | 3% | 1572957 | 4 | 171 | 171 | 158716 | 158716 | 28330 | 28330 | 9199 | 9199 | 89.60% | 11 | 2092 | 92.39 | 1.94 | 0 |
| | | GCA_000299365.1_ASM29936v1_genomic | GCA_000299365.1 | | | | Ca. Nitrosopumilus koreensis AR1 | <75 | 93 | yes | free-living | 34.18% | 0% | 1639964 | 0 | 1 | 1 | 1639964 | 1639964 | 1639964 | 1639964 | 1639964 | 1639964 | 90.15% | 11 | 2031 | 94.66 | 0 | 0 |
| | | GCA_000299395.1_ASM29939v1_genomic | GCA_000299395.1 | | | | Ca. Nitrosopumilus sediminis AR2 | <75 | 93 | yes | free-living | 33.59% | 0% | 1690905 | 0 | 1 | 1 | 1690905 | 1690905 | 1690905 | 1690905 | 1690905 | 1690905 | 89.97% | 11 | 2112 | 97.09 | 0 | 0 |
| | | GCA_000303155.1_ASM30315v1_genomic | GCA_000303155.1 | | | | Nitrososphaera gargensis | <75 | 84 | outgroup | no | 48.35% | 0% | 2833868 | 0 | 1 | 1 | 2833868 | 2833868 | 2833868 | 2833868 | 2833868 | 2833868 | 83.13% | 11 | 3768 | 100 | 2.91 | 0 |
| | | GCA_000402075.1_ASM40207v1_genomic | GCA_000402075.1 | | | | Thaumarchaeota archaeon SCGC AAA007-O23 | <75 | 92 | yes | free-living | 35.66% | 2% | 1104438 | 0 | 32 | 32 | 217386 | 217386 | 79020 | 79020 | 34514 | 34514 | 92.61% | 11 | 1354 | 97.33 | 2.09 | 100 |
| | | GCA_000685395.1_NKMY2_genomic | GCA_000685395.1 | | | | Ca. Nitrosotenuis chungbukensis MY2 | <75 | 92 | yes | no | 41.76% | 3% | 1762971 | 4 | 24 | 24 | 204441 | 204441 | 145899 | 145899 | 73457 | 73457 | 90.46% | 11 | 2047 | 99.03 | 0.97 | 0 |
| | | GCA_000723185.1_APF_1883_PRJEB4650_genomic | GCA_000723185.1 | | | | Ca. Nitrosotenuis uzonensis N4 | <75 | 91 | yes | no | 42.25% | 3% | 1636125 | 0 | 14 | 14 | 485325 | 485325 | 441459 | 441459 | 116866 | 116866 | 91.37% | 11 | 1921 | 100 | 0.97 | 0 |
| | | GCA_000812185.1_ASM81218v1_genomic | GCA_000812185.1 | | | | Ca. Nitrosopelagicus brevis CN25 | <75 | 91 | yes | free-living | 33.16% | 0% | 1232128 | 0 | 1 | 1 | 1232128 | 1232128 | 1232128 | 1232128 | 1232128 | 1232128 | 93.85% | 11 | 1470 | 99.51 | 0 | 0 |
| | | GCA_000955905.3_ASM95590v3_genomic | GCA_000955905.3 | | | | Ca. Nitrosotenuis cloacae SAT1 | <75 | 92 | yes | no | 41.00% | 0% | 1620156 | 0 | 1 | 1 | 1620156 | 1620156 | 1620156 | 1620156 | 1620156 | 1620156 | 92.10% | 11 | 1877 | 100 | 1.94 | 0 |
| | | GCA_000956175.1_ASM95617v1_genomic | GCA_000956175.1 | | | | Ca. Nitrosopumilus adriaticus NF5 | <75 | 94 | yes | free-living | 33.41% | 0% | 1803090 | 0 | 1 | 1 | 1803090 | 1803090 | 1803090 | 1803090 | 1803090 | 1803090 | 90.19% | 11 | 2210 | 100 | 0 | 0 |
| | | GCA_001437625.1_ASM143762v1_genomic | GCA_001437625.1 | | | | Nitrosopumilus sp. BACL13 MAG-121220-bin23 | <75 | 93 | yes | free-living | 31.01% | 2% | 1270387 | 0 | 183 | 183 | 67392 | 67392 | 13618 | 13618 | 6942 | 6942 | 90.05% | 11 | 1698 | 99.51 | 0.97 | 100 |
| | | GCA_001443365.1_ASM144336v1_genomic | GCA_001443365.1 | | | | Thaumarchaeota archaeon CSP1-1 | <75 | NA | yes | free-living | 34.42% | 1% | 1358434 | 81 | 10 | 12 | 249686 | 249685 | 222484 | 204537 | 135843 | 113196 | 89.93% | 11 | 1653 | 99.03 | 0 | 0 |
| | | GCA_001510275.1_ASM151027v1_genomic | GCA_001510275.1 | | | | Thaumarchaeota archaeon casp-thauma3 | <75 | NA | yes | free-living | 32.77% | 1% | 1221161 | 0 | 31 | 31 | 100507 | 100507 | 48651 | 48651 | 39392 | 39392 | 91.13% | 11 | 1532 | 94.17 | 0 | 0 |
| | | GCA_001627235.1_ASM162723v1_genomic | GCA_001627235.1 | | | | Nitrosopelagicus sp. REDSEA-S31_B2 | <75 | 91 | yes | free-living | 34.05% | 2% | 1009131 | 230 | 86 | 92 | 59797 | 59797 | 17107 | 17107 | 11734 | 10966 | 93.31% | 11 | 1346 | 92.64 | 1.05 | 100 |
| | | GCA_002156965.1_ASM215696v1_genomic | GCA_002156965.1 | | | | Ca. Nitrosomarinus catalina SPOT01 | <75 | 93 | yes | free-living | 31.44% | 0% | 1360076 | 0 | 1 | 1 | 1360076 | 1360076 | 1360076 | 1360076 | 1360076 | 1360076 | 91.88% | 11 | 1677 | 100 | 0 | 0 |
| | | GCA_002317795.1_ASM231779v1_genomic | GCA_002317795.1 | | | | Thaumarchaeota archaeon NORP47 | <75 | 93 | yes | free-living | 31.44% | 1% | 1906738 | 346 | 37 | 37 | 252867 | 252792 | 68949 | 68935 | 51533 | 51524 | 87.20% | 11 | 2373 | 93.85 | 0.97 | 100 |
| | | GCA_002506665.1_ASM250666v1_genomic | GCA_002506665.1 | | | | Nitrosopumilus sp. UBA526 | <75 | 94 | yes | free-living | 41.23% | 2% | 1389729 | 13984 | 89 | 236 | 111965 | 111965 | 25082 | 18613 | 15615 | 5829 | 83.77% | 11 | 1673 | 92.23 | 0 | 0 |
| | | GCA_002737445.1_ASM273744v1_genomic | GCA_002737445.1 | | | | Ca. Nitrosoarchaeum sp. Baikal-G1 | <75 | 93 | yes | free-living | 31.02% | 1% | 1193764 | 0 | 13 | 13 | 361196 | 361196 | 119318 | 119318 | 91828 | 91828 | 89.80% | 11 | 1461 | 99.03 | 1.94 | 0 |
| | | GCA_002737455.1_ASM273745v1_genomic | GCA_002737455.1 | | | | Thaumarchaeota archaeon Baikal-G2 | <75 | 92 | yes | free-living | 30.28% | 1% | 1139986 | 0 | 20 | 20 | 214392 | 214392 | 112971 | 112971 | 56999 | 56999 | 91.30% | 11 | 1444 | 99.03 | 0 | 0 |
| | | GCA_002787055.1_ASM278705v1_genomic | GCA_002787055.1 | | | | Ca. Nitrosotenuis aquarius AQ6f | <75 | 91 | yes | no | 42.18% | 0% | 1697207 | 0 | 1 | 1 | 1697207 | 1697207 | 1697207 | 1697207 | 1697207 | 1697207 | 92.42% | 11 | 2009 | 99.68 | 1.94 | 0 |
| | | GCA_003175215.1_ASM317521v1_genomic | GCA_003175215.1 | | | | Ca. Nitrosopumilus sp. NM25 | <75 | 93 | yes | free-living | 33.84% | 1% | 1757809 | 0 | 14 | 14 | 317662 | 317662 | 287590 | 287590 | 125558 | 125558 | 89.24% | 11 | 2114 | 100 | 0 | 0 |
| | | GCA_003331425.1_ASM333142v1_genomic | GCA_003331425.1 | | | | Nitrosopumilus sp. MED-G94 | <75 | 93 | yes | free-living | 31.19% | 1% | 1235315 | 0 | 16 | 16 | 290261 | 290261 | 102005 | 102005 | 77207 | 77207 | 91.51% | 11 | 1561 | 100 | 0 | 0 |
| | | GCA_003569705.1_ASM356970v1_genomic | GCA_003569705.1 | | | | Ca. Nitrosoarchaeum sp. BD3 | <75 | 93 | yes | free-living | 33.00% | 1% | 1462514 | 0 | 10 | 10 | 259406 | 259406 | 250506 | 250506 | 146251 | 146251 | 88.16% | 11 | 1869 | 98.54 | 1.94 | 66.67 |
| | | GCA_003702465.1_ASM370246v1_genomic | GCA_003702465.1 | | | | Ca. Nitrosomarinus sp. GoM_MAG2 | <75 | NA | yes | free-living | 30.91% | 2% | 1127829 | 1630 | 111 | 139 | 54501 | 41961 | 15297 | 10423 | 10161 | 8102 | 91.52% | 11 | 1519 | 93.2 | 0 | 0 |
| | | GCA_003702495.1_ASM370249v1_genomic | GCA_003702495.1 | | | | Nitrosopumilus sp. GoM_MAG4 | <75 | NA | yes | free-living | 34.55% | 2% | 1266765 | 580 | 93 | 106 | 101600 | 74425 | 22568 | 21766 | 13621 | 11945 | 91.74% | 11 | 1648 | 95 | 1.94 | 50 |
| | | GCA_003702525.1_ASM370252v1_genomic | GCA_003702525.1 | | | | Nitrosopumilus sp. GoM_MAG5 | <75 | NA | yes | free-living | 34.53% | 2% | 1279624 | 0 | 64 | 64 | 120092 | 120092 | 32296 | 32296 | 19994 | 19994 | 92.37% | 11 | 1637 | 92.72 | 2.91 | 66.67 |
| | | GCA_003702545.1_ASM370254v1_genomic | GCA_003702545.1 | | | | Nitrosopumilus sp. GoM_MAG6 | <75 | NA | yes | free-living | 34.87% | 1% | 1224048 | 1430 | 79 | 96 | 58201 | 58201 | 26433 | 20758 | 15494 | 12736 | 91.49% | 11 | 1580 | 91.26 | 0.97 | 0 |
| | | GCA_003724325.1_ASM372432v1_genomic | GCA_003724325.1 | | | | Nitrosopumilus sp. D6 | <75 | 93 | yes | free-living | 47.91% | 2% | 1171110 | 0 | 106 | 106 | 40077 | 40077 | 20317 | 20317 | 11048 | 11048 | 89.31% | 11 | 1470 | 92.23 | 0 | 0 |
| | | GCA_004322465.1_ASM432246v1_genomic | GCA_004322465.1 | | | | Ca. Nitrosotenuis sp. GW928_bin.17 | <75 | NA | yes | no | 42.65% | 3% | 1681024 | 0 | 126 | 126 | 71974 | 71974 | 24692 | 24692 | 13341 | 13341 | 90.95% | 11 | 2001 | 91.76 | 2.91 | 33.33 |
| | | GCA_005798405.1_ASM579840v1_genomic | GCA_005798405.1 | | | | Nitrosopumilaceae archaeon F1-80-MAGs016 | <75 | NA | yes | free-living | 33.42% | 2% | 1247066 | 0 | 66 | 66 | 198543 | 198543 | 30333 | 30333 | 18895 | 18895 | 90.31% | 11 | 1560 | 93.2 | 0 | 0 |
| | | GCA_005877205.1_ASM587720v1_genomic | GCA_005877205.1 | | | | Thaumarchaeota archaeon TA_9 | <75 | 88 | yes | no | 38.69% | 2% | 1562348 | 247 | 98 | 105 | 101168 | 101168 | 33414 | 33414 | 15942 | 14877 | 88.69% | 11 | 1885 | 97.5 | 0 | 0 |
| | | GCA_005877305.1_ASM587730v1_genomic | GCA_005877305.1 | | | | Thaumarchaeota archaeon TA_20 | <75 | NA | yes | no | 35.68% | 2% | 1329581 | 1003 | 112 | 126 | 56293 | 56293 | 17173 | 16153 | 11871 | 10544 | 89.30% | 11 | 1715 | 92.64 | 1.94 | 0 |
| | | GCA_006740685.1_ASM674068v1_genomic | GCA_006740685.1 | | | | Ca. Nitrosopumilus sp. SW | <75 | 93 | yes | free-living | 33.76% | 0% | 1499620 | 0 | 1 | 1 | 1499620 | 1499620 | 1499620 | 1499620 | 1499620 | 1499620 | 91.19% | 11 | 1833 | 100 | 0.97 | 0 |
| | | GCA_007280335.1_ASM728033v1_genomic | GCA_007280335.1 | | | | Nitrosopumilales archaeon PowLak16_MAG10 | <75 | 89 | yes | no | 35.51% | 1% | 1296374 | 0 | 84 | 84 | 58981 | 58981 | 20964 | 20964 | 15433 | 15433 | 90.38% | 11 | 1581 | 94.17 | 0 | 0 |
| | | GCA_900065925.1_NDEV.embl.gz_genomic | GCA_900065925.1 | | | | Ca. Nitrosotalea devanaterra ND1 | <75 | 88 | yes | no | 37.07% | 0% | 1805304 | 0 | 1 | 1 | 1805304 | 1805304 | 1805304 | 1805304 | 1805304 | 1805304 | 90.58% | 11 | 2074 | 98.54 | 0 | 0 |
| | | GCA_900143675.1_Candidatus_Nitrosotalea_sinensis_Nd2_whole_genome_genomic | GCA_900143675.1 | | | | Ca. Nitrosotalea sinensis ND2 | <75 | 88 | yes | no | 37.37% | 3% | 1598048 | 0 | 10 | 10 | 717560 | 717560 | 284249 | 284249 | 159805 | 159805 | 91.13% | 11 | 1880 | 99.51 | 0.97 | 0 |
| | | GCA_900167955.1_ASM90016795v1_genomic | GCA_900167955.1 | | | | Ca. Nitrosotalea bavarica SbT1 | <75 | 88 | yes | no | 35.98% | 1% | 1553159 | 182 | 19 | 24 | 424330 | 278831 | 164480 | 106636 | 81745 | 64707 | 89.64% | 11 | 1833 | 97.57 | 1.94 | 50 |
| | | GCA_900177045.1_Candidatus_Nitrosotalea_koreensis_CS_whole_genome_genomic | GCA_900177045.1 | | | | Ca. Nitrosotalea okcheonensis CS | <75 | 88 | yes | no | 37.47% | 0% | 1972585 | 250 | 1 | 8 | 1972585 | 922146 | 1972585 | 582823 | 1972585 | 246542 | 87.65% | 11 | 2330 | 99.51 | 0 | 0 |
| | | GCF_000698785.1_ASM69878v1_genomic | GCF_000698785.1 | | | | Nitrososphaera viennensis | <75 | 84 | outgroup | no | 52.72% | 0% | 2527938 | 0 | 1 | 1 | 2527938 | 2527938 | 2527938 | 2527938 | 2527938 | 2527938 | 85.86% | 11 | 3015 | 100 | 0.97 | 0 |
| nitrite-oxidizing bacterial genomes and MAGs | * | **CO36405bin_1** | **GCA_011523385.1** | **Glasl et al. (2020) ISME J** | **sponge** | **Coscinoderma matthewsi** | **Ca. Nitrosymbion coscinodermae** | **97.90** | **100** | **yes** | **new_genus** | **56.70%** | **6%** | **2506777** | **900** | **38** | **47** | **342572** | **342572** | **122077** | **104706** | **65968** | **53317** | **88.38%** | **11** | **2410** | **98.64** | **3.86** | **0** |
| | | GCA_001273775.1_ASM127377v1_genomic | GCA_001273775.1 | | | | Nitrospira moscoviensis | <75 | 89 | outgroup | no | 61.99% | 0% | 4589485 | 0 | 1 | 1 | 4589485 | 4589485 | 4589485 | 4589485 | 4589485 | 4589485 | 89.40% | 11 | 4507 | 95.91 | 6.55 | 0 |
| | | GCA_001458695.1_NiCh1_genomic | GCA_001458695.1 | | | | Ca. Nitrospira inopinata | <75 | 89 | outgroup | no | 59.23% | 0% | 3295117 | 0 | 1 | 1 | 3295117 | 3295117 | 3295117 | 3295117 | 3295117 | 3295117 | 88.82% | 11 | 3152 | 96.82 | 4.77 | 0 |
| | | GCA_001643555.1_ASM164355v1_genomic | GCA_001643555.1 | | | | Nitrospirae bacterium SPGG5 | <75 | NA | yes | IVb | 50.52% | 2% | 1328125 | 401 | 260 | 260 | 13890 | 13890 | 5385 | 5381 | 5108 | 5107 | 83.49% | 11 | 1470 | 61.31 | 1.41 | 0 |
| | | GCA_002238765.1_ASM223876v1_genomic | GCA_002238765.1 | Slaby et al. (2017) ISME J | sponge | Aplysina aerophoba | Ca. Nitrosymbion sp. | 84.55 | 99 | yes | new_genus | 56.24% | 3% | 3038306 | 0 | 115 | 115 | 176720 | 176720 | 44884 | 44884 | 26420 | 26420 | 86.36% | 11 | 3122 | 94.09 | 2.73 | 0 |
| | | GCA_002790795.1_ASM279079v1_genomic | GCA_002790795.1 | | | | Nitrospirae_bacterium_CG_4_9_14_3_um_filter_51_5 | <75 | 89 | yes | no | 50.92% | 3% | 1430058 | 14332 | 568 | 838 | 11257 | 9430 | 2883 | 2346 | 2518 | 1689 | 82.40% | 11 | 1745 | 60.02 | 0.1 | 0 |
| | | GCA_003695915.1_ASM369591v1_genomic | GCA_003695915.1 | | | | Nitrospirales bacterium J017 | <75 | 92 | yes | IVa | 54.77% | 2% | 3706191 | 0 | 135 | 135 | 170004 | 170004 | 58798 | 58798 | 27453 | 27453 | 86.75% | 11 | 3694 | 96.48 | 3.18 | 0 |
| | | GCA_003696975.1_ASM369697v1_genomic | GCA_003696975.1 | | | | Nitrospirales bacterium J031 | <75 | 95 | yes | IVb | 53.77% | 2% | 2769672 | 0 | 273 | 273 | 55721 | 55721 | 14752 | 14752 | 10145 | 10145 | 88.50% | 11 | 2805 | 95.59 | 3.28 | 14.29 |
| | | GCA_007570995.1_ASM757099v1_genomic | GCA_007570995.1 | Robbins et al. (2021) ISME J | sponge | Stylissa flabelliformis | Ca. Nitrosymbion sp. | 76.84 | 96 | yes | new_genus | 57.34% | 3% | 1976489 | 0 | 186 | 186 | 136189 | 136189 | 14721 | 14721 | 10626 | 10626 | 88.93% | 11 | 1912 | 83.96 | 0.91 | 0 |
| | | GCA_009840285.1_ASM984028v1_genomic | GCA_009840285.1 | Engelberts et al. (2020) ISME J | sponge | Ircinia ramosa | Ca. Nitrosymbion sp. | 88.84 | 100 | yes | new_genus | 57.08% | 2% | 2201017 | 240 | 113 | 119 | 78911 | 78911 | 27767 | 27767 | 19478 | 18494 | 88.73% | 11 | 2165 | 95.45 | 3.86 | 20 |
| | | GCA_009842645.1_ASM984264v1_genomic | GCA_009842645.1 | Engelberts et al. (2020) ISME J | sponge | Ircinia ramosa | Ca. Nitrosymbion sp. | 89.81 | 98 | yes | new_genus | 55.64% | 3% | 2549912 | 960 | 156 | 171 | 83283 | 83283 | 28164 | 23889 | 16346 | 14906 | 86.86% | 11 | 2476 | 95.71 | 2.5 | 0 |
| | | GCA_011088105.1_ASM1108810v1_genomic | GCA_011088105.1 | | sponge | Mycale hentscheli | Nitrospirota bacterium MH1_Bin19 | <75 | 93 | yes | IVb | 48.21% | 1% | 2595516 | 30 | 70 | 73 | 139627 | 137014 | 65867 | 65867 | 37079 | 35555 | 87.20% | 11 | 2580 | 95.55 | 0.91 | 0 |
| | | GCA_012270585.1_ASM1227058v1_genomic | GCA_012270585.1 | Robbins et al. (2021) ISME J | coral | Porites lutea | Ca. Nitrosymbion sp. | 85.11 | 99 | yes | new_genus | 55.80% | 2% | 2503252 | 361 | 89 | 95 | 153090 | 153090 | 51555 | 51555 | 28126 | 26346 | 87.19% | 11 | 2476 | 97.73 | 3.01 | 0 |
| | | GCA_020027205.1_ASM2002720v1_genomic | GCA_020027205.1 | | cold seep | - | Ca. Nitrosymbion sp. MEX2-MAG7 | 77.70 | 96 | yes | new_genus | 61.86% | 2% | 1914574 | 0 | 29 | 29 | 208821 | 208821 | 115745 | 115745 | 66020 | 66020 | 87.52% | 11 | 1855 | 93.68 | 0.91 | 0 |
| | | GCA_020027535.1_ASM2002753v1_genomic | GCA_020027535.1 | | cold seep | - | Ca. Nitrosymbion sp. MEX2-MAG12 | 85.02 | NA | yes | new_genus | 56.31% | 2% | 2228583 | 0 | 140 | 140 | 71013 | 71013 | 25042 | 25042 | 15918 | 15918 | 87.66% | 11 | 2297 | 97.16 | 2.8 | 25 |
| | | GCA_020348185.1_ASM2034818v1_genomic | GCA_020348185.1 | | | | Nitrospirota bacterium bin371 | <75 | NA | yes | IVa | 49.41% | 0% | 2456183 | 50500 | 1 | 505 | 2456183 | 27797 | 2456183 | 6445 | 2456183 | 4764 | 76.10% | 11 | 2345 | 76.41 | 0.91 | 0 |
| | | GCA_020632635.1_ASM2063263v1_genomic | GCA_020632635.1 | | | | Nitrospiraceae bacterium HK-STAS-NITR-1 | <75 | 93 | yes | IVa | 50.31% | 0% | 2991882 | 0 | 7 | 7 | 1019991 | 1019991 | 714656 | 714656 | 427412 | 427412 | 87.78% | 11 | 2762 | 92.03 | 0 | 0 |
| | | GCA_022824735.1_ASM2282473v1_genomic | GCA_022824735.1 | | sponge | Ircinia sp. | Ca. Nitrosymbion sp. | 89.38 | NA | yes | new_genus | 56.49% | 2% | 2315678 | 0 | 151 | 151 | 49235 | 49235 | 19262 | 19262 | 15336 | 15336 | 88.42% | 11 | 2365 | 91.31 | 4.55 | 20 |
| | | GCA_022825885.1_ASM2282588v1_genomic | GCA_022825885.1 | | sponge | Ircinia sp. | Ca. Nitrosymbion sp. | 85.89 | 99 | yes | new_genus | 56.19% | 1% | 2377051 | 0 | 50 | 50 | 171458 | 171458 | 78259 | 78259 | 47541 | 47541 | 87.82% | 11 | 2348 | 94.03 | 0 | 0 |
| | | GCA_022828225.1_ASM2282822v1_genomic | GCA_022828225.1 | | sponge | Ircinia sp. | Ca. Nitrosymbion sp. | 87.58 | 99 | yes | new_genus | 57.12% | 1% | 2362088 | 0 | 24 | 24 | 426321 | 426321 | 215793 | 215793 | 98420 | 98420 | 87.73% | 11 | 2254 | 97.67 | 1.82 | 0 |
| | | GCA_902500695.1_RBC003_genomic | GCA_902500695.1 | | | | uncultured Nitrospira sp. | <75 | 89 | yes | IVa | 50.07% | 2% | 4276258 | 11910 | 175 | 295 | 136597 | 93918 | 45187 | 24838 | 24436 | 14455 | 82.24% | 11 | 4052 | 94.03 | 1.82 | 0 |
| | | **metabat_nocov.4** | **TBA** | **from this study** | **sponge** | **Coscinoderma matthewsi** | **Ca. Nitrosymbion coscinodermae** | **100** | **100** | **yes** | **new_genus** | **56.53%** | **2%** | **2289212** | **680** | **328** | **396** | **49677** | **43986** | **9027** | **8295** | **6979** | **5779** | **87.59%** | **11** | **2471** | **96.59** | **5** | **50** |
| | | Nitrospira_denmark_2022_genomic | SAMN32641989 | Mueller et al. (2023) ISME J | marine | | Ca. Nitrospira neomarina | <75 | 92 | yes | IVa | 50.23% | 0% | 4797949 | 1298 | 1 | 5 | 4797949 | 2256338 | 4797949 | 1386234 | 4797949 | 959330 | 83.07% | 11 | 4369 | 96.71 | 1.82 | 0 |
| | | Nitrospira_elba_2022_genomic | SAMN32641646 | Mueller et al. (2023) ISME J | marine | | Ca. Nitronereus thalassa | <75 | 95 | yes | IVb | 48.50% | 0% | 4014579 | 0 | 2 | 2 | 3358140 | 3358140 | 3358140 | 3358140 | 2007290 | 2007290 | 86.86% | 11 | 3723 | 97.67 | 1.82 | 50 |
| | | Nitrospira_maine_2022_genomic | SAMN32641643 | Mueller et al. (2023) ISME J | marine | | Nitrospira sp. MA-1 Maine enrichment | <75 | 91 | yes | IVa | 49.91% | 1% | 4761458 | 0 | 7 | 7 | 1776099 | 1776099 | 1264914 | 1264914 | 680208 | 680208 | 82.92% | 11 | 4337 | 96.64 | 1.82 | 0 |
| | | Nitrospira_vancouver_2022_genomic | SAMN32641628 | Mueller et al. (2023) ISME J | marine | | Ca. Nitrospira allomarina | <75 | 92 | yes | IVa | 49.99% | 0% | 4553057 | 0 | 1 | 1 | 4553057 | 4553057 | 4553057 | 4553057 | 4553057 | 4553057 | 83.78% | 11 | 4167 | 96.76 | 0.91 | 0 |
| | | NSPALKALI_genomic | PRJEB34830 (ENA) | Daebeler et al (2020) ISME J | saline lake | | Ca. Nitrospira alkalitoleranse KS | <75 | 93 | yes | IVa | 51.40% | 2% | 4886114 | 0 | 87 | 87 | 341326 | 341326 | 109063 | 109063 | 56162 | 56162 | 81.63% | 11 | 4802 | 95.83 | 4.83 | 12.5 |
| | | Nitrospira_marina_genomic | 2596583682 (JGI IM | Bayer et al. (2020) ISME J | marine | | Nitrospira marina Nb-295 | <75 | 92 | yes | IVa | 49.93% | 1% | 4524875 | 30 | 9 | 12 | 1742493 | 1727571 | 805880 | 610556 | 502764 | 377070 | 83.75% | 11 | 4136 | 96.76 | 0.92 | 0 |

**Table S2:** Integrated Microbial Next Generation (IMNGS) output file.

| Sample_Name | IMNGS Description | Size (no. | Ca. Nitrosokoinonia sp. | Ca. Nitrosymbion | type | percent | Description_Checked |
|---|---|---|---|---|---|---|---|
| SRR3233807 | Xestospongia testudinaria | 16624 | 16536 | 0 | AOA | 99.47 | sponge |
| SRR3157231 | Xestospongia testudinaria | 3273 | 3230 | 0 | AOA | 98.69 | sponge |
| SRR3233805 | Xestospongia testudinaria | 9079 | 8889 | 0 | AOA | 97.91 | sponge |
| SRR3157230 | Xestospongia testudinaria | 3488 | 3406 | 0 | AOA | 97.65 | sponge |
| SRR3157215 | Aaptos suberitoides | 3036 | 2946 | 0 | AOA | 97.04 | sponge |
| SRR1585356 | marine metagenome | 2444 | 2370 | 0 | AOA | 96.97 | sponge |
| SRR1595984 | marine metagenome | 2426 | 2352 | 0 | AOA | 96.95 | sponge |
| SRR3157228 | Xestospongia testudinaria | 3033 | 2933 | 0 | AOA | 96.70 | sponge |
| SRR1595986 | marine metagenome | 1997 | 1860 | 0 | AOA | 93.14 | sponge |
| SRR1585358 | marine metagenome | 2023 | 1884 | 0 | AOA | 93.13 | sponge |
| SRR3157220 | Aaptos suberitoides | 3640 | 3358 | 0 | AOA | 92.25 | sponge |
| SRR1585355 | marine metagenome | 3535 | 3124 | 0 | AOA | 88.37 | sponge |
| SRR1595983 | marine metagenome | 3475 | 3069 | 0 | AOA | 88.32 | sponge |
| SRR1585357 | marine metagenome | 3824 | 3133 | 0 | AOA | 81.93 | sponge |
| SRR1595985 | marine metagenome | 3747 | 3060 | 0 | AOA | 81.67 | sponge |
| SRR3233806 | Xestospongia testudinaria | 9380 | 6960 | 0 | AOA | 74.20 | sponge |
| SRR3157218 | Aaptos suberitoides | 3278 | 1513 | 0 | AOA | 46.16 | sponge |
| SRR3233789 | Petrosia spheroida | 5393 | 2024 | 0 | AOA | 37.53 | sponge |
| SRR8306885 | air metagenome | 1437 | 0 | 257 | NOB | 17.88 | other |
| SRR2976121 | sponge metagenome | 5036 | 0 | 869 | NOB | 17.26 | sponge |
| SRR2976120 | sponge metagenome | 4603 | 0 | 793 | NOB | 17.23 | sponge |
| SRR3147495 | Stylissa carteri | 4293 | 0 | 733 | NOB | 17.07 | sponge |
| SRR2976119 | sponge metagenome | 4156 | 0 | 622 | NOB | 14.97 | sponge |
| SRR2976108 | sponge metagenome | 3397 | 0 | 505 | NOB | 14.87 | sponge |
| SRR2976111 | sponge metagenome | 3625 | 0 | 521 | NOB | 14.37 | sponge |
| SRR2976126 | sponge metagenome | 5546 | 0 | 787 | NOB | 14.19 | sponge |
| SRR7591410 | coral metagenome | 23453 | 3307 | 121 | AOA and NOB | 14.10 | coral |
| SRR7036626 | marine metagenome | 15606 | 0 | 2194 | NOB | 14.06 | sponge |
| SRR7598645 | marine metagenome | 15613 | 0 | 2194 | NOB | 14.05 | sponge |
| SRR7036625 | marine metagenome | 18411 | 0 | 2512 | NOB | 13.64 | sponge |
| SRR7598644 | marine metagenome | 18449 | 0 | 2512 | NOB | 13.62 | sponge |
| SRR2976112 | sponge metagenome | 4069 | 0 | 550 | NOB | 13.52 | sponge |
| SRR2976109 | sponge metagenome | 4020 | 0 | 532 | NOB | 13.23 | sponge |
| SRR2976122 | sponge metagenome | 5281 | 0 | 693 | NOB | 13.12 | sponge |
| SRR2976113 | sponge metagenome | 3903 | 0 | 496 | NOB | 12.71 | sponge |
| SRR2976095 | sponge metagenome | 778 | 0 | 93 | NOB | 11.95 | sponge |
| SRR3233787 | Petrosia spheroida | 10620 | 1258 | 0 | AOA | 11.85 | sponge |
| SRR2976123 | sponge metagenome | 8723 | 0 | 989 | NOB | 11.34 | sponge |
| SRR2976103 | sponge metagenome | 6240 | 0 | 678 | NOB | 10.87 | sponge |
| SRR2976114 | sponge metagenome | 3966 | 0 | 403 | NOB | 10.16 | sponge |
| SRR2976125 | sponge metagenome | 6169 | 0 | 563 | NOB | 9.13 | sponge |
| SRR2976116 | sponge metagenome | 4402 | 0 | 392 | NOB | 8.91 | sponge |
| SRR2976118 | sponge metagenome | 1585 | 0 | 141 | NOB | 8.90 | sponge |
| SRR2972730 | sponge metagenome | 146930 | 0 | 12790 | NOB | 8.70 | sponge |
| ERR173982 | symbiont metagenome | 7293 | 0 | 606 | NOB | 8.31 | sponge |
| SRR3147492 | Stylissa carteri | 3331 | 0 | 260 | NOB | 7.81 | sponge |
| SRR8359898 | coral reef metagenome | 48450 | 0 | 3765 | NOB | 7.77 | coral |
| SRR1593647 | coral metagenome | 205680 | 14959 | 8 | AOA and NOB | 7.27 | coral |
| SRR2976115 | sponge metagenome | 7541 | 0 | 512 | NOB | 6.79 | sponge |
| SRR2976106 | sponge metagenome | 6015 | 0 | 406 | NOB | 6.75 | sponge |
| SRR7598649 | marine metagenome | 20528 | 0 | 1364 | NOB | 6.64 | sponge |
| SRR7036620 | marine metagenome | 20530 | 0 | 1364 | NOB | 6.64 | sponge |
| SRR7598650 | marine metagenome | 15680 | 0 | 1039 | NOB | 6.63 | sponge |
| SRR7036621 | marine metagenome | 15707 | 0 | 1022 | NOB | 6.51 | sponge |
| ERR2073760 | marine plankton metagenome | 2436996 | 157328 | 0 | AOA | 6.46 | sponge |
| SRR7598651 | marine metagenome | 18568 | 0 | 1173 | NOB | 6.32 | sponge |
| SRR7036622 | marine metagenome | 18577 | 0 | 1173 | NOB | 6.31 | sponge |
| SRR7598648 | marine metagenome | 17090 | 0 | 1076 | NOB | 6.30 | sponge |
| SRR7036619 | marine metagenome | 17097 | 0 | 1076 | NOB | 6.29 | sponge |
| SRR1593660 | coral metagenome | 228658 | 14363 | 4 | AOA and NOB | 6.28 | coral |
| SRR2976127 | sponge metagenome | 4800 | 0 | 300 | NOB | 6.25 | sponge |
| SRR2976102 | sponge metagenome | 6274 | 0 | 387 | NOB | 6.17 | sponge |
| SRR8359899 | coral reef metagenome | 62162 | 0 | 3668 | NOB | 5.90 | coral |
| SRR1992963 | coral metagenome | 66069 | 3868 | 228 | AOA and NOB | 5.85 | coral |
| SRR7598419 | marine metagenome | 10929 | 0 | 638 | NOB | 5.84 | sponge |
| SRR7598575 | marine metagenome | 10253 | 0 | 589 | NOB | 5.74 | sponge |
| SRR7598532 | marine metagenome | 11494 | 0 | 657 | NOB | 5.72 | sponge |
| SRR7036428 | marine metagenome | 11495 | 0 | 657 | NOB | 5.72 | sponge |
| SRR7036427 | marine metagenome | 20905 | 0 | 1174 | NOB | 5.62 | sponge |
| SRR7598533 | marine metagenome | 20913 | 0 | 1174 | NOB | 5.61 | sponge |
| ERR2073756 | Aplysina cauliformis | 677175 | 37882 | 2113 | AOA and NOB | 5.59 | sponge |
| SRR2976124 | sponge metagenome | 3823 | 0 | 209 | NOB | 5.47 | sponge |
| SRR7036422 | marine metagenome | 10886 | 0 | 594 | NOB | 5.46 | sponge |
| SRR7598705 | marine metagenome | 10886 | 0 | 594 | NOB | 5.46 | sponge |
| SRR7598420 | marine metagenome | 20709 | 0 | 1124 | NOB | 5.43 | sponge |
| SRR7036764 | marine metagenome | 20717 | 0 | 1120 | NOB | 5.41 | sponge |
| SRR1593663 | coral metagenome | 183336 | 9745 | 21 | AOA and NOB | 5.32 | coral |
| ERR173981 | symbiont metagenome | 6490 | 0 | 343 | NOB | 5.29 | sponge |
| SRR7598574 | marine metagenome | 20790 | 0 | 1097 | NOB | 5.28 | sponge |
| SRR2976117 | sponge metagenome | 10335 | 0 | 545 | NOB | 5.27 | sponge |
| SRR8359870 | coral reef metagenome | 79248 | 0 | 4169 | NOB | 5.26 | coral |
| SRR7036421 | marine metagenome | 25803 | 0 | 1326 | NOB | 5.14 | sponge |
| SRR6401606 | marine metagenome | 59724 | 0 | 3069 | NOB | 5.14 | sponge |
| SRR7598706 | marine metagenome | 25830 | 0 | 1326 | NOB | 5.13 | sponge |
| SRR7598579 | marine metagenome | 13441 | 0 | 685 | NOB | 5.10 | sponge |
| SRR7598776 | marine metagenome | 12725 | 0 | 642 | NOB | 5.05 | sponge |
| SRR6401425 | marine metagenome | 62740 | 0 | 3160 | NOB | 5.04 | sponge |
| SRR7598526 | marine metagenome | 19067 | 0 | 958 | NOB | 5.02 | sponge |
| SRR2559795 | sponge metagenome | 16951 | 2 | 849 | AOA and NOB | 5.01 | sponge |
| SRR3741959 | marine metagenome | 25537 | 1256 | 0 | AOA | 4.92 | marine sediment |
| ERR2073757 | Aplysina cauliformis | 2479361 | 121901 | 1324 | AOA and NOB | 4.92 | sponge |
| SRR2559765 | sponge metagenome | 15421 | 0 | 753 | NOB | 4.88 | sponge |
| SRR1593661 | coral metagenome | 94432 | 4512 | 0 | AOA | 4.78 | coral |
| SRR7598775 | marine metagenome | 18186 | 0 | 866 | NOB | 4.76 | sponge |

| | | | | | | | |
|---|---|---|---|---|---|---|---|
| SRR7598333 | marine metagenome | 15120 | 0 | 717 | NOB | 4.74 | sponge |
| SRR3147491 | Stylissa carteri | 1718 | 0 | 81 | NOB | 4.71 | sponge |
| SRR7598578 | marine metagenome | 25448 | 0 | 1180 | NOB | 4.64 | sponge |
| SRR2559771 | sponge metagenome | 49911 | 9 | 2275 | AOA and NOB | 4.56 | sponge |
| SRR8359822 | coral reef metagenome | 37798 | 0 | 1719 | NOB | 4.55 | coral |
| SRR6401609 | marine metagenome | 35492 | 0 | 1614 | NOB | 4.55 | sponge |
| SRR2559767 | sponge metagenome | 16657 | 1 | 755 | AOA and NOB | 4.53 | sponge |
| SRR2559775 | sponge metagenome | 46644 | 0 | 2085 | NOB | 4.47 | sponge |
| SRR7598418 | marine metagenome | 17132 | 0 | 752 | NOB | 4.39 | sponge |
| SRR6401462 | marine metagenome | 42464 | 0 | 1850 | NOB | 4.36 | sponge |
| SRR7598781 | marine metagenome | 18111 | 0 | 788 | NOB | 4.35 | sponge |
| SRR1030316 | sponge metagenome | 7042 | 0 | 304 | NOB | 4.32 | sponge |
| ERR2073758 | marine plankton metagenome | 2513842 | 107933 | 0 | AOA | 4.29 | sponge |
| SRR7598652 | marine metagenome | 16498 | 0 | 707 | NOB | 4.29 | sponge |
| SRR7036623 | marine metagenome | 16515 | 0 | 707 | NOB | 4.28 | sponge |
| SRR7598576 | marine metagenome | 21004 | 0 | 894 | NOB | 4.26 | sponge |
| SRR7598653 | marine metagenome | 21857 | 0 | 929 | NOB | 4.25 | sponge |
| SRR7036624 | marine metagenome | 21860 | 0 | 929 | NOB | 4.25 | sponge |
| SRR1791583 | coral metagenome | 975 | 0 | 41 | NOB | 4.21 | coral |
| SRR7598777 | marine metagenome | 21563 | 0 | 904 | NOB | 4.19 | sponge |
| SRR7598417 | marine metagenome | 12888 | 0 | 538 | NOB | 4.17 | sponge |
| SRR7598778 | marine metagenome | 14179 | 0 | 591 | NOB | 4.17 | sponge |
| SRR7598782 | marine metagenome | 15243 | 0 | 623 | NOB | 4.09 | sponge |
| SRR2976107 | sponge metagenome | 3985 | 0 | 159 | NOB | 3.99 | sponge |
| SRR7598525 | marine metagenome | 15896 | 0 | 626 | NOB | 3.94 | sponge |
| SRR6401424 | marine metagenome | 50679 | 0 | 1987 | NOB | 3.92 | sponge |
| SRR1503484 | sponge metagenome | 41785 | 0 | 1637 | NOB | 3.92 | sponge |
| SRR7036706 | marine metagenome | 10376 | 0 | 406 | NOB | 3.91 | sponge |
| SRR7598577 | marine metagenome | 10383 | 0 | 406 | NOB | 3.91 | sponge |
| SRR2559811 | sponge metagenome | 22080 | 1 | 853 | AOA and NOB | 3.86 | sponge |
| SRR2559805 | sponge metagenome | 17936 | 1 | 691 | AOA and NOB | 3.85 | sponge |
| SRR2976105 | sponge metagenome | 14831 | 0 | 564 | NOB | 3.80 | sponge |
| SRR6401429 | marine metagenome | 147217 | 0 | 5589 | NOB | 3.80 | sponge |
| SRR8359894 | coral reef metagenome | 30043 | 0 | 1138 | NOB | 3.79 | coral |
| SRR7598529 | marine metagenome | 13121 | 0 | 486 | NOB | 3.70 | sponge |
| SRR2559817 | sponge metagenome | 66542 | 1 | 2420 | AOA and NOB | 3.64 | sponge |
| SRR2559789 | sponge metagenome | 40045 | 5 | 1456 | AOA and NOB | 3.64 | sponge |
| SRR7598528 | marine metagenome | 16962 | 0 | 614 | NOB | 3.62 | sponge |
| SRR7036770 | marine metagenome | 16990 | 0 | 614 | NOB | 3.61 | sponge |
| SRR1593655 | coral metagenome | 131533 | 4709 | 4 | AOA and NOB | 3.58 | coral |
| SRR7598524 | marine metagenome | 14325 | 0 | 506 | NOB | 3.53 | sponge |
| SRR7036766 | marine metagenome | 14349 | 0 | 506 | NOB | 3.53 | sponge |
| ERR173983 | symbiont metagenome | 8451 | 0 | 293 | NOB | 3.47 | sponge |
| SRR8359896 | coral reef metagenome | 78128 | 0 | 2708 | NOB | 3.47 | coral |
| SRR7598531 | marine metagenome | 11518 | 0 | 398 | NOB | 3.46 | sponge |
| SRR3204474 | coral metagenome | 89981 | 3105 | 158 | AOA and NOB | 3.45 | coral |
| SRR2559778 | sponge metagenome | 29357 | 0 | 1003 | NOB | 3.42 | sponge |
| SRR8359871 | coral reef metagenome | 48096 | 0 | 1621 | NOB | 3.37 | coral |
| SRR2559781 | sponge metagenome | 48225 | 0 | 1612 | NOB | 3.34 | sponge |
| SRR7598530 | marine metagenome | 19849 | 0 | 663 | NOB | 3.34 | sponge |
| SRR2559802 | sponge metagenome | 8412 | 1 | 277 | AOA and NOB | 3.29 | sponge |
| SRR2976110 | sponge metagenome | 4337 | 0 | 142 | NOB | 3.27 | sponge |
| SRR6401615 | marine metagenome | 40926 | 0 | 1337 | NOB | 3.27 | sponge |
| SRR2559806 | sponge metagenome | 34742 | 1 | 1134 | AOA and NOB | 3.26 | sponge |
| SRR2559818 | sponge metagenome | 27023 | 0 | 871 | NOB | 3.22 | sponge |
| SRR6401428 | marine metagenome | 56570 | 0 | 1814 | NOB | 3.21 | sponge |
| SRR2559824 | sponge metagenome | 14016 | 0 | 448 | NOB | 3.20 | sponge |
| SRR2559820 | sponge metagenome | 11866 | 1 | 375 | AOA and NOB | 3.16 | sponge |
| SRR1561181 | marine metagenome | 5866 | 0 | 184 | NOB | 3.14 | seawater |
| SRR1557141 | marine metagenome | 5891 | 0 | 184 | NOB | 3.12 | seawater |
| SRR2559773 | sponge metagenome | 21438 | 0 | 666 | NOB | 3.11 | sponge |
| SRR2976104 | sponge metagenome | 1267 | 0 | 39 | NOB | 3.08 | sponge |
| SRR709467 | sponge metagenome | 43846 | 0 | 1347 | NOB | 3.07 | sponge |
| SRR2559797 | sponge metagenome | 15260 | 0 | 466 | NOB | 3.05 | sponge |
| SRR869518 | sponge metagenome | 14033 | 0 | 428 | NOB | 3.05 | sponge |
| SRR2559785 | sponge metagenome | 34932 | 7 | 1042 | AOA and NOB | 2.98 | sponge |
| SRR2559815 | sponge metagenome | 21112 | 2 | 625 | AOA and NOB | 2.96 | sponge |
| SRR2559768 | sponge metagenome | 14065 | 1 | 411 | AOA and NOB | 2.92 | sponge |
| SRR2559816 | sponge metagenome | 28575 | 4 | 821 | AOA and NOB | 2.87 | sponge |
| SRR1560270 | marine metagenome | 5621 | 0 | 161 | NOB | 2.86 | seawater |
| SRR2559819 | sponge metagenome | 5519 | 2 | 158 | AOA and NOB | 2.86 | sponge |
| SRR7178011 | sponge metagenome | 135617 | 0 | 3871 | NOB | 2.85 | sponge |
| SRR2559791 | sponge metagenome | 20329 | 3 | 577 | AOA and NOB | 2.84 | sponge |
| SRR1593657 | coral metagenome | 88065 | 2493 | 0 | AOA | 2.83 | coral |
| SRR8359873 | coral reef metagenome | 50253 | 0 | 1416 | NOB | 2.82 | coral |
| ERR2073755 | Aplysina cauliformis | 621338 | 17305 | 748 | AOA and NOB | 2.79 | sponge |
| SRR2559799 | sponge metagenome | 24687 | 4 | 686 | AOA and NOB | 2.78 | sponge |
| SRR709466 | sponge metagenome | 53632 | 0 | 1487 | NOB | 2.77 | sponge |
| SRR3233788 | Petrosia spheroida | 6315 | 174 | 0 | AOA | 2.76 | sponge |
| SRR3204413 | coral metagenome | 37458 | 1032 | 40 | AOA and NOB | 2.76 | coral |
| SRR2559796 | sponge metagenome | 30477 | 2 | 838 | AOA and NOB | 2.75 | sponge |
| SRR1593658 | coral metagenome | 188297 | 5111 | 0 | AOA | 2.71 | coral |
| SRR2559766 | sponge metagenome | 56992 | 8 | 1531 | AOA and NOB | 2.69 | sponge |
| SRR2559813 | sponge metagenome | 24942 | 1 | 670 | AOA and NOB | 2.69 | sponge |
| SRR869546 | sponge metagenome | 12104 | 0 | 322 | NOB | 2.66 | sponge |
| SRR2559779 | sponge metagenome | 10136 | 0 | 269 | NOB | 2.65 | sponge |
| SRR8359841 | coral reef metagenome | 55546 | 0 | 1462 | NOB | 2.63 | coral |
| SRR8359877 | coral reef metagenome | 55778 | 0 | 1468 | NOB | 2.63 | coral |
| SRR7036618 | marine metagenome | 21215 | 0 | 558 | NOB | 2.63 | sponge |
| SRR2559786 | sponge metagenome | 23528 | 6 | 618 | AOA and NOB | 2.63 | sponge |
| SRR7598647 | marine metagenome | 21220 | 0 | 557 | NOB | 2.62 | sponge |
| SRR2559826 | sponge metagenome | 49810 | 1 | 1292 | AOA and NOB | 2.59 | sponge |
| SRR6401607 | marine metagenome | 64509 | 0 | 1668 | NOB | 2.59 | sponge |
| SRR2559764 | sponge metagenome | 42330 | 2 | 1094 | AOA and NOB | 2.58 | sponge |
| SRR2559772 | sponge metagenome | 25609 | 0 | 660 | NOB | 2.58 | sponge |
| SRR709465 | sponge metagenome | 54664 | 0 | 1399 | NOB | 2.56 | sponge |
| ERR173984 | symbiont metagenome | 7522 | 0 | 192 | NOB | 2.55 | sponge |

| | | | | | | | |
|---|---|---|---|---|---|---|---|
| SRR2559809 | sponge metagenome | 17272 | 5 | 440 | AOA and NOB | 2.55 | sponge |
| SRR7598646 | marine metagenome | 16508 | 0 | 419 | NOB | 2.54 | sponge |
| SRR7036617 | marine metagenome | 16509 | 0 | 419 | NOB | 2.54 | sponge |
| SRR1565480 | sponge metagenome | 21364 | 0 | 540 | NOB | 2.53 | sponge |
| SRR4244557 | marine metagenome | 70928 | 0 | 1788 | NOB | 2.52 | sponge |
| SRR2559800 | sponge metagenome | 13151 | 0 | 330 | NOB | 2.51 | sponge |
| SRR6401671 | marine metagenome | 2139 | 0 | 53 | NOB | 2.48 | sponge |
| SRR869519 | sponge metagenome | 13993 | 0 | 346 | NOB | 2.47 | sponge |
| SRR2559769 | sponge metagenome | 35658 | 4 | 874 | AOA and NOB | 2.45 | sponge |
| SRR8359874 | coral reef metagenome | 11896 | 0 | 287 | NOB | 2.41 | coral |
| SRR2559782 | sponge metagenome | 6597 | 0 | 157 | NOB | 2.38 | sponge |
| SRR869515 | sponge metagenome | 13458 | 0 | 320 | NOB | 2.38 | sponge |
| SRR8359862 | coral reef metagenome | 30010 | 0 | 710 | NOB | 2.37 | coral |
| SRR7598675 | marine metagenome | 12036 | 0 | 283 | NOB | 2.35 | sponge |
| SRR2559794 | sponge metagenome | 21738 | 0 | 509 | NOB | 2.34 | sponge |
| SRR3204414 | coral metagenome | 47933 | 1119 | 51 | AOA and NOB | 2.33 | coral |
| SRR8359876 | coral reef metagenome | 98444 | 0 | 2298 | NOB | 2.33 | coral |
| SRR7598415 | marine metagenome | 11492 | 0 | 267 | NOB | 2.32 | sponge |
| SRR2559812 | sponge metagenome | 12278 | 1 | 285 | AOA and NOB | 2.32 | sponge |
| SRR8359869 | coral reef metagenome | 28045 | 0 | 650 | NOB | 2.32 | coral |
| SRR6401477 | marine metagenome | 62882 | 0 | 1455 | NOB | 2.31 | sponge |
| SRR869526 | sponge metagenome | 10563 | 0 | 244 | NOB | 2.31 | sponge |
| SRR7598773 | marine metagenome | 27152 | 0 | 627 | NOB | 2.31 | sponge |
| SRR869551 | sponge metagenome | 13823 | 0 | 319 | NOB | 2.31 | sponge |
| SRR8359887 | coral reef metagenome | 68418 | 0 | 1559 | NOB | 2.28 | coral |
| SRR2559822 | sponge metagenome | 6764 | 0 | 154 | NOB | 2.28 | sponge |
| SRR7598416 | marine metagenome | 28261 | 0 | 641 | NOB | 2.27 | sponge |
| ERR951520 | metagenome | 4671 | 0 | 104 | NOB | 2.23 | coral |
| SRR7598411 | marine metagenome | 11352 | 0 | 247 | NOB | 2.18 | sponge |
| SRR7598412 | marine metagenome | 27487 | 0 | 593 | NOB | 2.16 | sponge |
| SRR869521 | sponge metagenome | 12764 | 0 | 275 | NOB | 2.15 | sponge |
| SRR1992871 | coral metagenome | 46997 | 1011 | 630 | AOA and NOB | 2.15 | coral |
| SRR7598674 | marine metagenome | 19063 | 0 | 410 | NOB | 2.15 | sponge |
| SRR6401610 | marine metagenome | 55432 | 0 | 1182 | NOB | 2.13 | sponge |
| SRR8359781 | coral reef metagenome | 46607 | 0 | 991 | NOB | 2.13 | coral |
| SRR7598580 | marine metagenome | 20541 | 0 | 432 | NOB | 2.10 | sponge |
| SRR7598774 | marine metagenome | 10635 | 0 | 223 | NOB | 2.10 | sponge |
| SRR2559810 | sponge metagenome | 24797 | 2 | 501 | AOA and NOB | 2.02 | sponge |
| SRR2559787 | sponge metagenome | 43255 | 1 | 858 | AOA and NOB | 1.98 | sponge |
| SRR869517 | sponge metagenome | 12229 | 0 | 242 | NOB | 1.98 | sponge |
| SRR2559823 | sponge metagenome | 21928 | 2 | 430 | AOA and NOB | 1.96 | sponge |
| SRR1565479 | sponge metagenome | 18465 | 0 | 362 | NOB | 1.96 | sponge |
| SRR8529235 | marine metagenome | 462324 | 9025 | 0 | AOA | 1.95 | seawater |
| SRR7598685 | marine metagenome | 10862 | 0 | 211 | NOB | 1.94 | sponge |
| SRR7178010 | sponge metagenome | 199530 | 1 | 3814 | AOA and NOB | 1.91 | sponge |
| SRR2559827 | sponge metagenome | 22355 | 5 | 427 | AOA and NOB | 1.91 | sponge |
| SRR2559798 | sponge metagenome | 44143 | 4 | 842 | AOA and NOB | 1.91 | sponge |
| SRR2559770 | sponge metagenome | 51188 | 4 | 976 | AOA and NOB | 1.91 | sponge |
| SRR768462 | marine metagenome | 3432 | 0 | 65 | NOB | 1.89 | sponge |
| SRR869528 | sponge metagenome | 9385 | 0 | 177 | NOB | 1.89 | sponge |
| SRR869516 | sponge metagenome | 12927 | 0 | 243 | NOB | 1.88 | sponge |
| ERR951517 | metagenome | 6302 | 0 | 118 | NOB | 1.87 | coral |
| SRR8359897 | coral reef metagenome | 85033 | 0 | 1592 | NOB | 1.87 | coral |
| SRR6401431 | marine metagenome | 53431 | 0 | 995 | NOB | 1.86 | sponge |
| SRR2748655 | sponge metagenome | 1156584 | 21511 | 5668 | AOA and NOB | 1.86 | sponge |
| SRR8359823 | coral reef metagenome | 54219 | 0 | 1006 | NOB | 1.86 | coral |
| SRR7591412 | coral metagenome | 4039 | 74 | 4 | AOA and NOB | 1.83 | coral |
| SRR2559807 | sponge metagenome | 9924 | 0 | 179 | NOB | 1.80 | sponge |
| SRR869520 | sponge metagenome | 15408 | 0 | 276 | NOB | 1.79 | sponge |
| SRR8359863 | coral reef metagenome | 52225 | 0 | 935 | NOB | 1.79 | coral |
| SRR7591434 | coral metagenome | 2370 | 0 | 42 | NOB | 1.77 | coral |
| SRR7591404 | coral metagenome | 567 | 3 | 10 | AOA and NOB | 1.76 | coral |
| SRR1039445 | sponge metagenome | 8637 | 0 | 151 | NOB | 1.75 | sponge |
| SRR869523 | sponge metagenome | 12257 | 0 | 212 | NOB | 1.73 | sponge |
| SRR7598779 | marine metagenome | 21236 | 0 | 363 | NOB | 1.71 | sponge |
| SRR4244552 | marine metagenome | 52895 | 111 | 902 | AOA and NOB | 1.71 | sponge |
| SRR2559821 | sponge metagenome | 7279 | 2 | 124 | AOA and NOB | 1.70 | sponge |
| SRR869525 | sponge metagenome | 9775 | 0 | 166 | NOB | 1.70 | sponge |
| SRR2970442 | sponge metagenome | 3425 | 0 | 58 | NOB | 1.69 | sponge |
| SRR2976101 | sponge metagenome | 3425 | 0 | 58 | NOB | 1.69 | sponge |
| SRR869556 | sponge metagenome | 18763 | 0 | 302 | NOB | 1.61 | sponge |
| SRR2559774 | sponge metagenome | 5581 | 0 | 89 | NOB | 1.59 | sponge |
| SRR2559784 | sponge metagenome | 9843 | 0 | 156 | NOB | 1.58 | sponge |
| SRR869543 | sponge metagenome | 10396 | 0 | 163 | NOB | 1.57 | sponge |
| SRR1565481 | sponge metagenome | 14610 | 0 | 229 | NOB | 1.57 | sponge |
| SRR7598780 | marine metagenome | 14135 | 0 | 217 | NOB | 1.54 | sponge |
| SRR2559788 | sponge metagenome | 20292 | 0 | 309 | NOB | 1.52 | sponge |
| SRR869544 | sponge metagenome | 11568 | 0 | 176 | NOB | 1.52 | sponge |
| SRR1565484 | sponge metagenome | 17513 | 6 | 266 | AOA and NOB | 1.52 | sponge |
| SRR869522 | sponge metagenome | 12174 | 0 | 182 | NOB | 1.49 | sponge |
| SRR7178004 | sponge metagenome | 143226 | 0 | 2128 | NOB | 1.49 | sponge |
| SRR6401430 | marine metagenome | 56647 | 0 | 837 | NOB | 1.48 | sponge |
| SRR2559780 | sponge metagenome | 61077 | 4 | 890 | AOA and NOB | 1.46 | sponge |
| SRR2559804 | sponge metagenome | 55880 | 3 | 814 | AOA and NOB | 1.46 | sponge |
| SRR1040519 | sponge metagenome | 8016 | 0 | 116 | NOB | 1.45 | sponge |
| SRR869548 | sponge metagenome | 12244 | 0 | 177 | NOB | 1.45 | sponge |
| SRR871431 | sponge metagenome | 3539 | 0 | 51 | NOB | 1.44 | sponge |
| SRR869549 | sponge metagenome | 9399 | 0 | 134 | NOB | 1.43 | sponge |
| SRR8285244 | algae metagenome | 90727 | 978 | 1293 | AOA and NOB | 1.43 | algae |
| SRR4244551 | marine metagenome | 55276 | 489 | 781 | AOA and NOB | 1.41 | sponge |
| SRR2559803 | sponge metagenome | 8717 | 0 | 123 | NOB | 1.41 | sponge |
| SRR6401662 | marine metagenome | 4477 | 0 | 63 | NOB | 1.41 | sponge |
| SRR2559825 | sponge metagenome | 28905 | 2 | 394 | AOA and NOB | 1.36 | sponge |
| SRR869545 | sponge metagenome | 11546 | 0 | 156 | NOB | 1.35 | sponge |
| SRR1992871 | coral metagenome | 46997 | 1011 | 630 | AOA and NOB | 1.34 | coral |
| SRR869530 | sponge metagenome | 11077 | 0 | 146 | NOB | 1.32 | sponge |
| SRR2559793 | sponge metagenome | 5768 | 1 | 76 | AOA and NOB | 1.32 | sponge |

| | | | | | | | |
|---|---|---|---|---|---|---|---|
| SRR6401611 | marine metagenome | 56222 | 0 | 740 | NOB | 1.32 | sponge |
| SRR6401426 | marine metagenome | 57600 | 0 | 756 | NOB | 1.31 | sponge |
| SRR1593656 | coral metagenome | 132916 | 1733 | 0 | AOA | 1.30 | coral |
| SRR2559801 | sponge metagenome | 24940 | 1 | 325 | AOA and NOB | 1.30 | sponge |
| SRR2559777 | sponge metagenome | 31218 | 1 | 406 | AOA and NOB | 1.30 | sponge |
| SRR709468 | sponge metagenome | 41038 | 0 | 532 | NOB | 1.30 | sponge |
| SRR7178023 | sponge metagenome | 200300 | 0 | 2596 | NOB | 1.30 | sponge |
| SRR869547 | sponge metagenome | 10431 | 0 | 131 | NOB | 1.26 | sponge |
| SRR709470 | sponge metagenome | 34480 | 0 | 430 | NOB | 1.25 | sponge |
| SRR709469 | sponge metagenome | 43749 | 0 | 529 | NOB | 1.21 | sponge |
| SRR768460 | marine metagenome | 6217 | 0 | 75 | NOB | 1.21 | sponge |
| SRR5469079 | sponge metagenome | 46801 | 0 | 558 | NOB | 1.19 | sponge |
| SRR6457253 | marine metagenome | 34424 | 409 | 107 | AOA and NOB | 1.19 | seawater |
| SRR7178007 | sponge metagenome | 150315 | 0 | 1760 | NOB | 1.17 | sponge |
| SRR6081891 | sediment metagenome | 17286 | 0 | 198 | NOB | 1.15 | marine sediment |
| SRR871434 | sponge metagenome | 4305 | 0 | 49 | NOB | 1.14 | sponge |
| SRR1565485 | sponge metagenome | 18463 | 0 | 210 | NOB | 1.14 | sponge |
| SRR869554 | sponge metagenome | 12743 | 0 | 143 | NOB | 1.12 | sponge |
| SRR6401727 | marine metagenome | 42973 | 0 | 471 | NOB | 1.10 | coral |
| SRR4244546 | marine metagenome | 75009 | 147 | 821 | AOA and NOB | 1.09 | sponge |
| SRR3197004 | coral metagenome | 51008 | 0 | 550 | NOB | 1.08 | coral |
| SRR8285244 | algae metagenome | 90727 | 978 | 1293 | AOA and NOB | 1.08 | algae |
| SRR1565477 | sponge metagenome | 14024 | 0 | 150 | NOB | 1.07 | sponge |
| SRR5437924 | marine metagenome | 8081 | 76 | 86 | AOA and NOB | 1.06 | coral |
| SRR869550 | sponge metagenome | 11012 | 0 | 114 | NOB | 1.04 | sponge |
| SRR3233455 | Petrosia spheroida | 2718 | 0 | 28 | NOB | 1.03 | sponge |
| SRR8359787 | coral reef metagenome | 62677 | 0 | 644 | NOB | 1.03 | coral |
| DRR036737 | uncultured bacterium | 16330 | 0 | 165 | NOB | 1.01 | sponge |
| SRR1393867 | sea squirt metagenome | 15421 | 0 | 155 | NOB | 1.01 | other marine invertebrates |
| SRR7036419 | marine metagenome | 17399 | 0 | 174 | NOB | 1.00 | sponge |
| SRR7598535 | marine metagenome | 17418 | 0 | 174 | NOB | 1.00 | sponge |
| SRR8359766 | coral reef metagenome | 24122 | 0 | 239 | NOB | 0.99 | coral |
| SRR3204470 | coral metagenome | 42433 | 418 | 188 | AOA and NOB | 0.99 | coral |
| SRR2559808 | sponge metagenome | 1429 | 0 | 14 | NOB | 0.98 | sponge |
| SRR8359814 | coral reef metagenome | 33086 | 0 | 318 | NOB | 0.96 | coral |
| SRR7598534 | marine metagenome | 12014 | 0 | 115 | NOB | 0.96 | sponge |
| SRR7036420 | marine metagenome | 12026 | 0 | 115 | NOB | 0.96 | sponge |
| SRR5437924 | marine metagenome | 8081 | 76 | 86 | AOA and NOB | 0.94 | coral |
| SRR1565482 | sponge metagenome | 17054 | 0 | 160 | NOB | 0.94 | sponge |
| SRR5469078 | sponge metagenome | 69041 | 0 | 637 | NOB | 0.92 | sponge |
| SRR8359906 | coral reef metagenome | 115659 | 0 | 1052 | NOB | 0.91 | coral |
| SRR6401663 | marine metagenome | 1323 | 0 | 12 | NOB | 0.91 | coral |
| SRR4471382 | marine metagenome | 3545 | 2 | 32 | AOA and NOB | 0.90 | coral |
| SRR7591405 | coral metagenome | 5433 | 49 | 42 | AOA and NOB | 0.90 | coral |
| SRR1565483 | sponge metagenome | 17172 | 0 | 152 | NOB | 0.89 | sponge |
| SRR4244551 | marine metagenome | 55276 | 489 | 781 | AOA and NOB | 0.88 | sponge |
| SRR7598413 | marine metagenome | 10445 | 0 | 91 | NOB | 0.87 | sponge |
| SRR2983135 | marine sediment metagenome | 1735 | 0 | 15 | NOB | 0.86 | marine sediment |
| SRR2559783 | sponge metagenome | 23465 | 0 | 202 | NOB | 0.86 | sponge |
| SRR4244550 | marine metagenome | 58447 | 165 | 501 | AOA and NOB | 0.86 | sponge |
| DRR036738 | uncultured bacterium | 24556 | 0 | 210 | NOB | 0.86 | sponge |
| SRR768463 | marine metagenome | 4904 | 0 | 41 | NOB | 0.84 | sponge |
| SRR869553 | sponge metagenome | 11229 | 0 | 93 | NOB | 0.83 | sponge |
| SRR3233442 | Dragmacidon coccineum | 4347 | 0 | 36 | NOB | 0.83 | sponge |
| SRR7598414 | marine metagenome | 23676 | 0 | 196 | NOB | 0.83 | sponge |
| SRR6081896 | sediment metagenome | 15539 | 0 | 128 | NOB | 0.82 | marine sediment |
| SRR869558 | sponge metagenome | 12049 | 0 | 99 | NOB | 0.82 | sponge |
| SRR8359900 | coral reef metagenome | 34812 | 0 | 281 | NOB | 0.81 | coral |
| SRR5469076 | sponge metagenome | 64223 | 0 | 507 | NOB | 0.79 | sponge |
| SRR3233443 | Dragmacidon coccineum | 6684 | 0 | 52 | NOB | 0.78 | sponge |
| SRR8359875 | coral reef metagenome | 74127 | 0 | 574 | NOB | 0.77 | coral |
| SRR7591405 | coral metagenome | 5433 | 49 | 42 | AOA and NOB | 0.77 | coral |
| SRR7598264 | metagenome | 20818 | 0 | 160 | NOB | 0.77 | coral |
| SRR8359861 | coral reef metagenome | 26510 | 0 | 201 | NOB | 0.76 | coral |
| SRR5437923 | marine metagenome | 60477 | 11 | 448 | AOA and NOB | 0.74 | coral |
| SRR768461 | marine metagenome | 5182 | 0 | 38 | NOB | 0.73 | sponge |
| SRR2559814 | sponge metagenome | 7318 | 0 | 53 | NOB | 0.72 | sponge |
| SRR5469084 | sponge metagenome | 59362 | 0 | 428 | NOB | 0.72 | sponge |
| SRR5469075 | sponge metagenome | 52668 | 0 | 379 | NOB | 0.72 | sponge |
| SRR5469081 | sponge metagenome | 72251 | 0 | 514 | NOB | 0.71 | sponge |
| SRR8359769 | coral reef metagenome | 12543 | 0 | 88 | NOB | 0.70 | coral |
| SRR3233470 | Xestospongia testudinaria | 5047 | 0 | 35 | NOB | 0.69 | sponge |
| SRR7178008 | sponge metagenome | 110047 | 0 | 757 | NOB | 0.69 | sponge |
| SRR869524 | sponge metagenome | 11372 | 0 | 76 | NOB | 0.67 | sponge |
| SRR5469073 | sponge metagenome | 56962 | 0 | 378 | NOB | 0.66 | sponge |
| SRR1565478 | sponge metagenome | 19986 | 1 | 130 | AOA and NOB | 0.65 | sponge |
| SRR869527 | sponge metagenome | 11207 | 0 | 71 | NOB | 0.63 | sponge |
| SRR5469077 | sponge metagenome | 78151 | 0 | 489 | NOB | 0.63 | sponge |
| SRR8661884 | coral metagenome | 32944 | 0 | 204 | NOB | 0.62 | coral |
| SRR869555 | sponge metagenome | 10066 | 0 | 61 | NOB | 0.61 | sponge |
| SRR2559790 | sponge metagenome | 28663 | 0 | 173 | NOB | 0.60 | sponge |
| SRR3741960 | marine metagenome | 63110 | 377 | 0 | AOA | 0.60 | marine sediment |
| SRR7356965 | coral metagenome | 19813 | 0 | 118 | NOB | 0.60 | coral |
| SRR5469082 | sponge metagenome | 51300 | 0 | 300 | NOB | 0.58 | sponge |
| SRR2559792 | sponge metagenome | 16058 | 1 | 92 | AOA and NOB | 0.57 | sponge |
| ERR951516 | metagenome | 4197 | 0 | 24 | NOB | 0.57 | coral |
| SRR8359821 | coral reef metagenome | 51420 | 0 | 290 | NOB | 0.56 | coral |
| SRR1565475 | sponge metagenome | 19549 | 2 | 110 | AOA and NOB | 0.56 | sponge |
| SRR4244568 | marine metagenome | 60430 | 90 | 329 | AOA and NOB | 0.54 | sponge |
| SRR1039451 | sponge metagenome | 7066 | 0 | 38 | NOB | 0.54 | sponge |
| SRR8640071 | algae metagenome | 70182 | 0 | 376 | NOB | 0.54 | algae |
| SRR651962 | sponge metagenome | 48652 | 0 | 260 | NOB | 0.53 | sponge |
| SRR3204449 | coral metagenome | 75161 | 399 | 33 | AOA and NOB | 0.53 | coral |
| SRR7591404 | coral metagenome | 567 | 3 | 10 | AOA and NOB | 0.53 | coral |
| SRR8359820 | coral reef metagenome | 53131 | 0 | 280 | NOB | 0.53 | coral |
| SRR8359865 | coral reef metagenome | 16681 | 0 | 87 | NOB | 0.52 | coral |
| SRR7034713 | marine metagenome | 22091 | 0 | 114 | NOB | 0.52 | sponge |

| | | | | | | | |
|---|---|---|---|---|---|---|---|
| SRR7598752 | marine metagenome | 22096 | 0 | 114 | NOB | 0.52 | sponge |
| SRR7591410 | coral metagenome | 23453 | 3307 | 121 | AOA and NOB | 0.52 | coral |
| SRR3233461 | Stylissa carteri | 13027 | 0 | 67 | NOB | 0.51 | sponge |
| SRR3233456 | Petrosia spheroida | 6340 | 0 | 32 | NOB | 0.50 | sponge |
| SRR1560096 | marine metagenome | 11293 | 0 | 56 | NOB | 0.50 | seawater |
| SRR2748655 | sponge metagenome | 1156584 | 21511 | 5668 | AOA and NOB | 0.49 | sponge |
| SRR8359827 | coral reef metagenome | 129350 | 0 | 614 | NOB | 0.47 | coral |
| SRR3233454 | Petrosia spheroida | 2875 | 0 | 13 | NOB | 0.45 | sponge |
| SRR3204470 | coral metagenome | 42433 | 418 | 188 | AOA and NOB | 0.44 | coral |
| SRR8359885 | coral reef metagenome | 30215 | 0 | 133 | NOB | 0.44 | coral |
| SRR413285 | marine metagenome | 62912 | 0 | 273 | NOB | 0.43 | coral |
| SRR2532851 | coral metagenome | 3244 | 0 | 14 | NOB | 0.43 | coral |
| SRR6401421 | marine metagenome | 137173 | 0 | 586 | NOB | 0.43 | sponge |
| SRR2983129 | marine sediment metagenome | 2372 | 0 | 10 | NOB | 0.42 | marine sediment |
| SRR8639894 | algae metagenome | 102504 | 0 | 431 | NOB | 0.42 | algae |
| SRR1039475 | sponge metagenome | 5731 | 0 | 24 | NOB | 0.42 | sponge |
| DRR036735 | uncultured bacterium | 20233 | 0 | 84 | NOB | 0.42 | sponge |
| SRR3742007 | marine metagenome | 42894 | 175 | 0 | AOA | 0.41 | marine sediment |
| SRR3233466 | Stylissa massa | 2723 | 0 | 11 | NOB | 0.40 | sponge |
| SRR5469083 | sponge metagenome | 41782 | 0 | 166 | NOB | 0.40 | sponge |
| SRR1039449 | sponge metagenome | 4307 | 0 | 17 | NOB | 0.39 | sponge |
| SRR2983130 | marine sediment metagenome | 2284 | 0 | 9 | NOB | 0.39 | marine sediment |
| SRR3233785 | Hyrtios erectus | 17835 | 70 | 0 | AOA | 0.39 | sponge |
| SRR8272691 | sponge metagenome | 34706 | 0 | 131 | NOB | 0.38 | seagrass |
| SRR3233460 | Stylissa carteri | 8234 | 0 | 31 | NOB | 0.38 | sponge |
| SRR7034366 | marine metagenome | 29700 | 0 | 110 | NOB | 0.37 | sponge |
| SRR7598751 | marine metagenome | 29701 | 0 | 110 | NOB | 0.37 | sponge |
| SRR2087834 | plant metagenome | 132388 | 485 | 23 | AOA and NOB | 0.37 | plant |
| SRR7178028 | sponge metagenome | 170210 | 0 | 617 | NOB | 0.36 | sponge |
| SRR3147482 | Aaptos suberitoides | 1656 | 0 | 6 | NOB | 0.36 | sponge |
| SRR869552 | sponge metagenome | 10571 | 0 | 38 | NOB | 0.36 | sponge |
| SRR8359788 | coral reef metagenome | 46466 | 0 | 167 | NOB | 0.36 | coral |
| SRR1039474 | sponge metagenome | 2531 | 0 | 9 | NOB | 0.36 | sponge |
| SRR8581535 | annelid metagenome | 101324 | 0 | 360 | NOB | 0.36 | other marine invertebrates |
| SRR5469074 | sponge metagenome | 78814 | 0 | 280 | NOB | 0.36 | sponge |
| SRR8640063 | algae metagenome | 67537 | 0 | 239 | NOB | 0.35 | algae |
| SRR8639947 | algae metagenome | 93596 | 0 | 324 | NOB | 0.35 | algae |
| SRR869557 | sponge metagenome | 12146 | 0 | 42 | NOB | 0.35 | sponge |
| SRR1992963 | coral metagenome | 66069 | 3868 | 228 | AOA and NOB | 0.35 | coral |
| SRR7178027 | sponge metagenome | 152273 | 0 | 525 | NOB | 0.34 | sponge |
| SRR7178024 | sponge metagenome | 177861 | 0 | 610 | NOB | 0.34 | sponge |
| SRR4471404 | marine metagenome | 588 | 0 | 2 | NOB | 0.34 | coral |
| SRR6081898 | sediment metagenome | 15364 | 0 | 52 | NOB | 0.34 | marine sediment |
| SRR1992937 | coral metagenome | 25275 | 0 | 83 | NOB | 0.33 | coral |
| SRR8359881 | coral reef metagenome | 30187 | 0 | 99 | NOB | 0.33 | coral |
| SRR1562083 | sponge metagenome | 23791 | 0 | 78 | NOB | 0.33 | sponge |
| SRR8359810 | coral reef metagenome | 22102 | 0 | 71 | NOB | 0.32 | coral |
| ERR2073756 | Aplysina cauliformis | 677175 | 37882 | 2113 | AOA and NOB | 0.31 | sponge |
| SRR6457253 | marine metagenome | 34424 | 409 | 107 | AOA and NOB | 0.31 | seawater |
| SRR1237906 | marine sediment metagenome | 1942 | 0 | 6 | NOB | 0.31 | marine sediment |
| SRR8359812 | coral reef metagenome | 16275 | 0 | 50 | NOB | 0.31 | coral |
| SRR7598486 | marine metagenome | 29666 | 0 | 91 | NOB | 0.31 | sponge |
| SRR7034513 | marine metagenome | 29667 | 0 | 91 | NOB | 0.31 | sponge |
| SRR6081897 | sediment metagenome | 27991 | 0 | 81 | NOB | 0.29 | marine sediment |
| SRR8359868 | coral reef metagenome | 76381 | 0 | 219 | NOB | 0.29 | coral |
| SRR8639897 | algae metagenome | 95650 | 0 | 274 | NOB | 0.29 | algae |
| SRR4471359 | marine metagenome | 7682 | 22 | 4 | AOA and NOB | 0.29 | coral |
| SRR4244550 | marine metagenome | 58447 | 165 | 501 | AOA and NOB | 0.28 | sponge |
| SRR8111657 | mollusc metagenome | 12409 | 8 | 35 | AOA and NOB | 0.28 | other marine invertebrates |
| SRR7598487 | marine metagenome | 24687 | 0 | 68 | NOB | 0.28 | sponge |
| SRR7034522 | marine metagenome | 24692 | 0 | 68 | NOB | 0.28 | sponge |
| SRR7036442 | marine metagenome | 6274 | 0 | 17 | NOB | 0.27 | coral |
| SRR2983133 | marine sediment metagenome | 2215 | 0 | 6 | NOB | 0.27 | marine sediment |
| SRR7598669 | marine metagenome | 6284 | 0 | 17 | NOB | 0.27 | coral |
| SRR8359825 | coral reef metagenome | 79121 | 0 | 214 | NOB | 0.27 | coral |
| SRR2559776 | sponge metagenome | 5208 | 0 | 14 | NOB | 0.27 | sponge |
| SRR8359826 | coral reef metagenome | 73885 | 0 | 197 | NOB | 0.27 | coral |
| SRR8243376 | coral reef metagenome | 34903 | 0 | 93 | NOB | 0.27 | coral |
| SRR2983128 | marine sediment metagenome | 2734 | 0 | 7 | NOB | 0.26 | marine sediment |
| SRR6457116 | marine metagenome | 54853 | 139 | 0 | AOA | 0.25 | seawater |
| SRR7034377 | marine metagenome | 43968 | 0 | 110 | NOB | 0.25 | sponge |
| SRR7598618 | marine metagenome | 43978 | 0 | 110 | NOB | 0.25 | sponge |
| SRR8359828 | coral reef metagenome | 47745 | 0 | 119 | NOB | 0.25 | coral |
| ERR951503 | metagenome | 4871 | 0 | 12 | NOB | 0.25 | coral |
| SRR3204478 | coral metagenome | 78006 | 191 | 11 | AOA and NOB | 0.24 | coral |
| SRR8359893 | coral reef metagenome | 49777 | 0 | 119 | NOB | 0.24 | coral |
| SRR8639891 | algae metagenome | 11439 | 0 | 27 | NOB | 0.24 | algae |
| SRR1992934 | coral metagenome | 26984 | 63 | 0 | AOA | 0.23 | coral |
| SRR2983134 | marine sediment metagenome | 2238 | 0 | 5 | NOB | 0.22 | marine sediment |
| SRR3147483 | Aaptos suberitoides | 1374 | 0 | 3 | NOB | 0.22 | sponge |
| SRR4244549 | marine metagenome | 89268 | 13 | 192 | AOA and NOB | 0.22 | sponge |
| SRR8359904 | coral reef metagenome | 98194 | 0 | 211 | NOB | 0.21 | coral |
| SRR1565476 | sponge metagenome | 30621 | 0 | 65 | NOB | 0.21 | sponge |
| SRR4244552 | marine metagenome | 52895 | 111 | 902 | AOA and NOB | 0.21 | sponge |
| SRR867624 | marine metagenome | 1909 | 4 | 0 | AOA | 0.21 | seawater |
| SRR1039471 | sponge metagenome | 5290 | 0 | 11 | NOB | 0.21 | sponge |
| SRR2087835 | plant metagenome | 196273 | 406 | 14 | AOA and NOB | 0.21 | plant |
| SRR7178026 | sponge metagenome | 208721 | 0 | 428 | NOB | 0.21 | sponge |
| SRR4471364 | marine metagenome | 1956 | 0 | 4 | NOB | 0.20 | coral |
| SRR3204477 | coral metagenome | 59562 | 120 | 17 | AOA and NOB | 0.20 | coral |
| SRR7596251 | marine metagenome | 124787 | 0 | 251 | NOB | 0.20 | coral |
| SRR7035663 | marine metagenome | 124830 | 0 | 251 | NOB | 0.20 | coral |
| SRR1659073 | mollusc metagenome | 3561 | 0 | 7 | NOB | 0.20 | other marine invertebrates |
| SRR4244546 | marine metagenome | 75009 | 147 | 821 | AOA and NOB | 0.20 | sponge |
| SRR7035534 | marine metagenome | 96732 | 0 | 189 | NOB | 0.20 | coral |
| SRR7596591 | marine metagenome | 97051 | 0 | 189 | NOB | 0.19 | coral |
| SRR8380135 | Porites lobata | 13881 | 0 | 27 | NOB | 0.19 | coral |

| | | | | | | | |
|---|---|---|---|---|---|---|---|
| SRR4471384 | marine metagenome | 15539 | 0 | 30 | NOB | 0.19 | coral |
| SRR3204423 | coral metagenome | 519 | 0 | 1 | NOB | 0.19 | coral |
| SRR6457410 | marine metagenome | 61699 | 116 | 0 | AOA | 0.19 | seawater |
| SRR8359903 | coral reef metagenome | 56387 | 0 | 106 | NOB | 0.19 | coral |
| SRR6393318 | marine metagenome | 33683 | 0 | 63 | NOB | 0.19 | seawater |
| SRR8640009 | algae metagenome | 1609 | 0 | 3 | NOB | 0.19 | algae |
| SRR8359815 | coral reef metagenome | 22194 | 0 | 41 | NOB | 0.18 | coral |
| SRR7591414 | coral metagenome | 9746 | 0 | 18 | NOB | 0.18 | coral |
| SRR8359785 | coral reef metagenome | 30932 | 0 | 57 | NOB | 0.18 | coral |
| SRR7598617 | marine metagenome | 25115 | 0 | 46 | NOB | 0.18 | sponge |
| SRR7034650 | marine metagenome | 25119 | 0 | 46 | NOB | 0.18 | sponge |
| SRR6393313 | marine metagenome | 29220 | 0 | 52 | NOB | 0.18 | seawater |
| SRR6081892 | sediment metagenome | 17983 | 0 | 32 | NOB | 0.18 | marine sediment |
| SRR3204474 | coral metagenome | 89981 | 3105 | 158 | AOA and NOB | 0.18 | coral |
| ERR951511 | metagenome | 576 | 0 | 1 | NOB | 0.17 | coral |
| ERR1419290 | marine metagenome | 85302 | 0 | 147 | NOB | 0.17 | other |
| SRR7034437 | marine metagenome | 13532 | 0 | 23 | NOB | 0.17 | marine sediment |
| SRR7598547 | marine metagenome | 13583 | 0 | 23 | NOB | 0.17 | marine sediment |
| SRR3204409 | coral metagenome | 1187 | 2 | 0 | AOA | 0.17 | coral |
| SRR1659074 | mollusc metagenome | 2396 | 0 | 4 | NOB | 0.17 | other marine invertebrates |
| SRR4471368 | marine metagenome | 3604 | 6 | 1 | AOA and NOB | 0.17 | coral |
| SRR7035661 | marine metagenome | 70895 | 0 | 117 | NOB | 0.17 | coral |
| SRR7596249 | marine metagenome | 71033 | 0 | 117 | NOB | 0.16 | coral |
| SRR8359793 | coral reef metagenome | 15789 | 0 | 26 | NOB | 0.16 | coral |
| SRR3147500 | Xestospongia testudinaria | 608 | 0 | 1 | NOB | 0.16 | sponge |
| SRR4471385 | marine metagenome | 8535 | 14 | 8 | AOA and NOB | 0.16 | coral |
| SRR1791584 | coral metagenome | 1837 | 0 | 3 | NOB | 0.16 | coral |
| SRR7035573 | marine metagenome | 145032 | 0 | 236 | NOB | 0.16 | coral |
| SRR7595745 | marine metagenome | 145435 | 0 | 236 | NOB | 0.16 | coral |
| ERR2016818 | metagenome | 121432 | 0 | 197 | NOB | 0.16 | marine sediment |
| SRR8639946 | algae metagenome | 34660 | 0 | 56 | NOB | 0.16 | algae |
| SRR4471392 | marine metagenome | 10580 | 12 | 17 | AOA and NOB | 0.16 | coral |
| SRR7598610 | marine metagenome | 13922 | 0 | 22 | NOB | 0.16 | coral |
| SRR7036490 | marine metagenome | 13925 | 0 | 22 | NOB | 0.16 | coral |
| SRR6401457 | marine metagenome | 2553 | 0 | 4 | NOB | 0.16 | coral |
| SRR7036425 | marine metagenome | 20093 | 0 | 31 | NOB | 0.15 | sponge |
| SRR7598710 | marine metagenome | 20149 | 0 | 31 | NOB | 0.15 | sponge |
| SRR1992897 | coral metagenome | 34244 | 52 | 1 | AOA and NOB | 0.15 | coral |
| SRR8359902 | coral reef metagenome | 107449 | 0 | 163 | NOB | 0.15 | coral |
| SRR3741979 | marine metagenome | 81642 | 0 | 123 | NOB | 0.15 | marine sediment |
| SRR3104274 | soil metagenome | 8633 | 0 | 13 | NOB | 0.15 | soil |
| SRR4244568 | marine metagenome | 60430 | 90 | 329 | AOA and NOB | 0.15 | sponge |
| SRR4471383 | marine metagenome | 678 | 1 | 1 | AOA and NOB | 0.15 | coral |
| SRR4471383 | marine metagenome | 678 | 1 | 1 | AOA and NOB | 0.15 | coral |
| SRR6401519 | marine metagenome | 36204 | 0 | 53 | NOB | 0.15 | marine sediment |
| ERR174002 | marine metagenome | 6155 | 0 | 9 | NOB | 0.15 | seawater |
| ERR951472 | metagenome | 684 | 0 | 1 | NOB | 0.15 | coral |
| SRR3233446 | Hyrtios erectus | 1371 | 0 | 2 | NOB | 0.15 | sponge |
| SRR7178022 | sponge metagenome | 178430 | 0 | 259 | NOB | 0.15 | sponge |
| SRR8359901 | coral reef metagenome | 99947 | 0 | 144 | NOB | 0.14 | coral |
| SRR3204434 | coral metagenome | 93747 | 135 | 109 | AOA and NOB | 0.14 | coral |
| SRR869531 | sponge metagenome | 12605 | 0 | 18 | NOB | 0.14 | sponge |
| SRR1659080 | mollusc metagenome | 2814 | 0 | 4 | NOB | 0.14 | other marine invertebrates |
| SRR7598609 | marine metagenome | 11586 | 0 | 16 | NOB | 0.14 | coral |
| SRR7036491 | marine metagenome | 11594 | 0 | 16 | NOB | 0.14 | coral |
| SRR6081894 | sediment metagenome | 19598 | 0 | 27 | NOB | 0.14 | marine sediment |
| SRR4471362 | marine metagenome | 1455 | 1 | 2 | AOA and NOB | 0.14 | coral |
| SRR8359883 | coral reef metagenome | 40866 | 0 | 56 | NOB | 0.14 | coral |
| SRR8380134 | Porites lobata | 12881 | 0 | 17 | NOB | 0.13 | coral |
| SRR1263017 | 2 | 118834 | 0 | 153 | NOB | 0.13 | coral |
| SRR651960 | sponge metagenome | 85292 | 109 | 0 | AOA | 0.13 | sponge |
| SRR6401458 | marine metagenome | 2363 | 0 | 3 | NOB | 0.13 | coral |
| SRR8359806 | coral reef metagenome | 38406 | 0 | 48 | NOB | 0.12 | coral |
| SRR6403601 | marine metagenome | 25608 | 0 | 32 | NOB | 0.12 | seawater |
| SRR4471381 | marine metagenome | 808 | 0 | 1 | NOB | 0.12 | coral |
| SRR7034298 | marine metagenome | 21913 | 0 | 27 | NOB | 0.12 | sponge |
| SRR7598350 | marine metagenome | 21939 | 0 | 27 | NOB | 0.12 | sponge |
| SRR8359801 | coral reef metagenome | 37486 | 0 | 46 | NOB | 0.12 | coral |
| SRR3147494 | Xestospongia testudinaria | 4941 | 0 | 6 | NOB | 0.12 | sponge |
| SRR8359776 | coral reef metagenome | 54794 | 0 | 66 | NOB | 0.12 | coral |
| ERR2073755 | Aplysina cauliformis | 621338 | 17305 | 748 | AOA and NOB | 0.12 | sponge |
| ERR951473 | metagenome | 831 | 0 | 1 | NOB | 0.12 | coral |
| SRR7034553 | marine metagenome | 11676 | 0 | 14 | NOB | 0.12 | marine sediment |
| SRR8639948 | algae metagenome | 85909 | 0 | 103 | NOB | 0.12 | algae |
| SRR7598510 | marine metagenome | 11713 | 0 | 14 | NOB | 0.12 | marine sediment |
| SRR6401591 | marine metagenome | 30174 | 0 | 36 | NOB | 0.12 | marine sediment |
| ERR951498 | metagenome | 3363 | 0 | 4 | NOB | 0.12 | coral |
| SRR8359824 | coral reef metagenome | 81104 | 0 | 96 | NOB | 0.12 | coral |
| SRR6081895 | sediment metagenome | 19542 | 0 | 23 | NOB | 0.12 | marine sediment |
| SRR8581542 | annelid metagenome | 92605 | 0 | 108 | NOB | 0.12 | other marine invertebrates |
| SRR3204434 | coral metagenome | 93747 | 135 | 109 | AOA and NOB | 0.12 | coral |
| SRR8359770 | coral reef metagenome | 13847 | 0 | 16 | NOB | 0.12 | coral |
| SRR651959 | sponge metagenome | 27181 | 0 | 31 | NOB | 0.11 | sponge |
| SRR7598322 | marine metagenome | 19306 | 0 | 22 | NOB | 0.11 | marine sediment |
| SRR4471392 | marine metagenome | 10580 | 12 | 17 | AOA and NOB | 0.11 | coral |
| SRR7591399 | coral metagenome | 22042 | 0 | 25 | NOB | 0.11 | coral |
| SRR6401589 | marine metagenome | 53876 | 0 | 61 | NOB | 0.11 | marine sediment |
| SRR7034591 | marine metagenome | 8839 | 0 | 10 | NOB | 0.11 | marine sediment |
| SRR1393908 | sea squirt metagenome | 6191 | 0 | 7 | NOB | 0.11 | other marine invertebrates |
| SRR7034269 | marine metagenome | 19502 | 0 | 22 | NOB | 0.11 | marine sediment |
| SRR5879046 | sediment metagenome | 54497 | 0 | 61 | NOB | 0.11 | marine sediment |
| SRR4471394 | marine metagenome | 904 | 0 | 1 | NOB | 0.11 | coral |
| SRR5469071 | seawater metagenome | 86163 | 15 | 95 | AOA and NOB | 0.11 | seawater |
| SRR6401401 | marine metagenome | 50926 | 0 | 56 | NOB | 0.11 | marine sediment |
| ERR1419304 | marine metagenome | 145097 | 0 | 159 | NOB | 0.11 | other |
| SRR6401517 | marine metagenome | 23883 | 0 | 26 | NOB | 0.11 | marine sediment |
| ERR1801519 | marine metagenome | 6527 | 0 | 7 | NOB | 0.11 | other |

| | | | | | | | |
|---|---|---|---|---|---|---|---|
| SRR7034501 | marine metagenome | 20560 | 0 | 22 | NOB | 0.11 | marine sediment |
| SRR3204413 | coral metagenome | 37458 | 1032 | 40 | AOA and NOB | 0.11 | coral |
| SRR7598735 | marine metagenome | 20656 | 0 | 22 | NOB | 0.11 | marine sediment |
| SRR3204414 | coral metagenome | 47933 | 1119 | 51 | AOA and NOB | 0.11 | coral |
| SRR7517646 | indoor metagenome | 1927 | 2 | 0 | AOA | 0.10 | other |
| SRR7034500 | marine metagenome | 8713 | 0 | 9 | NOB | 0.10 | marine sediment |
| SRR7034584 | marine metagenome | 9709 | 0 | 10 | NOB | 0.10 | marine sediment |
| SRR709456 | marine metagenome | 50615 | 0 | 52 | NOB | 0.10 | marine benthic and pelagic environments |
| SRR7598736 | marine metagenome | 8768 | 0 | 9 | NOB | 0.10 | marine sediment |
| SRR7036426 | marine metagenome | 18579 | 0 | 19 | NOB | 0.10 | sponge |
| SRR7598709 | marine metagenome | 18586 | 0 | 19 | NOB | 0.10 | sponge |
| SRR8640052 | algae metagenome | 80390 | 0 | 82 | NOB | 0.10 | algae |
| SRR6393361 | marine metagenome | 68958 | 0 | 70 | NOB | 0.10 | seawater |
| SRR8359784 | coral reef metagenome | 39026 | 0 | 39 | NOB | 0.10 | coral |
| SRR8359775 | coral reef metagenome | 78365 | 0 | 78 | NOB | 0.10 | coral |
| SRR8359860 | coral reef metagenome | 30205 | 0 | 30 | NOB | 0.10 | coral |
| SRR8581541 | annelid metagenome | 122082 | 0 | 121 | NOB | 0.10 | other marine invertebrates |
| SRR7591412 | coral metagenome | 4039 | 74 | 4 | AOA and NOB | 0.10 | coral |
| SRR1992929 | coral metagenome | 58595 | 57 | 0 | AOA | 0.10 | coral |
| SRR6401593 | marine metagenome | 44414 | 0 | 43 | NOB | 0.10 | marine sediment |
| SRR7598374 | marine metagenome | 7305 | 0 | 7 | NOB | 0.10 | coral |
| SRR7036535 | marine metagenome | 7324 | 0 | 7 | NOB | 0.10 | coral |
| SRR4471387 | marine metagenome | 3140 | 3 | 2 | AOA and NOB | 0.10 | coral |
| SRR7034538 | marine metagenome | 24103 | 0 | 23 | NOB | 0.10 | seawater |
| SRR7599232 | marine metagenome | 24118 | 0 | 23 | NOB | 0.10 | seawater |
| ERR1419302 | marine metagenome | 65062 | 0 | 62 | NOB | 0.10 | other |
| SRR4471385 | marine metagenome | 8535 | 14 | 8 | AOA and NOB | 0.09 | coral |
| SRR8359886 | coral reef metagenome | 35289 | 0 | 33 | NOB | 0.09 | coral |
| SRR7598740 | marine metagenome | 9687 | 0 | 9 | NOB | 0.09 | marine sediment |
| SRR7598382 | marine metagenome | 18320 | 0 | 17 | NOB | 0.09 | marine sediment |
| SRR7034590 | marine metagenome | 18344 | 0 | 17 | NOB | 0.09 | marine sediment |
| SRR6401397 | marine metagenome | 33550 | 0 | 31 | NOB | 0.09 | marine sediment |
| SRR3233445 | Hyrtios erectus | 2197 | 0 | 2 | NOB | 0.09 | sponge |
| SRR3204455 | coral metagenome | 75944 | 69 | 31 | AOA and NOB | 0.09 | coral |
| SRR3147481 | Aaptos suberitoides | 6608 | 0 | 6 | NOB | 0.09 | sponge |
| SRR3204429 | coral metagenome | 56189 | 35 | 51 | AOA and NOB | 0.09 | coral |
| SRR8380146 | Porites lobata | 5554 | 0 | 5 | NOB | 0.09 | coral |
| ERR1419294 | marine metagenome | 117002 | 0 | 103 | NOB | 0.09 | other |
| SRR6401673 | marine metagenome | 43512 | 0 | 38 | NOB | 0.09 | marine sediment |
| SRR8380147 | Porites lobata | 5791 | 0 | 5 | NOB | 0.09 | coral |
| SRR8359851 | coral reef metagenome | 18622 | 0 | 16 | NOB | 0.09 | coral |
| SRR4244545 | marine metagenome | 101178 | 17 | 86 | AOA and NOB | 0.08 | sponge |
| SRR7598351 | marine metagenome | 21471 | 0 | 18 | NOB | 0.08 | sponge |
| SRR3233801 | Stylissa massa | 3584 | 3 | 0 | AOA | 0.08 | sponge |
| SRR7034756 | marine metagenome | 21570 | 0 | 18 | NOB | 0.08 | sponge |
| SRR8359796 | coral reef metagenome | 68438 | 0 | 57 | NOB | 0.08 | coral |
| SRR8359884 | coral reef metagenome | 33759 | 0 | 28 | NOB | 0.08 | coral |
| SRR6401403 | marine metagenome | 52603 | 0 | 43 | NOB | 0.08 | sponge |
| SRR8548154 | coral metagenome | 13493 | 0 | 11 | NOB | 0.08 | coral |
| SRR8359882 | coral reef metagenome | 113745 | 0 | 92 | NOB | 0.08 | coral |
| SRR5927070 | marine metagenome | 12412 | 0 | 10 | NOB | 0.08 | marine sediment |
| SRR6401365 | marine metagenome | 57287 | 0 | 46 | NOB | 0.08 | coral |
| SRR8359907 | coral reef metagenome | 129563 | 0 | 104 | NOB | 0.08 | coral |
| SRR6457411 | marine metagenome | 66034 | 53 | 0 | AOA | 0.08 | seawater |
| SRR6401675 | marine metagenome | 47660 | 0 | 38 | NOB | 0.08 | marine sediment |
| SRR3233447 | Hyrtios erectus | 3763 | 0 | 3 | NOB | 0.08 | sponge |
| SRR7598680 | marine metagenome | 12587 | 0 | 10 | NOB | 0.08 | marine sediment |
| SRR3742013 | marine metagenome | 81200 | 1 | 64 | AOA and NOB | 0.08 | marine sediment |
| SRR8596052 | coral metagenome | 25415 | 20 | 12 | AOA and NOB | 0.08 | coral |
| ERR2113131 | metagenome | 426566 | 0 | 335 | NOB | 0.08 | marine sediment |
| SRR3204454 | coral metagenome | 105934 | 83 | 63 | AOA and NOB | 0.08 | coral |
| SRR6403308 | marine metagenome | 24367 | 0 | 19 | NOB | 0.08 | seawater |
| SRR4471400 | marine metagenome | 1283 | 0 | 1 | NOB | 0.08 | coral |
| SRR7034755 | marine metagenome | 25773 | 0 | 20 | NOB | 0.08 | sponge |
| SRR7598352 | marine metagenome | 25774 | 0 | 20 | NOB | 0.08 | sponge |
| SRR8359908 | coral reef metagenome | 110971 | 0 | 86 | NOB | 0.08 | coral |
| SRR7034392 | marine metagenome | 11628 | 0 | 9 | NOB | 0.08 | marine sediment |
| SRR7598489 | marine metagenome | 6476 | 0 | 5 | NOB | 0.08 | sponge |
| SRR7034520 | marine metagenome | 6494 | 0 | 5 | NOB | 0.08 | sponge |
| SRR6401805 | marine metagenome | 54600 | 0 | 42 | NOB | 0.08 | coral |
| SRR7034581 | marine metagenome | 16962 | 0 | 13 | NOB | 0.08 | marine sediment |
| SRR7598394 | marine metagenome | 17026 | 0 | 13 | NOB | 0.08 | marine sediment |
| SRR3184451 | coral metagenome | 27688 | 0 | 21 | NOB | 0.08 | coral |
| SRR8581536 | annelid metagenome | 132203 | 0 | 100 | NOB | 0.08 | other marine invertebrates |
| SRR7034757 | marine metagenome | 10631 | 0 | 8 | NOB | 0.08 | marine sediment |
| SRR8359840 | coral reef metagenome | 22624 | 0 | 17 | NOB | 0.08 | coral |
| SRR7513544 | sponge metagenome | 77504 | 0 | 58 | NOB | 0.07 | sponge |
| SRR8359905 | coral reef metagenome | 79130 | 0 | 59 | NOB | 0.07 | coral |
| SRR7177973 | sponge metagenome | 212267 | 0 | 158 | NOB | 0.07 | sponge |
| SRR6401511 | marine metagenome | 33788 | 0 | 25 | NOB | 0.07 | marine sediment |
| SRR8359768 | coral reef metagenome | 13529 | 0 | 10 | NOB | 0.07 | coral |
| ERR1419297 | marine metagenome | 88358 | 0 | 65 | NOB | 0.07 | other |
| SRR4471390 | marine metagenome | 1367 | 1 | 1 | AOA and NOB | 0.07 | coral |
| SRR4471390 | marine metagenome | 1367 | 1 | 1 | AOA and NOB | 0.07 | coral |
| ERR951519 | metagenome | 4111 | 0 | 3 | NOB | 0.07 | coral |
| SRR6401399 | marine metagenome | 46740 | 0 | 34 | NOB | 0.07 | marine sediment |
| SRR7598325 | marine metagenome | 9637 | 0 | 7 | NOB | 0.07 | marine sediment |
| SRR8751341 | coral metagenome | 27736 | 0 | 20 | NOB | 0.07 | coral |
| SRR8359836 | coral reef metagenome | 51503 | 0 | 37 | NOB | 0.07 | coral |
| SRR8359790 | coral reef metagenome | 26451 | 0 | 19 | NOB | 0.07 | coral |
| SRR8359774 | coral reef metagenome | 15335 | 0 | 11 | NOB | 0.07 | coral |
| SRR6401647 | marine metagenome | 32249 | 0 | 23 | NOB | 0.07 | marine sediment |
| SRR869534 | sponge metagenome | 15670 | 0 | 11 | NOB | 0.07 | sponge |
| SRR869529 | sponge metagenome | 9990 | 0 | 7 | NOB | 0.07 | sponge |
| SRR6401331 | marine metagenome | 41537 | 0 | 29 | NOB | 0.07 | marine sediment |
| SRR8359855 | coral reef metagenome | 34549 | 0 | 24 | NOB | 0.07 | coral |
| SRR4471362 | marine metagenome | 1455 | 1 | 2 | AOA and NOB | 0.07 | coral |

| | | | | | | | |
|---|---|---|---|---|---|---|---|
| SRR8359839 | coral reef metagenome | 16065 | 0 | 11 | NOB | 0.07 | coral |
| SRR8359838 | coral reef metagenome | 18996 | 0 | 13 | NOB | 0.07 | coral |
| SRR8380145 | Porites lobata | 8802 | 0 | 6 | NOB | 0.07 | coral |
| SRR8639806 | algae metagenome | 41494 | 0 | 28 | NOB | 0.07 | algae |
| ERR951507 | metagenome | 4495 | 0 | 3 | NOB | 0.07 | coral |
| SRR4471373 | marine metagenome | 1503 | 1 | 0 | AOA | 0.07 | coral |
| SRR7591426 | coral metagenome | 9163 | 0 | 6 | NOB | 0.07 | coral |
| SRR8359891 | coral reef metagenome | 20118 | 0 | 13 | NOB | 0.06 | coral |
| SRR2622048 | coral metagenome | 136478 | 0 | 88 | NOB | 0.06 | coral |
| SRR8111657 | mollusc metagenome | 12409 | 8 | 35 | AOA and NOB | 0.06 | other marine invertebrates |
| SRR7598443 | marine metagenome | 7760 | 0 | 5 | NOB | 0.06 | coral |
| SRR6401515 | marine metagenome | 40419 | 0 | 26 | NOB | 0.06 | marine sediment |
| ERR1894941 | metagenome | 40488 | 0 | 26 | NOB | 0.06 | other |
| SRR8596081 | coral metagenome | 9357 | 6 | 0 | AOA | 0.06 | coral |
| SRR8359797 | coral reef metagenome | 67281 | 0 | 43 | NOB | 0.06 | coral |
| SRR4471387 | marine metagenome | 3140 | 3 | 2 | AOA and NOB | 0.06 | coral |
| SRR3147496 | Xestospongia testudinaria | 3159 | 0 | 2 | NOB | 0.06 | sponge |
| SRR3204429 | coral metagenome | 56189 | 35 | 51 | AOA and NOB | 0.06 | coral |
| SRR6401575 | marine metagenome | 46635 | 0 | 29 | NOB | 0.06 | marine sediment |
| SRR3160544 | algae metagenome | 193347 | 0 | 120 | NOB | 0.06 | algae |
| SRR4471401 | marine metagenome | 1615 | 1 | 0 | AOA | 0.06 | coral |
| SRR5879081 | sediment metagenome | 64608 | 0 | 40 | NOB | 0.06 | marine sediment |
| SRR8359800 | coral reef metagenome | 17848 | 0 | 11 | NOB | 0.06 | coral |
| ERR951487 | metagenome | 4914 | 0 | 3 | NOB | 0.06 | coral |
| SRR4244560 | marine metagenome | 55930 | 0 | 34 | NOB | 0.06 | sponge |
| SRR7591395 | coral metagenome | 4940 | 3 | 0 | AOA | 0.06 | coral |
| SRR7598398 | marine metagenome | 14910 | 0 | 9 | NOB | 0.06 | marine sediment |
| SRR8380143 | Porites lobata | 13375 | 0 | 8 | NOB | 0.06 | coral |
| ERR1894942 | metagenome | 45212 | 0 | 27 | NOB | 0.06 | other |
| SRR3204454 | coral metagenome | 105934 | 83 | 63 | AOA and NOB | 0.06 | coral |
| SRR7598358 | marine metagenome | 11944 | 0 | 7 | NOB | 0.06 | marine sediment |
| SRR8359803 | coral reef metagenome | 41887 | 0 | 24 | NOB | 0.06 | coral |
| SRR1040545 | gut metagenome | 1748 | 0 | 1 | NOB | 0.06 | other |
| SRR6081893 | sediment metagenome | 27981 | 0 | 16 | NOB | 0.06 | marine sediment |
| SRR4244580 | marine metagenome | 45664 | 3 | 26 | AOA and NOB | 0.06 | sponge |
| SRR7036539 | marine metagenome | 14067 | 0 | 8 | NOB | 0.06 | coral |
| ERR173988 | marine metagenome | 7037 | 0 | 4 | NOB | 0.06 | seawater |
| SRR7035525 | marine metagenome | 81238 | 0 | 46 | NOB | 0.06 | marine sediment |
| SRR4471398 | marine metagenome | 5305 | 0 | 3 | NOB | 0.06 | coral |
| SRR4471382 | marine metagenome | 3545 | 2 | 32 | AOA and NOB | 0.06 | coral |
| SRR7513570 | sponge metagenome | 51477 | 0 | 29 | NOB | 0.06 | sponge |
| SRR7598484 | marine metagenome | 14221 | 0 | 8 | NOB | 0.06 | coral |
| SRR869559 | sponge metagenome | 19624 | 0 | 11 | NOB | 0.06 | sponge |
| SRR4471380 | marine metagenome | 3572 | 2 | 1 | AOA and NOB | 0.06 | coral |
| SRR7595840 | marine metagenome | 82398 | 0 | 46 | NOB | 0.06 | marine sediment |
| SRR6401333 | marine metagenome | 42387 | 0 | 23 | NOB | 0.05 | marine sediment |
| DRR095250 | estuary metagenome | 24065 | 0 | 13 | NOB | 0.05 | other |
| SRR8359782 | coral reef metagenome | 22238 | 0 | 12 | NOB | 0.05 | coral |
| SRR7034438 | marine metagenome | 9363 | 0 | 5 | NOB | 0.05 | marine sediment |
| ERR2073757 | Aplysina cauliformis | 2479361 | 121901 | 1324 | AOA and NOB | 0.05 | sponge |
| SRR3741983 | marine metagenome | 37608 | 20 | 0 | AOA | 0.05 | marine sediment |
| SRR8359808 | coral reef metagenome | 28405 | 0 | 15 | NOB | 0.05 | coral |
| SRR4244559 | marine metagenome | 95050 | 0 | 50 | NOB | 0.05 | sponge |
| SRR7598546 | marine metagenome | 9531 | 0 | 5 | NOB | 0.05 | marine sediment |
| SRR4471359 | marine metagenome | 7682 | 22 | 4 | AOA and NOB | 0.05 | coral |
| SRR2087831 | plant metagenome | 150364 | 77 | 0 | AOA | 0.05 | plant |
| SRR6401436 | marine metagenome | 25816 | 0 | 13 | NOB | 0.05 | marine sediment |
| SRR7598676 | marine metagenome | 13950 | 0 | 7 | NOB | 0.05 | marine sediment |
| SRR867625 | marine metagenome | 3988 | 2 | 0 | AOA | 0.05 | seawater |
| SRR8359837 | coral reef metagenome | 34133 | 0 | 17 | NOB | 0.05 | coral |
| SRR8661901 | coral metagenome | 20113 | 0 | 10 | NOB | 0.05 | coral |
| SRR7036537 | marine metagenome | 10278 | 0 | 5 | NOB | 0.05 | coral |
| SRR7598482 | marine metagenome | 10281 | 0 | 5 | NOB | 0.05 | coral |
| SRR3000457 | coral metagenome | 12415 | 0 | 6 | NOB | 0.05 | coral |
| SRR7034331 | marine metagenome | 10354 | 0 | 5 | NOB | 0.05 | marine sediment |
| SRR8359767 | coral reef metagenome | 47975 | 0 | 23 | NOB | 0.05 | coral |
| SRR8359853 | coral reef metagenome | 58498 | 0 | 28 | NOB | 0.05 | coral |
| ERR951513 | metagenome | 4223 | 0 | 2 | NOB | 0.05 | coral |
| SRR3204464 | coral metagenome | 71935 | 34 | 30 | AOA and NOB | 0.05 | coral |
| SRR8596052 | coral metagenome | 25415 | 20 | 12 | AOA and NOB | 0.05 | coral |
| ERR1894950 | metagenome | 55861 | 0 | 26 | NOB | 0.05 | other |
| SRR1503493 | marine metagenome | 12894 | 0 | 6 | NOB | 0.05 | seawater |
| SRR6401540 | marine metagenome | 2157 | 0 | 1 | NOB | 0.05 | coral |
| SRR6393388 | marine metagenome | 13119 | 0 | 6 | NOB | 0.05 | seawater |
| SRR6309327 | skin metagenome | 46409 | 0 | 21 | NOB | 0.05 | other |
| SRR4244590 | marine metagenome | 66979 | 0 | 30 | NOB | 0.04 | sponge |
| SRR7034521 | marine metagenome | 6728 | 0 | 3 | NOB | 0.04 | sponge |
| SRR7598488 | marine metagenome | 6737 | 0 | 3 | NOB | 0.04 | sponge |
| ERR951485 | metagenome | 4500 | 0 | 2 | NOB | 0.04 | coral |
| SRR8359789 | coral reef metagenome | 9065 | 0 | 4 | NOB | 0.04 | coral |
| SRR3204449 | coral metagenome | 75161 | 399 | 33 | AOA and NOB | 0.04 | coral |
| SRR8359802 | coral reef metagenome | 39341 | 0 | 17 | NOB | 0.04 | coral |
| SRR5878996 | sediment metagenome | 55615 | 0 | 24 | NOB | 0.04 | marine sediment |
| SRR6403551 | marine metagenome | 28094 | 0 | 12 | NOB | 0.04 | seawater |
| SRR1039444 | sponge metagenome | 7041 | 0 | 3 | NOB | 0.04 | sponge |
| SRR8639749 | algae metagenome | 147948 | 0 | 63 | NOB | 0.04 | algae |
| SRR8359835 | coral reef metagenome | 28225 | 0 | 12 | NOB | 0.04 | coral |
| SRR3000446 | coral metagenome | 4717 | 0 | 2 | NOB | 0.04 | coral |
| ERR1419298 | marine metagenome | 161034 | 0 | 68 | NOB | 0.04 | other |
| SRR5879041 | sediment metagenome | 97411 | 0 | 41 | NOB | 0.04 | marine sediment |
| SRR8359778 | coral reef metagenome | 11922 | 0 | 5 | NOB | 0.04 | coral |
| SRR8111681 | mollusc metagenome | 7154 | 3 | 3 | AOA and NOB | 0.04 | other marine invertebrates |
| SRR8111681 | mollusc metagenome | 7154 | 3 | 3 | AOA and NOB | 0.04 | other marine invertebrates |
| SRR2087830 | plant metagenome | 86256 | 36 | 0 | AOA | 0.04 | plant |
| SRR3204464 | coral metagenome | 71935 | 34 | 30 | AOA and NOB | 0.04 | coral |
| SRR8359864 | coral reef metagenome | 24034 | 0 | 10 | NOB | 0.04 | coral |
| SRR4458031 | marine metagenome | 36073 | 0 | 15 | NOB | 0.04 | seawater |

| | | | | | | | |
|---|---|---|---|---|---|---|---|
| SRR869538 | sponge metagenome | 12069 | 0 | 5 | NOB | 0.04 | sponge |
| SRR8359799 | coral reef metagenome | 14500 | 0 | 6 | NOB | 0.04 | coral |
| SRR7598331 | marine metagenome | 17071 | 0 | 7 | NOB | 0.04 | marine sediment |
| SRR3742012 | marine metagenome | 75659 | 31 | 0 | AOA | 0.04 | marine sediment |
| SRR3204455 | coral metagenome | 75944 | 69 | 31 | AOA and NOB | 0.04 | coral |
| ERR1777344 | sediment metagenome | 22063 | 0 | 9 | NOB | 0.04 | marine sediment |
| SRR8359772 | coral reef metagenome | 19664 | 0 | 8 | NOB | 0.04 | coral |
| SRR8751291 | coral metagenome | 7403 | 0 | 3 | NOB | 0.04 | coral |
| SRR8243359 | coral reef metagenome | 12394 | 1 | 5 | AOA and NOB | 0.04 | coral |
| SRR7513545 | sponge metagenome | 69418 | 0 | 28 | NOB | 0.04 | sponge |
| SRR6403320 | marine metagenome | 27301 | 0 | 11 | NOB | 0.04 | seawater |
| SRR8751369 | coral metagenome | 9994 | 0 | 4 | NOB | 0.04 | coral |
| ERR1894951 | metagenome | 32490 | 0 | 13 | NOB | 0.04 | other |
| SRR8359830 | coral reef metagenome | 37619 | 0 | 15 | NOB | 0.04 | coral |
| SRR6401513 | marine metagenome | 32632 | 0 | 13 | NOB | 0.04 | marine sediment |
| SRR3204472 | coral metagenome | 45664 | 2 | 18 | AOA and NOB | 0.04 | coral |
| SRR4244553 | marine metagenome | 66014 | 0 | 26 | NOB | 0.04 | sponge |
| SRR8359834 | coral reef metagenome | 59499 | 0 | 23 | NOB | 0.04 | coral |
| SRR8361252 | marine sediment metagenome | 67324 | 0 | 26 | NOB | 0.04 | marine sediment |
| SRR3233472 | Xestospongia testudinaria | 5179 | 0 | 2 | NOB | 0.04 | sponge |
| SRR3742052 | marine metagenome | 62727 | 0 | 24 | NOB | 0.04 | marine sediment |
| SRR7034272 | marine metagenome | 10461 | 0 | 4 | NOB | 0.04 | marine sediment |
| SRR1595988 | marine metagenome | 5293 | 2 | 0 | AOA | 0.04 | sponge |
| SRR7513600 | marine metagenome | 23847 | 0 | 9 | NOB | 0.04 | seawater |
| SRR8751352 | coral metagenome | 15912 | 0 | 6 | NOB | 0.04 | coral |
| SRR1585360 | marine metagenome | 5313 | 2 | 0 | AOA | 0.04 | sponge |
| SRR1734272 | beach sand metagenome | 173051 | 0 | 65 | NOB | 0.04 | marine sediment |
| SRR4244555 | marine metagenome | 72026 | 0 | 27 | NOB | 0.04 | sponge |
| SRR7513546 | sponge metagenome | 77518 | 0 | 29 | NOB | 0.04 | sponge |
| SRR8380133 | Porites lobata | 16071 | 0 | 6 | NOB | 0.04 | coral |
| SRR6309341 | skin metagenome | 34924 | 13 | 0 | AOA | 0.04 | other |
| SRR8359795 | coral reef metagenome | 37654 | 0 | 14 | NOB | 0.04 | coral |
| SRR7598511 | marine metagenome | 13547 | 0 | 5 | NOB | 0.04 | marine sediment |
| SRR7034551 | marine metagenome | 13549 | 0 | 5 | NOB | 0.04 | marine sediment |
| ERR1777346 | sediment metagenome | 35339 | 0 | 13 | NOB | 0.04 | marine sediment |
| SRR7034580 | marine metagenome | 10883 | 0 | 4 | NOB | 0.04 | marine sediment |
| SRR7513601 | marine metagenome | 87155 | 0 | 32 | NOB | 0.04 | seawater |
| SRR7034614 | marine metagenome | 16395 | 0 | 6 | NOB | 0.04 | sponge |
| SRR7598353 | marine metagenome | 16444 | 0 | 6 | NOB | 0.04 | sponge |
| SRR7034764 | marine metagenome | 93277 | 0 | 34 | NOB | 0.04 | marine sediment |
| SRR8639804 | algae metagenome | 88090 | 0 | 32 | NOB | 0.04 | algae |
| SRR2559819 | sponge metagenome | 5519 | 2 | 158 | AOA and NOB | 0.04 | sponge |
| SRR7597117 | marine metagenome | 94045 | 0 | 34 | NOB | 0.04 | marine sediment |
| SRR8359879 | coral reef metagenome | 44455 | 0 | 16 | NOB | 0.04 | coral |
| SRR7035205 | marine metagenome | 98634 | 0 | 35 | NOB | 0.04 | marine sediment |
| SRR1237903 | marine sediment metagenome | 19732 | 0 | 7 | NOB | 0.04 | marine sediment |
| SRR7597039 | marine metagenome | 98945 | 0 | 35 | NOB | 0.04 | marine sediment |
| SRR8359765 | coral reef metagenome | 14192 | 0 | 5 | NOB | 0.04 | coral |
| SRR8639950 | algae metagenome | 105593 | 0 | 37 | NOB | 0.04 | algae |
| SRR1653170 | bioreactor metagenome | 86259 | 0 | 30 | NOB | 0.03 | other |
| SRR8359890 | coral reef metagenome | 25955 | 0 | 9 | NOB | 0.03 | coral |
| SRR8487597 | coral metagenome | 28947 | 0 | 10 | NOB | 0.03 | coral |
| SRR4471397 | marine metagenome | 2900 | 0 | 1 | NOB | 0.03 | coral |
| SRR1565484 | sponge metagenome | 17513 | 6 | 266 | AOA and NOB | 0.03 | sponge |
| SRR2040969 | aquatic metagenome | 59007 | 0 | 20 | NOB | 0.03 | seawater |
| ERR2113134 | metagenome | 144819 | 0 | 49 | NOB | 0.03 | marine sediment |
| SRR7513542 | sponge metagenome | 44349 | 0 | 15 | NOB | 0.03 | sponge |
| SRR8639736 | algae metagenome | 47424 | 0 | 16 | NOB | 0.03 | algae |
| SRR8359811 | coral reef metagenome | 62501 | 0 | 21 | NOB | 0.03 | coral |
| SRR8359818 | coral reef metagenome | 41741 | 0 | 14 | NOB | 0.03 | coral |
| SRR8359858 | coral reef metagenome | 56680 | 0 | 19 | NOB | 0.03 | coral |
| SRR3742021 | marine metagenome | 75992 | 0 | 25 | NOB | 0.03 | marine sediment |
| SRR8639770 | algae metagenome | 39528 | 0 | 13 | NOB | 0.03 | algae |
| SRR6401329 | marine metagenome | 45623 | 0 | 15 | NOB | 0.03 | marine sediment |
| SRR7591422 | coral metagenome | 3061 | 1 | 0 | AOA | 0.03 | coral |
| SRR7035381 | marine metagenome | 123093 | 0 | 40 | NOB | 0.03 | marine sediment |
| SRR2000143 | marine sediment metagenome | 21555 | 0 | 7 | NOB | 0.03 | seagrass |
| SRR7596313 | marine metagenome | 123302 | 0 | 40 | NOB | 0.03 | marine sediment |
| SRR2975887 | coral metagenome | 9329 | 0 | 3 | NOB | 0.03 | coral |
| SRR7535354 | sediment metagenome | 28131 | 0 | 9 | NOB | 0.03 | marine sediment |
| SRR8359846 | coral reef metagenome | 87855 | 0 | 28 | NOB | 0.03 | coral |
| SRR7513543 | sponge metagenome | 69298 | 0 | 22 | NOB | 0.03 | sponge |
| SRR3180805 | coral metagenome | 118159 | 0 | 37 | NOB | 0.03 | coral |
| SRR7598506 | marine metagenome | 9872 | 0 | 3 | NOB | 0.03 | marine sediment |
| SRR7034583 | marine metagenome | 9941 | 0 | 3 | NOB | 0.03 | marine sediment |
| SRR8359854 | coral reef metagenome | 49828 | 0 | 15 | NOB | 0.03 | coral |
| SRR7035013 | marine metagenome | 123135 | 0 | 37 | NOB | 0.03 | marine sediment |
| SRR7597081 | marine metagenome | 123217 | 0 | 37 | NOB | 0.03 | marine sediment |
| SRR867623 | marine metagenome | 3332 | 1 | 0 | AOA | 0.03 | seawater |
| SRR8751314 | coral metagenome | 23436 | 0 | 7 | NOB | 0.03 | coral |
| SRR7598465 | marine metagenome | 13514 | 0 | 4 | NOB | 0.03 | coral |
| SRR5879043 | sediment metagenome | 57566 | 0 | 17 | NOB | 0.03 | marine sediment |
| SRR8359845 | coral reef metagenome | 115318 | 0 | 34 | NOB | 0.03 | coral |
| SRR1039439 | sponge metagenome | 6805 | 0 | 2 | NOB | 0.03 | sponge |
| SRR3233794 | Stylissa carteri | 10210 | 3 | 0 | AOA | 0.03 | sponge |
| SRR8359813 | coral reef metagenome | 23939 | 0 | 7 | NOB | 0.03 | coral |
| SRR6401578 | marine metagenome | 78694 | 0 | 23 | NOB | 0.03 | coral |
| SRR7513541 | sponge metagenome | 37678 | 0 | 11 | NOB | 0.03 | sponge |
| ERR951504 | metagenome | 3438 | 1 | 1 | AOA and NOB | 0.03 | coral |
| ERR951504 | metagenome | 3438 | 1 | 1 | AOA and NOB | 0.03 | coral |
| SRR7598754 | marine metagenome | 20628 | 0 | 6 | NOB | 0.03 | seawater |
| SRR7034371 | marine metagenome | 20654 | 0 | 6 | NOB | 0.03 | seawater |
| SRR7034444 | marine metagenome | 10339 | 0 | 3 | NOB | 0.03 | marine sediment |
| SRR2559809 | sponge metagenome | 17272 | 5 | 440 | AOA and NOB | 0.03 | sponge |
| SRR3204477 | coral metagenome | 59562 | 120 | 17 | AOA and NOB | 0.03 | coral |
| SRR4471408 | marine metagenome | 17522 | 0 | 5 | NOB | 0.03 | coral |
| SRR6309339 | skin metagenome | 35078 | 1 | 10 | AOA and NOB | 0.03 | other |

| | | | | | | | |
|---|---|---|---|---|---|---|---|
| SRR7513547 | sponge metagenome | 45696 | 0 | 13 | NOB | 0.03 | sponge |
| SRR5469072 | seawater metagenome | 119940 | 6 | 34 | AOA and NOB | 0.03 | seawater |
| SRR8359829 | coral reef metagenome | 24792 | 0 | 7 | NOB | 0.03 | coral |
| SRR4471380 | marine metagenome | 3572 | 2 | 1 | AOA and NOB | 0.03 | coral |
| SRR4471368 | marine metagenome | 3604 | 6 | 1 | AOA and NOB | 0.03 | coral |
| SRR4471367 | marine metagenome | 14419 | 0 | 4 | NOB | 0.03 | coral |
| SRR8359833 | coral reef metagenome | 32545 | 0 | 9 | NOB | 0.03 | coral |
| SRR7598461 | marine metagenome | 14548 | 0 | 4 | NOB | 0.03 | coral |
| SRR2559821 | sponge metagenome | 7279 | 2 | 124 | AOA and NOB | 0.03 | sponge |
| SRR7513539 | sponge metagenome | 54866 | 0 | 15 | NOB | 0.03 | sponge |
| SRR3233800 | Stylissa massa | 14649 | 4 | 0 | AOA | 0.03 | sponge |
| SRR3204420 | coral metagenome | 7437 | 2 | 0 | AOA | 0.03 | coral |
| DRR014865 | soil metagenome | 11191 | 0 | 3 | NOB | 0.03 | soil |
| SRR8359878 | coral reef metagenome | 26128 | 0 | 7 | NOB | 0.03 | coral |
| SRR2032895 | soil metagenome | 41304 | 0 | 11 | NOB | 0.03 | soil |
| ERR1777337 | sediment metagenome | 60080 | 0 | 16 | NOB | 0.03 | marine sediment |
| SRR1734243 | beach sand metagenome | 93886 | 0 | 25 | NOB | 0.03 | marine sediment |
| SRR8359892 | coral reef metagenome | 37601 | 0 | 10 | NOB | 0.03 | coral |
| SRR8751261 | coral metagenome | 18810 | 0 | 5 | NOB | 0.03 | coral |
| SRR1145124 | sediment metagenome | 3790 | 0 | 1 | NOB | 0.03 | marine sediment |
| SRR7513613 | marine metagenome | 57074 | 0 | 15 | NOB | 0.03 | seawater |
| SRR8548000 | coral metagenome | 19097 | 0 | 5 | NOB | 0.03 | coral |
| SRR1734245 | beach sand metagenome | 80362 | 0 | 21 | NOB | 0.03 | marine sediment |
| ERR2113123 | metagenome | 122728 | 0 | 32 | NOB | 0.03 | marine sediment |
| SRR8359831 | coral reef metagenome | 50627 | 0 | 13 | NOB | 0.03 | coral |
| SRR2559786 | sponge metagenome | 23528 | 6 | 618 | AOA and NOB | 0.03 | sponge |
| ERR951486 | metagenome | 3952 | 0 | 1 | NOB | 0.03 | coral |
| ERR1419301 | marine metagenome | 67217 | 0 | 17 | NOB | 0.03 | other |
| SRR8359807 | coral reef metagenome | 11864 | 0 | 3 | NOB | 0.03 | coral |
| SRR1734260 | beach sand metagenome | 150977 | 0 | 38 | NOB | 0.03 | marine sediment |
| SRR3157219 | Cinachyrella | 3988 | 1 | 0 | AOA | 0.03 | sponge |
| SRR8380144 | Porites lobata | 3990 | 0 | 1 | NOB | 0.03 | coral |
| SRR1503494 | marine metagenome | 19968 | 0 | 5 | NOB | 0.03 | seawater |
| SRR8639939 | algae metagenome | 56575 | 0 | 14 | NOB | 0.02 | algae |
| SRR8359847 | coral reef metagenome | 40420 | 0 | 10 | NOB | 0.02 | coral |
| SRR4471363 | marine metagenome | 4080 | 0 | 1 | NOB | 0.02 | coral |
| SRR7034978 | marine metagenome | 114823 | 0 | 28 | NOB | 0.02 | marine sediment |
| SRR7595248 | marine metagenome | 114900 | 0 | 28 | NOB | 0.02 | marine sediment |
| ERR1419296 | marine metagenome | 152820 | 0 | 37 | NOB | 0.02 | other |
| SRR1734237 | beach sand metagenome | 86858 | 0 | 21 | NOB | 0.02 | marine sediment |
| SRR8751299 | coral metagenome | 8299 | 0 | 2 | NOB | 0.02 | coral |
| SRR5469070 | seawater metagenome | 116297 | 4 | 28 | AOA and NOB | 0.02 | seawater |
| ERR1419300 | marine metagenome | 145519 | 1 | 35 | AOA and NOB | 0.02 | other |
| SRR6457373 | marine metagenome | 29297 | 0 | 7 | NOB | 0.02 | seawater |
| SRR8359859 | coral reef metagenome | 37741 | 0 | 9 | NOB | 0.02 | coral |
| SRR2622335 | coral metagenome | 157308 | 0 | 37 | NOB | 0.02 | coral |
| SRR6457331 | marine metagenome | 72498 | 17 | 0 | AOA | 0.02 | seawater |
| SRR1734292 | beach sand metagenome | 98128 | 0 | 23 | NOB | 0.02 | marine sediment |
| SRR7357004 | coral metagenome | 38461 | 0 | 9 | NOB | 0.02 | coral |
| SRR869536 | sponge metagenome | 12876 | 0 | 3 | NOB | 0.02 | sponge |
| SRR7513592 | sponge metagenome | 73078 | 0 | 17 | NOB | 0.02 | sponge |
| SRR8491591 | metagenome | 30121 | 0 | 7 | NOB | 0.02 | coral |
| SRR7513594 | sponge metagenome | 55977 | 0 | 13 | NOB | 0.02 | sponge |
| SRR1653167 | bioreactor metagenome | 81845 | 0 | 19 | NOB | 0.02 | other |
| SRR2032948 | soil metagenome | 60558 | 0 | 14 | NOB | 0.02 | soil |
| SRR8640010 | algae metagenome | 78059 | 0 | 18 | NOB | 0.02 | algae |
| SRR3185597 | sand metagenome | 99968 | 0 | 23 | NOB | 0.02 | marine sediment |
| SRR7598544 | marine metagenome | 13046 | 0 | 3 | NOB | 0.02 | coral |
| SRR2087833 | plant metagenome | 156558 | 36 | 0 | AOA | 0.02 | plant |
| SRR7036586 | marine metagenome | 13055 | 0 | 3 | NOB | 0.02 | coral |
| SRR7513548 | sponge metagenome | 43930 | 0 | 10 | NOB | 0.02 | sponge |
| SRR5437928 | marine metagenome | 26460 | 1 | 6 | AOA and NOB | 0.02 | coral |
| SRR5150196 | marine metagenome | 106150 | 0 | 24 | NOB | 0.02 | seawater |
| SRR8640045 | algae metagenome | 48845 | 0 | 11 | NOB | 0.02 | algae |
| SRR6309411 | skin metagenome | 35526 | 0 | 8 | NOB | 0.02 | other |
| SRR8380148 | Porites lobata | 4442 | 0 | 1 | NOB | 0.02 | coral |
| SRR1734233 | aquatic metagenome | 80112 | 0 | 18 | NOB | 0.02 | marine sediment |
| SRR7513551 | sponge metagenome | 62478 | 0 | 14 | NOB | 0.02 | sponge |
| SRR8359895 | coral reef metagenome | 62513 | 0 | 14 | NOB | 0.02 | coral |
| SRR2559827 | sponge metagenome | 22355 | 5 | 427 | AOA and NOB | 0.02 | sponge |
| SRR8359832 | coral reef metagenome | 80513 | 0 | 18 | NOB | 0.02 | coral |
| SRR8359805 | coral reef metagenome | 8951 | 0 | 2 | NOB | 0.02 | coral |
| SRR7598621 | marine metagenome | 17957 | 0 | 4 | NOB | 0.02 | sponge |
| SRR7034647 | marine metagenome | 17964 | 0 | 4 | NOB | 0.02 | sponge |
| SRR8639975 | algae metagenome | 26948 | 0 | 6 | NOB | 0.02 | algae |
| SRR3113142 | aquatic metagenome | 229200 | 0 | 51 | NOB | 0.02 | seawater |
| SRR3204486 | coral metagenome | 63208 | 0 | 14 | NOB | 0.02 | coral |
| SRR7035173 | marine metagenome | 85859 | 0 | 19 | NOB | 0.02 | marine sediment |
| SRR7596945 | marine metagenome | 85961 | 0 | 19 | NOB | 0.02 | marine sediment |
| SRR2130166 | marine sediment metagenome | 27200 | 0 | 6 | NOB | 0.02 | marine sediment |
| SRR8359783 | coral reef metagenome | 13816 | 0 | 3 | NOB | 0.02 | coral |
| SRR6393354 | marine metagenome | 36989 | 0 | 8 | NOB | 0.02 | marine sediment |
| SRR1734273 | beach sand metagenome | 171127 | 0 | 37 | NOB | 0.02 | marine sediment |
| SRR8639942 | algae metagenome | 41815 | 0 | 9 | NOB | 0.02 | algae |
| SRR7595137 | marine metagenome | 107315 | 0 | 23 | NOB | 0.02 | marine sediment |
| SRR7513611 | sponge metagenome | 60811 | 0 | 13 | NOB | 0.02 | sponge |
| SRR8359809 | coral reef metagenome | 47067 | 0 | 10 | NOB | 0.02 | coral |
| ERR1801521 | marine metagenome | 9450 | 0 | 2 | NOB | 0.02 | other |
| SRR1734283 | beach sand metagenome | 157303 | 0 | 33 | NOB | 0.02 | marine sediment |
| SRR7513596 | marine metagenome | 47920 | 0 | 10 | NOB | 0.02 | seawater |
| SRR2994396 | coral metagenome | 9626 | 0 | 2 | NOB | 0.02 | coral |
| SRR5878977 | sediment metagenome | 24101 | 0 | 5 | NOB | 0.02 | marine sediment |
| SRR7598457 | marine metagenome | 9738 | 0 | 2 | NOB | 0.02 | coral |
| SRR3741977 | marine metagenome | 92806 | 19 | 0 | AOA | 0.02 | marine sediment |
| ERR173986 | symbiont metagenome | 4913 | 0 | 1 | NOB | 0.02 | sponge |
| SRR8661902 | coral metagenome | 24676 | 0 | 5 | NOB | 0.02 | coral |
| SRR6403426 | marine metagenome | 29617 | 0 | 6 | NOB | 0.02 | seawater |

| | | | | | | | |
|---|---|---|---|---|---|---|---|
| SRR7513612 | marine metagenome | 49878 | 0 | 10 | NOB | 0.02 | seawater |
| SRR2559785 | sponge metagenome | 34932 | 7 | 1042 | AOA and NOB | 0.02 | sponge |
| SRR4244589 | marine metagenome | 75013 | 0 | 15 | NOB | 0.02 | sponge |
| SRR1734285 | beach sand metagenome | 145355 | 0 | 29 | NOB | 0.02 | marine sediment |
| ERR1056334 | marine metagenome | 111074 | 0 | 22 | NOB | 0.02 | marine sediment |
| SRR8359786 | coral reef metagenome | 20258 | 0 | 4 | NOB | 0.02 | coral |
| SRR709460 | sponge metagenome | 55976 | 0 | 11 | NOB | 0.02 | sponge |
| SRR3185599 | sand metagenome | 96993 | 0 | 19 | NOB | 0.02 | marine sediment |
| SRR4471402 | marine metagenome | 5110 | 0 | 1 | NOB | 0.02 | coral |
| SRR5970253 | coral metagenome | 102443 | 0 | 20 | NOB | 0.02 | coral |
| SRR8111658 | mollusc metagenome | 5154 | 1 | 1 | AOA and NOB | 0.02 | other marine invertebrates |
| SRR8111658 | mollusc metagenome | 5154 | 1 | 1 | AOA and NOB | 0.02 | other marine invertebrates |
| SRR8661803 | coral metagenome | 20999 | 0 | 4 | NOB | 0.02 | coral |
| SRR7517643 | subsurface metagenome | 31559 | 0 | 6 | NOB | 0.02 | other |
| SRR3632952 | marine metagenome | 100535 | 19 | 15 | AOA and NOB | 0.02 | coral |
| SRR8111680 | mollusc metagenome | 5293 | 1 | 1 | AOA and NOB | 0.02 | other marine invertebrates |
| SRR8111680 | mollusc metagenome | 5293 | 1 | 1 | AOA and NOB | 0.02 | other marine invertebrates |
| SRR2917919 | Scleractinia | 1783850 | 1 | 337 | AOA and NOB | 0.02 | coral |
| SRR3624191 | marine metagenome | 100637 | 19 | 15 | AOA and NOB | 0.02 | coral |
| SRR8359773 | coral reef metagenome | 31821 | 0 | 6 | NOB | 0.02 | coral |
| SRR7591418 | coral metagenome | 5330 | 1 | 0 | AOA | 0.02 | coral |
| SRR1653249 | bioreactor metagenome | 85427 | 0 | 16 | NOB | 0.02 | other |
| SRR7034440 | marine metagenome | 10683 | 0 | 2 | NOB | 0.02 | marine sediment |
| SRR5878973 | sediment metagenome | 80376 | 0 | 15 | NOB | 0.02 | marine sediment |
| SRR7598550 | marine metagenome | 10717 | 0 | 2 | NOB | 0.02 | marine sediment |
| SRR8547997 | coral metagenome | 16209 | 3 | 3 | AOA and NOB | 0.02 | coral |
| SRR8547997 | coral metagenome | 16209 | 3 | 3 | AOA and NOB | 0.02 | coral |
| SRR5879073 | sediment metagenome | 38368 | 0 | 7 | NOB | 0.02 | marine sediment |
| SRR8359857 | coral reef metagenome | 32953 | 0 | 6 | NOB | 0.02 | coral |
| SRR5437923 | marine metagenome | 60477 | 11 | 448 | AOA and NOB | 0.02 | coral |
| SRR2559771 | sponge metagenome | 49911 | 9 | 2275 | AOA and NOB | 0.02 | sponge |
| ERR1894966 | metagenome | 463628 | 0 | 83 | NOB | 0.02 | other |
| ERR173991 | symbiont metagenome | 5632 | 0 | 1 | NOB | 0.02 | sponge |
| SRR7517638 | subsurface metagenome | 33807 | 0 | 6 | NOB | 0.02 | other |
| SRR7598551 | marine metagenome | 16945 | 0 | 3 | NOB | 0.02 | marine sediment |
| SRR7036492 | marine metagenome | 22710 | 0 | 4 | NOB | 0.02 | coral |
| SRR7598612 | marine metagenome | 22717 | 0 | 4 | NOB | 0.02 | coral |
| SRR5640865 | marine metagenome | 5701 | 1 | 0 | AOA | 0.02 | sponge |
| SRR7598228 | metagenome | 11461 | 0 | 2 | NOB | 0.02 | coral |
| SRR5469071 | seawater metagenome | 86163 | 15 | 95 | AOA and NOB | 0.02 | seawater |
| SRR2087834 | plant metagenome | 132388 | 485 | 23 | AOA and NOB | 0.02 | plant |
| SRR2559793 | sponge metagenome | 5768 | 1 | 76 | AOA and NOB | 0.02 | sponge |
| SRR7513614 | marine metagenome | 40399 | 0 | 7 | NOB | 0.02 | seawater |
| SRR7178014 | sponge metagenome | 104044 | 0 | 18 | NOB | 0.02 | sponge |
| SRR5437922 | marine metagenome | 34761 | 0 | 6 | NOB | 0.02 | coral |
| SRR7036444 | marine metagenome | 17422 | 0 | 3 | NOB | 0.02 | coral |
| SRR7598671 | marine metagenome | 17427 | 0 | 3 | NOB | 0.02 | coral |
| SRR7035563 | marine metagenome | 52410 | 0 | 9 | NOB | 0.02 | marine sediment |
| SRR7595857 | marine metagenome | 52629 | 0 | 9 | NOB | 0.02 | marine sediment |
| SRR6309440 | skin metagenome | 41468 | 0 | 7 | NOB | 0.02 | other |
| SRR8639960 | algae metagenome | 5944 | 0 | 1 | NOB | 0.02 | algae |
| ERR2403181 | metagenome | 47560 | 8 | 0 | AOA | 0.02 | other marine invertebrates |
| SRR4244545 | marine metagenome | 101178 | 17 | 86 | AOA and NOB | 0.02 | sponge |
| SRR3204451 | coral metagenome | 108422 | 0 | 18 | NOB | 0.02 | coral |
| SRR8491639 | metagenome | 18218 | 0 | 3 | NOB | 0.02 | coral |
| SRR5879042 | sediment metagenome | 97168 | 0 | 16 | NOB | 0.02 | marine sediment |
| SRR8359843 | coral reef metagenome | 30647 | 0 | 5 | NOB | 0.02 | coral |
| SRR3196999 | coral metagenome | 30695 | 0 | 5 | NOB | 0.02 | coral |
| ERR173985 | symbiont metagenome | 6145 | 0 | 1 | NOB | 0.02 | sponge |
| SRR2559799 | sponge metagenome | 24687 | 4 | 686 | AOA and NOB | 0.02 | sponge |
| SRR7595065 | marine metagenome | 105028 | 0 | 17 | NOB | 0.02 | marine sediment |
| SRR8359780 | coral reef metagenome | 86881 | 0 | 14 | NOB | 0.02 | coral |
| SRR1734270 | beach sand metagenome | 167943 | 0 | 27 | NOB | 0.02 | marine sediment |
| SRR4471395 | marine metagenome | 6227 | 0 | 1 | NOB | 0.02 | coral |
| SRR4471377 | marine metagenome | 6283 | 1 | 0 | AOA | 0.02 | coral |
| SRR8548009 | coral metagenome | 37809 | 6 | 0 | AOA | 0.02 | coral |
| SRR1030296 | sponge metagenome | 6324 | 0 | 1 | NOB | 0.02 | sponge |
| SRR3197006 | coral metagenome | 25330 | 0 | 4 | NOB | 0.02 | coral |
| SRR869542 | sponge metagenome | 19150 | 0 | 3 | NOB | 0.02 | sponge |
| SRR2087832 | plant metagenome | 274716 | 43 | 3 | AOA and NOB | 0.02 | plant |
| SRR7595639 | marine metagenome | 19291 | 0 | 3 | NOB | 0.02 | coral |
| SRR7036566 | marine metagenome | 19296 | 0 | 3 | NOB | 0.02 | coral |
| SRR8661903 | coral metagenome | 25928 | 0 | 4 | NOB | 0.02 | coral |
| SRR8359850 | coral reef metagenome | 39090 | 0 | 6 | NOB | 0.02 | coral |
| SRR8640070 | algae metagenome | 163084 | 0 | 25 | NOB | 0.02 | algae |
| SRR2040996 | aquatic metagenome | 6531 | 0 | 1 | NOB | 0.02 | seawater |
| SRR6081899 | sediment metagenome | 13075 | 0 | 2 | NOB | 0.02 | marine sediment |
| SRR2032906 | soil metagenome | 33053 | 0 | 5 | NOB | 0.02 | soil |
| SRR8639755 | algae metagenome | 132907 | 0 | 20 | NOB | 0.02 | algae |
| SRR3742053 | marine metagenome | 66647 | 10 | 0 | AOA | 0.02 | marine sediment |
| SRR3632952 | marine metagenome | 100535 | 19 | 15 | AOA and NOB | 0.01 | coral |
| SRR3624191 | marine metagenome | 100637 | 19 | 15 | AOA and NOB | 0.01 | coral |
| SRR2917918 | Scleractinia | 1718634 | 26 | 255 | AOA and NOB | 0.01 | coral |
| SRR2559791 | sponge metagenome | 20329 | 3 | 577 | AOA and NOB | 0.01 | sponge |
| SRR8751319 | coral metagenome | 6778 | 0 | 1 | NOB | 0.01 | coral |
| SRR7595421 | marine metagenome | 142690 | 0 | 21 | NOB | 0.01 | marine sediment |
| SRR8554642 | soil metagenome | 6579006 | 963 | 0 | AOA | 0.01 | soil |
| SRR7650181 | gut metagenome | 47910 | 0 | 7 | NOB | 0.01 | other |
| SRR1734259 | beach sand metagenome | 191967 | 0 | 28 | NOB | 0.01 | marine sediment |
| SRR4244549 | marine metagenome | 89268 | 13 | 192 | AOA and NOB | 0.01 | sponge |
| SRR3098353 | marine metagenome | 55566 | 0 | 8 | NOB | 0.01 | seawater |
| SRR7035000 | marine metagenome | 97405 | 0 | 14 | NOB | 0.01 | marine sediment |
| SRR7517642 | subsurface metagenome | 20873 | 0 | 3 | NOB | 0.01 | other |
| SRR7596929 | marine metagenome | 98020 | 0 | 14 | NOB | 0.01 | marine sediment |
| SRR3204478 | coral metagenome | 78006 | 191 | 11 | AOA and NOB | 0.01 | coral |
| SRR2559766 | sponge metagenome | 56992 | 8 | 1531 | AOA and NOB | 0.01 | sponge |
| SRR2559816 | sponge metagenome | 28575 | 4 | 821 | AOA and NOB | 0.01 | sponge |

| | | | | | | | |
|---|---|---|---|---|---|---|---|
| SRR7513554 | sponge metagenome | 57156 | 0 | 8 | NOB | 0.01 | sponge |
| SRR7036470 | marine metagenome | 7172 | 0 | 1 | NOB | 0.01 | coral |
| SRR7598445 | marine metagenome | 7185 | 0 | 1 | NOB | 0.01 | coral |
| SRR8639801 | algae metagenome | 93723 | 0 | 13 | NOB | 0.01 | algae |
| SRR7598277 | metagenome | 21650 | 0 | 3 | NOB | 0.01 | coral |
| SRR8359792 | coral reef metagenome | 43368 | 0 | 6 | NOB | 0.01 | coral |
| SRR5879053 | sediment metagenome | 28917 | 0 | 4 | NOB | 0.01 | marine sediment |
| SRR8751300 | coral metagenome | 7232 | 0 | 1 | NOB | 0.01 | coral |
| SRR8380111 | Porites rus | 14874 | 0 | 2 | NOB | 0.01 | coral |
| SRR5640878 | marine metagenome | 14875 | 2 | 0 | AOA | 0.01 | sponge |
| SRR8359794 | coral reef metagenome | 37192 | 0 | 5 | NOB | 0.01 | coral |
| SRR7035450 | marine metagenome | 143493 | 0 | 19 | NOB | 0.01 | coral |
| SRR7596450 | marine metagenome | 143736 | 0 | 19 | NOB | 0.01 | coral |
| ERR951482 | metagenome | 7581 | 0 | 1 | NOB | 0.01 | coral |
| SRR8224392 | seagrass metagenome | 53103 | 0 | 7 | NOB | 0.01 | seagrass |
| SRR7035157 | marine metagenome | 137599 | 0 | 18 | NOB | 0.01 | marine sediment |
| SRR8640017 | algae metagenome | 68919 | 0 | 9 | NOB | 0.01 | algae |
| SRR5877410 | sediment metagenome | 30708 | 0 | 4 | NOB | 0.01 | marine sediment |
| SRR7595069 | marine metagenome | 138402 | 0 | 18 | NOB | 0.01 | marine sediment |
| SRR6403216 | marine metagenome | 23109 | 0 | 3 | NOB | 0.01 | seawater |
| SRR8359880 | coral reef metagenome | 92482 | 0 | 12 | NOB | 0.01 | coral |
| SRR8639945 | algae metagenome | 38751 | 0 | 5 | NOB | 0.01 | algae |
| SRR8639767 | algae metagenome | 54371 | 0 | 7 | NOB | 0.01 | algae |
| SRR5150268 | marine metagenome | 31110 | 0 | 4 | NOB | 0.01 | seawater |
| SRR869535 | sponge metagenome | 7782 | 0 | 1 | NOB | 0.01 | sponge |
| ERR2403183 | metagenome | 39116 | 5 | 0 | AOA | 0.01 | other marine invertebrates |
| SRR7178016 | sponge metagenome | 180333 | 0 | 23 | NOB | 0.01 | sponge |
| SRR3234039 | aquatic metagenome | 307795 | 39 | 0 | AOA | 0.01 | other |
| SRR7598622 | marine metagenome | 23711 | 0 | 3 | NOB | 0.01 | sponge |
| SRR7034515 | marine metagenome | 23719 | 0 | 3 | NOB | 0.01 | sponge |
| SRR8639896 | algae metagenome | 79338 | 0 | 10 | NOB | 0.01 | algae |
| SRR1653120 | bioreactor metagenome | 79877 | 0 | 10 | NOB | 0.01 | other |
| SRR7596721 | marine metagenome | 40042 | 0 | 5 | NOB | 0.01 | marine sediment |
| SRR2559789 | sponge metagenome | 40045 | 5 | 1456 | AOA and NOB | 0.01 | sponge |
| SRR8359852 | coral reef metagenome | 56081 | 0 | 7 | NOB | 0.01 | coral |
| SRR8751298 | coral metagenome | 8015 | 0 | 1 | NOB | 0.01 | coral |
| ERR330035 | sediment metagenome | 56178 | 0 | 7 | NOB | 0.01 | marine sediment |
| SRR6401427 | marine metagenome | 90276 | 0 | 11 | NOB | 0.01 | sponge |
| SRR7513604 | sponge metagenome | 90308 | 0 | 11 | NOB | 0.01 | sponge |
| SRR3136160 | bioreactor metagenome | 8309 | 0 | 1 | NOB | 0.01 | other |
| SRR8639944 | algae metagenome | 41664 | 0 | 5 | NOB | 0.01 | algae |
| SRR8751263 | coral metagenome | 8388 | 0 | 1 | NOB | 0.01 | coral |
| SRR1503495 | marine metagenome | 41993 | 0 | 5 | NOB | 0.01 | seawater |
| SRR2559802 | sponge metagenome | 8412 | 1 | 277 | AOA and NOB | 0.01 | sponge |
| SRR5437921 | marine metagenome | 33752 | 0 | 4 | NOB | 0.01 | coral |
| SRR2559795 | sponge metagenome | 16951 | 2 | 849 | AOA and NOB | 0.01 | sponge |
| SRR8491587 | metagenome | 17131 | 0 | 2 | NOB | 0.01 | coral |
| SRR8111684 | mollusc metagenome | 8668 | 0 | 1 | NOB | 0.01 | other marine invertebrates |
| SRR5878988 | sediment metagenome | 17377 | 0 | 2 | NOB | 0.01 | marine sediment |
| SRR1593663 | coral metagenome | 183336 | 9745 | 21 | AOA and NOB | 0.01 | coral |
| SRR7595081 | marine metagenome | 104870 | 0 | 12 | NOB | 0.01 | marine sediment |
| SRR7035083 | marine metagenome | 105347 | 0 | 12 | NOB | 0.01 | marine sediment |
| ERR1894959 | metagenome | 527242 | 0 | 60 | NOB | 0.01 | other |
| SRR1734241 | beach sand metagenome | 87912 | 0 | 10 | NOB | 0.01 | marine sediment |
| SRR2041115 | aquatic metagenome | 26374 | 0 | 3 | NOB | 0.01 | seawater |
| SRR1927881 | seawater metagenome | 53317 | 5 | 6 | AOA and NOB | 0.01 | seawater |
| SRR2559769 | sponge metagenome | 35658 | 4 | 874 | AOA and NOB | 0.01 | sponge |
| SRR8243337 | coral reef metagenome | 26822 | 0 | 3 | NOB | 0.01 | coral |
| SRR7513593 | sponge metagenome | 53692 | 0 | 6 | NOB | 0.01 | sponge |
| SRR7598260 | metagenome | 17962 | 0 | 2 | NOB | 0.01 | coral |
| SRR709454 | marine metagenome | 45414 | 0 | 5 | NOB | 0.01 | marine benthic and pelagic environments |
| ERR1056312 | marine metagenome | 55049 | 0 | 6 | NOB | 0.01 | marine sediment |
| ERR1056318 | marine metagenome | 248004 | 0 | 27 | NOB | 0.01 | marine sediment |
| SRR1593662 | coral metagenome | 158298 | 0 | 17 | NOB | 0.01 | coral |
| SRR7596848 | marine metagenome | 28061 | 0 | 3 | NOB | 0.01 | coral |
| SRR8359777 | coral reef metagenome | 112315 | 0 | 12 | NOB | 0.01 | coral |
| SRR8359856 | coral reef metagenome | 37779 | 0 | 4 | NOB | 0.01 | coral |
| SRR8639768 | algae metagenome | 28353 | 0 | 3 | NOB | 0.01 | algae |
| SRR7513549 | sponge metagenome | 56776 | 0 | 6 | NOB | 0.01 | sponge |
| SRR7034717 | marine metagenome | 9497 | 0 | 1 | NOB | 0.01 | marine sediment |
| SRR7598379 | marine metagenome | 9508 | 0 | 1 | NOB | 0.01 | marine sediment |
| SRR3091958 | coral metagenome | 152587 | 0 | 16 | NOB | 0.01 | coral |
| ERR951470 | metagenome | 9555 | 0 | 1 | NOB | 0.01 | coral |
| SRR6309405 | skin metagenome | 38715 | 0 | 4 | NOB | 0.01 | other |
| SRR1565475 | sponge metagenome | 19549 | 2 | 110 | AOA and NOB | 0.01 | sponge |
| ERR2906734 | Methanobacterium beijingense | 69285 | 0 | 7 | NOB | 0.01 | other |
| SRR3633253 | marine metagenome | 198279 | 0 | 20 | NOB | 0.01 | coral |
| SRR7598291 | metagenome | 19898 | 0 | 2 | NOB | 0.01 | coral |
| SRR7513550 | sponge metagenome | 49891 | 0 | 5 | NOB | 0.01 | sponge |

**Table S3:** Dissolved inorganic nitrogen (DIN) concentrations measured in the individual sponge and abiotic control incubation jars over time.

| Sample | individual # | treatment | group | time | sponge wet weight | NO2- produced [µM/L] | NO3- produced [µM/L] | NH4+ produced [µM/L] | NO2- produced per | NO3- produced per | NH4+ produced per |
|---|---|---|---|---|---|---|---|---|---|---|---|
| 0h-1-1-S | 1 | ambient | sponge | 0h | 36.19 | 0.035 | 1.990 | 0.175 | 0.001 | 0.055 | 0.005 |
| 1h-1-1-S | 1 | ambient | sponge | 1h | 36.19 | 0.040 | 7.945 | 0.805 | 0.001 | 0.220 | 0.022 |
| 3h-1-1-S | 1 | ambient | sponge | 3h | 36.19 | 0.045 | 13.654 | 1.015 | 0.001 | 0.377 | 0.028 |
| 6h-1-1-S | 1 | ambient | sponge | 6h | 36.19 | 0.045 | 24.137 | 1.437 | 0.001 | 0.667 | 0.040 |
| 0h-1-2-S | 2 | ambient | sponge | 0h | 17.79 | 0.035 | 1.960 | 0.110 | 0.002 | 0.110 | 0.006 |
| 1h-1-2-S | 2 | ambient | sponge | 1h | 17.79 | 0.045 | 3.920 | 1.070 | 0.003 | 0.220 | 0.060 |
| 3h-1-2-S | 2 | ambient | sponge | 3h | 17.79 | 0.045 | 5.841 | 1.865 | 0.003 | 0.328 | 0.105 |
| 6h-1-2-S | 2 | ambient | sponge | 6h | 17.79 | 0.045 | 9.902 | 3.385 | 0.003 | 0.557 | 0.190 |
| 0h-1-3-S | 3 | ambient | sponge | 0h | 31.71 | 0.040 | 2.030 | 0.130 | 0.001 | 0.064 | 0.004 |
| 1h-1-3-S | 3 | ambient | sponge | 1h | 31.71 | 0.045 | 7.440 | 1.415 | 0.001 | 0.235 | 0.045 |
| 3h-1-3-S | 3 | ambient | sponge | 3h | 31.71 | 0.040 | 13.373 | 2.200 | 0.001 | 0.422 | 0.069 |
| 6h-1-3-S | 3 | ambient | sponge | 6h | 31.71 | 0.073 | 25.143 | 4.238 | 0.002 | 0.793 | 0.134 |
| 0h-1-1-A | 4 | ambient | abiotic | 0h | 0 | 0.025 | 1.785 | 0.060 | NA | NA | NA |
| 1h-1-1-A | 4 | ambient | abiotic | 1h | 0 | 0.030 | 1.930 | 0.170 | NA | NA | NA |
| 3h-1-1-A | 4 | ambient | abiotic | 3h | 0 | 0.030 | 1.925 | 0.350 | NA | NA | NA |
| 6h-1-1-A | 4 | ambient | abiotic | 6h | 0 | 0.025 | 1.977 | 0.289 | NA | NA | NA |
| 0h-1-2-A | 5 | ambient | abiotic | 0h | 0 | 0.035 | 1.990 | 0.060 | NA | NA | NA |
| 1h-1-2-A | 5 | ambient | abiotic | 1h | 0 | 0.040 | 2.090 | 0.100 | NA | NA | NA |
| 3h-1-2-A | 5 | ambient | abiotic | 3h | 0 | 0.050 | 2.075 | 0.105 | NA | NA | NA |
| 6h-1-2-A | 5 | ambient | abiotic | 6h | 0 | 0.040 | 1.900 | 0.167 | NA | NA | NA |
| 0h-1-3-A | 6 | ambient | abiotic | 0h | 0 | 0.030 | 1.780 | 0.060 | NA | NA | NA |
| 1h-1-3-A | 6 | ambient | abiotic | 1h | 0 | 0.030 | 1.890 | 0.095 | NA | NA | NA |
| 3h-1-3-A | 6 | ambient | abiotic | 3h | 0 | 0.025 | 1.895 | 0.144 | NA | NA | NA |
| 6h-1-3-A | 6 | ambient | abiotic | 6h | 0 | 0.030 | 1.919 | 0.139 | NA | NA | NA |
| 0h-2-1-S | 1 | 5uM | sponge | 0h | 33.29 | 0.030 | 1.780 | 5.100 | 0.001 | 0.053 | 0.153 |
| 1h-2-1-S | 1 | 5uM | sponge | 1h | 33.29 | 0.035 | 5.720 | 3.260 | 0.001 | 0.172 | 0.098 |
| 3h-2-1-S | 1 | 5uM | sponge | 3h | 33.29 | 0.040 | 11.146 | 3.221 | 0.001 | 0.335 | 0.097 |
| 6h-2-1-S | 1 | 5uM | sponge | 6h | 33.29 | 0.054 | 20.646 | 4.309 | 0.002 | 0.620 | 0.129 |
| 0h-2-2-S | 2 | 5uM | sponge | 0h | 22.41 | 0.040 | 1.955 | 4.280 | 0.002 | 0.087 | 0.191 |
| 1h-2-2-S | 2 | 5uM | sponge | 1h | 22.41 | 0.080 | 5.230 | 4.515 | 0.004 | 0.233 | 0.201 |
| 3h-2-2-S | 2 | 5uM | sponge | 3h | 22.41 | 0.095 | 10.359 | 4.271 | 0.004 | 0.462 | 0.191 |
| 6h-2-2-S | 2 | 5uM | sponge | 6h | 22.41 | 0.133 | 19.198 | 4.770 | 0.006 | 0.857 | 0.213 |
| 0h-2-3-S | 3 | 5uM | sponge | 0h | 27.17 | 0.030 | 1.795 | 4.750 | 0.001 | 0.066 | 0.175 |
| 1h-2-3-S | 3 | 5uM | sponge | 1h | 27.17 | 0.070 | 5.885 | 3.990 | 0.003 | 0.217 | 0.147 |
| 3h-2-3-S | 3 | 5uM | sponge | 3h | 27.17 | 0.090 | 11.506 | 3.839 | 0.003 | 0.423 | 0.141 |
| 6h-2-3-S | 3 | 5uM | sponge | 6h | 27.17 | 0.161 | 19.134 | 5.411 | 0.006 | 0.704 | 0.199 |
| 0h-2-1-A | 4 | 5uM | abiotic | 0h | 0 | 0.040 | 1.975 | 4.360 | NA | NA | NA |
| 1h-2-1-A | 4 | 5uM | abiotic | 1h | 0 | 0.050 | 2.075 | 5.065 | NA | NA | NA |
| 3h-2-1-A | 4 | 5uM | abiotic | 3h | 0 | 0.079 | 2.031 | 5.011 | NA | NA | NA |
| 6h-2-1-A | 4 | 5uM | abiotic | 6h | 0 | 0.041 | 1.979 | 5.106 | NA | NA | NA |
| 0h-2-2-A | 5 | 5uM | abiotic | 0h | 0 | 0.030 | 1.790 | 4.845 | NA | NA | NA |
| 1h-2-2-A | 5 | 5uM | abiotic | 1h | 0 | 0.035 | 1.890 | 5.035 | NA | NA | NA |
| 3h-2-2-A | 5 | 5uM | abiotic | 3h | 0 | 0.064 | 1.861 | 5.020 | NA | NA | NA |
| 6h-2-2-A | 5 | 5uM | abiotic | 6h | 0 | 0.026 | 1.785 | 4.925 | NA | NA | NA |
| 0h-2-3-A | 6 | 5uM | abiotic | 0h | 0 | 0.030 | 1.780 | 4.875 | NA | NA | NA |
| 1h-2-3-A | 6 | 5uM | abiotic | 1h | 0 | 0.030 | 1.895 | 5.015 | NA | NA | NA |
| 3h-2-3-A | 6 | 5uM | abiotic | 3h | 0 | 0.025 | 1.915 | 5.010 | NA | NA | NA |
| 6h-2-3-A | 6 | 5uM | abiotic | 6h | 0 | 0.025 | 1.782 | 4.858 | NA | NA | NA |
| 0h-3-1-S | 1 | 25uM | sponge | 0h | 24.23 | 0.040 | 1.970 | 26.580 | 0.002 | 0.081 | 1.097 |
| 1h-3-1-S | 1 | 25uM | sponge | 1h | 24.23 | 0.055 | 10.290 | 17.860 | 0.002 | 0.425 | 0.737 |
| 3h-3-1-S | 1 | 25uM | sponge | 3h | 24.23 | 0.055 | 19.660 | 9.865 | 0.002 | 0.811 | 0.407 |
| 6h-3-1-S | 1 | 25uM | sponge | 6h | 24.23 | 0.060 | 28.134 | 5.172 | 0.002 | 1.161 | 0.213 |
| 0h-3-2-S | 2 | 25uM | sponge | 0h | 30.4 | 0.045 | 1.965 | 28.655 | 0.001 | 0.065 | 0.943 |
| 1h-3-2-S | 2 | 25uM | sponge | 1h | 30.4 | 0.035 | 6.425 | 15.600 | 0.001 | 0.211 | 0.513 |
| 3h-3-2-S | 2 | 25uM | sponge | 3h | 30.4 | 0.040 | 11.480 | 14.503 | 0.001 | 0.378 | 0.477 |
| 6h-3-2-S | 2 | 25uM | sponge | 6h | 30.4 | 0.045 | 22.638 | 13.278 | 0.001 | 0.745 | 0.437 |
| 0h-3-3-S | 3 | 25uM | sponge | 0h | 31.97 | 0.045 | 1.965 | 30.405 | 0.001 | 0.061 | 0.951 |
| 1h-3-3-S | 3 | 25uM | sponge | 1h | 31.97 | 0.055 | 6.750 | 17.150 | 0.002 | 0.211 | 0.536 |
| 3h-3-3-S | 3 | 25uM | sponge | 3h | 31.97 | 0.050 | 12.980 | 15.332 | 0.002 | 0.406 | 0.480 |
| 6h-3-3-S | 3 | 25uM | sponge | 6h | 31.97 | 0.055 | 26.485 | 14.239 | 0.002 | 0.828 | 0.445 |
| 0h-3-1-A | 4 | 25uM | abiotic | 0h | 0 | 0.020 | 1.710 | 27.720 | NA | NA | NA |
| 1h-3-1-A | 4 | 25uM | abiotic | 1h | 0 | 0.040 | 1.890 | 27.190 | NA | NA | NA |
| 3h-3-1-A | 4 | 25uM | abiotic | 3h | 0 | 0.030 | 1.866 | 26.407 | NA | NA | NA |
| 6h-3-1-A | 4 | 25uM | abiotic | 6h | 0 | 0.025 | 1.828 | 26.440 | NA | NA | NA |
| 0h-3-2-A | 5 | 25uM | abiotic | 0h | 0 | 0.020 | 1.740 | 30.080 | NA | NA | NA |
| 1h-3-2-A | 5 | 25uM | abiotic | 1h | 0 | 0.040 | 1.890 | 28.180 | NA | NA | NA |
| 3h-3-2-A | 5 | 25uM | abiotic | 3h | 0 | 0.030 | 1.866 | 25.863 | NA | NA | NA |
| 6h-3-2-A | 5 | 25uM | abiotic | 6h | 0 | 0.025 | 1.800 | 26.286 | NA | NA | NA |
| 0h-3-3-A | 6 | 25uM | abiotic | 0h | 0 | 0.030 | 1.900 | 23.680 | NA | NA | NA |
| 1h-3-3-A | 6 | 25uM | abiotic | 1h | 0 | 0.060 | 2.090 | 25.820 | NA | NA | NA |
| 3h-3-3-A | 6 | 25uM | abiotic | 3h | 0 | 0.045 | 2.040 | 26.970 | NA | NA | NA |
| 6h-3-3-A | 6 | 25uM | abiotic | 6h | 0 | 0.040 | 1.964 | 25.768 | NA | NA | NA |

**Table S4:** GTDB-tk output of the recovered ammonia-oxidizing archaeal (AOA) and nitrite-oxidizing bacterial (NOB) metagenome assembled genomes (MAG) recovered in this study.

| | AOA MAG | NOB MAG |
|---|---|---|
| **proposed name** | *Ca* . Nitrosokoinonia keratosae gen. nov. sp. nov. | *Ca* . Nitrosymbion coscinodermae gen. nov. sp. nov |
| **MAG_ID** | metabat_nocov.1 | metabat_nocov.4 |
| **classification** | d__Archaea;p__Thermoproteota;c__Nitrososphaeria;o__Nitrososphaerales; | d__Bacteria;p__Nitrospirota;c__Nitrospiria;o__Nitrospirales; |
| **fastani_reference** | GCA_009843815.1 | GCA_011523385.1 |
| **fastani_reference_radius** | 95 | 95 |
| **fastani_taxonomy** | d__Archaea;p__Thermoproteota;c__Nitrososphaeria;o__Nitrososphaerales; | d__Bacteria;p__Nitrospirota;c__Nitrospiria;o__Nitrospirales; |
| **fastani_ani** | 97.47 | 97.9 |
| **fastani_af** | 0.901 | 0.899 |
| **closest_placement_reference** | GCA_009843815.1 | GCA_011523385.1 |
| **closest_placement_radius** | 95 | 95 |
| **closest_placement_taxonomy** | d__Archaea;p__Thermoproteota;c__Nitrososphaeria;o__Nitrososphaerales; | d__Bacteria;p__Nitrospirota;c__Nitrospiria;o__Nitrospirales; |
| **closest_placement_ani** | 97.47 | 97.9 |
| **closest_placement_af** | 0.901 | 0.899 |
| **pplacer_taxonomy** | d__Archaea;p__Thermoproteota;c__Nitrososphaeria;o__Nitrososphaerales; | d__Bacteria;p__Nitrospirota;c__Nitrospiria;o__Nitrospirales; |
| **classification_method** | taxonomic classification defined by topology and ANI | taxonomic classification defined by topology and ANI |
| **note** | topological placement and ANI have congruent species assignments | topological placement and ANI have congruent species assignments |
| **other_related_references** | GCA_009840065.1, s__VYCS01 sp009840065, 95.0, 79.78, 0.589; | GCA_009842645.1, s__Bin75 sp009842645, 95.0, 89.93, 0.811; |
| **msa_percent** | 80.92 | 84.57 |
| **translation_table** | 11 | 11 |
| **red_value** | N/A | N/A |
| **warnings** | N/A | N/A |

**Table S5:** Unique orthologs (eggNOG OGs) associated with genomes of *Ca* . Nitrosokoinonia and shared between other sponge-associated AOA and *Ca* . Nitrosokoinonia.

| eggNOG OG (root) | other sponge-associated AOA (n=8) | free-living (n=29) | Ca. Nitrosokoinonia sp. (n=5) | Description |
|---|---|---|---|---|
| COG0663@1\|root | 0 | 0 | 4 | COG0663 Carbonic anhydrases acetyltransferases, isoleucine patch superfamily |
| COG1043@1\|root | 0 | 0 | 4 | **Involved in the biosynthesis of lipid A, a phosphorylated glycolipid that anchors the lipopolysaccharide to the outer membrane of the cell** |
| COG3592@1\|root | 0 | 0 | 4 | Divergent 4Fe-4S mono-cluster |
| 2BVTZ@1\|root | 0 | 0 | 3 | - |
| arCOG04936@1\|root | 0 | 0 | 3 | Glycosyl transferase family 2 |
| arCOG02754@1\|root | 0 | 0 | 2 | SPTR Transposase, IS4 family protein |
| COG0190@1\|root | 0 | 0 | 2 | Catalyzes the oxidation of 5,10-methylenetetrahydrofolate to 5,10-methenyltetrahydrofolate and then the hydrolysis of 5,10-methenyltetrahydrofolate to 10- formyltetrahydrofolate |
| COG1373@1\|root | 0 | 0 | 2 | AAA domain |
| COG1487@1\|root | 0 | 0 | 2 | nuclease activity |
| COG1889@1\|root | 0 | 0 | 2 | **rRNA 2'-O-methyltransferase fibrillarin** |
| COG2944@1\|root | 0 | 0 | 2 | sequence-specific DNA binding |
| COG3039@1\|root | 0 | 0 | 2 | Transposase |
| COG3316@1\|root | 0 | 0 | 2 | DDE domain |
| COG3676@1\|root | 0 | 0 | 2 | ISXO2-like transposase domain |
| KOG3539@1\|root | 0 | 0 | 2 | **Spondin_N** |
| 28H5D@1\|root | 0 | 0 | 1 | **Restriction endonuclease EcoRV** |
| 28HBW@1\|root | 0 | 0 | 1 | Domain of unknown function (DUF1998) |
| 28JGZ@1\|root | 0 | 0 | 1 | - |
| 28JT9@1\|root | 0 | 0 | 1 | Domain of unknown function (DUF4338) |
| 28KC2@1\|root | 0 | 0 | 1 | CRISPR-associated protein GSU0053 (Cas_GSU0053) |
| 2B1FT@1\|root | 0 | 0 | 1 | - |
| 2B1Z8@1\|root | 0 | 0 | 1 | **TIR domain** |
| 2C5PK@1\|root | 0 | 0 | 1 | **MTH538 TIR-like domain (DUF1863)** |
| 2DBGM@1\|root | 0 | 0 | 1 | **TIR domain** |
| 2DBSW@1\|root | 0 | 0 | 1 | PFAM CRISPR-associated protein, GSU0054 family (Cas_GSU0054) |
| 2F0GY@1\|root | 0 | 0 | 1 | Domain of unknown function (DUF4868) |
| 4QD0J@10239\|Viruses | 0 | 0 | 1 | - |
| arCOG06565@1\|root | 0 | 0 | 1 | - |
| COG1502@1\|root | 0 | 0 | 1 | Catalyzes the reversible phosphatidyl group transfer from one phosphatidylglycerol molecule to another to form cardiolipin (CL) (diphosphatidylglycerol) and glycerol |
| COG2230@1\|root | 0 | 0 | 1 | Cyclopropane fatty acid synthase |

| | | | | |
|---|---|---|---|---|
| COG2865@1\|root | 0 | 0 | 1 | Transcriptional regulator |
| COG2907@1\|root | 0 | 0 | 1 | Flavin containing amine oxidoreductase |
| COG3325@1\|root | 0 | 0 | 1 | PFAM chitin-binding domain 3 protein |
| KOG0908@1\|root | 0 | 0 | 1 | PITH domain |
| COG2801@1\|root | 3 | 0 | 5 | transposase |
| COG2176@1\|root | 1 | 0 | 5 | Exonuclease |
| COG4279@1\|root | 2 | 0 | 4 | zinc finger |
| COG3677@1\|root | 1 | 0 | 4 | Transposase and inactivated derivatives |
| COG1432@1\|root | 2 | 0 | 3 | OST-HTH/LOTUS domain |
| COG0323@1\|root | 2 | 0 | 2 | Histidine kinase-, DNA gyrase B-, and HSP90-like ATPase |
| COG2963@1\|root | 2 | 0 | 2 | COG2963, Transposase and inactivated derivatives |
| 28MTH@1\|root | 1 | 0 | 2 | Protein of unknown function (DUF4007) |
| COG2337@1\|root | 1 | 0 | 2 | PemK-like, MazF-like toxin of type II toxin-antitoxin system |
| COG4412@1\|root | 1 | 0 | 2 | peptidase activity, acting on L-amino acid peptides |
| COG0616@1\|root | 4 | 0 | 1 | Serine dehydrogenase proteinase |
| COG1343@1\|root | 3 | 0 | 1 | CRISPR (clustered regularly interspaced short palindromic repeat), is an adaptive immune system that provides protection against mobile genetic elements (viruses, transposable elements and conjugative plasmids). |
| COG0078@1\|root | 2 | 0 | 1 | Reversibly catalyzes the transfer of the carbamoyl group from carbamoyl phosphate (CP) to the N(epsilon) atom of ornithine (ORN) to produce L-citrulline |
| COG1203@1\|root | 2 | 0 | 1 | helicase superfamily c-terminal domain |
| COG2062@1\|root | 2 | 0 | 1 | PFAM Phosphoglycerate mutase family |
| 28IJY@1\|root | 1 | 0 | 1 | CRISPR-associated protein GSU0053 (Cas_GSU0053) |
| 28KAG@1\|root | 1 | 0 | 1 | Abortive infection C-terminus |
| 29YK4@1\|root | 1 | 0 | 1 | Type II restriction enzyme SfiI |
| COG0210@1\|root | 1 | 0 | 1 | PFAM UvrD REP helicase |
| COG0631@1\|root | 1 | 0 | 1 | protein phosphatase 2C domain protein |
| COG1132@1\|root | 1 | 0 | 1 | ABC transporter |
| COG1401@1\|root | 1 | 0 | 1 | GTPase subunit of restriction endonuclease |
| COG2267@1\|root | 1 | 0 | 1 | Alpha beta hydrolase |
| COG2856@1\|root | 1 | 0 | 1 | Pfam:DUF955 |
| COG3335@1\|root | 1 | 0 | 1 | DDE superfamily endonuclease |
| COG3410@1\|root | 1 | 0 | 1 | Uncharacterized conserved protein (DUF2075) |
| COG3481@1\|root | 1 | 0 | 1 | nucleic acid binding OB-fold tRNA helicase-type |
| COG4268@1\|root | 1 | 0 | 1 | McrBC 5-methylcytosine restriction system component |
| COG4886@1\|root | 1 | 0 | 1 | Mycoplasma protein of unknown function, DUF285 |

**Table S6:** Unique orthologs (eggNOG OGs) associated with genomes of *Ca* . Nitrosymbion.

| eggNOG OG (root) | Ca. Nitrosymbion (n=12) | IVa (n=9) | Ivb (n=4) | Description provided by eggNOG |
|---|---|---|---|---|
| 28NHW@1\|root | 10 | 0 | 0 | - |
| COG4147@1\|root | 10 | 0 | 0 | **Sodium:solute symporter family** |
| COG4430@1\|root | 10 | 0 | 0 | **Bacteriocin-protection, YdeI or OmpD-Associated** |
| 2BE1Z@1\|root | 9 | 0 | 0 | - |
| 2DB8Y@1\|root | 9 | 0 | 0 | Uncharacterized protein conserved in bacteria (DUF2236) |
| 2DFN7@1\|root | 9 | 0 | 0 | - |
| 2DT5B@1\|root | 9 | 0 | 0 | - |
| COG4327@1\|root | 9 | 0 | 0 | **solute sodium symporter, small subunit; membrane; membrane; Domain of unknown function (DUF4212)** |
| 28H5T@1\|root | 8 | 0 | 0 | - |
| 2EDCN@1\|root | 8 | 0 | 0 | - |
| 2EJAI@1\|root | 8 | 0 | 0 | - |
| COG1432@1\|root | 8 | 0 | 0 | NYN domain |
| COG3514@1\|root | 8 | 0 | 0 | **BrnA antitoxin of type II toxin-antitoxin system** |
| COG3943@1\|root | 8 | 0 | 0 | **COG3943 Virulence protein; Virulence protein RhuM family, Toxin-antitoxin system, toxin component, Fic** |
| 298AC@1\|root | 7 | 0 | 0 | **Antitoxin Phd_YefM, type II toxin-antitoxin system; Antitoxin Phd_YefM, type II toxin-antitoxin system** |
| 2BF9B@1\|root | 7 | 0 | 0 | Cytochrome c554 and c-prime; Tetraheme-Cyt. c |
| 2D74M@1\|root | 7 | 0 | 0 | - |
| 2DETT@1\|root | 7 | 0 | 0 | - |
| 2DGZ3@1\|root | 7 | 0 | 0 | **Antitoxin Phd_YefM, type II toxin-antitoxin system** |
| 2DR8X@1\|root | 7 | 0 | 0 | -; -; - |
| 2EF1J@1\|root | 7 | 0 | 0 | - |
| COG3077@1\|root | 7 | 0 | 0 | **Addiction module antitoxin, RelB DinJ family; RelB antitoxin; bacterial-type proximal promoter sequence-specific DNA binding; RelB antitoxin** |
| COG3248@1\|root | 7 | 0 | 0 | **Nucleoside-binding outer membrane; Nucleoside-binding outer membrane protein; Nucleoside-binding outer membrane** |
| COG3742@1\|root | 7 | 0 | 0 | **Toxic component of a toxin-antitoxin (TA) module. An Rnase** |
| COG4422@1\|root | 7 | 0 | 0 | Protein of unknown function (DUF5131) |
| COG4423@1\|root | 7 | 0 | 0 | **PFAM Rv0623 family protein transcription factor** |
| 2A6UU@1\|root | 6 | 0 | 0 | -; - |
| 2DNRA@1\|root | 6 | 0 | 0 | -; - |
| 2EKGF@1\|root | 6 | 0 | 0 | Domain of unknown function (DUF1902) |
| arCOG14015@1\|root | 6 | 0 | 0 | -; -; - |
| COG2405@1\|root | 6 | 0 | 0 | nucleic acid-binding protein contains PIN domain |
| COG3240@1\|root | 6 | 0 | 0 | esterase |
| COG3324@1\|root | 6 | 0 | 0 | glyoxalase bleomycin resistance protein dioxygenase; enzyme related to lactoylglutathione lyase; translation initiation factor activity; translation initiation factor activity |
| COG5499@1\|root | 6 | 0 | 0 | transcriptional regulator; regulator; regulator; transcription regulator containing HTH domain; Helix-turn-helix XRE-family like proteins |
| COG5550@1\|root | 6 | 0 | 0 | -; - |
| 28HGF@1\|root | 5 | 0 | 0 | **PFAM TraU** |
| 28IG0@1\|root | 5 | 0 | 0 | -; -; -; - |
| 28IWY@1\|root | 5 | 0 | 0 | -; -; - |
| 28JT9@1\|root | 5 | 0 | 0 | Domain of unknown function (DUF4338); Domain of unknown function (DUF4338); Domain of unknown function (DUF4338) |
| 28MTB@1\|root | 5 | 0 | 0 | PFAM Conjugative relaxosome accessory transposon protein |
| 28N2S@1\|root | 5 | 0 | 0 | P63C domain; P63C domain; P63C domain; P63C domain |
| 28PQH@1\|root | 5 | 0 | 0 | - |
| 299JR@1\|root | 5 | 0 | 0 | HicA toxin of bacterial toxin-antitoxin, |
| 29JYC@1\|root | 5 | 0 | 0 | Antitoxin Phd_YefM, type II toxin-antitoxin system |
| 2ASHK@1\|root | 5 | 0 | 0 | - |
| 2B9NZ@1\|root | 5 | 0 | 0 | Protein of unknown function (DUF1579) |
| 2CDV1@1\|root | 5 | 0 | 0 | Type II site-specific deoxyribonuclease |
| 2CEYG@1\|root | 5 | 0 | 0 | Ribbon-helix-helix domain; Ribbon-helix-helix domain; Ribbon-helix-helix domain |

| | | | | |
|---|---|---|---|---|
| 2CGUX@1\|root | 5 | 0 | 0 | Type-F conjugative transfer system protein TraW |
| 2CK24@1\|root | 5 | 0 | 0 | - |
| 2DPTZ@1\|root | 5 | 0 | 0 | -; -; - |
| 2DPXT@1\|root | 5 | 0 | 0 | - |
| 2EA3G@1\|root | 5 | 0 | 0 | Domain of unknown function (DUF4177) |
| 2EKK5@1\|root | 5 | 0 | 0 | Putative addiction module component |
| COG1662@1\|root | 5 | 0 | 0 | PFAM IS1 transposase; PFAM IS1 transposase; PFAM IS1 transposase |
| COG3041@1\|root | 5 | 0 | 0 | Bacterial toxin of type II toxin-antitoxin system, YafQ; Bacterial toxin of type II toxin-antitoxin system, YafQ; Bacterial toxin of type II toxin-antitoxin system, YafQ; Bacterial toxin of type II toxin-antitoxin system, YafQ |
| COG3306@1\|root | 5 | 0 | 0 | Glycosyltransferase family 25 (LPS biosynthesis protein) |
| COG3600@1\|root | 5 | 0 | 0 | Protein of unknown function (DUF4065) |
| COG4680@1\|root | 5 | 0 | 0 | regulation of mRNA catabolic process; HigB_toxin, RelE-like toxic component of a toxin-antitoxin system |
| COG4683@1\|root | 5 | 0 | 0 | Phage derived protein Gp49-like (DUF891) |
| COG4938@1\|root | 5 | 0 | 0 | Protein of unknown function (DUF3696) |
| 28H70@1\|root | 4 | 0 | 0 | Multiubiquitin |
| 28IJG@1\|root | 4 | 0 | 0 | XamI restriction endonuclease |
| 29PHP@1\|root | 4 | 0 | 0 | Type-F conjugative transfer system pilin assembly protein |
| 29R2N@1\|root | 4 | 0 | 0 | - |
| 29TB7@1\|root | 4 | 0 | 0 | - |
| 2A9IN@1\|root | 4 | 0 | 0 | - |
| 2AHTU@1\|root | 4 | 0 | 0 | Antitoxin Phd_YefM, type II toxin-antitoxin system |
| 2BB2W@1\|root | 4 | 0 | 0 | Nuclease-related domain |
| 2C1FP@1\|root | 4 | 0 | 0 | -; -; - |
| 2CF90@1\|root | 4 | 0 | 0 | Nucleotidyltransferase substrate binding protein like |
| 2DMZH@1\|root | 4 | 0 | 0 | Protein of unknown function (DUF3800) |
| 2DNYB@1\|root | 4 | 0 | 0 | RloB-like protein; |
| 2DP6R@1\|root | 4 | 0 | 0 | -; - |
| 2E5P1@1\|root | 4 | 0 | 0 | Protein of unknown function (DUF2442); |
| 2EH2T@1\|root | 4 | 0 | 0 | -; - |
| arCOG06889@1\|root | 4 | 0 | 0 | MTH538 TIR-like domain (DUF1863); |
| COG1141@1\|root | 4 | 0 | 0 | 4Fe-4S single cluster domain of Ferredoxin I |
| COG3321@1\|root | 4 | 0 | 0 | polyketide synthase; polyketide synthase; PKS_KR; polyketide synthase |
| COG4861@1\|root | 4 | 0 | 0 | Protein conserved in bacteria; Transcriptional regulator, AbiEi antitoxin, Type IV TA system |
| 28HPH@1\|root | 3 | 0 | 0 | -; -; - |
| 28HYK@1\|root | 3 | 0 | 0 | Protein of unknown function (DUF3800); |
| 28I67@1\|root | 3 | 0 | 0 | -; - |
| 28I6N@1\|root | 3 | 0 | 0 | -; -; - |
| 28MI1@1\|root | 3 | 0 | 0 | Nucleotidyl transferase AbiEii toxin, Type IV TA system; |
| 2918Q@1\|root | 3 | 0 | 0 | T3SS negative regulator,GrlR |
| 294PF@1\|root | 3 | 0 | 0 | - |
| 296TN@1\|root | 3 | 0 | 0 | Sulfotransferase domain |
| 29RCQ@1\|root | 3 | 0 | 0 | Domain of unknown function (DUF4276); |
| 2A03C@1\|root | 3 | 0 | 0 | - |
| 2A1D2@1\|root | 3 | 0 | 0 | Protein of unknown function (DUF2442) |
| 2A1G6@1\|root | 3 | 0 | 0 | - |
| 2AEIE@1\|root | 3 | 0 | 0 | - |
| 2C1VC@1\|root | 3 | 0 | 0 | -; -; - |
| 2C72C@1\|root | 3 | 0 | 0 | Domain of unknown function (DUF4926); |
| 2DM5T@1\|root | 3 | 0 | 0 | Domain of unknown function (DUF4126); |
| 2DPG5@1\|root | 3 | 0 | 0 | Protein of unknown function (DUF2442); |
| 2DRQZ@1\|root | 3 | 0 | 0 | Restriction endonuclease; |
| 2E46S@1\|root | 3 | 0 | 0 | -; -; - |
| 2E4D4@1\|root | 3 | 0 | 0 | - |
| 2E8SU@1\|root | 3 | 0 | 0 | -; - |
| 2E9JM@1\|root | 3 | 0 | 0 | - |
| 2EANF@1\|root | 3 | 0 | 0 | - |
| 2EAQB@1\|root | 3 | 0 | 0 | -; - |
| 2EFH4@1\|root | 3 | 0 | 0 | -; - |

| | | | | |
|---|---|---|---|---|
| 2EGWS@1\|root | 3 | 0 | 0 | - |
| 2EPQB@1\|root | 3 | 0 | 0 | Putative addiction module component |
| 2EUCM@1\|root | 3 | 0 | 0 | - |
| 2EUJ6@1\|root | 3 | 0 | 0 | -; -; - |
| 2F194@1\|root | 3 | 0 | 0 | - |
| arCOG06613@1\|root | 3 | 0 | 0 | AIPR protein; AIPR protein; AIPR protein |
| arCOG06916@1\|root | 3 | 0 | 0 | Domain of unknown function (DUF4276); |
| arCOG10846@1\|root | 3 | 0 | 0 | nuclease activity; |
| COG0641@1\|root | 3 | 0 | 0 | Radical SAM superfamily; radical SAM |
| COG1431@1\|root | 3 | 0 | 0 | Peptidase, M16; PFAM Stem cell self-renewal protein Piwi; - |
| COG1630@1\|root | 3 | 0 | 0 | NurA; |
| COG2301@1\|root | 3 | 0 | 0 | Belongs to the HpcH HpaI aldolase family; Belongs to the HpcH HpaI aldolase family; Belongs to the HpcH HpaI aldolase family |
| COG2357@1\|root | 3 | 0 | 0 | Region found in RelA / SpoT proteins; guanosine tetraphosphate metabolic process; Region found in RelA / SpoT proteins |
| COG2886@1\|root | 3 | 0 | 0 | Uncharacterised protein family (UPF0175); PFAM Uncharacterised protein family UPF0175 |
| COG3239@1\|root | 3 | 0 | 0 | Alkane 1-monooxygenase; Fatty acid desaturase; Alkane 1-monooxygenase |
| COG3512@1\|root | 3 | 0 | 0 | CRISPR (clustered regularly interspaced short palindromic repeat), is an adaptive immune system that provides protection against mobile genetic elements (viruses, transposable elements and conjugative plasmids). |
| COG3513@1\|root | 3 | 0 | 0 | CRISPR (clustered regularly interspaced short palindromic repeat) is an adaptive immune system that provides protection against mobile genetic elements (viruses, transposable elements and conjugative plasmids). |
| COG3645@1\|root | 3 | 0 | 0 | DNA-damage-inducible protein d; Psort location Cytoplasmic, score 8.96; SOS response |
| COG3886@1\|root | 3 | 0 | 0 | PLD-like domain; PLD-like domain; PLD-like domain |
| COG5378@1\|root | 3 | 0 | 0 | PIN domain |
| 28H5D@1\|root | 2 | 0 | 0 | Restriction endonuclease EcoRV; Restriction endonuclease EcoRV |
| 28J7V@1\|root | 2 | 0 | 0 | NgoMIV restriction enzyme |
| 28JMM@1\|root | 2 | 0 | 0 | -; - |
| 28K3C@1\|root | 2 | 0 | 0 | - |
| 28KC2@1\|root | 2 | 0 | 0 | CRISPR-associated protein GSU0053 (Cas_GSU0053); CRISPR-associated protein GSU0053 (Cas_GSU0053) |
| 28M2G@1\|root | 2 | 0 | 0 | Protein of unknown function (DUF3800) |
| 28M70@1\|root | 2 | 0 | 0 | PFAM Restriction endonuclease, type II, Bpu10I |
| 28PTH@1\|root | 2 | 0 | 0 | -; -; - |
| 28WCA@1\|root | 2 | 0 | 0 | - |
| 28Z22@1\|root | 2 | 0 | 0 | - |
| 294EY@1\|root | 2 | 0 | 0 | - |
| 29ATZ@1\|root | 2 | 0 | 0 | - |
| 29X54@1\|root | 2 | 0 | 0 | Protein of unknown function (DUF2442); Protein of unknown function (DUF2442) |
| 2AG75@1\|root | 2 | 0 | 0 | -; - |
| 2B3FR@1\|root | 2 | 0 | 0 | - |
| 2BHNR@1\|root | 2 | 0 | 0 | Domain of unknown function (DUF4160) |
| 2BK4D@1\|root | 2 | 0 | 0 | -; - |
| 2BNF2@1\|root | 2 | 0 | 0 | -; - |
| 2BQUW@1\|root | 2 | 0 | 0 | - |
| 2BWMX@1\|root | 2 | 0 | 0 | -; - |
| 2C10S@1\|root | 2 | 0 | 0 | Restriction endonuclease NotI |
| 2C5KY@1\|root | 2 | 0 | 0 | - |
| 2C8CT@1\|root | 2 | 0 | 0 | - |
| 2C8HQ@1\|root | 2 | 0 | 0 | -; - |
| 2CCKR@1\|root | 2 | 0 | 0 | - |
| 2CHG4@1\|root | 2 | 0 | 0 | -; - |
| 2CM4P@1\|root | 2 | 0 | 0 | - |
| 2CNI4@1\|root | 2 | 0 | 0 | -; - |
| 2CT64@1\|root | 2 | 0 | 0 | HicA toxin of bacterial toxin-antitoxin, |

| | | | | |
|---|---|---|---|---|
| 2DBEM@1\|root | 2 | 0 | 0 | Domain of unknown function (DUF4433); Domain of unknown function (DUF4433) |
| 2DGKD@1\|root | 2 | 0 | 0 | - |
| 2DM15@1\|root | 2 | 0 | 0 | Methyltransferase FkbM domain |
| 2DMID@1\|root | 2 | 0 | 0 | - |
| 2DMWM@1\|root | 2 | 0 | 0 | -; - |
| 2DP1W@1\|root | 2 | 0 | 0 | - |
| 2DQR1@1\|root | 2 | 0 | 0 | - |
| 2DR05@1\|root | 2 | 0 | 0 | Peptidase inhibitor I78 family |
| 2DRAI@1\|root | 2 | 0 | 0 | - |
| 2DS95@1\|root | 2 | 0 | 0 | - |
| 2DSUP@1\|root | 2 | 0 | 0 | - |
| 2DW5W@1\|root | 2 | 0 | 0 | - |
| 2DYDA@1\|root | 2 | 0 | 0 | PIN domain |
| 2E3CI@1\|root | 2 | 0 | 0 | ParE toxin of type II toxin-antitoxin system, parDE; ParE toxin of type II toxin-antitoxin system, parDE |
| 2E4HV@1\|root | 2 | 0 | 0 | -; - |
| 2E9J1@1\|root | 2 | 0 | 0 | - |
| 2EAFZ@1\|root | 2 | 0 | 0 | Domain of unknown function (DUF4393) |
| 2EAZ1@1\|root | 2 | 0 | 0 | Evidence 3 Function proposed based on presence of conserved amino acid motif, structural feature or limited homology |
| 2ED35@1\|root | 2 | 0 | 0 | - |
| 2EFXV@1\|root | 2 | 0 | 0 | Nitrile hydratase; Nitrile hydratase; Nitrile hydratase; Nitrile hydratase |
| 2EH8B@1\|root | 2 | 0 | 0 | - |
| 2EHW9@1\|root | 2 | 0 | 0 | -; - |
| 2EPPT@1\|root | 2 | 0 | 0 | - |
| 2F4SE@1\|root | 2 | 0 | 0 | - |
| 2FFE4@1\|root | 2 | 0 | 0 | - |
| arCOG12551@1\|root | 2 | 0 | 0 | Putative PD-(D/E)XK family member, (DUF4420); Putative PD-(D/E)XK family member, (DUF4420) |
| COG1260@1\|root | 2 | 0 | 0 | Myo-inositol-1-phosphate synthase |
| COG1546@1\|root | 2 | 0 | 0 | TIGRFAM competence damage-inducible protein CinA; Belongs to the CinA family |
| COG1637@1\|root | 2 | 0 | 0 | Domain of unknown function (DUF4268); Domain of unknown function (DUF4268) |
| COG1746@1\|root | 2 | 0 | 0 | tRNA cytidylyltransferase activity; tRNA cytidylyltransferase activity |
| COG1879@1\|root | 2 | 0 | 0 | PFAM periplasmic binding protein LacI transcriptional regulator; PFAM periplasmic binding protein LacI transcriptional regulator |
| COG3013@1\|root | 2 | 0 | 0 | YfbU domain; YfbU domain |
| COG3972@1\|root | 2 | 0 | 0 | UvrD-like helicase C-terminal domain; DNA helicase |
| COG5619@1\|root | 2 | 0 | 0 | Uncharacterized conserved protein (DUF2290); Uncharacterized conserved protein (DUF2290) |
| KOG1435@1\|root | 2 | 0 | 0 | sterol delta7 reductase activity |
| 28H52@1\|root | 1 | 0 | 0 | Protein of unknown function (DUF1329) |
| 28HJT@1\|root | 1 | 0 | 0 | PFAM Restriction endonuclease BamHI |
| 28ICZ@1\|root | 1 | 0 | 0 | SIR2-like domain |
| 28IYQ@1\|root | 1 | 0 | 0 | Type II restriction endonuclease EcoO109I |
| 28J59@1\|root | 1 | 0 | 0 | - |
| 28J8Z@1\|root | 1 | 0 | 0 | - |
| 28J9C@1\|root | 1 | 0 | 0 | Restriction endonuclease BsobI |
| 28JFH@1\|root | 1 | 0 | 0 | TIGRFAM CRISPR-associated protein, Cse1 family |
| 28JYP@1\|root | 1 | 0 | 0 | RES domain |
| 28KC1@1\|root | 1 | 0 | 0 | F plasmid TraN appears to recognize OmpA in the recipient cell |
| 28KU8@1\|root | 1 | 0 | 0 | STELLO glycosyltransferases |
| 28KYP@1\|root | 1 | 0 | 0 | Domain of unknown function DUF1828 |
| 28M0F@1\|root | 1 | 0 | 0 | - |
| 28MBM@1\|root | 1 | 0 | 0 | - |
| 28MDY@1\|root | 1 | 0 | 0 | - |
| 28MTH@1\|root | 1 | 0 | 0 | Protein of unknown function (DUF4007) |
| 28NJ6@1\|root | 1 | 0 | 0 | Beta protein |
| 28NVF@1\|root | 1 | 0 | 0 | - |

| | | | | |
|---|---|---|---|---|
| 28NWR@1\|root | 1 | 0 | 0 | PD-(D/E)XK nuclease superfamily; PD-(D/E)XK nuclease superfamily |
| 28PA4@1\|root | 1 | 0 | 0 | - |
| 28Q19@1\|root | 1 | 0 | 0 | Protein of unknown function (DUF1524) |
| 28QM9@1\|root | 1 | 0 | 0 | Protein of unknown function (DUF3800) |
| 28WYZ@1\|root | 1 | 0 | 0 | - |
| 29WR2@1\|root | 1 | 0 | 0 | Protein of unknown function (DUF2442) |
| 29WR8@1\|root | 1 | 0 | 0 | Uncharacterized protein conserved in bacteria (DUF2236) |
| 29YK4@1\|root | 1 | 0 | 0 | Type II restriction enzyme SfiI |
| 2A0GS@1\|root | 1 | 0 | 0 | - |
| 2A3Z6@1\|root | 1 | 0 | 0 | - |
| 2A4D2@1\|root | 1 | 0 | 0 | Protein of unknown function (DUF2905) |
| 2AGFQ@1\|root | 1 | 0 | 0 | - |
| 2AHDV@1\|root | 1 | 0 | 0 | - |
| 2AHYN@1\|root | 1 | 0 | 0 | - |
| 2AJDX@1\|root | 1 | 0 | 0 | - |
| 2ANBB@1\|root | 1 | 0 | 0 | - |
| 2AP99@1\|root | 1 | 0 | 0 | - |
| 2AUQT@1\|root | 1 | 0 | 0 | - |
| 2AWGV@1\|root | 1 | 0 | 0 | - |
| 2AY5K@1\|root | 1 | 0 | 0 | - |
| 2B6ZF@1\|root | 1 | 0 | 0 | - |
| 2BFBB@1\|root | 1 | 0 | 0 | - |
| 2BJH7@1\|root | 1 | 0 | 0 | - |
| 2BK2H@1\|root | 1 | 0 | 0 | Restriction endonuclease BglII |
| 2BM5T@1\|root | 1 | 0 | 0 | - |
| 2BMG5@1\|root | 1 | 0 | 0 | - |
| 2BPIW@1\|root | 1 | 0 | 0 | - |
| 2BR06@1\|root | 1 | 0 | 0 | - |
| 2BV38@1\|root | 1 | 0 | 0 | SIR2-like domain |
| 2BVMY@1\|root | 1 | 0 | 0 | - |
| 2BWTI@1\|root | 1 | 0 | 0 | - |
| 2BX5R@1\|root | 1 | 0 | 0 | - |
| 2BXAG@1\|root | 1 | 0 | 0 | - |
| 2BXCR@1\|root | 1 | 0 | 0 | - |
| 2BZTS@1\|root | 1 | 0 | 0 | - |
| 2C10P@1\|root | 1 | 0 | 0 | - |
| 2C1S9@1\|root | 1 | 0 | 0 | - |
| 2C1W3@1\|root | 1 | 0 | 0 | - |
| 2C20N@1\|root | 1 | 0 | 0 | - |
| 2C2MK@1\|root | 1 | 0 | 0 | Restriction endonuclease NotI |
| 2C40F@1\|root | 1 | 0 | 0 | - |
| 2C5WF@1\|root | 1 | 0 | 0 | Restriction endonuclease |
| 2C62F@1\|root | 1 | 0 | 0 | - |
| 2C6PE@1\|root | 1 | 0 | 0 | - |
| 2C851@1\|root | 1 | 0 | 0 | - |
| 2C86X@1\|root | 1 | 0 | 0 | - |
| 2C9UE@1\|root | 1 | 0 | 0 | - |
| 2CB4H@1\|root | 1 | 0 | 0 | - |
| 2CE0P@1\|root | 1 | 0 | 0 | - |
| 2CEEW@1\|root | 1 | 0 | 0 | - |
| 2CEMP@1\|root | 1 | 0 | 0 | - |
| 2CH9B@1\|root | 1 | 0 | 0 | CRISPR-associated protein (Cas_Cas5) |
| 2CI7T@1\|root | 1 | 0 | 0 | - |
| 2CKI7@1\|root | 1 | 0 | 0 | - |
| 2CKQA@1\|root | 1 | 0 | 0 | - |
| 2CV9G@1\|root | 1 | 0 | 0 | - |
| 2CWAR@1\|root | 1 | 0 | 0 | - |
| 2D16E@1\|root | 1 | 0 | 0 | Motility quorum-sensing regulator, toxin of MqsA |
| 2DAFR@1\|root | 1 | 0 | 0 | ParD-like antitoxin of type II bacterial toxin-antitoxin system |
| 2DB8R@1\|root | 1 | 0 | 0 | - |
| 2DB9A@1\|root | 1 | 0 | 0 | Psort location Cytoplasmic, score |
| 2DBHK@1\|root | 1 | 0 | 0 | CRISPR-associated protein, GSU0054 family (Cas_GSU0054) |
| 2DBHY@1\|root | 1 | 0 | 0 | - |

| | | | | |
|---|---|---|---|---|
| 2DBJ6@1\|root | 1 | 0 | 0 | KilA-N |
| 2DBQ2@1\|root | 1 | 0 | 0 | - |
| 2DC1X@1\|root | 1 | 0 | 0 | CRISPR associated protein |
| 2DCA2@1\|root | 1 | 0 | 0 | - |
| 2DE59@1\|root | 1 | 0 | 0 | - |
| 2DJN0@1\|root | 1 | 0 | 0 | MTH538 TIR-like domain (DUF1863) |
| 2DM29@1\|root | 1 | 0 | 0 | Abortive infection C-terminus |
| 2DMGG@1\|root | 1 | 0 | 0 | - |
| 2DMPM@1\|root | 1 | 0 | 0 | -; - |
| 2DN1S@1\|root | 1 | 0 | 0 | Golgi phosphoprotein 3 (GPP34) |
| 2DNAB@1\|root | 1 | 0 | 0 | Domain of unknown function (DUF4433) |
| 2DNVU@1\|root | 1 | 0 | 0 | Protein of unknown function (DUF2442) |
| 2DP4H@1\|root | 1 | 0 | 0 | Protein of unknown function (DUF2442) |
| 2DPB2@1\|root | 1 | 0 | 0 | - |
| 2DPCE@1\|root | 1 | 0 | 0 | - |
| 2DPDF@1\|root | 1 | 0 | 0 | - |
| 2DQUE@1\|root | 1 | 0 | 0 | - |
| 2DRB5@1\|root | 1 | 0 | 0 | Putative addiction module component |
| 2DT95@1\|root | 1 | 0 | 0 | - |
| 2DTZU@1\|root | 1 | 0 | 0 | - |
| 2DU51@1\|root | 1 | 0 | 0 | - |
| 2DU5G@1\|root | 1 | 0 | 0 | Antitoxin Phd_YefM, type II toxin-antitoxin system |
| 2DW4I@1\|root | 1 | 0 | 0 | - |
| 2DYKZ@1\|root | 1 | 0 | 0 | LAGLIDADG endonuclease |
| 2DZQY@1\|root | 1 | 0 | 0 | - |
| 2DZT9@1\|root | 1 | 0 | 0 | - |
| 2DZZK@1\|root | 1 | 0 | 0 | - |
| 2E0UP@1\|root | 1 | 0 | 0 | - |
| 2E0WH@1\|root | 1 | 0 | 0 | restriction endonuclease |
| 2E10U@1\|root | 1 | 0 | 0 | - |
| 2E24U@1\|root | 1 | 0 | 0 | - |
| 2E25F@1\|root | 1 | 0 | 0 | - |
| 2E2T3@1\|root | 1 | 0 | 0 | - |
| 2E3P3@1\|root | 1 | 0 | 0 | - |
| 2E3ZN@1\|root | 1 | 0 | 0 | - |
| 2E45P@1\|root | 1 | 0 | 0 | - |
| 2E5ZM@1\|root | 1 | 0 | 0 | - |
| 2E6MH@1\|root | 1 | 0 | 0 | Domain of unknown function (DUF4160) |
| 2E7J7@1\|root | 1 | 0 | 0 | - |
| 2E8MC@1\|root | 1 | 0 | 0 | - |
| 2E953@1\|root | 1 | 0 | 0 | - |
| 2E9PK@1\|root | 1 | 0 | 0 | Domain of unknown function (DUF4258) |
| 2EA28@1\|root | 1 | 0 | 0 | Domain of unknown function (DUF4326) |
| 2EB1N@1\|root | 1 | 0 | 0 | - |
| 2EBHJ@1\|root | 1 | 0 | 0 | - |
| 2EE2G@1\|root | 1 | 0 | 0 | - |
| 2EG2X@1\|root | 1 | 0 | 0 | - |
| 2EGD8@1\|root | 1 | 0 | 0 | - |
| 2EGI4@1\|root | 1 | 0 | 0 | - |
| 2EIHU@1\|root | 1 | 0 | 0 | Protein of unknown function (DUF2442) |
| 2EIIA@1\|root | 1 | 0 | 0 | CRISPR associated protein Cas1 |
| 2EKA7@1\|root | 1 | 0 | 0 | Multiubiquitin |
| 2EKBW@1\|root | 1 | 0 | 0 | - |
| 2EMZW@1\|root | 1 | 0 | 0 | - |
| 2EN6U@1\|root | 1 | 0 | 0 | STAS-like domain of unknown function (DUF4325) |
| 2ENFS@1\|root | 1 | 0 | 0 | - |
| 2EPC0@1\|root | 1 | 0 | 0 | - |
| 2EQ8V@1\|root | 1 | 0 | 0 | - |
| 2ERZX@1\|root | 1 | 0 | 0 | - |
| 2ESEF@1\|root | 1 | 0 | 0 | - |
| 2ETCA@1\|root | 1 | 0 | 0 | TIR domain |
| 2EXE2@1\|root | 1 | 0 | 0 | - |
| 2F57W@1\|root | 1 | 0 | 0 | Protein of unknown function (DUF3800) |
| 2FBW8@1\|root | 1 | 0 | 0 | - |

| | | | | |
|---|---|---|---|---|
| 2FBYB@1\|root | 1 | 0 | 0 | - |
| 2FE6T@1\|root | 1 | 0 | 0 | -; - |
| 2FGA9@1\|root | 1 | 0 | 0 | - |
| 2FK18@1\|root | 1 | 0 | 0 | - |
| arCOG01426@1\|root | 1 | 0 | 0 | - |
| arCOG04629@1\|root | 1 | 0 | 0 | Uncharacterised protein family (UPF0175) |
| arCOG05255@1\|root | 1 | 0 | 0 | STAS-like domain of unknown function (DUF4325) |
| arCOG05276@1\|root | 1 | 0 | 0 | - |
| arCOG08210@1\|root | 1 | 0 | 0 | -; - |
| arCOG08856@1\|root | 1 | 0 | 0 | Psort location Cytoplasmic, score 8.96 |
| arCOG11468@1\|root | 1 | 0 | 0 | -; - |
| arCOG13036@1\|root | 1 | 0 | 0 | - |
| arCOG13189@1\|root | 1 | 0 | 0 | - |
| arCOG13199@1\|root | 1 | 0 | 0 | - |
| COG0143@1\|root | 1 | 0 | 0 | Is required not only for elongation of protein synthesis but also for the initiation of all mRNA translation through initiator tRNA(fMet) aminoacylation |
| COG1195@1\|root | 1 | 0 | 0 | DNA synthesis involved in DNA repair |
| COG1205@1\|root | 1 | 0 | 0 | DEAD DEAH box helicase |
| COG1372@1\|root | 1 | 0 | 0 | intein-mediated protein splicing |
| COG1567@1\|root | 1 | 0 | 0 | CRISPR-associated RAMP protein, Csm4 family |
| COG1574@1\|root | 1 | 0 | 0 | Amidohydrolase family |
| COG1788@1\|root | 1 | 0 | 0 | Coenzyme A transferase |
| COG1809@1\|root | 1 | 0 | 0 | (2R)-phospho-3-sulfolactate synthase (ComA); (2R)-phospho-3-sulfolactate synthase (ComA) |
| COG1910@1\|root | 1 | 0 | 0 | DNA binding domain, excisionase family |
| COG2057@1\|root | 1 | 0 | 0 | Acyl CoA acetate 3-ketoacid CoA transferase, beta subunit |
| COG2233@1\|root | 1 | 0 | 0 | xanthine |
| COG2515@1\|root | 1 | 0 | 0 | Catalyzes a cyclopropane ring-opening reaction, the irreversible conversion of 1-aminocyclopropane-1-carboxylate (ACC) to ammonia and alpha-ketobutyrate. Allows growth on ACC as a nitrogen source |
| COG2522@1\|root | 1 | 0 | 0 | sequence-specific DNA binding |
| COG2895@1\|root | 1 | 0 | 0 | Belongs to the TRAFAC class translation factor GTPase superfamily. Classic translation factor GTPase family. CysN NodQ subfamily |
| COG2927@1\|root | 1 | 0 | 0 | DNA polymerase III, chi subunit |
| COG2932@1\|root | 1 | 0 | 0 | sequence-specific DNA binding |
| COG3038@1\|root | 1 | 0 | 0 | cytochrome b561 |
| COG3203@1\|root | 1 | 0 | 0 | Forms passive diffusion pores that allow small molecular weight hydrophilic materials across the outer membrane; Protein of unknown function (DUF1302) |
| COG3392@1\|root | 1 | 0 | 0 | D12 class N6 adenine-specific DNA methyltransferase |
| COG3410@1\|root | 1 | 0 | 0 | Uncharacterized conserved protein (DUF2075) |
| COG3421@1\|root | 1 | 0 | 0 | Type III restriction enzyme, res subunit |
| COG3428@1\|root | 1 | 0 | 0 | Bacterial PH domain |
| COG3465@1\|root | 1 | 0 | 0 | - |
| COG3470@1\|root | 1 | 0 | 0 | protein probably involved in high-affinity Fe2 transport |
| COG3503@1\|root | 1 | 0 | 0 | Protein of unknown function (DUF1624) |
| COG3571@1\|root | 1 | 0 | 0 | Dienelactone hydrolase family |
| COG3740@1\|root | 1 | 0 | 0 | Caudovirus prohead serine protease |
| COG3793@1\|root | 1 | 0 | 0 | Tellurite resistance protein TerB |
| COG4341@1\|root | 1 | 0 | 0 | phosphohydrolase |
| COG4644@1\|root | 1 | 0 | 0 | Transposase |
| COG4770@1\|root | 1 | 0 | 0 | PFAM biotin lipoyl attachment domain-containing protein |
| COG4849@1\|root | 1 | 0 | 0 | Nucleotidyl transferase AbiEii toxin, Type IV TA system |
| COG4928@1\|root | 1 | 0 | 0 | KAP family P-loop domain |
| COG4933@1\|root | 1 | 0 | 0 | rRNA binding |
| COG5316@1\|root | 1 | 0 | 0 | Domain of unknown function (DUF4139) |
| COG5433@1\|root | 1 | 0 | 0 | transposase activity |
| COG5464@1\|root | 1 | 0 | 0 | double-stranded DNA endodeoxyribonuclease activity |

**Table S7:** Functional annotation of hallmark genes for chemolithoautotrophic ammonia-oxidation and carbon fixation.

| | | | arCOG | thaumarchaeal eggnog OG | notes | Nmar_loci to compare | Ca. sponge IRC_PAM_S | Ca. sponge IRC_PAM_S | Ca. spong GCA_01612 | Ca. sponge GCA_02129 | Cenarcheau sponge AXIM_halla | Ca. sponge IB2_shallow | Nitrosopum sponge LOPHE_tian | Nitrosopum sponge CYMC_moiti | Nitrosopum sponge SUB_tianQi | Cenarchea sponge STY3_bin_6 | Nitrosopum sponge APA_bin_56 | Nitrosopum sponge RHO2_bin | Nitrosoarch soil GCA_00022 | Nitrosopum marine GCA_00087 | Nitrosopum marine GCA_00001 |
|---|---|---|---|---|---|---|---|---|---|---|---|---|---|---|---|---|---|---|---|---|---|
| Energy metabolism & ammonia utilization | Ammonia-monooxygenase | | | | | | | | | | | | | | | | | | | | |
| | amoA | ammonia monooxygenase subunit A | arCOG08676 | 41SCF@651137\|Thaumarchaeota | key gene AOAs | Nmar_1500 | 1 | 1 | 1 | 0 | 1 | 1 | 1 | 1 | 1 | 1 | 1 | 1 | 1 | 1 | 1 |
| | amoX | ammonia monooxygenase subunit X | arCOG10586 | 41STH@651137\|Thaumarchaeota | | Nmar_1501 | 1 | 1 | 1 | 0 | 1 | 1 | 1 | 1 | 1 | 1 | 1 | 1 | 1 | 1 | 1 |
| | amoB | ammonia monooxygenase subunit B | arCOG08675 | 41SI5@651137\|Thaumarchaeota | | Nmar_1503 | 1 | 1 | 1 | 0 | 1 | 1 | 1 | 1 | 1 | 1 | 1 | 1 | 1 | 2 | 1 |
| | amoC | ammonia monooxygenase subunit C | arCOG08699 | 41S8N@651137\|Thaumarchaeota | | Nmar_1502 | 1 | 1 | 0 | 0 | 2 | 1 | 1 | 1 | 4 | 1 | 1 | 1 | 1 | 1 | 1 |
| | amoY | ammonia monooxygenase subunit Y | arCOG08672 | 41SU2@651137\|Thaumarchaeota | | Nmar_1506 | 1 | 0 | 1 | 0 | 1 | 1 | 1 | 1 | 1 | 1 | 1 | 1 | 1 | 1 | 1 |
| | amoZ | ammonia monooxygenase subunit Z | arCOG10587 | 41SVM@651137\|Thaumarchaeota | | Nmar_1507 | 1 | 1 | 1 | 1 | 1 | 1 | 1 | 1 | 1 | 1 | 1 | 2 | 1 | 2 | 1 |
| | periplasmic multicopper oxidases (oxidation of NH2OH) | | | | | | | | | | | | | | | | | | | | |
| | | PFAM blue (type 1) copper domain protein / Multicopper | arCOG02926 / arCOG03914 | 41SWV@651137\|Thaumarchaeota | (Cu-oxidase_2, Cu-oxidase_3, Copper-bind, Cupredoxin), multicopper | Nmar_1354 | 0 | 0 | 0 | 0 | 0 | 0 | 0 | 0 | 0 | 0 | 0 | 0 | 0 | 0 | 1 |
| | | PFAM blue (type 1) copper domain protein / Multicopper | arCOG02926 / arCOG03914 | 41SY3@651137\|Thaumarchaeota | (Cu-oxidase_2, Cu-oxidase_3, Cupredoxin_1, Copper_bind) | - | 0 | 0 | 0 | 0 | 0 | 0 | 0 | 0 | 0 | 0 | 0 | 0 | 1 | 1 | 0 |
| | Cu-HAO | multicopper oxidase | arCOG03914 | 41S77@651137\|Thaumarchaeota | putatively involved in the hydroxylamine oxidation - NcgA-like (nirK | Nmar_1663, Nmar_1131 | 1 | 1 | 1 | 1 | 1 | 1 | 1 | 1 | 1 | 1 | 1 | 1 | 1 | 1 | 2 |
| | mco | multicopper oxidase | arCOG03914 | 41T22@651137\|Thaumarchaeota | located next to genes of Complex IV; Copper_bind, Cu_oxidase_3 | | 1 | 1 | 1 | 1 | 0 | 1 | 0 | 0 | 1 | 1 | 1 | 1 | 1 | 1 | 0 |
| | mco | multicopper oxidase | seed_ortholog@436308.Nmar_1136\|- | arCOG03914@2157\|Archaea | 3dMCO (Cu_oxidase_2, Cu_oxidase_3, Cu_oxidase, Copper_bind, | Nmar_1136 | 0 | 0 | 0 | 0 | 0 | 0 | 0 | 0 | 0 | 0 | 0 | 0 | 0 | 1 | 1 |
| | Cu-nirK | putative copper-containig nitrite reductase | arCOG03914 | 41SZT@651137\|Thaumarchaeota | KEGG: K00368, Cu_oxidase_3 | Nmar_1667, Nmar_1259 | 1 | 1 | 2 | 2 | 0 | 1 | 1 | 1 | 1 | 1 | 1 | 1 | 1 | 1 | 2 |
| | Ammonia transporters | | | | | | | | | | | | | | | | | | | | |
| | amt2 | ammonia transporter | arCOG04397 | 41SY0@651137\|Thaumarchaeota | ammonia transporter (high affinity) | Nmar_1698 | 1 | 1 | 1 | 1 | 2 | 1 | 1 | 1 | 1 | 1 | 0 | 1 | 1 | 1 | 1 |
| | amt1 | ammonia transporter | arCOG04397 | 41SZ0@651137\|Thaumarchaeota | ammonia transporter (low affinity) | Nmar_0588 | 0 | 0 | 0 | 0 | 0 | 0 | 0 | 0 | 1 | 0 | 0 | 0 | 1 | 1 | 1 |
| | Urease and alternative ammonia sources | | | | | | | | | | | | | | | | | | | | |
| | ureD | urease accessory protein ureD | arCOG04529 | 41SBH@651137\|Thaumarchaeota | urea | | 0 | 1 | 1 | 1 | 1 | 0 | 0 | 1 | 0 | 0 | 0 | 0 | 0 | 1 | 0 |
| | ureG | urease accessory protein ureG | arCOG01231 | 41SAZ@651137\|Thaumarchaeota | | | 1 | 1 | 1 | 1 | 1 | 1 | 0 | 1 | 0 | 0 | 1 | 1 | 0 | 1 | 0 |
| | ureF | urease accessory protein ureF | arCOG04530 | 41SG4@651137\|Thaumarchaeota | | | 0 | 1 | 1 | 1 | 1 | 0 | 0 | 1 | 0 | 0 | 0 | 1 | 0 | 1 | 0 |
| | ureE | urease accessory protein ureE | arCOG06188 | 41STJ@651137\|Thaumarchaeota | | | 0 | 0 | 0 | 0 | 1 | 0 | 0 | 1 | 0 | 0 | 1 | 0 | 0 | 1 | 0 |
| | ureC | urea amidohydrolase (urease), subunit alpha | arCOG00698 | 41S7M@651137\|Thaumarchaeota | | | 1 | 1 | 1 | 1 | 1 | 1 | 0 | 1 | 0 | 0 | 1 | 1 | 0 | 1 | 0 |
| | ureB | urea amidohydrolase (urease), subunit beta | arCOG04527 | 41T4P@651137\|Thaumarchaeota | | | 0 | 0 | 1 | 1 | 1 | 0 | 0 | 0 | 0 | 0 | 1 | 0 | 0 | 1 | 0 |
| | ureA | urea amidohydrolase (urease), subunit gamma | arCOG04528 | 41SPI@651137\|Thaumarchaeota | | | 0 | 0 | 0 | 0 | 1 | 0 | 0 | 1 | 0 | 0 | 1 | 1 | 0 | 1 | 0 |
| | nitA | Nitrilase cyanide hydratase and apolipoprotein N-acyltransferase | arCOG00062 | 41SF8@651137\|Thaumarchaeota | Nitrilase/Cyanide hydratase | Nmar_1633, Nmar_0650 | 1 | 1 | 1 | 1 | 1 | 1 | 2 | 1 | 2 | 0 | 1 | 2 | 1 | 2 | 2 |
| | cre1/arfB | Creatininase amidohydralase | arCOG04536 | 41SNC@651137\|Thaumarchaeota | Creatiniase / Creatinine amidohyrolase | Nmar_0222 | 1 | 1 | 1 | 1 | 1 | 1 | 1 | 1 | 1 | 1 | 1 | 1 | 1 | 1 | 1 |
| | | Cytosine deaminase | arCOG00692 | 41S61@651137\|Thaumarchaeota | Creatinin degradation Step 1 | Nmar_1613 | 1 | 1 | 0 | 1 | 1 | 1 | 1 | 1 | 1 | 1 | 1 | 1 | 1 | 1 | 1 |
| | | N-methylhydantoin amidohyrolase | arCOG01511 | 41SZ7@651137\|Thaumarchaeota | Creatinin degradation step 2 | | 0 | 0 | 1 | 1 | 0 | 0 | 0 | 0 | 1 | 1 | 1 | 1 | 1 | 0 | 0 |
| | | Creatinase amidohydralase | arCOG01000 | 41T14@651137\|Thaumarchaeota | creatinase (creatine to urea) | | 1 | 0 | 1 | 1 | 1 | 1 | 1 | 1 | 1 | 1 | 0 | 1 | 1 | 1 | 1 |
| | Electron-transfer flavoproteins | | | | | | | | | | | | | | | | | | | | |
| | (fixCX) | putative electron-transfer flavoprotein /FAD-dependent / fixCX-like | arCOG00570 | 41S82@651137\|Thaumarchaeota | geranylgeranyl reductase | Nmar_0631 | 0 | 0 | 1 | 1 | 1 | 1 | 1 | 1 | 1 | 1 | 1 | 1 | 1 | 1 | 1 |
| | (fixCX) | putative electron-transfer flavoprotein /FAD-dependent / fixCX-like | arCOG00570 | 41SBB@651137\|Thaumarchaeota | FAD_binding_2 | Nmar_1514 | 1 | 1 | 1 | 1 | 1 | 1 | 1 | 1 | 1 | 1 | 1 | 1 | 1 | 1 | 1 |
| | (fixCX) | putative electron-transfer flavoprotein /FAD-dependent / fixCX-like | arCOG00570 | 41SCG@651137\|Thaumarchaeota | Trp_halogenase | Nmar_0359 | 1 | 1 | 1 | 1 | 1 | 1 | 1 | 2 | 1 | 1 | 1 | 1 | 1 | 1 | 1 |
| | (fixCX) | putative electron-transfer flavoprotein /FAD-dependent / fixCX-like | arCOG00570 | 41SB5@651137\|Thaumarchaeota | FAD_binding_3 | Nmar_1220 | 1 | 1 | 1 | 1 | 1 | 1 | 1 | 1 | 1 | 1 | 2 | 1 | 1 | 1 | 1 |
| | etfA (fixB) | electron transfer flavoprotein, alpha subunit | arCOG00448 | 41SZJ@651137\|Thaumarchaeota | | | 0 | 0 | 0 | 0 | 0 | 0 | 0 | 0 | 0 | 0 | 0 | 0 | 0 | 0 | 0 |
| | eftB (fixA) | electron transfer flavoprotein, beta subunit | arCOG00446 | 41SX2@651137\|Thaumarchaeota | | | 0 | 0 | 0 | 0 | 0 | 0 | 0 | 0 | 0 | 0 | 0 | 0 | 0 | 0 | 0 |
| | Rieske proteins and ferredoxins | | | | | | | | | | | | | | | | | | | | |
| | sdx | rieske [2Fe-2S] iron-sulphur domain protein | arCOG02854 | 41STC@651137\|Thaumarchaeota | | Nmar_0965 | 1 | 1 | 1 | 1 | 1 | 0 | 1 | 1 | 1 | 1 | 1 | 1 | 1 | 1 | 1 |
| | | 4Fe-4S ferredoxin iron-sulfur binding domain protein | arCOG00291 | 41SNB@651137\|Thaumarchaeota | | Nmar_1537 | 2 | 2 | 1 | 1 | 1 | 1 | 1 | 1 | 1 | 1 | 1 | 1 | 1 | 1 | 1 |
| | | 4Fe-4S ferredoxin iron-sulfur binding domain protein | arCOG00291 | 41T34@651137\|Thaumarchaeota | | - | 0 | 0 | 1 | 1 | 1 | 1 | 0 | 1 | 1 | 1 | 1 | 1 | 1 | 0 | 0 |
| | | 4Fe-4S ferredoxin iron-sulfur binding domain protein | arCOG00291 | arCOG00291@2157\|Archaea | | Nmar_1765 | 0 | 0 | 0 | 0 | 0 | 0 | 1 | 0 | 0 | 0 | 0 | 0 | 0 | 1 | 1 |
| | | ferredoxin | arCOG04548 | 41SH5@651137\|Thaumarchaeota | | Nmar_0443 | 1 | 1 | 1 | 1 | 1 | 1 | 1 | 1 | 1 | 1 | 1 | 1 | 1 | 1 | 1 |
| | | COG2146 Ferredoxin | arCOG02852 | arCOG02852 | COG2146 Ferredoxin subunits of nitrite reductase and ring-hydroxy | Nmar_0162 | 1 | 0 | 1 | 0 | 0 | 1 | 1 | 0 | 0 | 0 | 0 | 0 | 0 | 0 | 1 |
| | ferB | PFAM 4Fe-4S dicluster domain | arCOG04548 | 41SI0@651137\|Thaumarchaeota | | Nmar_0238, Nmar_0239 | 1 | 1 | 1 | 1 | 2 | 2 | 2 | 2 | 3 | 1 | 2 | 2 | 2 | 2 | 2 |
| | | PFAM Rieske 2Fe-2S domain | arCOG01720 | 41SCE@651137\|Thaumarchaeota | | Nmar_1544 | 1 | 1 | 1 | 1 | 1 | 1 | 1 | 1 | 1 | 1 | 1 | 1 | 1 | 1 | 1 |
| | Electron carriers, plastocyanines | | | | | | | | | | | | | | | | | | | | |
| | | blue (type 1) copper domain protein | arCOG02926 | 41SPG@651137\|Thaumarchaeota | PFAM Rieske 2Fe-2S domain | Nmar_1542, Nmar_1142 | 1 | 2 | 2 | 1 | 1 | 1 | 1 | 1 | 1 | 0 | 1 | 0 | 1 | 2 | 3 |
| | | blue (type 1) copper domain protein | arCOG02926 | 41T8E@651137\|Thaumarchaeota | PFAM Copper binding proteins, plastocyanin azurin family | Nmar_0918 | 0 | 0 | 0 | 0 | 0 | 0 | 1 | 1 | 0 | 0 | 0 | 0 | 0 | 1 | 1 |
| | | blue (type 1) copper domain protein | arCOG02926 | 41TAN@651137\|Thaumarchaeota | PFAM Copper binding proteins, plastocyanin azurin family | Nmar_1443 | 1 | 1 | 0 | 1 | 0 | 0 | 1 | 1 | 1 | 0 | 1 | 1 | 1 | 1 | 1 |
| | | PFAM blue (type 1) copper domain protein / Blue copper protein | arCOG01594 | 41SBF@651137\|Thaumarchaeota | MGS-like domain | Nmar_0185 | 1 | 1 | 1 | 1 | 2 | 1 | 1 | 1 | 1 | 1 | 2 | 1 | 1 | 1 | 1 |
| | | PFAM blue (type 1) copper domain protein / - | arCOG02926 / arCOG06255 | arCOG06255,arCOG02926,arCOG06255 | | Nmar_1102 | 1 | 1 | 1 | 1 | 0 | 1 | 0 | 1 | 0 | 0 | 1 | 0 | 1 | 1 | 1 |
| | | PFAM blue (type 1) copper domain protein / Fibronectin type III domain protein | arCOG02926 / arCOG05978 | arCOG05978,arCOG02926,arCOG05978 | | Nmar_1040 | 0 | 0 | 0 | 0 | 0 | 0 | 0 | 0 | 0 | 0 | 0 | 0 | 1 | 1 | 0 |
| | | PFAM blue (type 1) copper domain protein / copper ion binding | arCOG02926 / arCOG08795 | arCOG02926@2157\|Archaea / arCOG08795@2157\|Archaea | | - | 0 | 0 | 0 | 0 | 0 | 0 | 1 | 0 | 1 | 0 | 1 | 0 | 2 | 1 | 0 |
| | | PFAM blue (type 1) copper domain protein / sister chromatid segregation | arCOG02926 / arCOG10350 | arCOG02926@2157\|Archaea / arCOG10350@2157\|Archaea | | - | 0 | 1 | 0 | 0 | 0 | 0 | 1 | 0 | 1 | 0 | 1 | 0 | 1 | 0 | 1 |
| | | PFAM blue (type 1) copper domain protein / Fibronectin type III domain protein / | arCOG02926 / arCOG05978 / arCOG07813 | arCOG02926@2157\|Archaea / arCOG05978@2157\|Archaea / | | Nmar_1073 | 0 | 0 | 0 | 0 | 0 | 0 | 0 | 0 | 0 | 0 | 0 | 0 | 0 | 0 | 1 |
| | | blue (type 1) copper domain protein | arCOG02926 | arCOG02926@2157\|Archaea | cupredoxin | Nmar_1650, Nmar_1665, | 2 | 2 | 3 | 1 | 3 | 2 | 3 | 3 | 5 | 5 | 1 | 5 | 3 | 5 | 11 |
| | QRED | plastocyanin | arCOG02926 / arCOG02929 | 41SHN@651137\|Thaumarchaeota | QRED - quinone reductase | Nmar_1226 | 1 | 2 | 1 | 1 | 1 | 2 | 1 | 1 | 1 | 1 | 1 | 1 | 1 | 1 | 1 |
| | | plastocyanin | arCOG02929 | 41T61@651137\|Thaumarchaeota | | - | 0 | 0 | 0 | 0 | 0 | 0 | 0 | 0 | 0 | 0 | 0 | 0 | 1 | 0 | 0 |
| | | putative cupredoxin | arCOG11327 | 41T5V@651137\|Thaumarchaeota | | Nmar_1365 | 0 | 0 | 0 | 0 | 0 | 0 | 0 | 0 | 0 | 0 | 1 | 0 | 2 | 1 | 1 |
| | | Periplasmic protein of plastocyanin/azurin family protein | arCOG08798 | 41SZ3@651137\|Thaumarchaeota | | Nmar_0730 | 0 | 0 | 0 | 0 | 0 | 1 | 1 | 1 | 0 | 1 | 2 | 1 | 0 | 1 | 1 |
| | | Periplasmic protein of plastocyanin/azurin family protein | arCOG08764 | arCOG08764@2157\|Archaea | | Nmar_1753 | 1 | 1 | 2 | 1 | 1 | 1 | 1 | 0 | 0 | 1 | 0 | 0 | 1 | 1 | 1 |
| | Nitrogen regulatory protein PII | | | | | | | | | | | | | | | | | | | | |
| | | Belongs to the P(II) protein family | arCOG02305 | 41TB5@651137\|Thaumarchaeota | Belongs to the P(II) protein family | Nmar_1523 | 1 | 0 | 1 | 1 | 0 | 0 | 1 | 1 | 1 | 0 | 1 | 1 | 1 | 1 | 1 |
| | | Belongs to the P(II) protein family | arCOG02305 | arCOG02305@2157\|Archaea | Belongs to the P(II) protein family | Nmar_0587, Nmar_1317 | 0 | 0 | 0 | 0 | 1 | 0 | 2 | 2 | 2 | 0 | 0 | 0 | 4 | 3 | 3 |
| | | Nitrogen regulatory protein P-II | arCOG11286 | arCOG11286@2157\|Archaea | | Nmar_0486 | 0 | 0 | 0 | 0 | 0 | 1 | 1 | 1 | 1 | 0 | 0 | 0 | 0 | 1 | 1 |
| | alternative NADH dehydrogenases, type-II NADH dehydrogenase, NADH-dependent FMN reductases, FAD dependent oxidoreductases | | | | | | | | | | | | | | | | | | | | |
| | pndr | putative FAD-dependent pyridine nucleotide-disulphide oxidoreductase | arCOG01064 | 41SZR@651137\|Thaumarchaeota | PFAM FAD-dependent pyridine nucleotide-disulphide oxidoreductase | Nmar_1785 | 1 | 1 | 1 | 1 | 0 | 0 | 1 | 0 | 1 | 0 | 1 | 0 | 1 | 1 | 1 |
| | | COG1252 NADH dehydrogenase, FAD-containing subunit | arCOG01067 | arCOG01067@2157\|Archaea | NADH dehydrogenase | Nmar_0008 | 2 | 1 | 1 | 2 | 1 | 1 | 2 | 2 | 1 | 0 | 1 | 2 | 1 | 2 | 1 |
| | | NAD(P)H-dependent FMN reductase | arCOG04624 | 41T2F@651137\|Thaumarchaeota | NADH dehydrogenase, NAD(P)H-FMN reductase | Nmar_1737 | 1 | 1 | 1 | 1 | 1 | 1 | 1 | 1 | 3 | 1 | 1 | 1 | 1 | 1 | 1 |
| | | NAD(P)H-dependent FMN reductase | arCOG04624 | arCOG04624@2157\|Archaea | | Nmar_0654 | 0 | 0 | 0 | 0 | 0 | 0 | 1 | 0 | 0 | 0 | 0 | 0 | 0 | 1 | 1 |
| | ssuE1 | NAD(P)H-dependent FMN reductase | arCOG04624 | 41T7Y@651137\|Thaumarchaeota | | | 0 | 0 | 0 | 0 | 0 | 0 | 0 | 0 | 0 | 0 | 0 | 0 | 1 | 0 | 0 |
| | fnr | COG1018 Flavodoxin reductases (ferredoxin-NADPH reductases) family 1 | arCOG02200 | 41S72@651137\|Thaumarchaeota | oxidoreductase FAD NAD(P)-binding | Nmar_0253, Nmar_0267 | 1 | 1 | 1 | 1 | 2 | 2 | 2 | 2 | 2 | 1 | 2 | 2 | 2 | 2 | 2 |
| | glcD | COG0277 FAD FMN-containing dehydrogenases | arCOG00337 | 41T0E@651137\|Thaumarchaeota | | Nmar_0734 | 2 | 1 | 1 | 1 | 0 | 0 | 1 | 0 | 1 | 0 | 2 | 1 | 0 | 1 | 1 |
| | | FAD dependent oxidoreductase | arCOG00754 | 41SXA@651137\|Thaumarchaeota | | Nmar_0409 | 1 | 1 | 1 | 1 | 1 | 1 | 1 | 1 | 1 | 1 | 1 | 1 | 1 | 1 | 1 |
| | Glutamate and Glutamine | | | | | | | | | | | | | | | | | | | | |
| | gdhA1 | Glutamate dehydrogenase | arCOG01352 | 41S7T@651137\|Thaumarchaeota | | Nmar_1312 | 1 | 1 | 1 | 1 | 1 | 1 | 1 | 1 | 1 | 1 | 1 | 1 | 1 | 1 | 1 |
| | glnA | glutamine synthetase | arCOG01909 | 41SD6@651137\|Thaumarchaeota | | Nmar_1790, Nmar_1771 | 2 | 1 | 2 | 1 | 2 | 2 | 2 | 2 | 2 | 2 | 2 | 2 | 2 | 2 | 2 |
| 3-HP/4-HB cycle | 3-hydroxypropionate-4-hydroxybutyrate pathway | | | | | | | | | | | | | | | | | | | | |
| | accB/pccB | Acetyl-CoA carboxylase, carboxyltransferase component (subunits alpha and beta) | arCOG02705 | 41SEH@651137\|Thaumarchaeota | Acetyl-CoA carboxylase, carboxyltransferase component (subunits | Nmar_0272 | 1 | 1 | 1 | 1 | 1 | 1 | 1 | 1 | 1 | 1 | 1 | 1 | 1 | 1 | 1 |
| | accC/pccC | acetyl-CoA/propionyl-CoA carboxylase, biotin carboxylation subunit | arCOG01590 | 41SXC@651137\|Thaumarchaeota | Carbamoyl-phosphate synthetase large chain domain protein | Nmar_0273 | 1 | 1 | 1 | 1 | 1 | 1 | 1 | 1 | 1 | 1 | 1 | 1 | 1 | 1 | 1 |
| | accA/pccA | biotin carboxyl carrier protein of put. acetyl-CoA/ propionyl-CoA carboxylase | arCOG02095 | 41SKA@651137\|Thaumarchaeota | Biotin-requiring enzyme | Nmar_0274 | 1 | 1 | 1 | 1 | 1 | 1 | 1 | 1 | 1 | 1 | 1 | 1 | 1 | 1 | 1 |
| | mcr | malonyl-CoA reductase | arCOG00494 | 41SFD@651137\|Thaumarchaeota | | Nmar_1586 | 1 | 1 | 1 | 1 | 1 | 1 | 1 | 1 | 1 | 1 | 1 | 1 | 1 | 1 | 1 |
| | | 3-hydroxypropionate dehydrogenase (NADP+) | | | | | | | | | | | | | | | | | | | |
| | msr | malonic semialdehyde reductase | arCOG00984 | 41SFC@651137\|Thaumarchaeota | | Nmar_0161, Nmar_1110 | 1 | 1 | 1 | 1 | 1 | 1 | 1 | 1 | 1 | 1 | 1 | 1 | 1 | 1 | 2 |
| | hpcs | 3-hydroxypropionyl-CoA synthetase (ADP-forming) | arCOG01338 | 41S7I@651137\|Thaumarchaeota | Acyl-CoA synthetase (NDP forming) / Acyl-CoA synthetase (NDP forming) | Nmar_1309 | 1 | 1 | 1 | 1 | 1 | 1 | 1 | 1 | 1 | 1 | 1 | 1 | 1 | 1 | 1 |
| | mce | methylmalonyl-CoA epimerase | arCOG02706 | 41SQS@651137\|Thaumarchaeota | Methylmalonyl-CoA epimerase | Nmar_0953 | 1 | 1 | 0 | 1 | 1 | 1 | 1 | 1 | 1 | 1 | 0 | 0 | 1 | 1 | 1 |
| | mut | methylmalonyl-CoA mutase, N-terminal domain | arCOG04232 | 41SBW@651137\|Thaumarchaeota | Methylmalonyl-CoA mutase | Nmar_0954 | 1 | 1 | 1 | 1 | 1 | 1 | 1 | 1 | 1 | 1 | 1 | 1 | 1 | 1 | 1 |
| | mcm | methylmalonyl-CoA mutase, C-terminal domain | arCOG01710 | 41SJN@651137\|Thaumarchaeota | | Nmar_0958 | 1 | 1 | 1 | 1 | 1 | 1 | 1 | 1 | 1 | 1 | 0 | 1 | 1 | 1 | 1 |
| | | succinyl-CoA reductase | | | | | | | | | | | | | | | | | | | |
| | | succinate semialdehyde reductase | | | | | | | | | | | | | | | | | | | |
| | | 4-hydroxybutyrate---CoA ligase (AMP-forming) | | | | | | | | | | | | | | | | | | | |
| | hbcd | 4-hydroxybutyryl-CoA dehydratase / vinylacetyl-CoA-Delta-isomerase | arCOG02143 | 41SA0@651137\|Thaumarchaeota | PFAM 4-hydroxyphenylacetate 3-hydroxylase | Nmar_0207 | 1 | 1 | 1 | 1 | 1 | 1 | 1 | 1 | 1 | 1 | 1 | 1 | 1 | 1 | 1 |
| | | Acyl-CoA synthetase (NDP forming) | arCOG01340 | 41S6N@651137\|Thaumarchaeota | | Nmar_0206 | 0 | 1 | 1 | 1 | 1 | 1 | 1 | 1 | 1 | 1 | 1 | 1 | 1 | 1 | 1 |
| | cch | enoyl-CoA hydratase | arCOG00239 | 41SH7@651137\|Thaumarchaeota | 3-hydroxybutyryl-CoA dehydratase | Nmar_1308 | 1 | 1 | 1 | 1 | 1 | 1 | 1 | 1 | 1 | 1 | 2 | 1 | 1 | 1 | 1 |
| | hbd2 | 3-hydroxyacyl-CoA dehydrogenase | arCOG00249 | 41S92@651137\|Thaumarchaeota | | Nmar_1028 | 1 | 1 | 1 | 1 | 1 | 1 | 1 | 1 | 1 | 1 | 1 | 1 | 1 | 1 | 1 |
| | atoB | acetyl-CoA C-acetyltransferase | arCOG01278 | 41SB7@651137\|Thaumarchaeota | | Nmar_0841 | 1 | 1 | 1 | 1 | 1 | 1 | 1 | 1 | 1 | 1 | 1 | 1 | 1 | 1 | 1 |
| | atoB | acetyl-CoA C-acetyltransferase | arCOG01278 | 41SCI@651137\|Thaumarchaeota | | Nmar_1631 | 1 | 1 | 1 | 1 | 1 | 1 | 1 | 1 | 1 | 1 | 1 | 1 | 1 | 1 | 1 |
| | Candidate genes for 3-hydroxypropionate-4-hydroxybutyrate carbon fixation pathway | | | | | | | | | | | | | | | | | | | | |
| | | 3-hydroxypropionyl-coenzyme A synthetase | arCOG01529 | 41S8Z@651137\|Thaumarchaeota | | | 1 | 1 | 1 | 1 | 0 | 0 | 0 | 1 | 0 | 0 | 0 | 1 | 0 | 0 | 0 |
| | | 3-hydroxypropionyl-coenzyme A synthetase | arCOG01529 | 41SZA@651137\|Thaumarchaeota | | Nmar_0866 | 0 | 0 | 0 | 0 | 0 | 0 | 1 | 1 | 1 | 0 | 0 | 0 | 1 | 1 | 1 |
| | asd | aspartate-semialdehyde dehydrogenase | arCOG00494 | 41SFD@651137\|Thaumarchaeota | succinyl-CoA reductase / succinate semialdehyde reductase? | Nmar_1586 | 1 | 1 | 1 | 1 | 1 | 1 | 1 | 1 | 1 | 1 | 1 | 1 | 1 | 1 | 1 |
| | acs | Acyl-coenzyme A synthetase/AMP-(fatty) acid ligase | arCOG04201 | arCOG04201@2157\|Archaea | | Nmar_0701 | 0 | 0 | 0 | 0 | 0 | 0 | 1 | 0 | 1 | 0 | 1 | 0 | 1 | 1 | 1 |
| | gabD (SSADH) | succinate-semialdehyde dehydrogenase | arCOG01252 | 41T0F@651137\|Thaumarchaeota | succinate semialdehyde reductase? | Nmar_1608 | 1 | 1 | 1 | 1 | 0 | 0 | 1 | 1 | 0 | 0 | 1 | 1 | 1 | 1 | 1 |
| | Alcohol dehydrogenases, EC 1.1.1.- | | | | | | | | | | | | | | | | | | | | |
| | | NAD-dependent alcohol dehydrogenase | arCOG01459 | 41S98@651137\|Thaumarchaeota | Glucose dehydrogenase C-terminus | Nmar_1079 | 1 | 1 | 1 | 1 | 1 | 1 | 1 | 1 | 1 | 1 | 1 | 1 | 2 | 1 | 1 |
| | fdh | NAD-dependent alcohol dehydrogenase | arCOG01459 | arCOG01459@2157\|Archaea | Catalyzes the NAD(P)( )-dependent oxidation of D-glucose to D- | Nmar_0369 | 1 | 0 | 1 | 0 | 0 | 0 | 0 | 0 | 1 | 0 | 1 | 1 | 0 | 1 | 1 |
| | | COG0604 NADPH quinone reductase and related Zn-dependent oxidoreductases | arCOG01458 | 41SX7@651137\|Thaumarchaeota | | Nmar_1565 | 0 | 0 | 0 | 0 | 1 | 1 | 1 | 1 | 1 | 0 | 1 | 1 | 1 | 1 | 1 |
| | | COG0604 NADPH quinone reductase and related Zn-dependent oxidoreductases | arCOG01458 | 41SZ9@651137\|Thaumarchaeota | | Nmar_1622 | 1 | 1 | 1 | 1 | 1 | 1 | 1 | 1 | 1 | 1 | 1 | 1 | 1 | 1 | 1 |
| | adh4 | alcohol dehydrogenase | arCOG01455 | 41SA9@651137\|Thaumarchaeota | | Nmar_0523 | 1 | 1 | 1 | 1 | 1 | 1 | 1 | 1 | 1 | 1 | 1 | 1 | 1 | 1 | 1 |
| | | alcohol dehydrogenase | arCOG01455 | 41SK3@651137\|Thaumarchaeota | | Nmar_1193 | 0 | 0 | 1 | 1 | 1 | 1 | 1 | 1 | 1 | 1 | 1 | 1 | 1 | 1 | 1 |
| | | alcohol dehydrogenase | arCOG00984 | 41SFC@651137\|Thaumarchaeota | | Nmar_0161, Nmar_1110 | 1 | 1 | 1 | 1 | 1 | 1 | 1 | 1 | 1 | 1 | 1 | 1 | 1 | 1 | 2 |
| | Tricarboxylic acid cycle (oxidative) | | | | | | | | | | | | | | | | | | | | |
| | gltA | citrate synthase | arCOG04237 | 41SAN@651137\|Thaumarchaeota | | Nmar_1099 | 1 | 1 | 1 | 1 | 1 | 1 | 1 | 1 | 1 | 1 | 1 | 1 | 1 | 1 | 1 |
| | acnA | aconitate hydratase A | arCOG01697 | 41SE6@651137\|Thaumarchaeota | | Nmar_1482 | 1 | 2 | 2 | 2 | 1 | 1 | 1 | 1 | 1 | 1 | 1 | 1 | 1 | 1 | 1 |
| | icd | isocitrate/isopropylmalate dehydrogenase | arCOG01163 | 41S5U@651137\|Thaumarchaeota | | Nmar_1379 | 1 | 1 | 2 | 0 | 1 | 1 | 1 | 1 | 1 | 1 | 1 | 1 | 1 | 1 | 1 |
| | icd | Isocitrate/isopropylmalate dehydrogenase | arCOG01163 | 41SAA@651137\|Thaumarchaeota | | Nmar_1069 | 1 | 1 | 1 | 1 | 1 | 1 | 1 | 1 | 1 | 1 | 0 | 1 | 1 | 1 | 1 |
| | korAC | 2:oxoacid-ferredoxin oxidoreductase, fused alpha and gamma subunit | arCOG01606 | 41S60@651137\|Thaumarchaeota | | Nmar_0413 | 1 | 1 | 1 | 1 | 1 | 1 | 1 | 1 | 1 | 1 | 1 | 1 | 1 | 1 | 1 |
| | korB | 2:oxoacid-ferredoxin oxidoreductase, beta subunit | arCOG01599 | 41SAB@651137\|Thaumarchaeota | | Nmar_0414 | 1 | 1 | 1 | 1 | 1 | 1 | 1 | 1 | 1 | 1 | 1 | 1 | 1 | 1 | 1 |
| | sucD | succinyl-CoA ligase, subunit alpha, ADP-forming | arCOG01339 | 41SEU@651137\|Thaumarchaeota | Succinyl-CoA synthetase functions in the citric acid cycle (TCA), | Nmar_1124 | 1 | 1 | 1 | 1 | 1 | 1 | 1 | 1 | 1 | 1 | 0 | 1 | 1 | 1 | 1 |
| | sucC | succinyl-CoA ligase, subunit beta, ADP-forming | arCOG01337 | 41S5M@651137\|Thaumarchaeota | | Nmar_1123 | 1 | 1 | 1 | 1 | 1 | 1 | 1 | 1 | 1 | 1 | 0 | 1 | 1 | 1 | 1 |
| | sdhA | succinate dehydrogenase (ubiquinone) flavoprotein subunit | arCOG00571 | 41SAM@651137\|Thaumarchaeota | | Nmar_0258 | 1 | 1 | 1 | 1 | 1 | 1 | 1 | 1 | 1 | 1 | 1 | 1 | 1 | 1 | 1 |
| | sdhB | succinate dehydrogenase (ubiquinone) iron-sulfur subunit | arCOG00962 | 41SAW@651137\|Thaumarchaeota | | Nmar_0261 | 1 | 1 | 1 | 1 | 1 | 1 | 1 | 1 | 1 | 1 | 1 | 1 | 1 | 1 | 1 |
| | sdhC | succinate dehydrogenase (ubiquinone) cytochrome b560 subunit | arCOG02244 | 41SKZ@651137\|Thaumarchaeota | | Nmar_0259 | 1 | 1 | 1 | 1 | 1 | 1 | 1 | 1 | 1 | 1 | 1 | 1 | 1 | 1 | 1 |
| | sdhD | succinate dehydrogenase (ubiquinone) membrane anchor subunit | arCOG04162 | 41SRA@651137\|Thaumarchaeota | | Nmar_0260 | 1 | 1 | 1 | 1 | 1 | 1 | 1 | 1 | 1 | 1 | 1 | 1 | 1 | 1 | 1 |
| | fumC | Fumarase C | arCOG01750 | 41S9C@651137\|Thaumarchaeota | Fumarase C C-terminus | Nmar_1260 | 1 | 1 | 1 | 1 | 1 | 1 | 1 | 1 | 1 | 1 | 1 | 1 | 1 | 1 | 1 |
| | mdh | malate dehydrogenase | arCOG00246 | 41S7Z@651137\|Thaumarchaeota | | Nmar_0338, Nmar_0676 | 1 | 1 | 1 | 1 | 1 | 1 | 2 | 1 | 2 | 1 | 1 | 1 | 2 | 2 | 2 |
| | Gluconeogenesis | | | | | | | | | | | | | | | | | | | | |
| | pckA | ATP-dependent phosphoenolpyruvate carboxykinase | arCOG06073 | 41SE5@651137\|Thaumarchaeota | | Nmar_0392 | 1 | 0 | 0 | 1 | 1 | 1 | 1 | 1 | 1 | 1 | 1 | 1 | 1 | 1 | 1 |
| | ppdk | pyruvate, phosphate dikinase | arCOG01111 | 41SYV@651137\|Thaumarchaeota | Catalyzes the phosphorylation of pyruvate to phosphoenolpyruvate | Nmar_0951 | 1 | 1 | 1 | 1 | 1 | 1 | 1 | 1 | 1 | 1 | 2 | 1 | 1 | 1 | 1 |
| | eno | enolase | arCOG01169 | 41SE9@651137\|Thaumarchaeota | essential for the degradation of carbohydrates via glyco | Nmar_0317 | 1 | 1 | 1 | 1 | 1 | 1 | 1 | 1 | 1 | 1 | 1 | 1 | 1 | 1 | 1 |
| | apgM | 2,3-bisphosphoglycerate-independent phosphoglycerate mutase | arCOG01696 | 41S6M@651137\|Thaumarchaeota | | Nmar_0537 | 1 | 1 | 1 | 1 | 1 | 1 | 1 | 1 | 1 | 1 | 1 | 1 | 1 | 1 | 1 |
| | gpmB | 2,3-bisphosphoglycerate-dependent phosphoglycerate mutase | arCOG01991 | 41SJM@651137\|Thaumarchaeota | Phosphoglycerate mutase | Nmar_0181 | 1 | 1 | 1 | 1 | 1 | 1 | 1 | 1 | 1 | 1 | 1 | 1 | 1 | 1 | 1 |
| | pgk | phosphoglycerate kinase | arCOG00496 | 41SMB@651137\|Thaumarchaeota | | Nmar_0500 | 1 | 1 | 1 | 1 | 1 | 1 | 1 | 1 | 1 | 1 | 1 | 1 | 1 | 1 | 1 |
| | (gap) | putative glyceraldehyde-3-phosphate dehydrogenase, phosphorylating | arCOG00493 | 41SDF@651137\|Thaumarchaeota | | Nmar_0831 | 1 | 1 | 1 | 1 | 1 | 1 | 1 | 1 | 1 | 1 | 1 | 1 | 1 | 1 | 1 |
| | tpiA | triosephosphate isomerase | arCOG01087 | 41SHT@651137\|Thaumarchaeota | | Nmar_0950 | 0 | 1 | 0 | 0 | 0 | 0 | 1 | 1 | 1 | 1 | 1 | 1 | 1 | 1 | 1 |
| | fbp | bifunctional fructose-1,6-bisphosphate aldolase/phosphatase | arCOG04180 | 41S67@651137\|Thaumarchaeota | | Nmar_1035 | 1 | 1 | 1 | 1 | 1 | 1 | 1 | 1 | 1 | 1 | 1 | 1 | 1 | 1 | 1 |
| | pgi | phosphoglucose isomerase | arCOG00052 | 41SYN@651137\|Thaumarchaeota | | | 1 | 1 | 1 | 1 | 1 | 1 | 1 | 1 | 0 | 1 | 0 | 1 | 1 | 1 | 0 |
| | pgi | phosphoglucose isomerase | arCOG00052 | arCOG00052@2157\|Archaea | | Nmar_0167 | 0 | 0 | 0 | 0 | 0 | 0 | 0 | 0 | 0 | 0 | 0 | 0 | 0 | 0 | 1 |
| | Non-oxidative pentose-phosphate pathway | | | | | | | | | | | | | | | | | | | | |
| | rpiA | ribose-5-phosphate isomerase | arCOG01122 | 41SJE@651137\|Thaumarchaeota | | Nmar_0218 | 1 | 1 | 1 | 1 | 0 | 1 | 1 | 1 | 1 | 0 | 1 | 1 | 1 | 1 | 1 |
| | tktA+rpe | ribulose-phosphate 3-epimerase protein with N-terminal transketolase domain | arCOG01053 / arCOG05046 | 41SBE@651137\|Thaumarchaeota | PFAM Transketolase domain protein / PFAM Ribulose-phosphate 3-epime | Nmar_0299 | 1 | 1 | 1 | 1 | 1 | 1 | 1 | 1 | 1 | 1 | 1 | 1 | 1 | 1 | 1 |
| | tktB | transketolase C-terminal subunit | arCOG01051 | 41SEP@651137\|Thaumarchaeota | | Nmar_0300 | 1 | 1 | 1 | 1 | 1 | 1 | 1 | 1 | 1 | 1 | 1 | 1 | 1 | 1 | 1 |
| | tal | transaldolase | arCOG05061 | 41S99@651137\|Thaumarchaeota | | Nmar_0301 | 1 | 1 | 1 | 1 | 1 | 1 | 1 | 1 | 1 | 1 | 1 | 1 | 1 | 1 | 1 |
| | prs | ribose-phosphate pyrophosphokinase | arCOG00067 | 41SH1@651137\|Thaumarchaeota | | Nmar_1774 | 0 | 1 | 1 | 1 | 1 | 0 | 1 | 1 | 1 | 1 | 1 | 1 | 1 | 1 | 1 |
| Branch-chain amino acid transporter | livF | ABC-type branched-chain amino acid transport systems ATPase component | arCOG00924 | 23REP@183939\|Methanococci | | | 1 | 1 | 0 | 0 | 0 | 1 | 0 | 1 | 0 | 0 | 0 | 0 | 0 | 0 | 0 |
| | livF | ABC-type branched-chain amino acid transport systems ATPase component | arCOG00924 | 245ZE@183980\|Archaeoglobi | | | 0 | 0 | 0 | 0 | 0 | 0 | 0 | 0 | 0 | 1 | 1 | 0 | 0 | 0 | 0 |
| | livF | ABC-type branched-chain amino acid transport systems ATPase component | arCOG00924 | 2XQDQ@28889\|Crenarchaeota | | | 0 | 0 | 0 | 0 | 1 | 0 | 2 | 0 | 0 | 0 | 0 | 1 | 0 | 0 | 0 |
| | livG | COG0411 ABC-type branched-chain amino acid transport systems ATPase | arCOG00925 | arCOG00925@2157\|Archaea | | | 0 | 0 | 0 | 0 | 0 | 1 | 2 | 0 | 0 | 0 | 0 | 0 | 0 | 0 | 0 |
| | livG | COG0411 ABC-type branched-chain amino acid transport systems ATPase | arCOG00925 | 2XQAX@28889\|Crenarchaeota | | | 1 | 1 | 1 | 1 | 1 | 0 | 0 | 0 | 0 | 0 | 0 | 0 | 0 | 0 | 0 |
| | livG | COG0411 ABC-type branched-chain amino acid transport systems ATPase | arCOG00925 | 2XSVZ@28890\|Euryarchaeota,2461J@183980\|Archaeoglobi | | | 0 | 0 | 1 | 0 | 0 | 0 | 0 | 1 | 0 | 0 | 1 | 0 | 0 | 0 | 0 |
| | livK | COG0683 ABC-type branched-chain amino acid transport systems, periplasmic | arCOG01021 | 2XQA8@28889\|Crenarchaeota | | | 0 | 1 | 0 | 0 | 0 | 0 | 1 | 0 | 0 | 1 | 0 | 1 | 0 | 0 | 0 |
| | livK | COG0683 ABC-type branched-chain amino acid transport systems, periplasmic | arCOG01021 | arCOG01021@2157\|Archaea | | | 0 | 0 | 0 | 0 | 0 | 0 | 0 | 0 | 0 | 0 | 1 | 0 | 0 | 0 | 0 |
| | livH | COG0559 Branched-chain amino acid ABC-type transport system, permease | arCOG01269 | 23RCV@183939\|Methanococci | | | 0 | 0 | | 0 | 0 | 1 | 2 | 0 | 0 | 1 | 1 | 0 | 0 | 0 | 0 |
| | livH | COG0559 Branched-chain amino acid ABC-type transport system, permease | arCOG01269 | 245U7@183980\|Archaeoglobi | | | 0 | 0 | 0 | 0 | 1 | 0 | 0 | 0 | 0 | 0 | 0 | 1 | 0 | 0 | 0 |
| | livH | COG0559 Branched-chain amino acid ABC-type transport system, permease | arCOG01269 | 2XQXU@28889\|Crenarchaeota | | | 1 | 1 | 1 | 1 | 0 | 0 | 0 | 1 | 0 | 0 | 0 | 0 | 0 | 0 | 0 |
| | livH | COG0559 Branched-chain amino acid ABC-type transport system, permease | arCOG01269 | 244XY@183968\|Thermococci | | | 0 | 1 | 0 | 0 | 0 | 0 | 0 | 0 | 0 | 0 | 1 | 0 | 0 | 0 | 0 |
| | livM | Branched-chain amino acid transport system, permease component | arCOG01273 | 245V9@183980\|Archaeoglobi | | | 0 | 1 | 1 | 1 | 0 | 0 | 2 | 1 | 0 | 0 | 0 | 1 | 0 | 0 | 0 |
| | livM | Branched-chain amino acid transport system, permease component | arCOG01273 | arCOG01273@2157\|Archaea | | | 0 | 0 | 0 | 0 | 0 | 1 | 0 | 0 | 0 | 1 | 0 | 0 | 0 | 0 | 0 |
| Electron transport chain | F420/FMN/Flavin-dependent oxidoreductases | | | | | | | | | | | | | | | | | | | | |
| | | Luciferase-like monooxygenase | arCOG02410 | 41SBM@651137\|Thaumarchaeota | | Nmar_1109 | 0 | 0 | 1 | 1 | 1 | 1 | 1 | 1 | 1 | 1 | 1 | 1 | 1 | 1 | 1 |
| | | Luciferase-like monooxygenase | arCOG02410 | 41SC0@651137\|Thaumarchaeota | | Nmar_0842 | 1 | 1 | 1 | 1 | 1 | 1 | 1 | 1 | 1 | 1 | 0 | 1 | 1 | 1 | 1 |
| | npdG | NADP oxidoreductase coenzyme F420-dependent | arCOG00457 | 41SH2@651137\|Thaumarchaeota | | Nmar_0180 | 1 | 1 | 1 | 1 | 1 | 1 | 1 | 1 | 1 | 1 | 1 | 1 | 1 | 1 | 1 |
| | | putative F420H(2)-dependent quinone reductase | arCOG07802 | 41T6C@651137\|Thaumarchaeota | Pfam:DUF385, F420H2_quin_red | Nmar_0473 | 1 | 1 | 1 | 1 | 1 | 0 | 1 | 1 | 1 | 0 | 1 | 1 | 1 | 1 | 1 |
| | 11 subunit version of Complex I: type I NADH dehydrogenase (nuoEFG absent) | | | | | | | | | | | | | | | | | | | | |
| | nuoA | NADH-quinone oxidoreductase, subunit A | arCOG01557 | 41SMI@651137\|Thaumarchaeota | | Nmar_0276 | 1 | 1 | 1 | 1 | 1 | 1 | 1 | 1 | 1 | 1 | 1 | 1 | 1 | 1 | 1 |
| | nuoB | NADH-quinone oxidoreductase, subunit B | arCOG01554 | 41S6D@651137\|Thaumarchaeota | | Nmar_0277 | 1 | 1 | 1 | 1 | 1 | 1 | 1 | 1 | 1 | 1 | 1 | 1 | 1 | 1 | 1 |
| | nuoC | NADH-quinone oxidoreductase, subunit C | arCOG01551 | 41SJS@651137\|Thaumarchaeota | | Nmar_0278 | 1 | 1 | 1 | 1 | 1 | 1 | 1 | 1 | 1 | 1 | 0 | 1 | 1 | 1 | 1 |
| | nuoD | NADH-quinone oxidoreductase, subunit D | arCOG01548 | 41S62@651137\|Thaumarchaeota | | Nmar_0279 | 1 | 1 | 1 | 1 | 1 | 1 | 1 | 1 | 1 | 1 | 1 | 1 | 1 | 1 | 1 |
| | nuoH | NADH-quinone oxidoreductase, subunit H | arCOG01546 | 41SAT@651137\|Thaumarchaeota | | Nmar_0280 | 1 | 1 | 1 | 1 | 1 | 1 | 1 | 1 | 1 | 1 | 1 | 1 | 1 | 1 | 1 |
| | nuoI | NADH-quinone oxidoreductase, subunit I | arCOG01543 | 41SGW@651137\|Thaumarchaeota | | Nmar_0281 | 1 | 1 | 1 | 1 | 1 | 1 | 1 | 1 | 1 | 1 | 1 | 1 | 1 | 1 | 1 |
| | nuoJ | NADH-quinone oxidoreductase, subunit J | arCOG04654 | 41SM8@651137\|Thaumarchaeota | | Nmar_0282 | 1 | 1 | 1 | 1 | 1 | 1 | 1 | 1 | 1 | 1 | 1 | 1 | 1 | 1 | 1 |
| | nuoK | NADH-quinone oxidoreductase, subunit K | arCOG03073 | 41SPM@651137\|Thaumarchaeota | | Nmar_0283 | 1 | 1 | 1 | 1 | 1 | 1 | 1 | 1 | 1 | 1 | 1 | 1 | 1 | 1 | 1 |
| | nuoM | NADH-quinone oxidoreductase, subunit M | arCOG01537 | 41SAV@651137\|Thaumarchaeota | | Nmar_0284 | 1 | 1 | 1 | 1 | 0 | 1 | 0 | 0 | 0 | 1 | 2 | 1 | 3 | 1 | 1 |
| | nuoL | NADH-quinone oxidoreductase, subunit L | arCOG01539 | 41SC4@651137\|Thaumarchaeota | | Nmar_0285 | 1 | 1 | 1 | 1 | 1 | 1 | 1 | 1 | 1 | 1 | 2 | 1 | 1 | 1 | 1 |
| | | NADH-quinone oxidoreductase, subunit L | arCOG01539 | 41TBF@651137\|Thaumarchaeota | | | 0 | 0 | 0 | 0 | 1 | 0 | 1 | 1 | 1 | 0 | 0 | 0 | 0 | 0 | 0 |
| | nuoN | NADH-quinone oxidoreductase, subunit N | arCOG01540 | 41SDU@651137\|Thaumarchaeota | | Nmar_0286 | 1 | 1 | 1 | 1 | 1 | 1 | 1 | 1 | 1 | 1 | 2 | 1 | 1 | 1 | 1 |
| | Complex II | | | | | | | | | | | | | | | | | | | | |
| | sdhA | succinate dehydrogenase flavoprotein subunit/fumarate reductase | arCOG00571 | 41SAM@651137\|Thaumarchaeota | | Nmar_0258 | 1 | 1 | 1 | 1 | 1 | 1 | 1 | 1 | 1 | 1 | 1 | 1 | 1 | 1 | 1 |
| | (sdhC) | putative succinate dehydrogenase/fumarate reductase | arCOG02244 | 41SKZ@651137\|Thaumarchaeota | | Nmar_0259 | 1 | 1 | 1 | 1 | 1 | 1 | 1 | 1 | 1 | 1 | 1 | 1 | 1 | 1 | 1 |
| | sdhD | succinate dehydrogenase | arCOG04162 | 41SRA@651137\|Thaumarchaeota | | Nmar_0260 | 1 | 1 | 1 | 1 | 1 | 1 | 1 | 1 | 1 | 1 | 1 | 1 | 1 | 1 | 1 |
| | sdhB | FeS-center protein of succinate dehydrogenase/fumarate reductase | arCOG00962 | 41SAW@651137\|Thaumarchaeota | TIGRFAM succinate dehydrogenase and fumarate reductase iron-sulfur pr | Nmar_0261 | 1 | 1 | 1 | 1 | 1 | 1 | 1 | 1 | 1 | 1 | 1 | 1 | 1 | 1 | 1 |
| | Complex III: 1.10.2.2, quinol:ferricytochrome-c oxidoreductase, bc1 complex | | | | | | | | | | | | | | | | | | | | |
| | soxC | Cytochrome b subunit of the bc complex | arCOG01721 | 41SDA@651137\|Thaumarchaeota | cytochrome b6-f complex iron-sulfur unit | Nmar_1543 | 1 | 1 | 1 | 1 | 1 | 1 | 1 | 1 | 1 | 1 | 1 | 1 | 1 | 1 | 1 |
| | | PFAM Rieske 2Fe-2S domain | arCOG01720 | 41SCE@651137\|Thaumarchaeota | | Nmar_1544 | 1 | 1 | 1 | 1 | 1 | 1 | 1 | 1 | 1 | 1 | 1 | 1 | 1 | 1 | 1 |
| | Complex IV | | | | | | | | | | | | | | | | | | | | |
| | | putative Heme/copper-type cytochrome/quinol oxidase, subunit | arCOG08824 | 41SW7@651137\|Thaumarchaeota | | Nmar_0182 | 1 | 1 | 1 | 1 | 1 | 1 | 1 | 1 | 1 | 1 | 1 | 1 | 1 | 1 | 1 |
| | soxA | Heme copper-type cytochrome quinol oxidases, subunit 2 | arCOG01236 | 41SMY@651137\|Thaumarchaeota | | Nmar_0183 | 1 | 1 | 1 | 1 | 1 | 1 | 1 | 1 | 1 | 1 | 1 | 1 | 1 | 1 | 1 |
| | coxA | Heme copper-type cytochrome quinol | arCOG01237 | 41S9Z@651137\|Thaumarchaeota | | Nmar_0184 | 1 | 1 | 1 | 1 | 1 | 1 | 1 | 1 | 1 | 1 | 1 | 1 | 1 | 1 | 1 |
| | | Cytochrome oxidase assembly protein | arCOG03103 | 41SNT@651137\|Thaumarchaeota | | Nmar_0186 | 1 | 1 | 1 | 0 | 1 | 1 | 1 | 1 | 1 | 1 | 1 | 1 | 1 | 1 | 1 |
| | Complex V: A1A0-type ATPse. | | | | | | | | | | | | | | | | | | | | |
| | atpK | ATP synthase subunit C | arCOG02455 | 41T6A@651137\|Thaumarchaeota | | Nmar_1688 | 1 | 1 | 1 | 1 | 0 | 1 | 1 | 0 | 1 | 1 | 0 | 1 | 1 | 0 | 1 |
| | atpD | Produces ATP from ADP in the presence of a proton gradient across the | arCOG04101 | 41T0N@651137\|Thaumarchaeota | | Nmar_1689 | 1 | 1 | 1 | 1 | 1 | 1 | 1 | 1 | 1 | 1 | 1 | 1 | 1 | 1 | 1 |
| | atpB | Produces ATP from ADP in the presence of a proton gradient across the | arCOG00865 | 41SPY@651137\|Thaumarchaeota | | Nmar_1690 | 1 | 1 | 1 | 1 | 1 | 1 | 1 | 1 | 1 | 1 | 1 | 1 | 1 | 1 | 1 |
| | atpA | Produces ATP from ADP in the presence of a proton gradient across the | arCOG00868 | 41SZU@651137\|Thaumarchaeota | | Nmar_1691 | 1 | 1 | 1 | 1 | 1 | 1 | 1 | 1 | 1 | 1 | 1 | 1 | 1 | 1 | 1 |
| | atpE | Produces ATP from ADP in the presence of a proton gradient across the | arCOG00869 | 41T4W@651137\|Thaumarchaeota | | Nmar_1692 | 1 | 1 | 1 | 1 | 1 | 1 | 1 | 1 | 1 | 1 | 1 | 1 | 1 | 1 | 1 |
| | atpI | Belongs to the V-ATPase 116 kDa subunit family | arCOG04138 | 41SRH@651137\|Thaumarchaeota | | Nmar_1693 | 1 | 1 | 1 | 1 | 1 | 1 | 1 | 1 | 1 | 1 | 1 | 1 | 1 | 1 | 1 |

**Table S8:** Functional annotation of hallmark genes for chemolithoautotrophic nitrite-oxidation and carbon fixation.

| Category | Function | Gene | Description | COG | Nitrospirae eggnog_OG | notes | Nide_loci to compare | Nitrospira sp. sponge CO36405bin_1.protein | Nitrospirales sponge GCA_022824735.1_AS | Nitrospirales sponge GCA_022825885.1_AS | Nitrospirales sponge GCA_022828225.1_AS | Nitrospira sp. bin75 of sponge GCA_002238765.1_AS | Nitrospirales sponge GCA_007570995.1_AS | Nitrospira_sp._SB0662 sponge GCA_009840285.1_AS | Nitrospira_sp._SB0677 sponge GCA_009842645.1_AS | Nitrospirales coral GCA_012270585.1_AS | Nitrospirales cold seep GCA_020027205.1_AS | Nitrospirales cold seep GCA_020027535.1_AS | Ca. Nitrospira marine Nitrospira_denmark_2 | Nitrospira sp. MA-1 marine Nitrospira_maine_202 | Nitrospira marina Nb- marine Nitrospira_marina_ge | Ca. Nitronereus marine Nitrospira_elba_2022 | Nitrospira_vancouver marine Nitrospira_vancouver | Ca. Nitrospira terrestrial NSPALKALI_genomic.p |
|---|---|---|---|---|---|---|---|---|---|---|---|---|---|---|---|---|---|---|---|---|---|---|---|---|
| Nitrogen metabolism | Nitrite oxidation | nxrA | Nitrite oxidoreductase, alpha subunit | COG0243 | | | **Nide3255**, Nide3237 | 1 | 1 | 1 | 1 | 1 | 1 | 0 | 1 | 1 | 1 | 1 | 3 | 3 | 3 | 3 | 3 | 2 |
| | Nitrite oxidation | nxrB | Nitrite oxidoreductase, beta subunit | COG1140 | **3J1FU@40117\|Nitrospirae** | | **Nide3256**, Nide3236 | 1 | 1 | 1 | 1 | 1 | 1 | 0 | 1 | 1 | 1 | 1 | 3 | 3 | 3 | 3 | 3 | 1 |
| | Nitrite oxidation | nxrC | Nitrite oxidoreductase, gamma subunit | **COG2180** | | | **Nide3271, Nide3278** | 1 | 3 | 5 | 2 | 2 | 2 | 1 | 1 | 2 | 2 | 3 | 3 | 3 | 3 | 3 | 3 | 2 |
| | Nitrite oxidation | alt_nxrC1 | Nitrite oxidoreductase, putative membrane subunit | 2ERPD@1\|root,33J8R@2\|Bacteria | | | **Nide0904** | 1 | 1 | 1 | 1 | 1 | 1 | 1 | 1 | 1 | 1 | 1 | 1 | 1 | 1 | 1 | 1 | 1 |
| | Nitrite oxidation | alt_nxrC2 | Nitrite oxidoreductase, putative membrane subunit | COG2010 | | | **Nide3293** | 1 | 1 | 1 | 1 | 1 | 1 | 1 | 1 | 1 | 1 | 1 | 1 | 1 | 1 | 1 | 1 | 1 |
| | Nitrite oxidation | napG | Ferredoxin-type protein NapG | COG0437 | | | **Nide3280** | 1 | 1 | 1 | 1 | 1 | 1 | 1 | 1 | 1 | 1 | 1 | 1 | 1 | 1 | 1 | 1 | 1 |
| | Nitrite reduction | nirK | copper-containing Nitrite reductase (NO forming) | COG3794 | | EC:1.7.2.2, PFAM blue ( | **Nide1412** | 2 | 1 | 1 | 2 | 1 | 0 | 2 | 1 | 2 | 0 | 2 | 3 | 2 | 3 | 4 | 3 | 3 |
| | Nitrite reduction | nirK | copper-containing Nitrite reductase (NO forming) | COG3794 | | EC:1.7.2.2, PFAM blue ( | **Nide1926** | 0 | 0 | 0 | 0 | 1 | 0 | 1 | 0 | 0 | 0 | 0 | 1 | 1 | 1 | 1 | 1 | 2 |
| | Nitrite reduction | nirK | copper-containing Nitrite reductase (NO forming) | COG2132 | | EC:1.7.2.2, Multicopper | Nide2534 | 0 | 0 | 0 | 0 | 0 | 0 | 0 | 0 | 0 | 0 | 0 | 0 | 0 | 0 | 1 | 0 | 0 |
| | Nitrite reduction/Nitrogen assimilation | nirA | Ferredoxin-nitrite reductase | COG0155 | 3J10S@40117\|Nitrospirae, 3J11D@40117\|Nitrospirae | | Nide1367, **Nide0407** | 1 | 1 | 1 | 1 | 2 | 1 | 1 | 1 | 1 | 1 | 2 | 2 | 2 | 2 | 2 | 2 | 2 |
| | Nitrogen assimilation | gltB | glutamate synthase, alpha subunit | COG0067, COG0069,COG0070 | 3J0SV@40117\|Nitrospirae | | **Nide3384** | 1 | 1 | 1 | 1 | 1 | 1 | 1 | 1 | 1 | 1 | 1 | 1 | 1 | 1 | 1 | 1 | 1 |
| | Nitrogen assimilation | gltD | glutamate synthase, beta subunit | COG0493 | 3J152@40117\|Nitrospirae | | **Nide3383** | 1 | 1 | 1 | 1 | 1 | 1 | 1 | 1 | 1 | 1 | 1 | 1 | 1 | 1 | 1 | 1 | 1 |
| | Nitrogen assimilation | glnB | Nitrogen regulatory protein PII | COG0347 | 3J0P1@40117\|Nitrospirae | | Nide1361 | 0 | 0 | 0 | 0 | 0 | 1 | 0 | 0 | 0 | 1 | 0 | 1 | 1 | 1 | 1 | 1 | 1 |
| | Nitrogen assimilation | glnB? | Nitrogen regulatory protein PII | COG0347 | 3J17N@40117\|Nitrospirae | | Nide4390, Nide1771 | 1 | 1 | 1 | 1 | 0 | 0 | 1 | 1 | 1 | 0 | 0 | 1 | 1 | 1 | 1 | 1 | 1 |
| | Nitrogen assimilation | gdhA1 | glutamate dehydrogenase | COG0334 | | | **Nide0440** | 1 | 1 | 1 | 1 | 1 | 0 | 1 | 1 | 1 | 0 | 1 | 1 | 1 | 1 | 1 | 1 | 1 |
| | Nitrogen assimilation | glnD | putative (Protein-PII) uridylyltransferase GlnD | COG2844 | | | **Nide1376, Nide1362** | 2 | 2 | 2 | 2 | 2 | 1 | 2 | 2 | 2 | 1 | 2 | 2 | 2 | 2 | 2 | 2 | 2 |
| | Nitrogen assimilation | nadE | Glutamine-dependent NAD(+) synthase | COG0388, COG0171 | 3J0EB@40117\|Nitrospirae | | **Nide1358** | 1 | 1 | 1 | 1 | 1 | 1 | 1 | 1 | 1 | 1 | 1 | 1 | 1 | 1 | 1 | 1 | 1 |
| | Nitrogen assimilation | gdhA2 | putative glutamate dehydrogenase | COG0334 | | | Nide0440 | 1 | 1 | 1 | 1 | 1 | 0 | 1 | 1 | 1 | 0 | 1 | 1 | 1 | 1 | 1 | 1 | 1 |
| | alternative substrates | cynS | Cyanate hydratase | COG1513 | | | Nide1365 | 1 | 1 | 1 | 1 | 1 | 0 | 1 | 1 | 1 | 0 | 1 | 0 | 0 | 0 | 0 | 0 | 1 |
| | alternative substrates | ureA | urease subunit gamma | COG0831, COG0832 | | | - | 0 | 1 | 0 | 1 | 1 | 1 | 1 | 1 | 0 | 0 | 0 | 0 | 1 | 0 | 0 | 0 | 0 |
| | alternative substrates | ureB | urease subunit beta | COG0831, COG0832 | | | - | 0 | 0 | 1 | 1 | 1 | 1 | 1 | 0 | 0 | 0 | 0 | 0 | 1 | 0 | 0 | 0 | 0 |
| | alternative substrates | ureC | urease subunit alpha | COG0804 | | | - | 0 | 1 | 1 | 1 | 1 | 1 | 1 | 1 | 0 | 0 | 0 | 0 | 1 | 0 | 0 | 0 | 0 |
| | alternative substrates | UT | urea transporter | COG4413 | | | - | 0 | 0 | 0 | 0 | 0 | 0 | 0 | 0 | 0 | 0 | 0 | 0 | 1 | 0 | 0 | 0 | 0 |
| citric acid cycle/carbon fixation | o/rTCA cycle | pdhA | Pyruvate dehydrogenase E1 component, alpha subunit (acetyl-transferring) | COG1071 | | | Nide3953 | 0 | 0 | 0 | 0 | 0 | 0 | 0 | 0 | 0 | 0 | 0 | 1 | 1 | 1 | 0 | 1 | 1 |
| | o/rTCA cycle | pdhB | Pyruvate dehydrogenase E1 component, beta subunit (Transketolase) | COG0022 | | | Nide3952 | 0 | 0 | 0 | 0 | 0 | 0 | 0 | 0 | 0 | 0 | 0 | 1 | 1 | 1 | 0 | 1 | 1 |
| | o/rTCA cycle | pdhC | putative Dihydrolipoamide acetyltransferase (E2) component of pyruvate dehydrogenase complex | COG0508 | | | Nide3854, Nide3951 | 0 | 0 | 0 | 0 | 0 | 0 | 0 | 0 | 0 | 0 | 0 | 1 | 1 | 1 | 0 | 1 | 1 |
| | o/rTCA cycle | aceE | pyruvate dehydrogenase E1 component | COG2010 | | | Nide3293 | 1 | 1 | 1 | 1 | 1 | 1 | 1 | 1 | 1 | 1 | 1 | 1 | 1 | 1 | 1 | 1 | 1 |
| | o/rTCA cycle | aceF | Pyruvate dehydrogenase complex, dihydrolipoyllysine-residue acetyltransferase (E2) component | COG2010 | | | Nide3294 | 1 | 1 | 1 | 1 | 1 | 1 | 1 | 1 | 1 | 1 | 1 | 1 | 1 | 1 | 1 | 1 | 1 |
| | o/rTCA cycle | lpd | Dihydrolipoyl dehydrogenase, E3 component of Pyruvate and 2-oxoglutarate dehydrogenase complexes | COG1249 | 3J0IM@40117\|Nitrospirae | | Nide2727 | 1 | 1 | 1 | 1 | 1 | 1 | 1 | 1 | 1 | 1 | 1 | 1 | 1 | 1 | 0 | 1 | 1 |
| | o/rTCA cycle | porA | Pyruvate:ferredoxin oxidoreductase, alpha subunit | COG0674 | 3J0WN@40117\|Nitrospirae | | Nide0971 | 1 | 1 | 1 | 1 | 1 | 1 | 1 | 1 | 1 | 1 | 1 | 1 | 1 | 1 | 1 | 1 | 1 |
| | o/rTCA cycle | porB | Pyruvate:ferredoxin oxidoreductase, beta subunit | COG1013 | 3J0Z2@40117\|Nitrospirae | | Nide0970 | 1 | 1 | 1 | 1 | 1 | 1 | 1 | 1 | 1 | 1 | 1 | 1 | 1 | 1 | 1 | 1 | 1 |
| | o/rTCA cycle | porC | Pyruvate:ferredoxin oxidoreductase, gamma subunit | COG1014 | 3J0Y6@40117\|Nitrospirae | | Nide0969 | 1 | 1 | 1 | 1 | 1 | 1 | 1 | 1 | 1 | 1 | 1 | 1 | 1 | 1 | 1 | 1 | 1 |
| | o/rTCA cycle | porD | Pyruvate:ferredoxin oxidoreductase, delta subunit | COG0226 | 3J0WN@40117\|Nitrospirae | | Nide0972 | 1 | 1 | 1 | 1 | 1 | 1 | 1 | 1 | 1 | 1 | 1 | 1 | 1 | 1 | 1 | 1 | 1 |
| | o/rTCA cycle | porE | Pyruvate:ferredoxin oxidoreductase, epsilon subunit | COG1144 | 3J1DG@40117\|Nitrospirae | | Nide0968 | 0 | 0 | 1 | 1 | 1 | 1 | 0 | 0 | 1 | 1 | 1 | 1 | 1 | 1 | 1 | 1 | 1 |
| | o/rTCA cycle | porA2 | Pyruvate:ferredoxin oxidoreductase, alpha subunit | COG0674 | | | Nide3874 | 0 | 0 | 0 | 0 | 0 | 0 | 0 | 0 | 0 | 0 | 0 | 1 | 1 | 1 | 1 | 1 | 2 |
| | o/rTCA cycle | porB2 | Pyruvate:ferredoxin oxidoreductase, beta subunit | COG1013 | | | Nide3873 | 0 | 0 | 0 | 0 | 0 | 0 | 0 | 0 | 0 | 0 | 0 | 1 | 1 | 1 | 1 | 1 | 2 |
| | o/rTCA cycle | porC2 | Pyruvate:ferredoxin oxidoreductase, gamma subunit | COG1014 | 3J0TG@40117\|Nitrospirae | | Nide3875 | 0 | 0 | 0 | 0 | 0 | 0 | 0 | 0 | 0 | 0 | 0 | 1 | 1 | 1 | 1 | 1 | 1 |
| | o/rTCA cycle | mdh | Malate dehydrogenase | COG0039 | 3J0EG@40117\|Nitrospirae | | Nide3562 | 1 | 1 | 1 | 1 | 1 | 1 | 1 | 0 | 1 | 1 | 1 | 1 | 1 | 1 | 1 | 1 | 1 |
| | o/rTCA cycle | mdh2 | Malate dehydrogenase | COG0039 | | | | 0 | 0 | 0 | 0 | 0 | 0 | 0 | 1 | 0 | 0 | 0 | 0 | 0 | 0 | 1 | 0 | 0 |
| | o/rTCA cycle | fumC | Fumarate hydratase, class II | COG0114 | 3J0FU@40117\|Nitrospirae | | Nide2454 | 0 | 0 | 0 | 0 | 0 | 0 | 0 | 0 | 0 | 0 | 0 | 1 | 1 | 1 | 0 | 1 | 1 |
| | o/rTCA cycle | fumA | Fumarate hydratase | COG1838 | | | | 1 | 1 | 0 | 1 | 1 | 0 | 1 | 1 | 1 | 1 | 1 | 1 | 1 | 1 | 1 | 1 | 0 |
| | o/rTCA cycle | sucC | Succinyl-CoA synthetase, beta subunit | COG0045 | 3J0ZB@40117\|Nitrospirae | | Nide0840 | 1 | 0 | 1 | 1 | 1 | 1 | 1 | 1 | 1 | 1 | 1 | 1 | 1 | 1 | 1 | 1 | 1 |
| | o/rTCA cycle | sucD | Succinyl-CoA synthetase, NAD(P)-binding, alpha subunit | COG0074 | 3J0WZ@40117\|Nitrospirae | | Nide0841 | 0 | 0 | 1 | 1 | 0 | 1 | 0 | 0 | 0 | 1 | 0 | 1 | 1 | 1 | 1 | 1 | 1 |
| | o/rTCA cycle | forB | 2-oxoglutarate:ferredoxin oxidoreductase, beta subunit | COG1013 | 3J10F@40117\|Nitrospirae | | Nide0823 | 1 | 1 | 1 | 1 | 1 | 1 | 1 | 1 | 1 | 1 | 1 | 1 | 1 | 1 | 1 | 1 | 1 |
| | o/rTCA cycle | forC | 2-oxoglutarate:ferredoxin oxidoreductase, gamma subunit | COG1014 | 3J0YE@40117\|Nitrospirae | | Nide0824 | 1 | 1 | 1 | 1 | 1 | 1 | 1 | 1 | 1 | 1 | 1 | 1 | 1 | 1 | 1 | 1 | 1 |
| | o/rTCA cycle | forE | 2-oxoglutarate:ferredoxin oxidoreductase, epsilon subunit | COG1144 | 3J1A9@40117\|Nitrospirae | | Nide0825 | 1 | 1 | 1 | 1 | 1 | 1 | 1 | 1 | 1 | 1 | 1 | 1 | 1 | 1 | 1 | 1 | 1 |
| | o/rTCA cycle | forD | 2-oxoglutarate:ferredoxin oxidoreductase, delta subunit | COG0226 | 3J0YH@40117\|Nitrospirae | | Nide0826 | 1 | 1 | 1 | 1 | 1 | 1 | 1 | 1 | 1 | 1 | 1 | 1 | 1 | 1 | 1 | 1 | 1 |
| | o/rTCA cycle | forA | 2-oxoglutarate:ferredoxin oxidoreductase, alpha subunit | COG0674 | 3J105@40117\|Nitrospirae | | Nide0827 | 1 | 1 | 1 | 1 | 1 | 1 | 1 | 1 | 1 | 1 | 1 | 1 | 1 | 1 | 1 | 1 | 1 |
| | o/rTCA cycle | korA | 2-oxoglutarate oxidoreductase subunit KorA | COG0674, COG1014 | | | | 0 | 0 | 0 | 0 | 0 | 0 | 0 | 0 | 0 | 0 | 0 | 0 | 0 | 0 | 1 | 0 | 0 |
| | o/rTCA cycle | korB | 2-oxoglutarate oxidoreductase subunit KorB | | | | | 0 | 0 | 0 | 0 | 0 | 0 | 0 | 0 | 0 | 0 | 0 | 0 | 0 | 0 | 1 | 0 | 0 |
| | o/rTCA cycle | icd | Isocitrate dehydrogenase (NADP(+)) | COG2838 | | | Nide0837 | 1 | 1 | 1 | 1 | 1 | 1 | 1 | 1 | 1 | 1 | 1 | 1 | 1 | 1 | 1 | 1 | 1 |
| | o/rTCA cycle | acnA | Aconitate hydratase | COG1048 | 3J0E6@40117\|Nitrospirae | | Nide0836 | 1 | 1 | 1 | 1 | 1 | 1 | 1 | 1 | 1 | 1 | 1 | 1 | 1 | 1 | 1 | 1 | 1 |
| | o/rTCA cycle | gltA | Citrate synthase | COG0372 | | | Nide2451 | 1 | 0 | 1 | 1 | 1 | 0 | 1 | 1 | 0 | 0 | 1 | 1 | 1 | 1 | 2 | 1 | 1 |
| | o/rTCA cycle | ppsA | putative Phosphoenolpyruvate synthase | COG0574, COG1080 | 3J101@40117\|Nitrospirae | | Nide0497 | 1 | 1 | 1 | 1 | 1 | 1 | 1 | 1 | 1 | 1 | 1 | 1 | 1 | 1 | 1 | 1 | 1 |
| | o/rTCA cycle | aclB | ATP-citrate lyase, beta subunit | COG0045 | 3J185@40117\|Nitrospirae | | Nide0834 | 1 | 1 | 1 | 1 | 1 | 1 | 1 | 1 | 1 | 1 | 1 | 1 | 1 | 1 | 1 | 1 | 1 |
| | o/rTCA cycle | aclA | ATP citrate lyase, alpha subunit | COG0074, COG0372 | | | Nide0835 | 1 | 1 | 1 | 1 | 1 | 1 | 1 | 1 | 1 | 1 | 1 | 1 | 1 | 1 | 1 | 1 | 1 |
| | o/rTCA cycle | pycB | Pyruvate carboxylase, subunit B | COG0511, COG5016 | 3J10I@40117\|Nitrospirae | | Nide1204 | 1 | 1 | 1 | 1 | 1 | 1 | 1 | 1 | 1 | 1 | 1 | 1 | 1 | 1 | 1 | 1 | 1 |
| | o/rTCA cycle | pycA | Pyruvate carboxylase, subunit A | COG0439 | 3J0YP@40117\|Nitrospirae | | Nide1205 | 1 | 1 | 1 | 0 | 1 | 1 | 0 | 1 | 1 | 1 | 1 | 1 | 1 | 1 | 1 | 1 | 1 |
| | o/rTCA cycle | forB | 2-oxoacid:ferredoxin oxidoreductase beta subunit | COG1013 | 3J10F@40117\|Nitrospirae | | Nide1465 | 0 | 0 | 0 | 0 | 0 | 0 | 0 | 0 | 0 | 0 | 0 | 0 | 0 | 0 | 0 | 0 | 0 |
| | o/rTCA cycle | lpd | putative Dihydrolipoyl dehydrogenase, E3 component of pyruvate and 2-oxoglutarate dehydrogenase complexes | COG1249 | 3J11P@40117\|Nitrospirae | | Nide4341 | | | | | | | | | | | | | | | | | |
| **Alternative metabolism** | | | | | | | | | | | | | | | | | | | | | | | | |
| | Formate oxidation | fdhD | formate dehydrogenase accessory protein FdhD | COG1526 | | | | 0 | 0 | 0 | 0 | 0 | 0 | 0 | 0 | 0 | 0 | 0 | 1 | 1 | 1 | 0 | 1 | 1 |
| | Formate oxidation | fdhA | putative formate dehydrogenase | COG1063 | | | | 0 | 0 | 0 | 0 | 0 | 0 | 0 | 0 | 0 | 0 | 0 | 1 | 1 | 1 | 0 | 1 | 0 |
| | hydrogen oxidation | hypE | Hydrogenase maturation protein HypE | COG0309 | | | | 0 | 0 | 0 | 0 | 0 | 0 | 0 | 0 | 0 | 0 | 0 | 1 | 1 | 1 | 1 | 1 | 1 |
| | hydrogen oxidation | hypD | Hydrogenase maturation protein HypD | COG0409 | | | | 0 | 0 | 0 | 0 | 0 | 0 | 0 | 0 | 0 | 0 | 0 | 1 | 1 | 1 | 1 | 1 | 1 |
| | hydrogen oxidation | hypC | Hydrogenase assembly chaperone HypC | COG0298 | | | | 0 | 0 | 0 | 0 | 0 | 0 | 0 | 0 | 0 | 0 | 0 | 1 | 1 | 1 | 1 | 1 | 1 |
| | hydrogen oxidation | hypF | Hydrogenase maturation protein HypF | COG0068 | | | | 0 | 0 | 0 | 0 | 0 | 0 | 0 | 0 | 0 | 0 | 0 | 1 | 1 | 1 | 1 | 1 | 1 |
| | hydrogen oxidation | hypA | Hydrogenase nickel incorporation protein HypA | COG0375 | | | | 0 | 0 | 0 | 0 | 0 | 0 | 0 | 0 | 0 | 0 | 0 | 1 | 1 | 1 | 1 | 1 | 1 |
| | hydrogen oxidation | hypB | hydrogenase isoenzymes nickel incorporation protein HypB | COG0378 | | | | 0 | 0 | 0 | 0 | 0 | 0 | 0 | 0 | 0 | 0 | 0 | 1 | 1 | 1 | 0 | 1 | 1 |
| | hydrogen oxidation | hybD | Hydrogenase maturation protease HybD | | | | | 0 | 0 | 0 | 0 | 0 | 0 | 0 | 0 | 0 | 0 | 0 | 0 | 0 | 1 | 1 | 0 | 1 |
| | hydrogen oxidation | hydA | Nickel-dependent hydrogenase, alpha subunit | | 3b [NiFe] hydrogenase | | | 0 | 0 | 0 | 0 | 0 | 0 | 0 | 0 | 0 | 0 | 0 | 1 | 1 | 1 | 1 | 1 | 1 |
| | hydrogen oxidation | hydD | Nickel-dependent hydrogenase, delta subunit | | 3b [NiFe] hydrogenase | | | 0 | 0 | 0 | 0 | 0 | 0 | 0 | 0 | 0 | 0 | 0 | 1 | 1 | 1 | 1 | 1 | 1 |
| | hydrogen oxidation | hydC | Nickel-dependent hydrogenase, gamma subunit | | 3b [NiFe] hydrogenase | | | 0 | 0 | 0 | 0 | 0 | 0 | 0 | 0 | 0 | 0 | 0 | 1 | 1 | 1 | 1 | 1 | 1 |
| | hydrogen oxidation | hydB | Nickel-dependent hydrogenase, beta subunit | | 3b [NiFe] hydrogenase | | | 0 | 0 | 0 | 0 | 0 | 0 | 0 | 0 | 0 | 0 | 0 | 1 | 1 | 1 | 1 | 1 | 1 |
| | hydrogen oxidation | hupS | NiFe hydrogenase, small subunit | COG1740 | 2a [NiFe] hydrogenase | | | 0 | 0 | 0 | 0 | 0 | 0 | 0 | 0 | 0 | 0 | 0 | 0 | 0 | 0 | 0 | 0 | 1 |
| | hydrogen oxidation | hupL | NiFe hydrogenase, large subunit | COG0374 | 2a [NiFe] hydrogenase | | | 0 | 0 | 0 | 0 | 0 | 0 | 0 | 0 | 0 | 0 | 0 | 0 | 0 | 0 | 0 | 0 | 1 |
| Respiratory Chain | Complex I | nuoN | NADH-quinone oxidoreductase, membrane subunit N | COG1007 | 3J0FN@40117\|Nitrospirae | | Nide0602 | 1 | 1 | 1 | 1 | 1 | 1 | 1 | 1 | 1 | 1 | 1 | 1 | 1 | 1 | 1 | 1 | 1 |
| | Complex I | nuoM | NADH-quinone oxidoreductase, membrane subunit M | COG1008 | 3J0EQ@40117\|Nitrospirae | | Nide0603 | 1 | 1 | 1 | 1 | 1 | 1 | 1 | 1 | 1 | 1 | 0 | 1 | 1 | 1 | 1 | 1 | 1 |
| | Complex I | nuoL | NADH-quinone oxidoreductase, membrane subunit L | COG1009 | 3J0AN@40117\|Nitrospirae | | Nide0604 | 1 | 1 | 1 | 1 | 1 | 1 | 1 | 1 | 1 | 1 | 1 | 1 | 1 | 1 | 1 | 1 | 1 |
| | Complex I | nuoK | NADH-quinone oxidoreductase, membrane subunit K | COG0713 | 3J0SN@40117\|Nitrospirae | | Nide0605 | 1 | 1 | 1 | 1 | 1 | 1 | 1 | 1 | 1 | 1 | 1 | 1 | 1 | 1 | 1 | 1 | 1 |
| | Complex I | nuoJ | NADH-quinone oxidoreductase, membrane subunit J | COG0839 | 3J0RP@40117\|Nitrospirae | | Nide0606 | 1 | 1 | 1 | 1 | 1 | 1 | 1 | 1 | 1 | 1 | 1 | 1 | 1 | 1 | 1 | 1 | 1 |
| | Complex I | nuoI | NADH-quinone oxidoreductase, subunit I | COG1143 | 3J0N0@40117\|Nitrospirae | | Nide0607 | 1 | 1 | 1 | 1 | 1 | 1 | 1 | 1 | 1 | 1 | 1 | 1 | 1 | 1 | 1 | 1 | 1 |
| | Complex I | nuoG | putative NADH-quinone oxidoreductase, subunit G | COG3383 | 3J0UR@40117\|Nitrospirae | | Nide0609 | 1 | 1 | 1 | 1 | 1 | 0 | 1 | 1 | 1 | 1 | 1 | 1 | 1 | 1 | 1 | 1 | 1 |
| | Complex I | nuoC/D | NADH-quinone oxidoreductase, subunit C/D | COG0649, COG0852 | 3J0FM@40117\|Nitrospirae | | Nide0613 | 0 | 0 | 0 | 0 | 0 | 0 | 0 | 0 | 0 | 0 | 0 | 0 | 0 | 0 | 0 | 0 | 0 |
| | Complex I | nuoB | NADH-quinone oxidoreductase, subunit B | COG0377 | 3J11S@40117\|Nitrospirae | | Nide0614 | 1 | 1 | 1 | 1 | 1 | 2 | 1 | 0 | 1 | 1 | 1 | 1 | 1 | 1 | 1 | 1 | 1 |
| | Complex I | nuoA | NADH-quinone oxidoreductase, membrane subunit A | COG0838 | 3J0MS@40117\|Nitrospirae | | Nide0615 | 1 | 1 | 1 | 1 | 1 | 1 | 1 | 1 | 1 | 1 | 1 | 1 | 1 | 1 | 1 | 1 | 1 |
| | Complex I | nuoH | NADH-quinone oxidoreductase, membrane subunit H | COG1005 | 3J0E2@40117\|Nitrospirae | | Nide0608 | 1 | 1 | 1 | 1 | 1 | 1 | 1 | 1 | 1 | 1 | 1 | 1 | 1 | 1 | 1 | 1 | 1 |
| | Complex I | nuoF | NADH-quinone oxidoreductase, subunit F | COG1894 | 3J0WX@40117\|Nitrospirae | | Nide0611 | 1 | 1 | 1 | 1 | 1 | 1 | 1 | 1 | 1 | 1 | 1 | 1 | 1 | 1 | 1 | 1 | 1 |
| | Complex I | nuoE | NADH-quinone oxidoreductase, subunit E | COG1905 | 3J1BD@40117\|Nitrospirae | | Nide0612 | 1 | 1 | 1 | 1 | 1 | 1 | 1 | 1 | 1 | 1 | 1 | 1 | 1 | 1 | 1 | 1 | 1 |
| | Alternative Complex I | nuoN | NADH-quinone oxidoreductase, membrane subunit N | COG1007 | | | Nide0225 | 1 | 1 | 1 | 1 | 1 | 1 | 1 | 1 | 1 | 1 | 1 | 1 | 1 | 1 | 1 | 1 | 1 |
| | Alternative Complex I | nuoM | NADH-quinone oxidoreductase, membrane subunit M | COG1008 | | | Nide0226, Nide0227 | 2 | 2 | 2 | 2 | 2 | 2 | 2 | 2 | 2 | 2 | 2 | 2 | 2 | 2 | 2 | 2 | 2 |
| | Alternative Complex I | nuoL | NADH-quinone oxidoreductase, membrane subunit L | COG1009 | | | Nide0228 | 1 | 1 | 1 | 1 | 1 | 1 | 1 | 1 | 1 | 1 | 1 | 1 | 1 | 1 | 1 | 1 | 1 |
| | Alternative Complex I | nuoK | NADH-quinone oxidoreductase, membrane subunit K | COG0713 | | | Nide0229 | 1 | 1 | 1 | 1 | 1 | 1 | 1 | 1 | 1 | 1 | 1 | 1 | 1 | 1 | 1 | 1 | 1 |
| | Alternative Complex I | nuoJ | NADH-quinone oxidoreductase, membrane subunit J | COG0839 | | | Nide0230 | 1 | 1 | 1 | 1 | 1 | 0 | 1 | 1 | 1 | 1 | 1 | 1 | 1 | 1 | 1 | 1 | 1 |
| | Alternative Complex I | nuoI | NADH-quinone oxidoreductase, subunit I | COG1143 | | | Nide0231 | 1 | 1 | 1 | 1 | 1 | 1 | 1 | 1 | 1 | 1 | 1 | 1 | 1 | 1 | 1 | 1 | 1 |
| | Alternative Complex I | nuoG | putative NADH-quinone oxidoreductase, subunit G | COG3383 | | | Nide0232 | 1 | 1 | 1 | 1 | 1 | 1 | 1 | 1 | 1 | 1 | 1 | 1 | 1 | 1 | 1 | 1 | 1 |
| | Alternative Complex I | nuoD | NADH-quinone oxidoreductase, subunit D | COG0649, COG0852 | | | Nide0233 | 1 | 1 | 1 | 1 | 1 | 1 | 1 | 1 | 1 | 1 | 2 | 1 | 1 | 1 | 1 | 1 | 1 |
| | Alternative Complex I | nuoC | NADH-quinone oxidoreductase, subunit C | COG0852 | | | Nide0234 | 1 | 1 | 1 | 1 | 1 | 1 | 1 | 1 | 1 | 1 | 1 | 1 | 1 | 1 | 1 | 1 | 1 |
| | Alternative Complex I | nuoB | NADH-quinone oxidoreductase, subunit B | COG0377 | | | Nide0235 | 1 | 1 | 1 | 1 | 1 | 1 | 1 | 1 | 1 | 1 | 1 | 1 | 1 | 1 | 1 | 1 | 1 |
| | Alternative Complex I | nuoA | NADH-quinone oxidoreductase, membrane subunit A | COG0838 | | | Nide0236 | 1 | 1 | 1 | 1 | 1 | 1 | 1 | 1 | 1 | 1 | 1 | 1 | 1 | 1 | 1 | 1 | 1 |
| | Alternative Complex I | nuoF | NADH-quinone oxidoreductase, subunit F | COG1894 | 3J0WA@40117\|Nitrospirae | | Nide3560 | 1 | 1 | 1 | 1 | 1 | 1 | 1 | 1 | 1 | 1 | 1 | 1 | 1 | 1 | 1 | 1 | 1 |
| | Complex II | sdhA | Succinate dehydrogenase/fumarate reductase, flavoprotein subunit | COG1053 | | | Nide0310 | 1 | 2 | 1 | 1 | 1 | 0 | 1 | 1 | 1 | 0 | 1 | 1 | 1 | 1 | 1 | 1 | 1 |
| | Complex II | fer | Ferredoxin | COG1152 | 3J11C@40117\|Nitrospirae | | Nide0838 | 1 | 1 | 1 | 1 | 1 | 1 | 1 | 1 | 1 | 1 | 1 | 1 | 1 | 1 | 1 | 1 | 1 |
| | Complex II | sdhA or nadB | Succinate dehydrogenase/fumarate reductase, flavoprotein subunit or L-aspartate oxidase | COG0029 | 3J0WB@40117\|Nitrospirae | | Nide0839 | 1 | 1 | 1 | 1 | 1 | 1 | 1 | 1 | 1 | 1 | 2 | 1 | 1 | 1 | 1 | 1 | 1 |
| | Complex II | sdhB | Succinate dehydrogenase/fumarate reductase, Fe-S protein subunit | COG0479 | | | Nide2517 | 1 | 1 | 1 | 1 | 1 | 0 | 1 | 1 | 1 | 0 | 1 | 1 | 1 | 1 | 1 | 1 | 1 |
| | Complex II | sdhE | Succinate dehydrogenase, subunit C | COG2048 | | | Nide2527 | 1 | 1 | 1 | 1 | 1 | 0 | 1 | 1 | 1 | 0 | 1 | 1 | 1 | 1 | 1 | 1 | 1 |
| | Complex III | qcrA | Cytochrome *bc1* complex, iron-sulfur subunit | COG0723 | 3J0TT@40117\|Nitrospirae | | Nide0898, Nide3890 | 1 | 1 | 1 | 1 | 1 | 1 | 1 | 1 | 2 | 1 | 1 | 2 | 2 | 2 | 3 | 2 | 2 |
| | Complex III | qcrB/C | Cytochrome *bc1* complex, cytochrome b/c subunit | COG1290 | 3J0Q6@40117\|Nitrospirae | | Nide0899, Nide3889 | 1 | 1 | 1 | 1 | 1 | 1 | 1 | 1 | 3 | 1 | 1 | 2 | 2 | 2 | 3 | 2 | 2 |
| | Alternative Complex III | alt_qcrC | putative Menaquinol oxidoreductase complex ACIII, iron-sulfur cluster-binding subunit | COG3258 | | NITMAR_v1_2632 | Nide3886 | 1 | 1 | 1 | 1 | 1 | 0 | 1 | 1 | 1 | 1 | 1 | 1 | 0 | 1 | 0 | 1 | 1 |
| | Alternative Complex III | *alt_qrcA* | putative Menaquinol oxidoreductase complex ACIII, cytochrome c subunit | | | NITMAR_v1_2633 | | 0 | 0 | | | 0 | | | | | | | 1 | 0 | 1 | 0 | 1 | 0 |
| | Alternative Complex III | *alt_qrcB* | putative Menaquinol oxidoreductase complex ACIII,, molybdopterin-binding subunit | | | NITMAR_v1_2634 | | 0 | 0 | | | 0 | | | | | | | 1 | 0 | 1 | 0 | 1 | 0 |
| | Alternative Complex III | *qrcD* | putative Menaquinol oxidoreductase complex ACIII, menaquinol-binding integral membrane subunit | | | NITMAR_v1_2635 | | 0 | 0 | | | 0 | | | | | | | 1 | 0 | 1 | 0 | 1 | 0 |
| | Alternative Complex III | *actD* | putative Menaquinol oxidoreductase complex ACIII, DUF3341 subunit | | | NITMAR_v1_2636 | | 0 | 0 | | | 0 | | | | | | | 1 | 0 | 1 | 0 | 1 | 0 |
| | Alternative Complex III | *actE* | putative Menaquinol oxidoreductase complex ACIII, monohaem cytochrome c subunit | | | NITMAR_v1_2637 | | 0 | 0 | | | 0 | | | | | | | 1 | 0 | 1 | 0 | 1 | 0 |
| | Complex V | atpH | putative ATP synthase F1, delta subunit | COG0712 | 3J1BY@40117\|Nitrospirae | | Nide0369 | 1 | 1 | 1 | 1 | 1 | 1 | 1 | 1 | 1 | 1 | 2 | 1 | 1 | 1 | 1 | 1 | 1 |
| | Complex V | atpA | ATP synthase F1, alpha subunit | COG0056 | 3J0CZ@40117\|Nitrospirae | | Nide0370 | 1 | 1 | 1 | 1 | 1 | 1 | 0 | 1 | 1 | 1 | 1 | 1 | 1 | 1 | 1 | 1 | 1 |
| | Complex V | atpG | ATP synthase F1, gamma subunit | COG0224 | 3J0J8@40117\|Nitrospirae | | Nide0371 | 1 | 1 | 1 | 1 | 1 | 1 | 1 | 1 | 1 | 1 | 1 | 1 | 1 | 1 | 1 | 1 | 1 |
| | Complex V | atpD | ATP synthase F1, beta subunit | COG0055 | 3J0BN@40117\|Nitrospirae | | Nide0372 | 1 | 1 | 1 | 1 | 1 | 1 | 1 | 1 | 1 | 1 | 1 | 1 | 1 | 1 | 1 | 1 | 1 |
| | Complex V | atpC | ATP synthase F1, epsilon subunit | COG0355 | 3J0SS@40117\|Nitrospirae | | Nide0373 | 1 | 1 | 1 | 1 | 1 | 1 | 1 | 1 | 1 | 1 | 1 | 1 | 1 | 1 | 1 | 1 | 1 |
| | Complex V | atpF | ATP synthase F0, subunit B | COG0711 | 3J1AE@40117\|Nitrospirae | | Nide3732 | 1 | 1 | 1 | 1 | 1 | 1 | 1 | 1 | 1 | 1 | 1 | 1 | 1 | 1 | 1 | 1 | 1 |
| | Complex V | atpE | ATP synthase F0, subunit C | COG0636 | 3J198@40117\|Nitrospirae | | Nide3733 | 1 | 1 | 1 | 1 | 1 | 1 | 1 | 1 | 1 | 1 | 1 | 1 | 1 | 1 | 1 | 1 | 1 |
| | Complex V | atpB | ATP synthase F0, subunit A | COG0356 | 3J0RW@40117\|Nitrospirae | | Nide3734 | 1 | 1 | 1 | 1 | 1 | 1 | 1 | 1 | 1 | 1 | 1 | 1 | 1 | 1 | 1 | 1 | 1 |
| | Complex V | atpI | putative ATP synthase F0, subunit I | COG5336 | | | Nide3735 | 1 | 1 | 1 | 1 | 1 | 1 | 1 | 1 | 1 | 1 | 1 | 1 | 1 | 1 | 1 | 1 | 1 |
| | Alternative Complex IV | alt_atpA | canonical cyt. *bd* quinol oxidase (CcoNOP cyt. cbb3 terminal oxidase) | COG0056 | | | | 0 | 0 | 0 | 0 | 0 | 0 | 1 | 0 | 0 | 0 | 0 | 0 | 1 | 1 | 0 | 1 | 1 |
| | Alternative Complex IV | alt_atpG | canonical cyt. *bd* quinol oxidase (CcoNOP cyt. cbb3 terminal oxidase) | COG0224 | | | | 0 | 0 | 0 | 0 | 0 | 0 | 0 | 0 | 0 | 0 | 0 | 0 | 1 | 1 | 0 | 1 | 2 |
| | Alternative Complex IV | alt_atpD | canonical cyt. *bd* quinol oxidase (CcoNOP cyt. cbb3 terminal oxidase) | COG0055 | | | | 0 | 0 | 0 | 0 | 0 | 1 | 0 | 0 | 0 | 0 | 0 | 0 | 1 | 1 | 0 | 1 | 1 |
| | Alternative Complex IV | alt_atpF | canonical cyt. *bd* quinol oxidase (CcoNOP cyt. cbb3 terminal oxidase) | COG0711 | | | | 0 | 0 | 0 | 0 | 0 | 0 | 0 | 0 | 1 | 0 | 0 | 0 | 1 | 1 | 0 | 1 | 1 |
| | Alternative Complex IV | alt_atpE | canonical cyt. *bd* quinol oxidase (CcoNOP cyt. cbb3 terminal oxidase) | COG0636 | | | | 0 | 0 | 0 | 0 | 0 | 0 | 0 | 0 | 1 | 0 | 0 | 0 | 1 | 1 | 0 | 1 | 1 |
| | Alternative Complex IV | *alt_atpC* | canonical cyt. *bd* quinol oxidase (CcoNOP cyt. cbb3 terminal oxidase) | COG0355 | | | | 0 | 0 | 0 | 0 | 0 | 0 | 0 | 0 | 0 | 0 | 0 | 0 | 1 | 1 | 0 | 1 | 1 |
| | Alternative Complex IV | *atpQ* | canonical cyt. *bd* quinol oxidase (CcoNOP pr terminal oxidase) | | | | | 0 | | | | | | | | | | | | 1 | 1 | | 1 | 1 |
| | Alternative Complex IV | *atpR* | canonical cyt. *bd* quinol oxidase (CcoNOP cyt. cbb3 terminal oxidase) | | | | | 0 | | | | | | | | | | | | 1 | 1 | | 1 | 1 |
| | Complex IV | | putative Cytochrome bd oxidase subunit I (cyt. bd-like ) | | | | | 4 | 4 | 4 | 4 | 4 | 4 | 4 | 4 | 4 | 4 | 4 | 7 | 5 | 7 | 4 | 7 | 6 |
| | Complex IV | *cydA* | Cytochrome bd-type quinol oxidase subunit I (cyt. bd-like ) | COG1271 | 3J0HH@40117\|Nitrospirae | Terminal oxidases | Nide2610 | 0 | 0 | 0 | 0 | 0 | 0 | 0 | 0 | 0 | 0 | 0 | 2 | 2 | 2 | 1 | 2 | 1 |
| | Complex IV | *cydB* | Cytochrome BD ubiquinol oxidase subunit II (cyt. bd-like ) | COG1294 | 3J0M4@40117\|Nitrospirae | Terminal oxidases | Nide2609 | 0 | 0 | 0 | 0 | 0 | 0 | 0 | 0 | 0 | 0 | 0 | 2 | 2 | 2 | 1 | 2 | 1 |
